# Performance of Rilpivirine-Based Hydrophobic Tags and PROTACs Directed Against HIV-1 Reverse Transcriptase

**DOI:** 10.64898/2026.09.24.753933

**Authors:** Ferdinand K. Amanor, Garrett D. Clements, Rabia Khurshid, Anna F. Howard, Diana Soto Martinez, Courtney Barkley, Zhengrong Yang, Kabita Gyawali, John C. Kappes, Robert C. Reynolds, Stephan C. Schürer, Timothy S. Snowden, Christina Ochsenbauer

**Affiliations:** Department of Medicine, University of Alabama at Birmingham, AL, USA; Department of Chemistry and Biochemistry, The University of Alabama Tuscaloosa, AL, USA; Department of Molecular and Cellular Pharmacology, Miller School of Medicine, University of Miami, Miami FL, USA; Department of Biochemistry & Molecular Genetics, University of Alabama at Birmingham, AL, USA; Sylvester Comprehensive Cancer Center, University of Miami, Miami FL, USA; Frost Institute for Data Science & Computing, University of Miami, Miami FL, USA

**Keywords:** HIV-1, PROTACs, Targeted Protein Degradation, Hydrophobic Tag, Reverse Transcriptase

## Abstract

We aimed to repurpose the non-nucleoside reverse transcriptase (RT) inhibitor (NNRTI) rilpivirine (RPV) as a targeted protein degrader (TPD) of HIV-1 RT. Structure-guided modeling identified a TPD bifunctional linker exit trajectory from RPV that threads the NNRTI entrance channel. Ten RPV analogs (RPV’) with varied warhead-linker attachments retained RT inhibition and antiviral activity, guiding selection of an amide-linked connector for degrader construction. We synthesized Proteolysis Targeting Chimeras (PROTACs) designed to recruit CUL4^CRBN^ and CUL2^VHL^, alongside adamantyl acetic acid-based Hydrophobic Tags (HyTs). TPDs were evaluated for RT inhibition, virus inhibition, biophysical target engagement, and proteasome-dependent degradation. Some PROTACs showed minimal antiviral activity or limited solubility for our assays, whereas the HyT with a tetraethylene glycol (PEG4) linker (PEG4-Ad) exhibited low-nanomolar potency to suppress HIV-1 replication and acceptable solubility. PEG4-Ad reduced RT levels in a proteasome-dependent manner without cytotoxicity but did not exhibit superior potency against RPV resistance mutations. MD simulations suggested that PEG4 linkers maximize episodic exposure of the hydrophobic tag, consistent with PEG4-Ad efficacy. These data highlight the promise—and constraints—of antiviral degraders, indicating that linker-controlled hydrophobic tag exposure and subcellular target accessibility may be critical design parameters for prospective therapeutics.

## 1. Introduction

HIV-1 reverse transcriptase (RT) is a heterodimeric enzyme (p66/p51) essential for converting the viral RNA genome into double-stranded DNA, making it a primary target in antiretroviral therapy (ART). RT inhibitors are a cornerstone of clinical ART regimens [1]. These compounds bind to RT and prevent the synthesis of the proviral DNA needed for productive infection. There are currently two classes of FDA approved RT inhibitors— nucleoside inhibitors (NRTIs) and non-nucleoside inhibitors (NNRTIs). NRTIs were the first discovered ART and continue to be important drugs for first line HIV ART regimens and pre-exposure prophylaxis (PrEP) [2]. These compounds competitively inhibit DNA synthesis. NNRTIs are allosteric inhibitors of RT that bind in a hydrophobic cavity located ∼10 Å from the polymerase active site at the junction of the p66 palm and thumb subdomains [3]. This pocket is not present in the apoenzyme but is induced upon inhibitor binding through conformational rearrangement of key residues – including Y181, Y188, K103, and V106 – forming a flexible hydrophobic cleft [4]. NNRTIs occupy this site and reposition the primer grip, locking RT in a catalytically inactive conformation [3–5]. Despite their clinical utility, RT inhibitors have the highest frequency of resistance mutations of all HIV drugs due to low genetic barriers for selection of resistant viral variants. Drug resistance presents a significant concern for effective HIV treatment, especially regarding mutations such as M184V and K65R, which confer cross-resistance to many NRTIs, as well as K103N and Y181V/C [1, 4, 6, 7]. Second generation NNRTIs, such as rilpivirine, have increased efficacy against many of these mutations due to their structural flexibility [3, 8]; however, additional mutations are already contributing to the resistance profiles of these compounds, thus highlighting the need for novel RT inhibitors [1, 9].

Targeted protein degradation harnesses the ubiquitin–proteasome system for 26S proteasomal elimination of disease-associated proteins and has matured into a clinically validated therapeutic strategy [10]. Proteolysis Targeting Chimeras (PROTACs) constitute the most prominent class of targeted protein degraders (TPDs) [11]. PROTACs are heterobifunctional small molecules that link a ligand for a protein of interest (POI) to a recruiter of an E3 ubiquitin ligase (E3). Their mechanism of action relies on stabilizing ternary POI–E3 ligase complexes, which facilitate proximity-induced polyubiquitination of the target and its subsequent recognition and degradation by the proteasome [12–14] **(Figure 1A)**. In recent years, TPD approaches have expanded to include other mechanisms of degradation outside the PROTAC approach [15, 16]. One such alternative is hydrophobic tag degraders (HyT degraders), which feature a hydrophobic group linked to the POI ligand [17]. When bound, the hydrophobic moiety sits on or near the surface of the protein – mimicking a partially unfolded or unstable protein, which is then recognized by the cell’s protein quality control machinery (e.g., including heat shock protein 70 (HSP70) and CHIP), leading to target degradation [17, 18] **(Figure 1B)**. TPDs have favorable characteristics compared to the parent occupancy-based inhibitors due to the TPD’s catalytic mechanism of action, which requires sub-stoichiometric doses for efficacy and generally features a longer duration of activity [19]. In addition, several TPDs show enhanced efficacy against treatment-resistant mutants compared to drug/inhibitor alone [20–23]. Targeted protein degraders have been most intensively investigated in oncology, where multiple PROTACs have advanced into clinical evaluation [12, 24]. By contrast, the application of targeted protein degradation to infectious diseases has been comparatively limited. One of the earliest published antiviral demonstrations, reported in 2019, described a telaprevir-based PROTAC capable of inducing degradation of the hepatitis C virus NS3/4A protease [25]. Since then, TPDs have been reported for viral targets such as influenza A neuraminidase, SARS-CoV-2 main protease, and the HIV-1 Nef and Vif proteins and host factors such as cyclophilin A [26–30].

**Figure 1.**
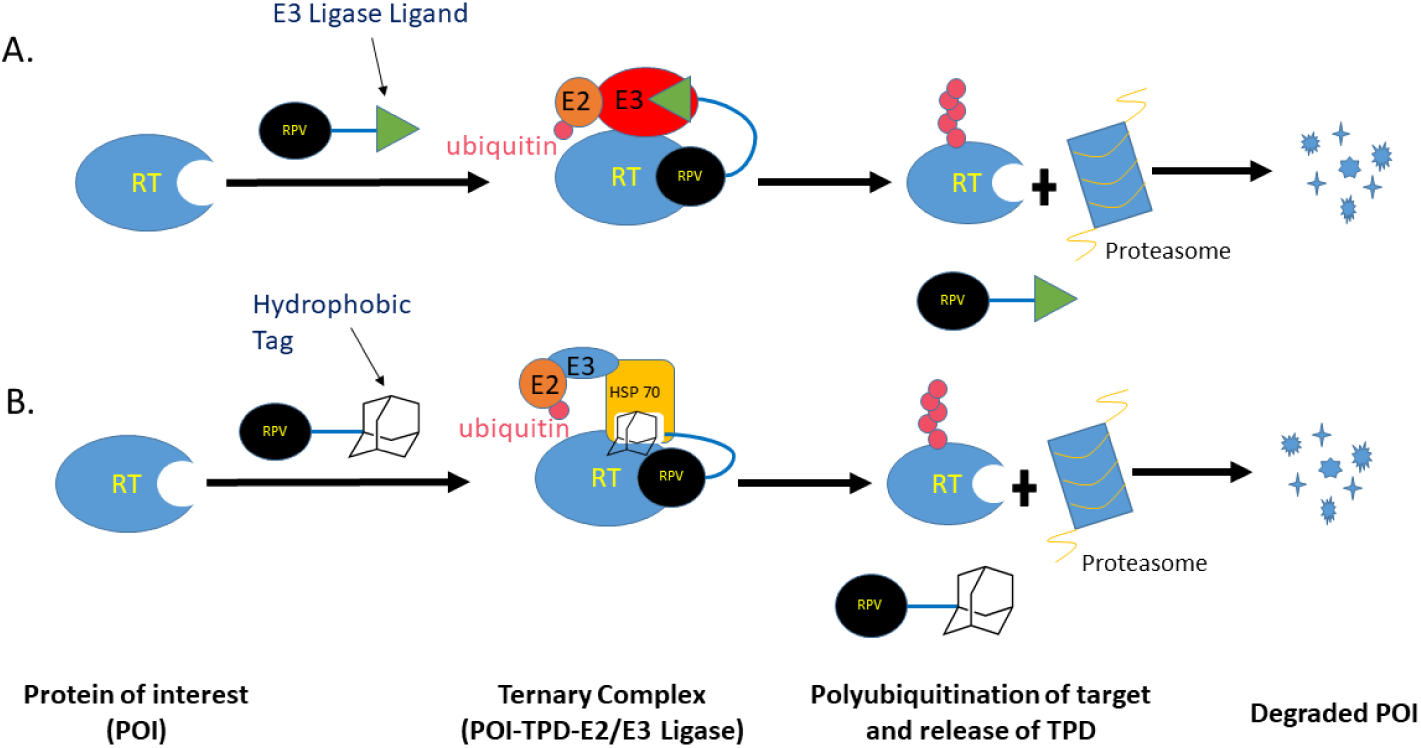
Schematic representation of canonical (A) PROTAC and (B) HyT mechanisms of targeted protein degradation. HSP70 is shown as one possible recognition factor for the exposed hydrophobic tag; the actual degradation machinery may involve alternative chaperones and ubiquitin–proteasome system components.

Herein, we describe our approach to rilpivirine (RPV, **6**, **Figure 4**)-based PROTACs and HyT degraders targeting HIV-1 reverse transcriptase. We utilized the high-resolution crystal structure of RT bound to RPV (PDB ID: 2ZD1) to inform our design for linker attachment sites and for E3/HSP recruitment. Eight synthetic compounds retained sub-micromolar IC₅₀ values in infection assays. The HyT designs generally displayed the highest antiviral activity in this series, and proteasome-dependent degradation of RT was observed with one compound. However, its relative antiviral activity against RT variants containing resistance mutations to rilpivirine (RPV) was comparable to the parent RPV.

## 2. Materials and Methods

### A. Combined Rational and Computational Design of PROTACs and HyTs

Detailed computational protocols, parameter settings, and analysis procedures are provided in the Supporting Information in addition to the most relevant method overview provided here.

#### Linker Enumeration and Constrained Docking

A focused virtual library of 232 rationally designed rilpivirine-based linker derivatives spanning multiple connector chemotypes and polyethylene glycol (PEG) lengths (1-6 units) was enumerated using ChemAxon Marvin Sketch. Linker variants were evaluated using constrained molecular docking in the Schrödinger Small-Molecule Drug Discovery software suite [31], where the warhead orientation was fixed while allowing conformational sampling of the appended linker. Docking scores and physicochemical property predictions generated with Schrödinger QikProp were used to prioritize linker chemotypes and lengths compatible with RT binding and spanning the p66—p51 inter-subunit channel. 12 of 35 prioritized RPV’ linker candidates were selected for chemical synthesis (Figure 3, first column). More details are found in **Supporting Information (2.2, 2.3, 2.4)**.

#### Modeling of PROTAC and HyT Constructs

Prioritized linker designs were combined either with E3 ligase recruiters (pomalidomide or VH032) to model 120 bifunctional TPDs (Figure 3, second column), or with 1-adamantane acetic acid to model HyT constructs Figure 3, fourth column). Predicted physicochemical properties of full PROTACs were benchmarked against empirically derived property distributions from PROTAC-DB [32] to assess optimal medicinal chemistry properties. More details are found in **Supporting Information (2.5, 2.8).**

#### PROTAC Ternary Complex Modeling and Molecular Dynamics (MD) Simulations

For PROTACs, ternary complexes comprising RT, the degrader, and the E3 ligase cereblon (CRBN; PDB ID: 4CI3) or VHL (PDB ID: 4W9H) were modeled using protein– protein docking in HADDOCK 2.4 [33] and P4ward [34]. Selected RT–PROTAC–CRBN complexes were refined and evaluated using explicit-solvent molecular dynamics simulations performed in Desmond with the OPLS4 force field [35, 36]. Production simulations of ternary complexes were 150 – 250 ns (analyzed over a uniform 50 – 150 ns window. Binary RT-CRBN simulations were performed as controls. A detailed description is found in **Supporting Information (2.6, 2.7, 2.9)**.

#### Hydrophobic Tag Modeling and Molecular Dynamics (MD) Simulations

Methods similar to those above were applied to RT-HyT complexes, which were refined and evaluated using explicit-solvent molecular dynamics simulations performed in Desmond with the OPLS4 force field [35, 36]. HyT constructs were simulated for 50 ns (analyzed over 20 – 50 ns). A detailed description is found in **Supporting Information (2.8, 2.9)**.

### B. Synthesis

#### Preparative routes and purification

All bio-assayed compounds were purified by flash and/or preparative thin layer chromatography and demonstrated >95% purity by NMR spectroscopy. Full preparative routes, detailed experimental procedures, and characterization data (^1^H and ^13^C NMR spectroscopy and high-resolution mass spectrometry (HRMS)) of compounds in Figures 2, 6, and 9, and all associated precursors are reported in Supporting Information Section 1 and the Supplementary NMR Spectra file.

**Figure 2.**
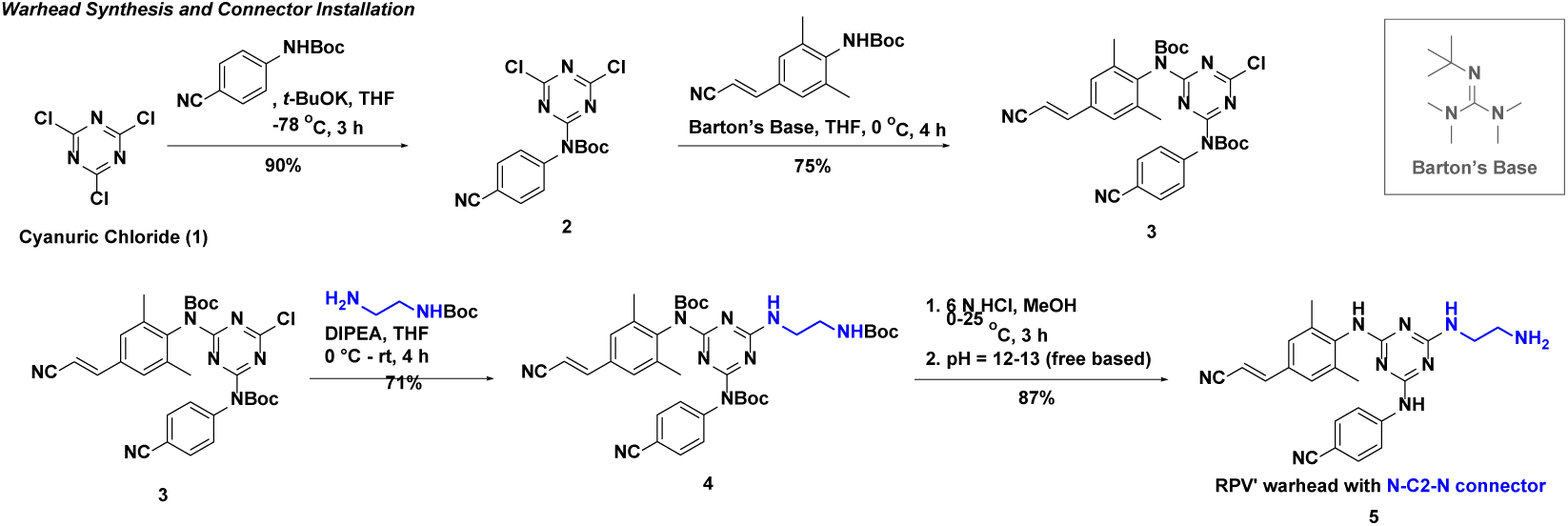
Synthetic route to prepare the warhead-connector segment of 10, 14, and all TPDs in Figure 9.

**Figure 3.**
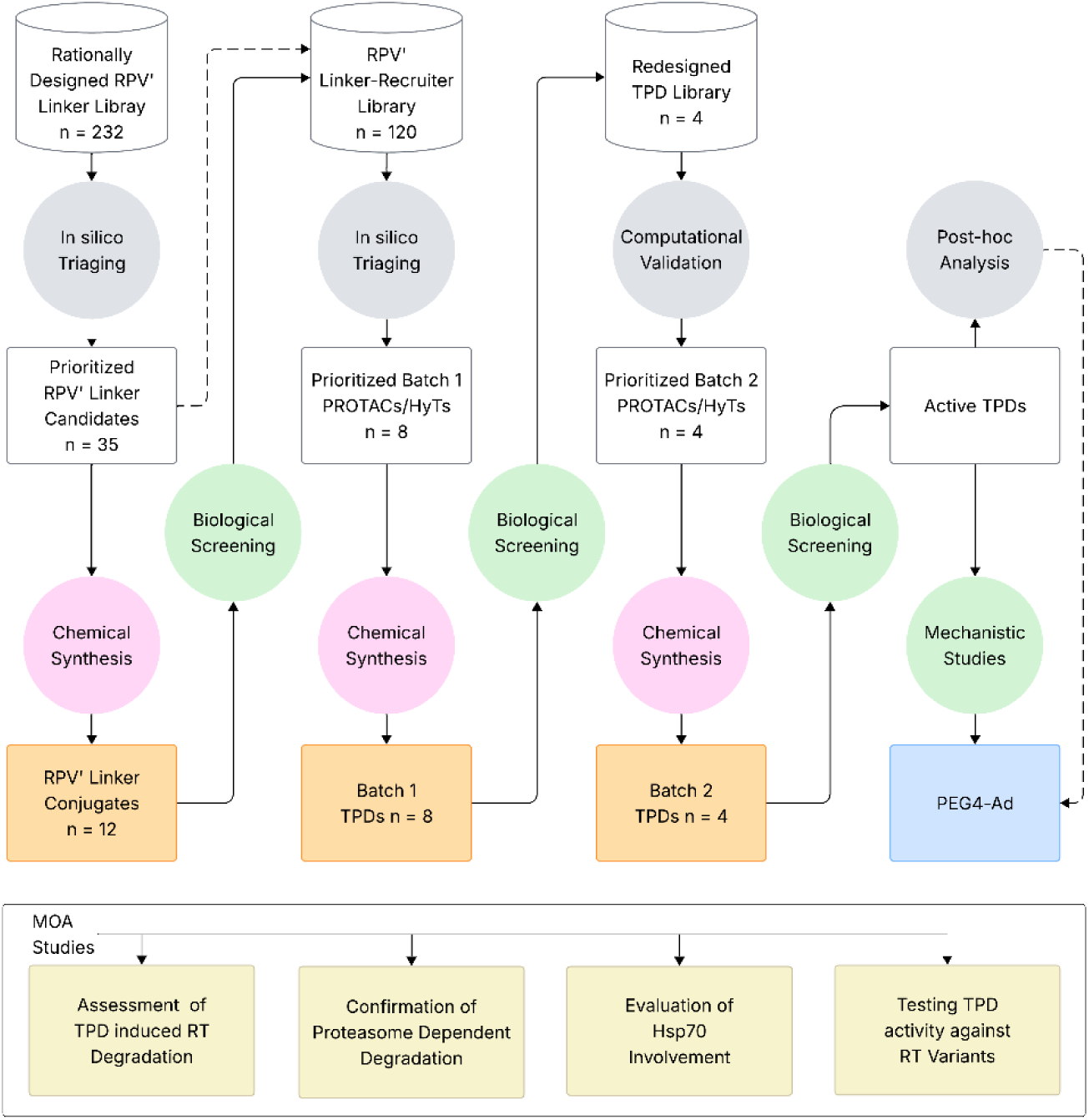
Workflow overview for the computationally guided design, iterative optimization, and mechanistic evaluation of RPV′-based reverse transcriptase degraders. Starting from a rationally designed library of 232 RPV′ linkers, we applied a computational triaging pipeline comprising molecular docking, molecular dynamics, and ADMET prediction to identify 35 prioritized candidates. Twelve candidates were synthesized as RPV′ linker conjugates and evaluated by biological screening. The prioritized linkers were also combinatorially coupled to CRBN-and VHL-recruiting ligands through distinct connector chemistries to generate a virtual RPV′ linker–recruiter library of PROTAC and hydrophobic-tag designs (n = 120). Computational triaging prioritized eight Batch 1 candidates that were synthesized and biologically screened. Screening results informed the design of a refined TPD library (n = 4), which underwent computational validation before all four Batch 2 candidates were synthesized and tested. Active compounds identified across the screening cycles were advanced to post-hoc computational analysis and biological characterization, leading to the selection of PEG4-Ad (26, Figure 9) for mechanistic studies. Mode-of-action studies independently assessed TPD-induced RT degradation, proteasome dependence, Hsp70 involvement, and activity against RT variants carrying rilpivirine-resistance mutations. Solid arrows indicate progression through the design–model–make–test workflow, whereas dashed arrows indicate transfer of computationally prioritized designs or feedback between computational analysis and experimental findings.

**Figure 4.**
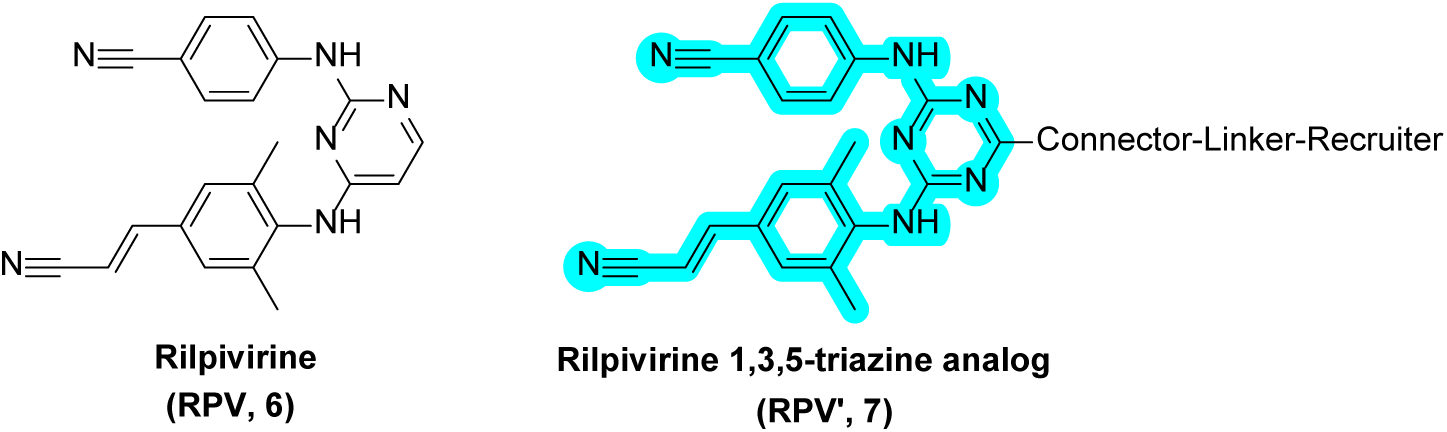
Structural comparison of the NNRTI rilpivirine (6) and the highlighted 1,3,5-triazine– based rilpivirine analog (RPV′, 7) employed as the warhead scaffold in this work. In RPV′, the triazine core replaces the native pyrimidine of rilpivirine, and schematic connector, linker, and recruiter segments are shown to indicate the segments of TPD elaboration.

To enable modular linker installation to the warhead, a 1,3,5-triazine-based rilpivirine analog (RPV’, **7**, **Figure 4**) was employed as the warhead scaffold. The 4-((2-aminoethyl)amino)-1,3,5-triazinyl RPV′-connector (**5**) ultimately selected was generated on multigram scale through successive nucleophilic aromatic mono-substitution reactions starting from cyanuric chloride (**1**) (**Figure 2**). Sequential installation of *tert*-butyl (4-cyanophenyl) carbamate to generate **2**, and *tert*-butyl (*E*)-(4-(2-cyanovinyl)-2,6-dimethylphenyl) carbamate to afford the trisubstituted triazine core bearing a remaining chloride (**3**), was conducted at low temperature under basic conditions. Subsequent chloride substitution in **3** was achieved by nucleophilic displacement by the appropriate oxygen-or amine-based connector of varied length (e.g., *tert*-butyl (2-aminoethyl)carbamate to create **4**), using conditions optimized for the specific connector employed (see **Supporting Information Section 1**). Global Boc removal using HCl in methanol provided a common connector-terminus synthetic handle, enabling installation of diverse linkers and subsequent elaboration to full TPDs, as detailed in the **Supporting Information Section 1**.

The VHL recruiting ligands VH032 and Me-VH032 in PROTACs **23**-**25** and **30**, respectively, were prepared as reported previously[37].

### C. Bioassays

#### Reverse Transcriptase Inhibition Assay

Assessment of inhibition of RT enzymatic activity in vitro was conducted using the Roche reverse transcriptase colorimetric assay following manufacturer’s instructions [38]. 10 mM stocks of RPV and RPV’-TPDs in DMSO were diluted in PBS pH 7.4 without magnesium or calcium (Gibco) so that the final concentrations in the 60 µl reactions were 40.0, 20.0,10.0,1.0, 0.1, and 0.01 µM respectively. Reactions were set up using 0.5 ng of purified RT supplied in the kit, following in-house optimization. Reaction mixtures consisting of the supplied template, RT, and inhibitors were incubated at 37°C for 1 hour. After incubation, the reaction mixtures were transferred to the streptavidin coated microplate modules, covered with self-adhesive foil, and incubated for 1 hour at 37°C. Solutions were then removed, and the modules were washed 5X with 250 µl of washing buffer. 200 µl of the anti-DIG-POD working solution was then added and modules were covered and incubated at 37°C for 1 hour. Wash steps were then repeated and 200 µl of the ABTS substrate solution was added. Modules were then incubated in the dark at room temperature for 15 minutes. Absorbance was read using a microplate reader at 405 nm and 495 nm (reference wavelength). Percent activity was calculated by multiplying the ratio of our inhibitor-treated conditions to the untreated control by 100 after background correction. Percent inhibition was calculated by subtracting the percent activity from 100. Dose response curves were generated by performing a nonlinear fit of the percent inhibition data after log transformation in GraphPad Prism software (Dotmatics).

#### Proviral plasmid DNA

We utilized our previously described HIV-1 strain pNL4-3-based proviral plasmid that encodes the NL4-3 *env* gene and the Renilla luciferase reporter gene, placed upstream of a modified IRES element driving Nef expression pNL-LucR.6ATRi/K5300 [39]. In addition to this infectious molecular clone (IMC) of HIV-1, several RT-mutant derivatives were generated as described below.

#### Generation of 293T cell-derived HIV-1 virus stocks

Virus stocks were produced by transfecting 293T cells (300,000 cells/6-well) with 2 µg of proviral plasmid DNA using FuGENE^®^HD Transfection Reagent (Promega) following manufacturer instructions. DMEM media supplemented with 10% FBS, 25 mM HEPES, L-glutamine (2 mM), streptomycin (100 µg/ml), and penicillin (100 U/ml) (DMEM10++) was changed 24 hours post transfection. 72 hours post transfection, viral supernatants were harvested and clarified at 300 x g for 10 minutes before freezing at –80°C. Virus stocks were titered on TZMbl cells by counting beta-galactosidase-stained cells as previously[40].

#### TZMbl single round infection assay

TZMbl assays were conducted essentially as previously described [41]. Briefly, RPV and RPV’-TPDs were 3-fold serially diluted eight time, encompassing a range of 0.0125-225 nM, depending on starting concentration, in quadruplicates in 96-well plates in 100 µl. 1 x 10^4^ freshly trypsinized TZMbl cells were then added into each well in 50 µl DMEM10++ containing 40 ug/ml DEAE-dextran (final concentration 10 µg/ml). Cells were then infected with 293T-transfection derived HIV-1 reporter virus, NL-LucR.6ATRi (Lab reference K5300) [39] at a TZMbl MOI of 2.5 in 50 µl. Cells were then incubated at 37°C in a humidified atmosphere of 5% CO_2_ for 48 hours after which cells were lysed using 50 µl of 5X Renilla Luciferase Assay Lysis Buffer (Promega). 96-well plates were kept frozen at –80°C until the Renilla luciferase signal of the lysed samples were analyzed using the Renilla Luciferase Assay System (Promega) and a GLOMAX Discover luminometer (Promega) equipped with injectors, essentially following manufacturer’s instructions.

#### HIV-1 Replication assay

To assess inhibition of replication, we utilized a CD4 T cell line derived from C8166 [42] modified to express CCR5 [43], which we further engineered with an LTR-driven, Tat –dependent reporter cassette for mCherry and secreted nanoluciferase (snLuc), named C8166-R5.JR.3g_LTR.mCherry.sNLuc /D1473, (to be described elsewhere; *Jones et al*., *in preparation*). 10 mM stocks of RPV and RPV’-HyTs were diluted to concentrations of 225, 25, and 2.75 nM in RPMI 1640 supplemented with 10% FBS, 25 mM HEPES, L-glutamine (2 mM), streptomycin (100 µg/ml), and penicillin (100 U/ml). For addition of compounds at t_0_, time of infection, per condition, 4.5 x 10^5^ C8166-R5.JR.3g_LTR.mCherry.sNLuc/D1473 cells were centrifuged at 300 x g for 5 min and resuspended in 1.0 ml of RPMI 1640 medium containing the appropriate drug dilution. Cells were then transferred to a 2.0 ml Sarstedt screw-cap tube, inoculated with 860 TZMbl IU of NL-LucR.6ATRi reporter virus, and placed on a vertical rotator for 2 hours at 37°C to facilitate infection. Cells were then transferred to 15 ml conical tubes, washed with 10 ml of cold PBS, and centrifuged at 300 x g for 5 min. Supernatants were discarded, and cell pellets were resuspended in 9.0 ml of RPMI medium containing the appropriate concentration of RPV or RPV’-HyTs. Cells were seeded into six 48-well plates in a final volume of 500 µl and 25,000 cells/well in triplicate and incubated at 37°C in a humidified atmosphere of 5% CO_2_. In cases when inhibitors were added 18 hrs post infection at a final concentration of 225 nM, 0.56 ul of a 200 µM stock concentration was added directly into each well containing infected cells and mixed by pipetting. An additional 500 µl of medium was added to cultures two days post infection. Four days post infection cultures were expanded into 24 well plates in a final volume of 1.2 ml. RPV, RPV’-HyTs, and the HIV protease inhibitor indinavir (IDV) were replenished on days two and four post infection. 50 µl of culture supernatants were sampled every 24 hours for 6-9 days for snLuc using the Nano-Glo^®^Luciferase Assay System (Promega) on the GLOMAX Discover luminometer.

#### Thermal shift assay

Thermal unfolding profiles of purified RT protein with or without compounds were obtained using the Prometheus NT.48 NanoDSF instrument, which has 48 capillary chambers (NanoTemper Technologies, LLC,San Francisco, CA). Samples were excited at 290 nm, and their emissions were concurrently measured at 330 and 350 nm. Protein or compound aggregation was monitored using the integrated back-reflection optics. All samples were heated from 15 to 95°C at a constant rate of 2°C/min. A stock solution of 3.34 mg/ml (28.55 µM) purified heterodimeric RT was generously provided by the lab of Dr. Stefan Sarafianos [44] in buffer comprising 50 mM TRIS, 25 mM NaCl, 1 mM EDTA, 1 mM DTT, and 50% glycerol. Initial assessment of the protein stability was performed using a serial dilution of the stock solution into the assay buffer (20 mM HEPES, pH 7.6, 150 mM NaCl, 0.5 mM TCEP) to final concentrations of 5.7 to 0.36 µM. It was determined that 1 µM was the optimal concentration for the assay. For protein-compound mixtures, 10 mM compound stocks were diluted in the assay buffer to yield compound solutions that contained 2% DMSO and 2x final compound concentration. The protein working stock was also diluted in the assay buffer to 2x the final protein concentration. The protein and compound were then mixed 1:1 (v/v) to make the final samples.

#### Sample preparation for microscale thermophoresis

The Monolith X instrument (NanoTemper Technologies, LLC, San Francisco, CA) was used to perform the Microscale thermophoresis (MST) experiment. To ensure sufficient dilution of TRIS, EDTA, and DTT to levels suitable for MST experiments, RT stocks described above were buffer exchanged into the MST assay buffer (PBS) by mixing 40 µl of the stock protein with 450 µl of PBS without calcium or magnesium and concentrated in a 0.5-ml spin concentrator with a MWCO of 10 kDa. This dilution-concentration step was repeated 6X, resulting in a protein concentration of 6 µM. The protein was labeled using the His-Tag Labeling Kit RED-tris-NTA 2nd Generation (NanoTemper Technologies). The dissociation constant of the His-Tag binding to the dye was 6 nM. Under these conditions, 100% of the dye would be bound to the protein (100 nM RT and a dye concentration of 50 nM). Labeled RT protein was incubated with serial dilutions of PEG4-Ad (26) ranging from 6 nM to 200 µM. The MST measurements were performed in premium capillaries with 40% LED/excitation power. All data analyses were done using Monolith proprietary software.

#### Cloning of IMCs encoding NNRTI resistant RT

RT resistance mutations E138K and Y181V were introduced into pNL-LucR.6ATRi/K5300 by PCR using mutagenic overlapping primers (E138K fwd primer: CCATACCTAGTATAAACAATAAGACACCAGGG; E138K rev primer: TTATTCTTGTCCCGAACCTTTCCTAAAACGATATTCGA; Y181V fwd primer: TCCAGACATAGTCATC GTTCAATACATGG ATGA; Y181V rev primer: same as E138K rev primer). The remainder of the IMC was PCR amplified in two halves segmented at the ampicillin resistance gene (fwd primer: ATTTTGCTATAAGCTAGCCACCATGGCTTCC, rev primer: TTCTGACAACGATCGGAGGACCGAAGGAGCTAACCGCT, and fwd primer: CGATCGTTGTCAGAAGTAAGTTGGCCGCAGTGTT, rev primer (E138K): GTCCTTCATATGACGTAAATGGTATGGATCATATTTGT, rev primer (Y181V): ATGACTATGTCTGGATTTTGTTTTCTAAAA). PCR fragments were resolved and isolated by agarose gel-electrophoresis and purified using the QIAquick^®^ Gel Extraction Kit (QIAGEN). A three-piece infusion ligation was performed using In-Fusion® Cloning Master Mix (Takara Bio) followed by transformation into Stellar™ competent cells. The resulting plasmids named pNL-LucR.6ATRi-RT(E138K)/ K5814 and pNL-LucR.6ATRi-RT(I181V)/K6332, respectively, were sequenced to confirm the presence of each mutation.

The K103N/Y181C mutations were introduced separately by PCR (K103N fwd primer: ATCCTGCAGGGTTAAAACAGAATAAATCAGTA; rev primer: ATGACTATGTCTGGATTTTGTTTTCTAAAA; Y181C fwd primer: TCCAGACATAGTCATCTGTCAATACATGGATGA; rev primer: AGCTTATAGCAAAATCCTTTCCAAGCCCTGTCTTATT). The 3’ end of the fragment encoding the K103N mutation was complimentary to the 5’ end of the fragment encoding the Y181C fragment. To join these two fragments, a fusion PCR reaction was performed using the K103N fwd primer and the Y181C rev primer. The remainder of the IMC was amplified in two halves segmented at the ampicillin resistance gene (fwd primer: CGATCGTTGTCAGAAGTAAGTTGGCCGCAGTGTT; rev primer: GTTTTAACCCTGCAGGATGTGGTATTCCTAATTGAAC; and fwd primer: ATTTTGCTATAAGCTAGCCACCATGGCTTCC rev primer: TTCTGACAACGATCGGAGGACCGAAGGAGCTAACCGCT). PCR fragments were purified as described above, followed by three-piece infusion ligation and transformation. The resulting plasmid, pNL-LucR.6ATRi-RT(K103N/Y181C)/K5816 was sequenced to confirm the presence of both mutations.

#### Cloning of RT-HiBiT expression vector

The following expression cassette was constructed: p51-P2A-p66-linker-HiBiT. The nucleotide sequences comprising p51 and p66 of NL4-3 RT were amplified by PCR with overlapping primers from pNL-LucR.6ATRi/K5300. The P2A peptide sequence [45] was introduced via the reverse primer during amplification of p51 (fwd primer: ATTCGGATCCGCCACCATGCCCATTAGTCCTATTGAGACTGT; rev primer: CACGTCGCCGGCCTGCTTCAGCAGGGAGAAGTTGGTGGCCAAAGTTTCTGCTCCT AT). The (GGSGG)_2_ linker and HiBiT peptide sequences [46] were appended to the p66 subunit by the reverse PCR primer (fwd primer: GCTGAAGCAGGCCGGCGAA CGTGGAGGAGAACCCCGGCCCCATTAGTCCTATTGAGACTGT; rev primer: CTAGCTAATCTTCTTGAACAGCCGCCAGCCGCTCACGCCCCTGATCCGCCCCCGC CTGAGCCTCCTAGTACTTTCCTGATTCCAG).

The pTRE3g-CH505.w53.e16.D8.gp120-IRS6A.Puro-T2a-GFP (K5029) lentiviral vector (LVV) was amplified in two halves segmented at the ampicillin resistance gene with overlapping primers (fwd primer 1: TTCTGACAACGATCGGAGGACCGAAGGA GCTAACCGCTT; rev primer1: TGGGCATGGTGGCGGATCCGAATTCAAGTAT AAGACAAAAGT; fwd primer 2: TGTTCAAGAAGATTAGCTCGAGTAATACG; rev primer 2: CGATCGTTGTCAGAAGTAAGTTGGCCGCAGTGTT). PCR fragments were purified as described above. A four-piece infusion ligation reaction was performed. The resulting plasmid, named LVV-LTR.TRE3g-p51-P2A-p66.HiBiT-T2A-EGFP-LTR (K5891) was sequenced confirmed.

We then amplified the p51-P2A-P66-HiBiT segment of K5891 (fwd primer: CAGAAGACAGTGGCAACTTGAATTCGGATCCGCCACC, rev primer: CTCGGTATCATTATTTTAGCTAATCTTCTTGAACA) for insertion into Lenti-TRE3g-Rev.AD8+FLE.2-GGHHHHHH-IRS6A.Puro-T2a-GFP (K5626) [47] with overlapping primers (fwd primer 3: AAGAAGATTAGCTAAAATAAT GATACCGAGACCTTT, rev primer: same as “rev primer 2”, fwd primer 4: same as “fwd primer 2”, rev primer: TGCCACTGTCTTCTGCTCTTTCTATTAGTCTAT. The first 1459 bp of the AD8+FL E.2 envelope were excluded during PCR to retain only sequence that encodes the second exon of Rev and the Rev Response Element. Amplicons were then treated with DpnI for 1 hour at 37°C to digest any methylated plasmid followed by DNA purification as described above. We then conducted a three-piece infusion ligation reaction using In-Fusion^®^ Cloning Master Mix (Takara Bio) followed by transformation into Stellar™ competent cells. The resulting plasmid, named LVV-TRE3g.Rev.RTp51.P2A.RTp66.HiBiT.En-IRES.puro.T2A.EGFP (K5932) was sequence confirmed.

#### Transductions with LVV RT-HiBiT

HEK 293T stably expressing the reverse tetracycline controlled trans activator (D1317) were transfected with 2 µg of K5932, 1.5 µg of packaging plasmid, pΔ8.2 (K4), and 1 µg of Vesicular Stomatitis Virus Glycoprotein (K1746) using FuGENE^®^HD Transfection Reagent (Promega) and 1 µg/ml doxycycline. 72 hours post transfection, supernatants were harvested, and particles were pelleted by ultracentrifugation at 17000 x g for 2 hours at 4°C. Pellets were resuspended in RPMI 1640 supplemented with 1% FBS, 25 mM HEPES, L-glutamine (2 mM), streptomycin (100 µg/ml), and penicillin (100 U/ml). 5 x 10^5^ HEK293 cells (D1041) were transduced with the packaged vector by tumbling overnight. Following transduction, cells were seeded into T-25 flasks with pre-warmed F12/DMEM supplemented with 10% FBS, 25 mM HEPES, L-glutamine (2 mM), streptomycin (100 µg/ml), and penicillin (100 U/ml). 48 hours after seeding, cells were trypsinized and seeded into 6-well plates at a density of 3 x 10^5^ cells/ well. Cells were then induced with 1 µg/ml of doxycycline and selected for puromycin resistance (1 µg/ml) for 72 hours. Cultures were then expanded into a T-75 flask and FACs sorted for the GFP^+^ population using a FACs Melody cytometer.

#### RT-HiBiT degradation assay

HEK293 cells stably transduced with the RT-HiBiT LVV construct (D1786) were seeded at 7×10^4^ cells/per well into 48-well plates in culture medium (F12/DMEM). The following day, the culture medium was replaced with fresh medium supplemented with 10 ng/ml of doxycycline to induce p51-p66-HiBiT expression. 24 hours later, the medium was replaced with fresh medium containing the indicated amounts of tested inhibitors and 10 ng/ml of doxycycline. 24 hours post drug treatment, medium was removed from all wells and cells were detached from the plate using 500 µl cold PBS and collected in a 1.5 ml Eppendorf tubes. 100 µl of cell suspension was then transferred to a white luminometer plate and the Nano-Glo HiBiT Lytic Detection System (Promega) was utilized to quantify the HiBiT signal following the manufacturer’s protocol.

#### Western blot

The remaining 400 µl of the cell suspensions from our degradation assay were centrifuged at 1200 rpm for 5 min. Cell pellets were resuspended in 100 µl of cold PBS. 2X Laemmli sample buffer was added, and samples were briefly vortexed. Samples were then placed in an OMNI Bead Ruptor 12 for 30 seconds prior to heating for 5 min at 95°C. SDS-PAGE was then conducted by loading equal volumes of each sample into a 4-20% Mini-PROTEAN^®^TGX^®^ precast gel (BIORAD). Following SDS-PAGE, samples were transferred unto a nitrocellulose membrane and blocked in 5% dry milk in PBS-T for 1 hour prior to an overnight incubation with anti-HIV immune globulin (HIV-IG) (1:100) on a shaker at 4°C. The following day, membranes were washed 3X in PBS-T and incubated with the goat anti-human HRP-conjugated secondary antibody (1:1000, Southern Biotech cat# 2040-05) for 1 hour at room temperature. Protein signals were produced using Clarity Western ECL Substrate (BioRad) and visualized using the Analytikjena VisionWorks^®^ Capture and Analysis Software. Protein signals were semi quantified using UN-SCAN-IT™ Graph Digitizing Software. HiBiT-tagged p66 was visualized using the Nano-Glo^®^ HiBiT Blotting System (Promega) on a replica membrane according to the manufacturer’s protocol.

### D. Comparative Kinetic Solubility Determinations

Intrinsic kinetic solubility measurements were conducted on the PEG-4 series compounds **21**, **24**, and **26** using a UV–visible spectroscopy–based method adapted from established UV–vis solubility methodologies employed in early drug discovery and recently applied to PROTACs [48, 49].

#### Calibration Curve Preparation

Calibration standards were prepared in 2-dram glass vials by sequential addition of phosphate buffered saline (PBS; 10 mM phosphate, 150 mM NaCl, pH 7.4), dimethyl sulfoxide (DMSO; 1% v/v final), and aliquots of compound stock solutions prepared in DMSO. Samples were prepared in duplicate over the indicated concentration ranges, sonicated for 2 min, and equilibrated at 25 °C for 15 minutes. Each sample was filtered through a 0.22 µm pore hydrophobic PTFE syringe filter prior to analysis. UV–visible spectra were collected using a Hewlett Packard 8452A diode array spectrophotometer with a 1 cm quartz cuvette. Full-scan spectra were acquired in 2 nm increments to determine the wavelength of maximum absorbance (λ_max_). Calibration curves were constructed by plotting average absorbance values versus concentration, and molar absorptivities (ε) were determined by linear regression using Beer’s law (A = εlc). For **RPV’-N-C2-AmPEG4-VH032** (**24**), calibration standards were prepared at 5.05, 15.15, 25.75, 37.87, 70.69, 85.84, and 90.89 µM, with a λ_max_ of 294 nm. For **RPV’-N-C2-AmPEG4-Ad** (**26**), standards were prepared at 1.69, 3.37, 8.43, 10.11, 14.16, 16.18, and 24.26 µM, with a λ_max_ of 292 nm. Following filtration, the **RPV’-N-C2-AmPEG4-AnPom** (**21**) exhibited absorbance below the UV–visible detection limit (<3 µM), and no further solubility measurements were performed for this compound. The calculated molar absorptivities were 24,090 M^-1^·cm^-1^ for **24** and 1,486 M^-1^·cm^-1^ for **26**.

#### Kinetic Solubility Measurements

Kinetic solubility measurements were performed in triplicate. Samples (2.5 mL) were prepared by combining PBS (pH 7.4) with compound stock solutions in DMSO, maintaining a final DMSO content of 1% (v/v). The suspensions were sonicated for 3 min, gently stirred for 1 h, and allowed to equilibrate at 25 °C for 18 h. Insoluble material was removed by filtration through a 0.22 µm pore hydrophobic PTFE syringe filter. Filtered solutions were analyzed by UV–visible spectroscopy at the previously determined λ_max_. Kinetic solubility was defined as the concentration of compound present in the filtrate at this apparent supersaturation plateau and was calculated using the corresponding molar absorptivity values (See: **Supporting Information Section 4** for details).

## 3. Results

### 3.1 Design and Evaluation of RPV’-Linker Chemotypes

The overall workflow for computational design and iterative experimental refinement of RT-targeted degraders is summarized in **Figure 3**.

Rilpivirine (RPV, **6**, **Figure 4**) was selected as a warhead among second-generation NNRTIs based on its high potency against RT, favorable clinical safety profile, and the availability of high-resolution RT co-crystal structures enabling the informed rational design and modeling-guided, focused library generation [50]. A 1,3,5-triazine-based analog of 6 (RPV′, **7**, **Figure 4**) was subsequently advanced as a superior warhead alternative to RPV, based on its comparable activity against both wild-type and drug-resistant RT and its improved solubility, as demonstrated for related triazine-based RPV derivatives by Jorgensen and Anderson [51], together with the relative ease and efficiency of linker installation at the triazine 6-position. In addition, the 6-morpholinoethyl attachment found in Jorgensen’s high-potency NNRTI served as a theoretical structural surrogate for the connector and proximal linker region envisioned in our idealized TPD design.

PROTACs and HyT constructs targeting HIV-1 reverse transcriptase (RT) were designed using a multi-stage workflow outlined in **Figure 3**. All structure-based modeling was performed using the full heterodimeric RT complex including both p66 and p51 subunits (PDB ID: 2ZD1) to preserve the native architecture of the NNRTI binding pocket and the adjacent inter-subunit channel relevant for linker placement [8].

To establish whether the RPV’ (**7**) scaffold could tolerate the connector and linker modifications required for degrader design, we computationally and experimentally evaluated a representative set of RPV’ derivatives containing varying connector and linker compositions designed for later E3 ligase ligand attachment (**Table S1**). Structural modeling using core-constrained docking of 232 RPV’-connector-linker constructs (**7, Figure 4**) from a rationally designed library (see workflow, first column, **Figure 3**) identified four favorable chemotypes (amides, ureas, triazoles, and PEG-based) that maintained the native binding position while projecting linkers toward solvent (**Figure 5**; **Supporting Information 2.2 – 2.5**).

**Figure 5.**
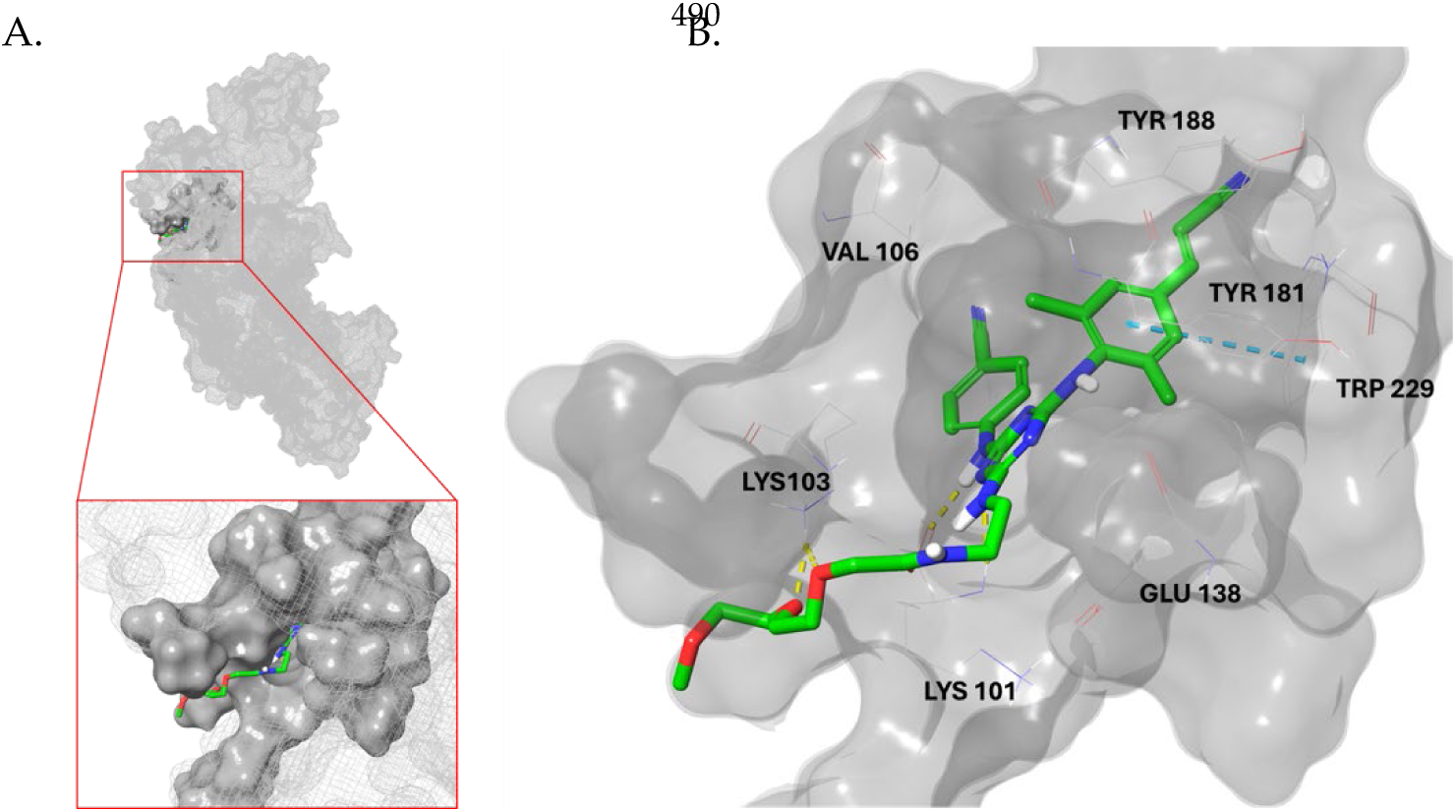
The HIV-1 NNRTI pocket and the narrow inter-subunit channel leveraged during rational linker design and modeling. (A) Surface representation of HIV-1 RT with the NNRTI binding pocket boxed in red; the magnified inset shows the RPV’ warhead seated in the pocket with its linker traversing the solvent-exposed channel toward the protein surface. (B) Close-up of the pocket showing RPV′-N-C2-Am-PEG4 (**10**) (green sticks) occupying the NNRTI site, with the diarylpyrimidine warhead held in the hydrophobic, aromatic core. Linkers that fully threaded the narrow inter-subunit channel toward solvent rather than folding back into the pocket were prioritized. The compound (**10**) shown here is a representative example, with its PEG4 linker extending through the constricted channel to make additional contacts with the channel lysines.

Candidates (n = 35) combining favorable docking scores and predicted solubilities (**Table S1, Figure S1**) were prioritized. From this group, 12 RPV’-Linker conjugate constructs (**Figure 6**) that spanned the computationally top-ranked chemotypes and attachment modes were selected for synthesis (**Supporting Information Section 1**). RPV’-Linker conjugates were tested for inhibition of in vitro activity of purified reverse transcriptase using the Roche Colorimetric Reverse Transcriptase Assay (Roche Diagnostics) to identify those that retained RT occupancy-based inhibitory activity [38]. Ten of the 12 RPV’-Linker conjugates displayed >70% inhibition of RT polymerase activity in a dose-dependent manner, with 50% inhibitory concentrations (IC₅₀) in the low micromolar range **(Figure 7A, Table S1**). The two RPV’-triazole adaptor-based compounds (**16**,**17**) did not exhibit any dose-dependent RT inhibition in this assay. Overall, the experimentally observed potencies were consistent with computational predictions for the synthesized compounds. One compound, N-C2-Am (**10**) had an IC₅₀ within 3-fold of that of RPV (1.75 µM vs. 0.98 µM, respectively) indicating similar potency of the two compounds.

**Figure 6.**
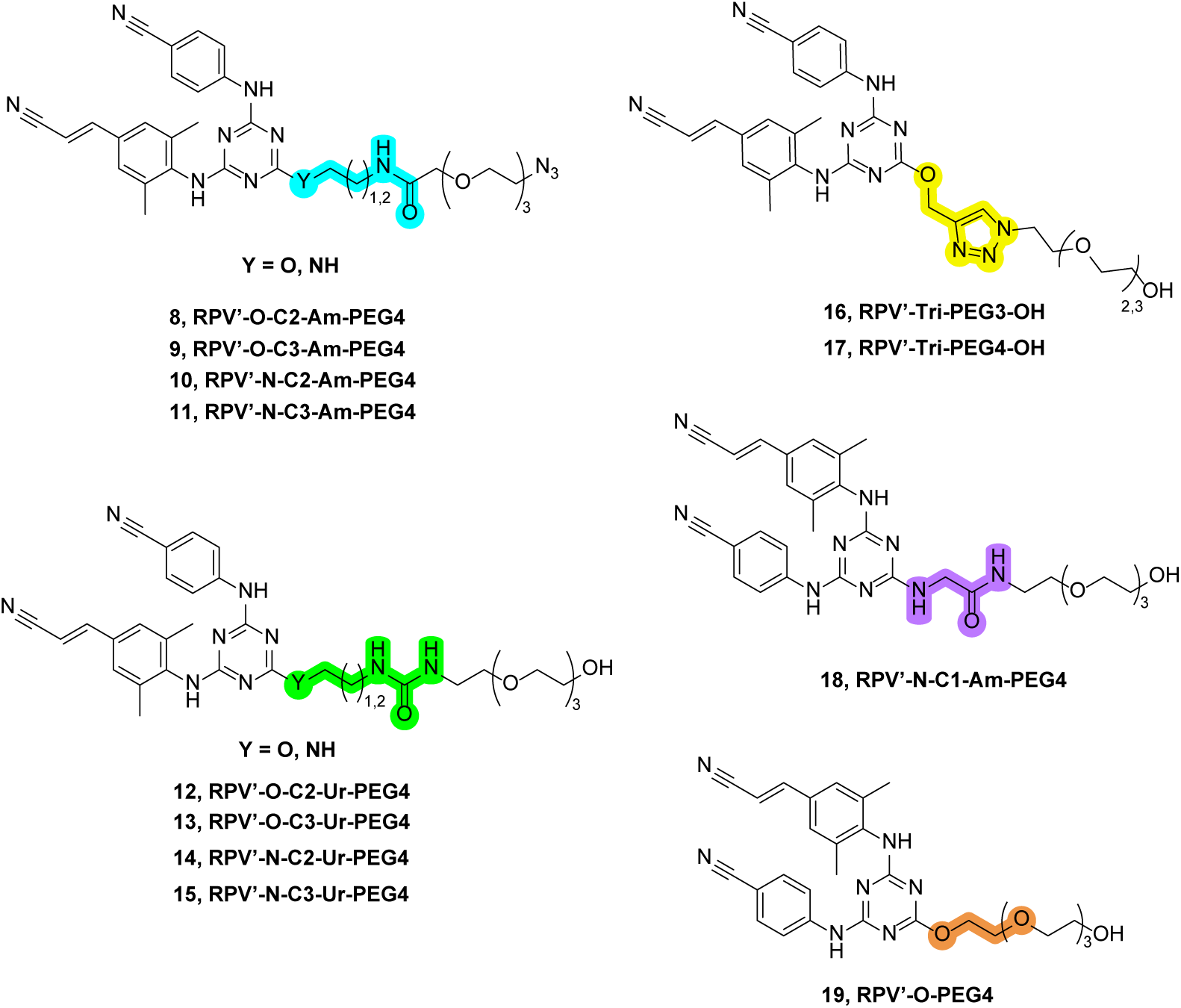
Monofunctional RPV’ warhead-connector-PEG linker constructs prepared to assess relative HIV-1 RT inhibition and single-round infectivity performance. The varied connector segments positioned within the NNRTI entrance channel are highlighted. Connector chemotypes are reflected within the coded names: AM – amide; UR – urea; and Tri – 1,2,3-triazole.

**Figure 7.**
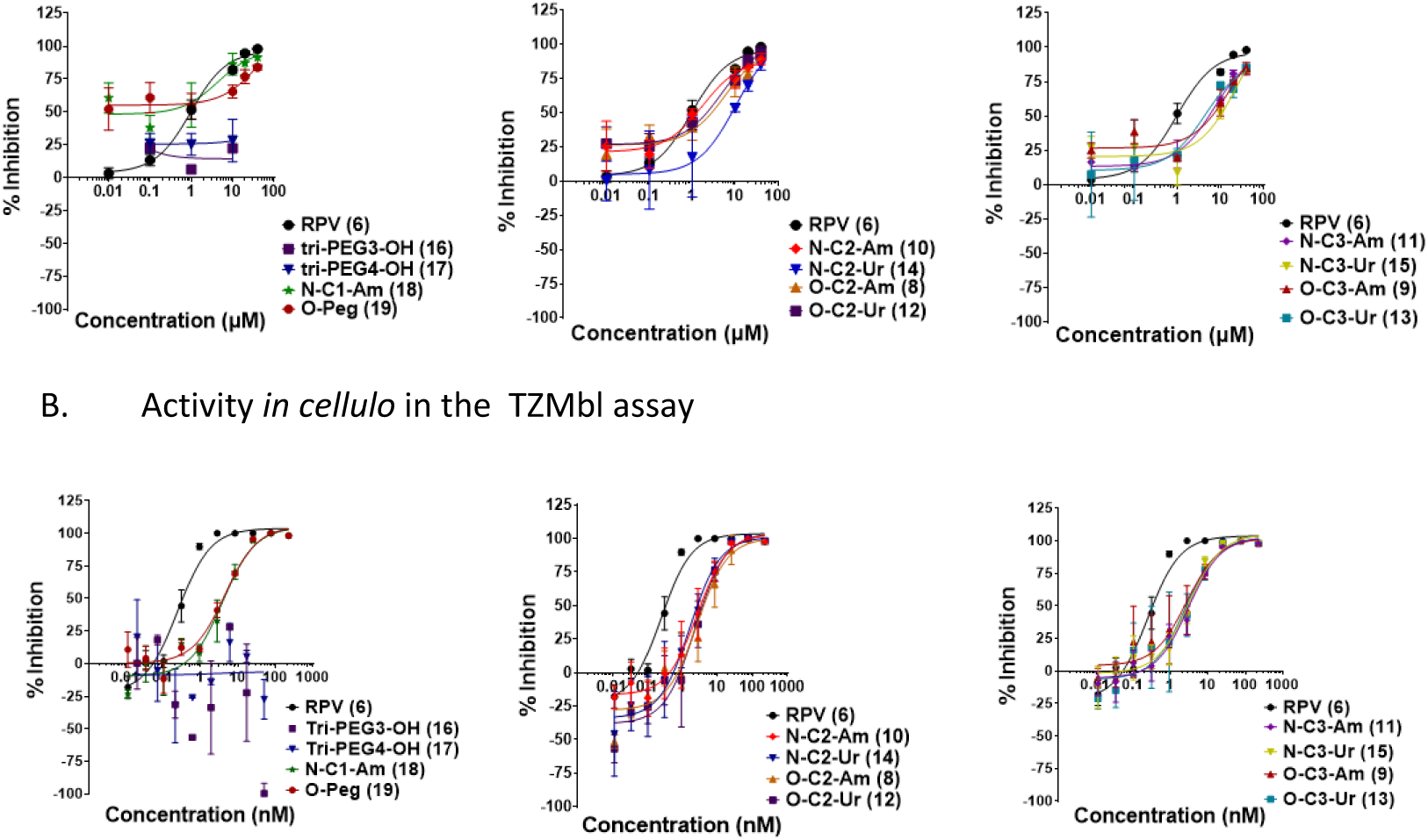
Screening of Monovalent TPD Linker Candidates to Identify Precursors that Demonstrate Activity. Precursor RPV’-TPDs modified with amide (Am), urea (Ur), 1,2,3-triazole (Tri), connectors attached to polyethylene glycol linkers (Figure 6) were screened in the in vitro RT assay (A) and in the TZMbl assay (B). Compounds were grouped by one (left), two (middle), or three (right) intervening methylene groups between the nitrogen or oxygen triazine anchor and the connector to the PEG linker (i.e., highlighted segments in Figure 6.) Compounds **16** and **17** exhibited visually limited solubility at <1 mM. Data is representative of n = 2 biological replicates.

Compounds were then assessed for inhibition of productive HIV-1 infection in the TZMbl single-round infection assay [40, 41] using an HIV-1 infectious molecular clone (IMC) of the reference strain, NL4-3 [52], which we had modified as previously described [39, 53] to encode a Renilla Luciferase reporter cassette (NL4-3-LucR.6ATRi/K5300, referred to as NL4-3-LucR). Apart from the two triazole based connectors, 10 of the 12 compounds displayed >95% inhibition of infection in a dose dependent manner when added at time of infection (t_0_), achieving IC₅₀ values between 2-5 nM (see **Table S2**). However, these IC₅₀ values were approximately 10-fold higher than that of RPV, indicating a loss of potency by the addition of a linker (**Figure 7B**). The most potent compound in the TZMbl assay was N-C2-Ur (**14**) with an IC₅₀ of 1.62 nM. Despite its performance in the TZMbl assay, we decided not to proceed with N-C2-Ur (**14**) for synthesis of bivalent TPD compounds given its comparatively modest activity in the Colorimetric Reverse Transcriptase Assay (IC₅₀=10.11 µM) and its additional hydrogen bond donor and acceptor atom relative to the high potency N-C2-Am (**10**), which performed similarly in the TZMbl assay (IC₅₀ of 2.63 nM), and superior in the RT assay (IC₅₀=1.75 µM) (see Supporting Information, **Table S1**).

### 3.2 Evaluation of RPV’-Amide Connector-based PROTAC and HyT Designs

Building on the 35 RPV′–PEG linkers prioritized through in silico triaging of the initial library (n = 232) and informed by first-round biological screening, 120 bifunctional degraders bearing either CRBN-or VHL-recruiting ligands were computationally enumerated (**Figure 3**, second column). Physicochemical properties were benchmarked against PROTAC-DB [32] to define ranges consistent with cell-permeable degraders. The candidates with PEG3, PEG4, or PEG5 architectures were selected based on favorable molecular weight, polarity, and linker flexibility profiles. Modeling of ternary complexes using protein-protein docking [33] indicated that multiple linker geometries could place the E3 ligase recognition domains (CRBN or VHL) in favorable orientations relative to the RPV binding pocket, with PEG4 to PEG5 architectures providing more favorable linker dynamics/movements and solvent exposure over PEG3 or PEG6 linkers. (**Figure S3**: Protein-protein docking and MD). Molecular dynamics simulations [35] of the modeled ternary complexes predicted that PEG3 would exhibit the most compact and stable geometry (degrader RMSD ∼1–1.6 Å), while PEG5 and longer linkers showed higher mobility and expanded conformations that could compromise ternary complex stability (**Figure 8)**

**Figure 8.**
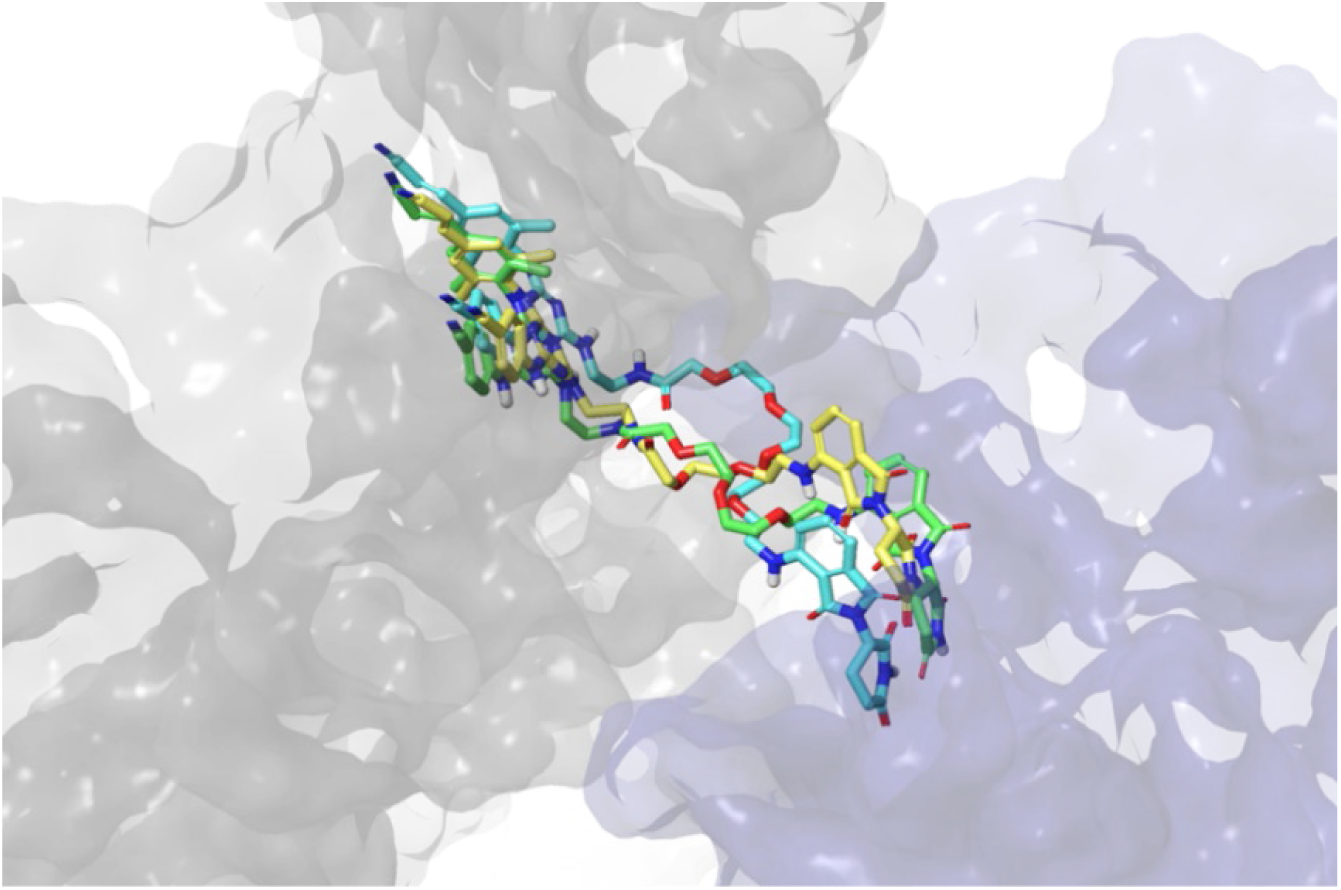
Overlay of representative ternary complex structures from the dominant MD cluster (50– 150 ns) for each system, highlighting the spatial orientation of PEG3 (yellow), PEG4 (green), and PEG5 (blue) linkers.

Based on the MD simulations, eight TPDs were synthesized from the RPV’-N-C2-Am framework (**10**). Three compounds (**20**-**22**) were appended with pomalidomide (Pom), a ligand for Cereblon, which is the recognition domain of the Cullin-4 RING E3 ligase [46, 54–56]: three (**23**-**25**) were appended to VH032, a ligand for the von Hippel-Lindau (VHL) recognition domain of the Cullin-2 RING E3 ligase [57, 58]; and two (**26**-**27**) were appended with an adamantane acetic acid tag to prompt the HyT mechanism [59, 60] of protein degradation (**Figure 9A**, TPD Batch 1). Screening in the in vitro RT assay and in the TZMbl infection assay (**Figure 10 A, B**) showed the VH032 compounds to be the least potent compounds with a maximum percent inhibition of <75% in the RT assay, and a lack of dose-dependent antiviral activity in the TZMbl assays. Compound PEG3-AnPom (**20**) was the most potent inhibitor in the RT assay (IC₅₀ = 6.39 µM); however, it failed to exhibit antiviral activity in TZMbl cells. Surprisingly, the PEG4 homolog **21** demonstrated only partial RT inhibition (55% maximal inhibition) yet showed potent antiviral activity against NL4-3-LucR (IC₅₀ = 23.98 nM; >90% maximal inhibition). The PEG5 analog **22** performed similarly, with a TZMBl IC₅₀ of 31.05 nM (**Figure 10**, middle). The hydrophobic-tagged compounds were active in both assay formats: PEG4-Ad (**26**) inhibited RT and viral infectivity with IC₅₀ values of 22.93 µM and 26.62 nM, respectively, while PEG5-Ad (**27**) displayed comparable antiviral activity with TZMbl IC₅₀ = 21.60 nM (Figure 10, right). IC_50_ values are tabulated in **Supporting Information, Table S1**.

**Figure 9.**
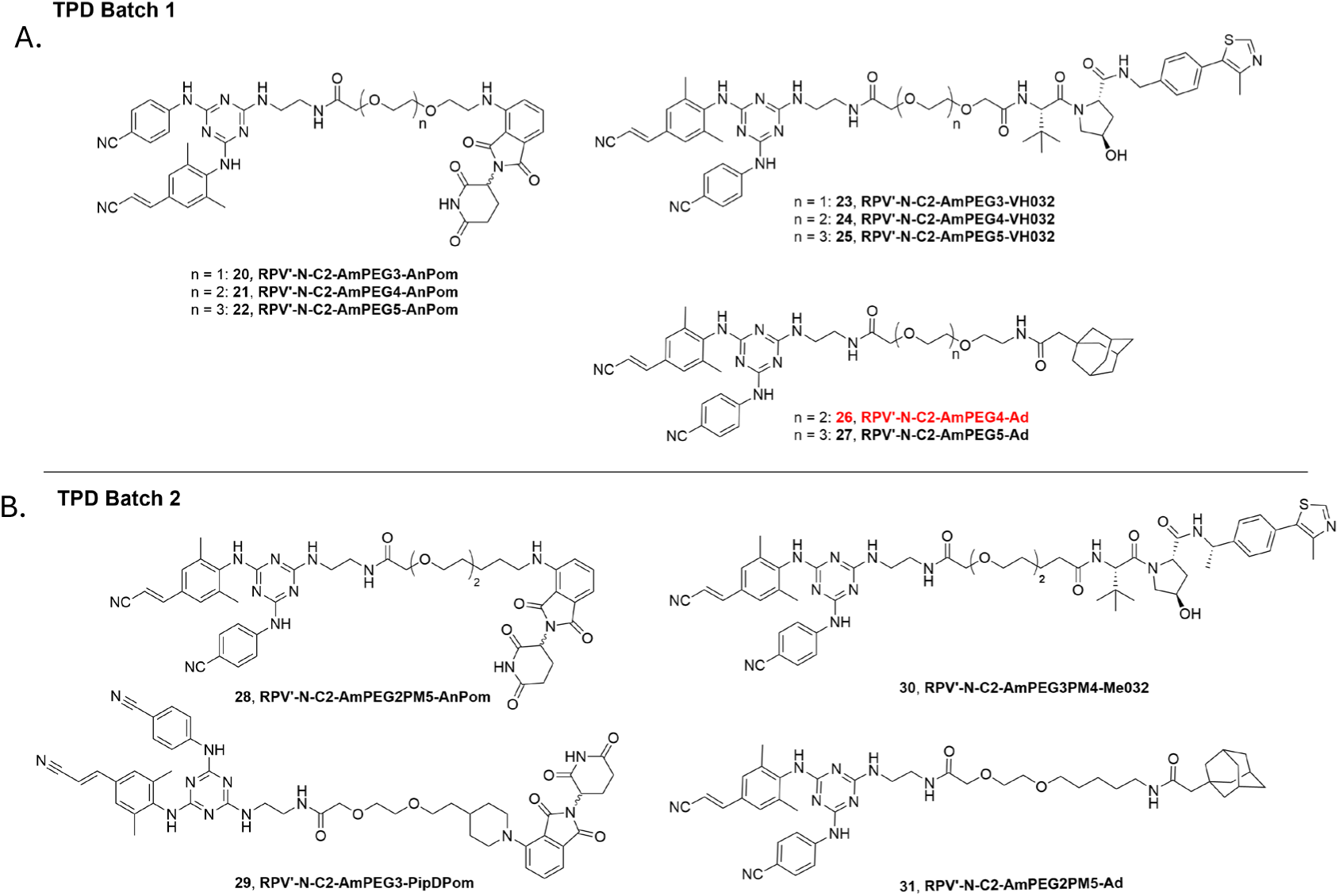
RT-Targeting PROTACs and Hydrophobic Tags (HyTs). (A) Batch 1: Cul-4^CRBN^-targeting (**20**-**22**), Cul-2^VHL^-targeting (**23**-**25**), and hydrophobic tag-bearing (**26**-**27**) TPDs featuring *N*-(aminoethyl)amide connectors and polyethylene glycol linkers of varying length. (B) Batch 2: a focused, four-member linker-tuning series of computationally vetted TPDs (**28**–**31**) designed to evaluate PEG–polymethylene hybrid linker architectures. Compound **26**, a high performing TPD in TZMBl infectivity assays and the compound selected for subsequent mechanism-of-action studies, is highlighted in red.

**Figure 10.**
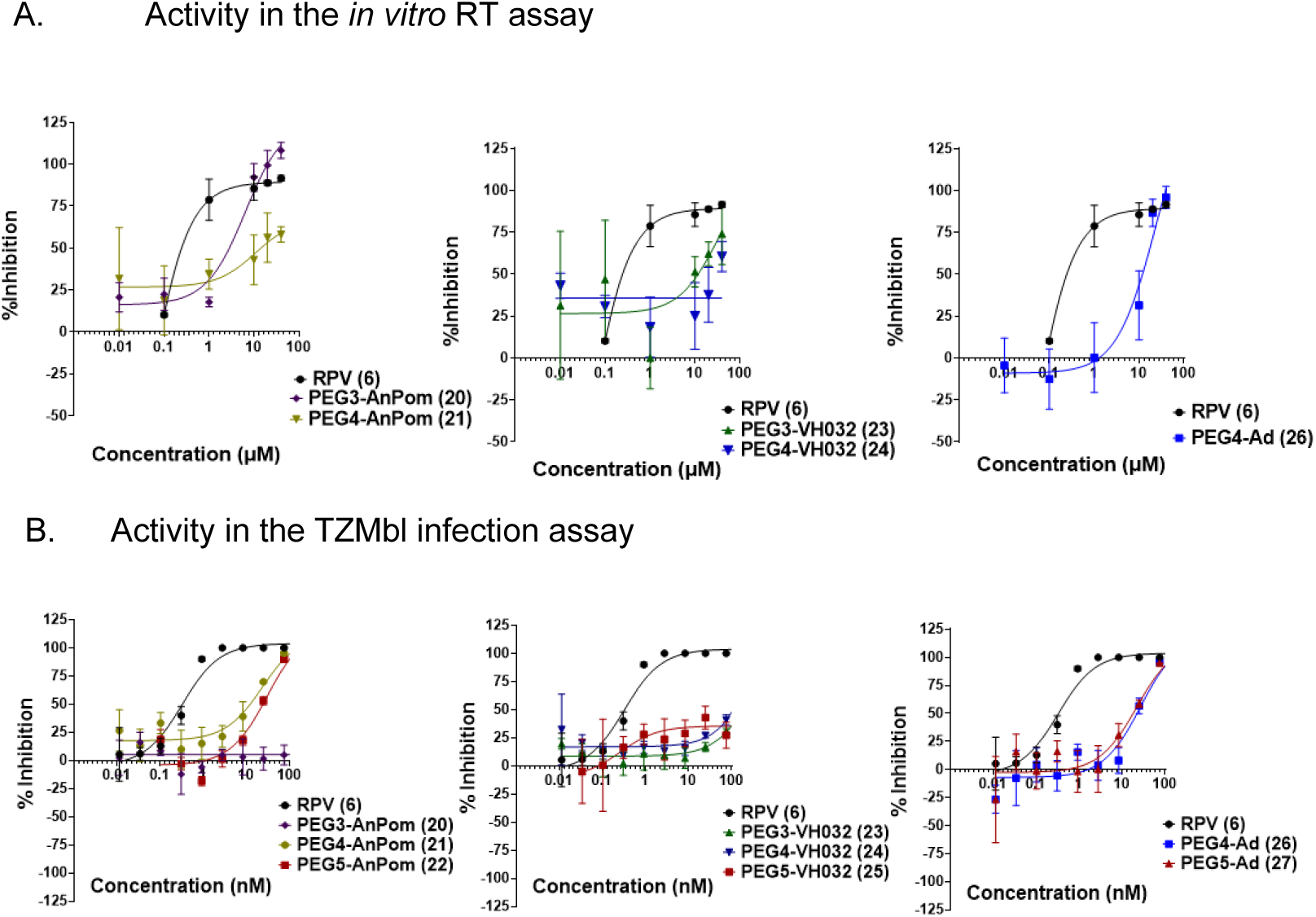
Screening of Batch 1 TPDs 20-27 (shown in Figure 9A). RPV’ appended with ligands to recruit Cereblon (left), VHL (middle) or with a hydrophobic tag (right). Compounds were screened for their ability to inhibit the in vitro polymerization activity of RT (A) and for inhibition of a single round of infection of the in the TZMbl assay (B). Data is representative of n = 2 independent biological replicates.

During the preparation of compound dilutions for the biochemical assays, visible cloudiness or precipitate formation was observed with both PEG4-AnPom (**21**) at concentrations above 1 µM and with PEG4-Ad (**26**) at approximately 25 µM. No turbidity was observed among the precursors lacking a recruiter (**Figure 6**) nor for PEG4-VH032 (**24**) or the PEG-3 or PEG-5 series TPDs (**Figure 9**A) at concentrations approaching 100 µM. These observations prompted a comparative UV-based kinetic solubility analysis of the PEG4 series in PBS (pH 7.4). Following filtration of supersaturated solutions, PEG4-AnPom (**21**) exhibited absorbance below the UV–visible detection limit (<3 µM), whereas the PEG4-Ad (**26**) and PEG4-VH032 (**24**) compounds displayed kinetic solubilities of 16.06 ± 0.88 µM and 101.84 ± 0.65 µM, respectively (**Supporting Information Section 3**). These solubility differences contextualize the inhibition data: The PEG4-AnPom (**21**) produced only partial inhibition (∼55%) in the RT inhibition assay with a shallow response across apparent concentrations from 10–100 µM, consistent with solubility-limited assay behavior. The PEG4-Ad (**26**) compound yielded a sigmoidal concentration–response curve with an IC₅₀ of 22.93 µM, near its measured kinetic solubility, whereas the PEG4-VH032 (**24**) showed no detectable inhibition up to 100 µM, a result not attributable to solubility limitations. Together, these findings illustrate the importance of accounting for compound specific solubility differences when interpreting potency measurements of TPDs at µM concentrations [49].

A focused second series of compounds (Batch 2, **Figure 9B**) was synthesized to explore the effect of subtle changes in linker composition and E3-ligase affinity on degrader performance. Compounds **28**–**31** all featured slightly more lipophilic PEG– polymethylene hybrid linkers. In addition, compound **29** incorporated a rigidified piperidyl attachment to the CRBN-recruiting ligand (PipD-Pom), and compound **30** employed a higher-affinity VHL ligand Me-VH032 [61]. Physicochemical predictions of the polymethylene-linked compounds revealed elevated log P values (3.3–3.9) compared to their PEG counterparts (2.3–3.2), consistent with increased lipophilicity and improved membrane permeability potential (**Supporting Information, Table S1**: Predicted physicochemical properties of connector chemotypes.). All compounds remained within predicted physicochemical property ranges typical of previously described active PROTACs, as assessed using a universal scale derived from PROTAC-DB [32].

To complement these physicochemical predictions with a dynamic assessment of ternary complex behavior, MD simulations of the Batch 2 TPDs (**Figure 9B**) showed that the polymethylene-linked variant PEG2PM5-AnPom (**28**) maintained ternary complex stability comparable to PEG3-AnPom (**20**) (**Figure S3:** Ternary Complex MD analysis). The VHL-recruiting PEG2PM4Me032 (**30)** showed appreciably improved antiviral activity compared to its PEG-linker counterpart **24**, with a TZMbl IC₅₀ of 73.60 nM (**Figure 11, left**). Pomalidomide-appended compounds featuring a PEG-polymethylene hybrid (PEG2PM5-AnPom (**28**)) and a piperazine recruiter attachment (PEG3-PipD-Pom (**29**)) linkers also displayed improved TZMbl antiviral activities with IC₅₀ values of 70.95 and 30.89 nM, respectively (**Figure 11, middle**). The adamantane-tagged compound featuring the polymethylene linker, PEG2PM5-Ad (**31)** demonstrated an IC₅₀ of 12.96 nM (**Figure 11, right**). IC_50_ values are tabulated in **Supporting Information, Table S1**.

**Figure 11.**
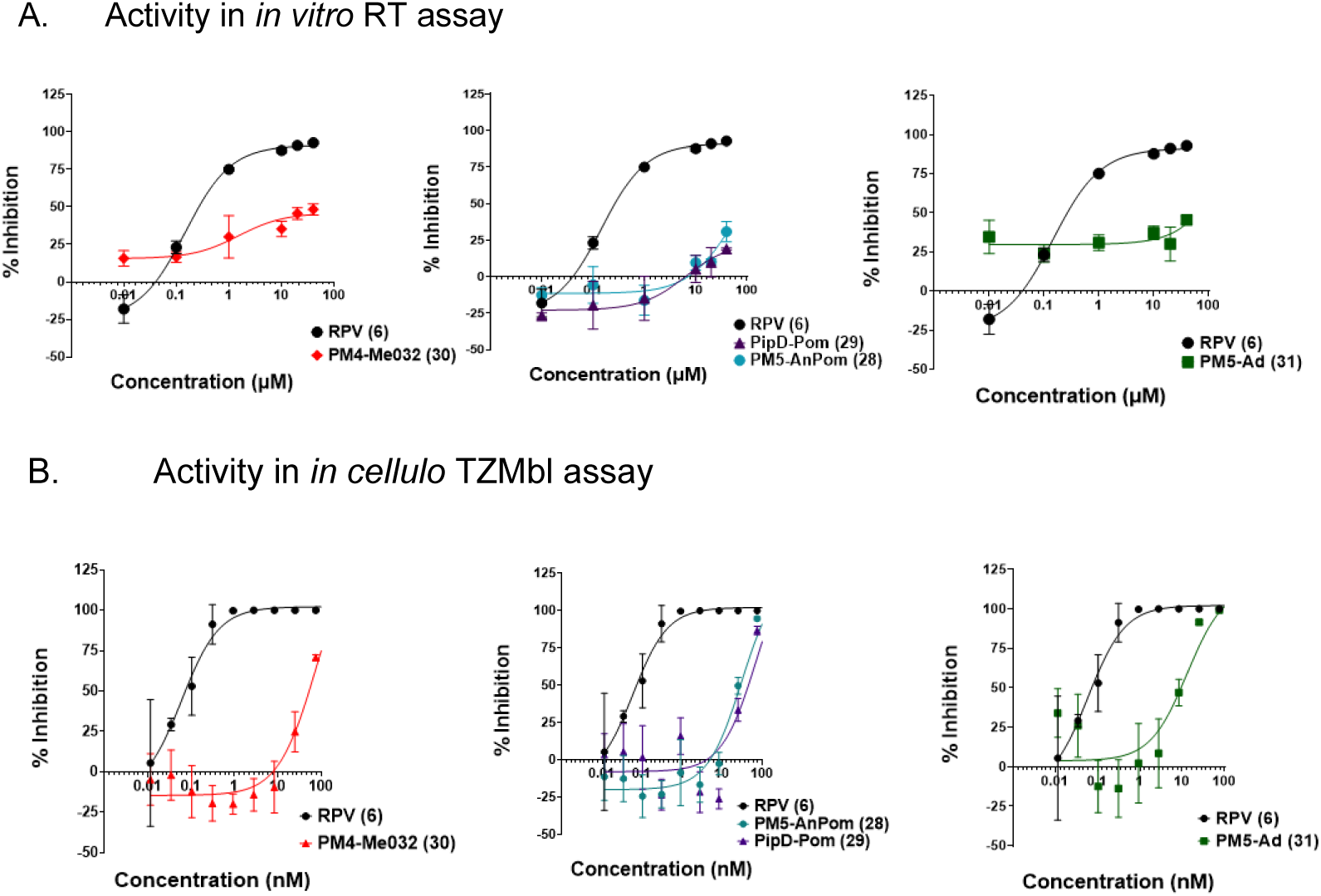
Screening of Batch 2 TPDs with Modified Linkers. RPV’ appended with ligands to recruit VHL (left), cereblon (middle) or with a hydrophobic tag (right) were synthesized with polymethylene linkers (PM4, PM5) and piperazine linkers (PipD). Compounds were screened for their ability to inhibit the in vitro polymerization activity of RT (A) and for inhibition of a single round of infection of the in the TZMbl assay (B). Data is representative of n=2 independent biological replicates

We performed MD simulations on variants of adamantane-tagged HyT compounds (**26**, **27**, and **31**) to evaluate how linker architecture influences the conformational dynamics and solvent exposure of the hydrophobic tag – critical features that static docking poses and calculated physicochemical properties fail to capture. A matched theoretical PEG3 analog was included to complete the PEG3–PEG5 series and isolate linker-length-dependent effects while retaining the same RPV′ warhead and adamantane tag. Physicochemical properties were comparable across the HyT series (Plog 4.2-4.5; **Table S3:** Predicted physicochemical properties and docking scores of HyTs). Using a threshold of ∼70 Å² exposed nonpolar surface area as a proxy for hydrophobic patch recognition by protein quality control machinery [62]. PEG4-Ad (**26**) exhibited the most favorable exposure dynamics with approximately 12% of simulation frames exceeding this threshold (**Figure 12**). PEG2PM5-Ad (**31**), which also showed activity *in cellulo*, exceeded the threshold in ∼5% of frames. In contrast, the theoretical PEG3-based analog remained compact with SASA centered near 25-40 Å², notably, rarely presenting the adamantane tag to solvent. PEG5-Ad (**27**), despite its longer linker, showed frequent back-folding that limited high-exposure events to ∼3% of frames (**Figure 12)**

**Figure 12.**
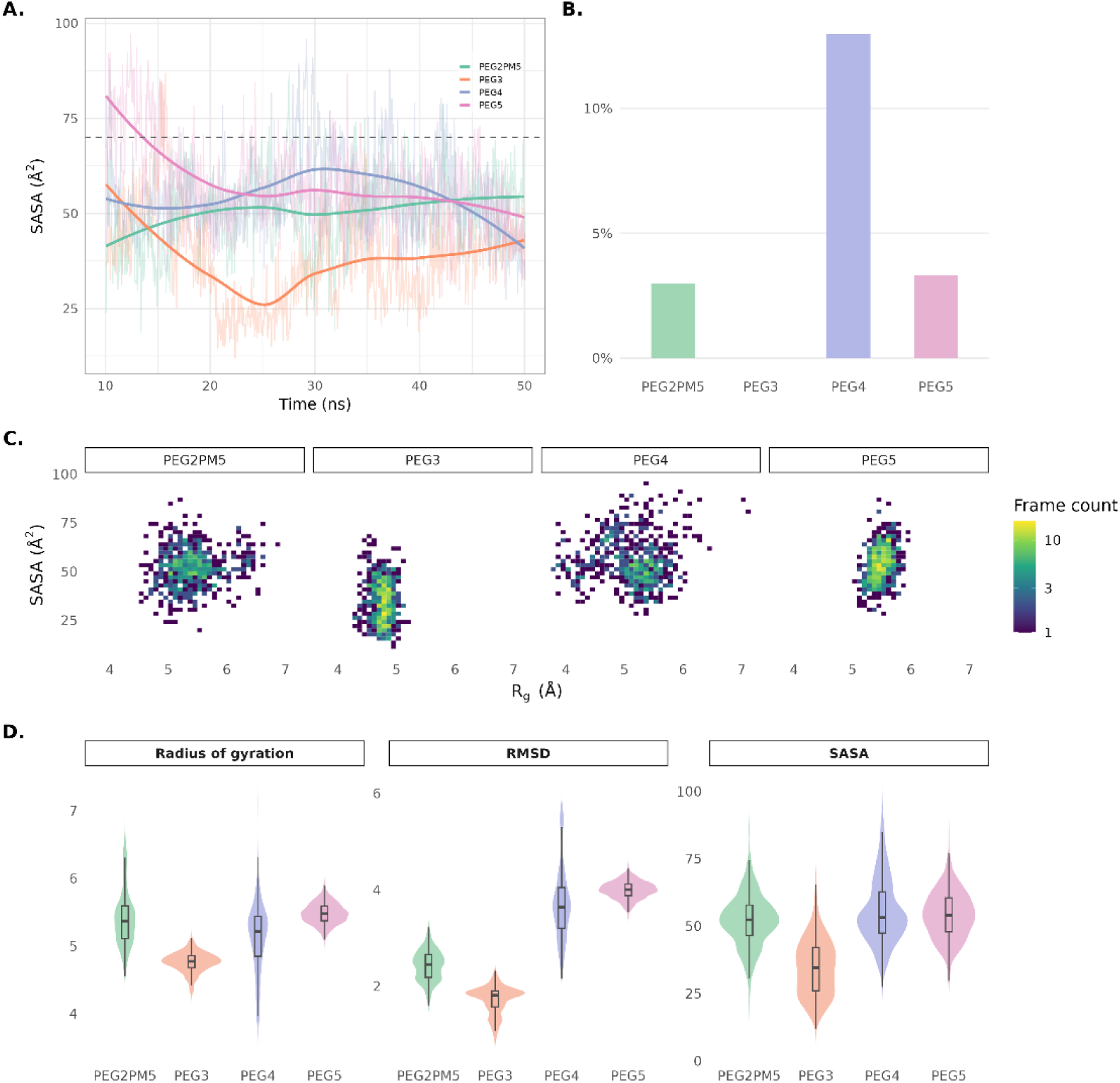
Post-equilibration molecular dynamics analysis of hydrophobic-tagged RPV′ conjugates bound to HIV-1 reverse transcriptase. The four HyTs analyzed were RPV′-N-C2-Am-PEG3-Ad (theoretical), RPV′-N-C2-Am-PEG4-Ad (**26**), RPV′-N-C2-Am-PEG5-Ad (**27**), and RPV′-N-C2-Am-PEG2PM5-Ad (**31**), simulated in explicit solvent. (A) Time evolution of linker solvent-accessible surface area (SASA) over the full 10–50 ns trajectory, with the dashed line marking an operationally defined 70 Å² cutoff used to flag solvent-exposed adamantane states. The early trajectory reflects relaxation from the initial docked poses, with the two exposure extremes relaxing in opposite directions: PEG5 (**27**), docked in an extended conformation, begins with its adamantane solvent-exposed and retracts into the pocket within ∼20 ns, whereas PEG4 (**26**), docked in a more compact pose, extends outward and sustains adamantane presentation for most of the trajectory. Quantitative comparisons in (B–D) are therefore restricted to the equilibrated 20–50 ns interval. (B) Percentage of trajectory frames over the 20–50 ns interval in which linker SASA exceeded the 70 Å² cutoff; the PEG4 conjugate (**26**) surpassed this cutoff in 13.0 % of frames (≈3.9 ns cumulative, distributed across ∼21–43 ns), roughly four-fold more than any other conjugate (PEG5, 3.3 %; PEG2PM5, 3.0 %; PEG3, 0 %). (C) Two-dimensional distributions of linker SASA versus radius of gyration (Rg*) over the 20–50 ns interval, with color indicating the number of frames within each bin. (D) Distributions of linker Rg, root-mean-square deviation (RMSD), and SASA over the 20–50 ns interval. Rg and RMSD are reported in Å, and SASA is reported in Å². *Rg describes linker compactness, with lower values indicating a more compact conformation.

The distinct conformational regimes underlying these exposure profiles are further illustrated by representative MD cluster medoids (**Figure 13**). PEG4-Ad (**26**) projects the adamantane toward bulk solvent, whereas the theoretical PEG3 analog, PEG5-Ad (**27**), and PEG2PM5-Ad (**31**) adopt more enclosed conformations within or adjacent to the RT pocket.

**Figure 13:**
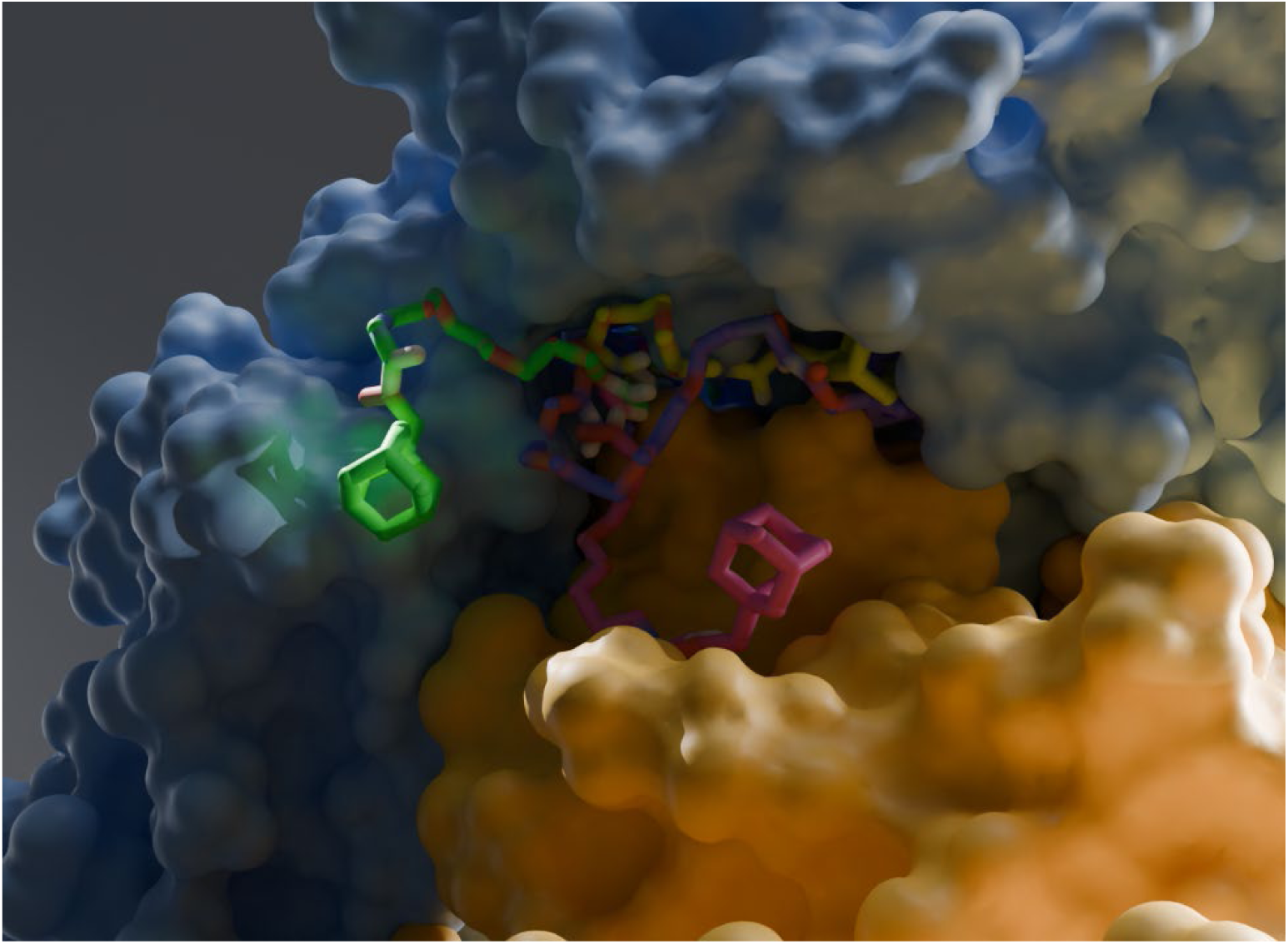
Overlaid molecular dynamics cluster medoids of hydrophobic-tagged RPV′ conjugates bound to HIV-1 reverse transcriptase. Representative conformations of the same four HyTs analyzed in Figure 12 are overlaid within the NNRTI binding channel of HIV-1 RT: RPV’-N-C2-Am-PEG3-Ad (theoretical), RPV’-N-C2-Am-PEG4-Ad (**26**), RPV’-N-C2-Am-PEG5-Ad (**27**), and RPV’-N-C2-Am-PEG2PM5-Ad (**31**). For each conjugate, the representative structure is the medoid of its explicit-solvent MD trajectory, obtained by clustering the linker+adamantane heavy-atom RMSD over the 10–50 ns interval (DBSCAN) and selecting the frame whose RMSD was closest to the cluster mean. The overlaid medoids were rendered in Blender 4.4 using the Molecular Nodes add-on; the two protein surfaces correspond to the p66 (blue) and p51 (peach) subunits of HIV-1 RT. Conjugates are shown as sticks colored by linker: PEG2PM5 (pink), PEG3 (yellow), PEG4 (green), and PEG5 (purple). The overlay highlights differences in linker orientation and adamantane positioning among the four HyT constructs.

Rilpivirine and other NNRTIs have been described to bind to RT precursors in GagPol during virion assembly with the effect of stabilizing the transient “encounter complexes” of GagPol/RT-p66 homodimers, which also results in prematurely stabilized precursor protease (PR) dimers and perturbed PR (auto)processing and virion maturation [42–44]. This poses an additional opportunity for the TPDs to engage dimeric RT during the late stage of the viral life cycle that is not captured in TZMbl single round infection assays. Thus, we sought to determine if RPV’-TPDs could have a deleterious effect on viral replication, possibly due to interactions with RT at the late stage of viral assembly.

Given the superior performance of the HyT compounds over the VHL compounds, and the more limited solubility of the Pom-based compounds, we selected the HyTs PEG4-Ad (**26**) and PEG5-Ad (**27**) for further characterization in multi-round infection assays for which we utilized a CD4 T cell line derived from C8166-R5 cells [43] modified with an LTR-driven secreted nano-Luciferase (snLuc) reporter gene (see Materials and Methods). The T cells were infected with NL4-3-LucR [39] in the presence of RPV’-HyTs **26** and **27**, respectively, at concentrations of 225, 25, and 2.75 nM and cultured over a course of six days with compounds replenished every two days. The protease inhibitor indinavir (IDV), which limits infection to a single round by preventing the maturation of newly formed virions [63], was used as a control for viral replication, which we quantified as the level of snLuc activity in the culture supernatant, relative to uninfected controls. We observed a dose-dependent reduction in the snLuc activity by the tested RPV’-HyTs (**Figure 14**). The T cell cultures treated with 225 nM of the RPV’-HyTs remained uninfected over the six days of infection, whereas those treated with 25 nM showed a 20-fold increase in snLuc activity over the uninfected control. RPV’-HyTs at 2.75 nM did not have an inhibitory effect, as we observed a greater than 100-fold increase in the snLuc activity to levels essentially identical to the inhibitor-free condition (**Figure 14A**).

**Figure 14.**
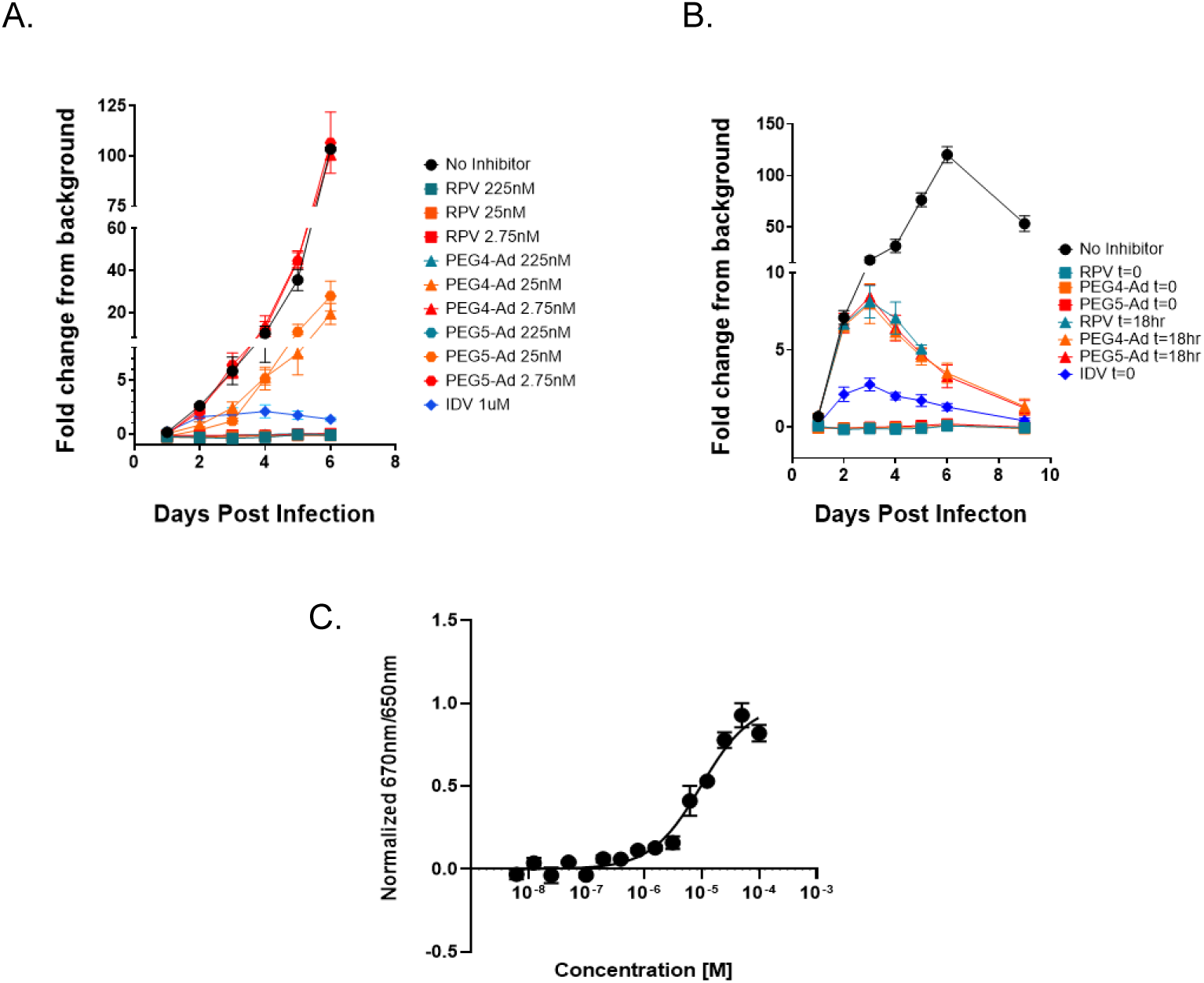
RPV’-HyTs Inhibit Infection in a Multi-Round T-Cell based Infection Assay. (A) C8166-R5 derived CD4 T cells (D1473) were infected with HIV NL4-3-LucR reporter virus in replica plates in the presence of RPV or RPV’-HyTs PEG4-Ad (**26)** and PEG5-Ad (**27)** at the indicated concentrations. Culture supernatants were sampled at the indicated time points and snLuc activity, measured as Relative Light Units (RLU) was determined. The fold change of RLU was calculated with regard to uninfected controls. (B) RPV or RPV’-HyTs were added either at the time of infection (t=0) or 18 hours post infection (t=18) at 225nM. snLuc RLU were recorded as done in (A). Decline in the fold-change over no-inhibitor condition at day 9 is due to cell death. N=2 biological replicates. (C) MST binding curve of PEG4-Ad **(26)** to purified RT.

Next, we assessed the ability of PEG4-Ad (**26**) and PEG5-Ad (**27**) to inhibit further cycles of cell-to-cell spread following the establishment of a first round of infection by inoculating cell cultures with HIV-1 18 hours prior to the addition of RPV’-HyTs at 225 nM (**Figure 14B**). In treated cultures, infection peaked three days post inoculation with a 9-fold increase of snLuc activity above the uninfected control, followed by subsequent decreases resulting in a snLuc levels similar to that of the IDV treated condition at day 9 post infection. The reduction in snLuc signal after day 3 in the IDV and RPV’-HyT treated groups suggests that the first round of HIV replication was complete after 72 hours and that PEG4-Ad (**26**) and PEG5-Ad (**27**) were able to inhibit further rounds of HIV replication following a single round of replication.

We next employed microscale thermophoresis (MST) to characterize PEG4-Ad (**26**) binding affinity for RT. By utilizing a purified 6X His-tagged RT [44] we first determined that our protein sample was suitable for using a His-Tag labeling system to determine the dissociation constants of ligands from RT. The Kd of PEG4-Ad (**26**) was determined to be 12.53 ± 3 µM after averaging the data across the duplicates and fitting it to the Hill model (**Figure 14C**). MST was found to be unsuitable for the calculation of Kd of RPV because the fluorescence ratio (670 nm/650 nm) at 12.5 nM exceeded that of PEG4-Ad (**26**) at saturation, indicating that the autofluorescence signal from RPV was likely masking the changes that would be observed due to RT/RPV binding interactions. Thus, we turned to nano-differential scanning fluorimetry to assess binding of PEG4-Ad (26) relative to RPV. RPV induced a 3°C shift in the thermal unfolding of RT whereas PEG4-Ad (26) induced a 0.8°C shift in the thermal unfolding of RT suggesting a lower binding affinity (data not shown). In summary, the results from experiments designed to estimate binding affinity underpin the notion that RPV’-TPDs do have the ability to bind RT, though likely with reduced affinity compared to RPV.

### 3.3 Mechanistic Studies

Next, we sought to determine whether the RPV’-HyTs may induce proteasomal degradation of RT. To do this, we aimed to cytosolically express the RT heterodimer, composed of p51 and p66 subunits, as a proof-of-principle model system for the degradation of RT mediated by PEG4-Ad (**26**) in the absence of the other HIV viral proteins. To circumvent the need for proteolytic processing of p66 by HIV Protease (PR) to derive p51, as is the case during the HIV replication cycle, we expressed both subunits mono-cistronically with a P2A sequence following p51 to allow for the separation of the subunits during translation. We also appended the p66 subunit with a HiBiT tag on the C-terminus (RT/ p51-p66.HiBiT) to enable the sensitive quantification of RT protein levels through bi-molecular complementation after the addition of the LrgBiT peptide to cell lysates to reconstitute nanoluciferase, as is schematically outlined in **Figure 15A**. We confirmed equimolar expression of both RT subunits from the p51-P2A-p66.HiBiT LVV in transfected 293T cells (**Figure 15B**). We then stably transduced HEK 293 cells to assess degradation of RT in the presence of PEG4-Ad (**26**). To establish optimized assay conditions, we titrated the amount of doxycycline used to induce gene expression and identified a concentration range that resulted in sensitively quantifiable p66.HiBiT levels above background, which were also low enough to not fully saturate the HiBiT signal (**Figure S4**). Using this approach, we found a statistically significant (p=0.0243) reduction in p66.HiBiT levels upon administration of 5 µM of PEG4-Ad (**26**) by HiBiT readout (**Figure 15C**). To verify that these results were not attributable to cytotoxicity, we assessed both live cell counts and cell viability and saw no differences in these parameters between the RPV’-HyT treated groups compared to RPV and the untreated control (**Figure 15F**). We next investigated whether the reduction of RT/p51-p66.HiBiT levels occurred via proteasomal degradation by including the reversible proteasome inhibitor MG132 in the assay. Addition of 5 µM MG132 restored RT/p51-p66.HiBiT levels to that of the 10 ng/mL doxycycline only treated control, indicating that the reduction in RT/p51-p66.HiBiT levels observed was dependent on an active proteasome (**Figure 15C**). To further validate these findings, western blots were performed to visualize p66.HiBiT protein expression (**Figure 15D**). Semi-quantification of the p66.HiBiT bands in Figure **15D** indicated a greater than 70% decrease in p66.HiBiT levels in the presence of 5 µM and 10 µM PEG4-Ad (**26**), whereas addition of MG132 restored RT to levels exceeding the 10ng/ml doxycycline only treated control (**Figure 15E**). Interestingly, the western blot analysis, using anti-HIV-IgG (panel a), failed to detect p51 in stably transduced cells in which gene expression was induced with the low concentration of 10 ng/ml doxycycline. This was surprising considering that western blot analysis after transfection of 293T with RT/p51-p66.HiBiT LVV plasmid indicated the presence of both p51 and p66 subunits. However, overall protein levels are likely much higher in transfected cells, in part to the much higher gene expression induction with 1µg/ml dox.

**Figure 15.**
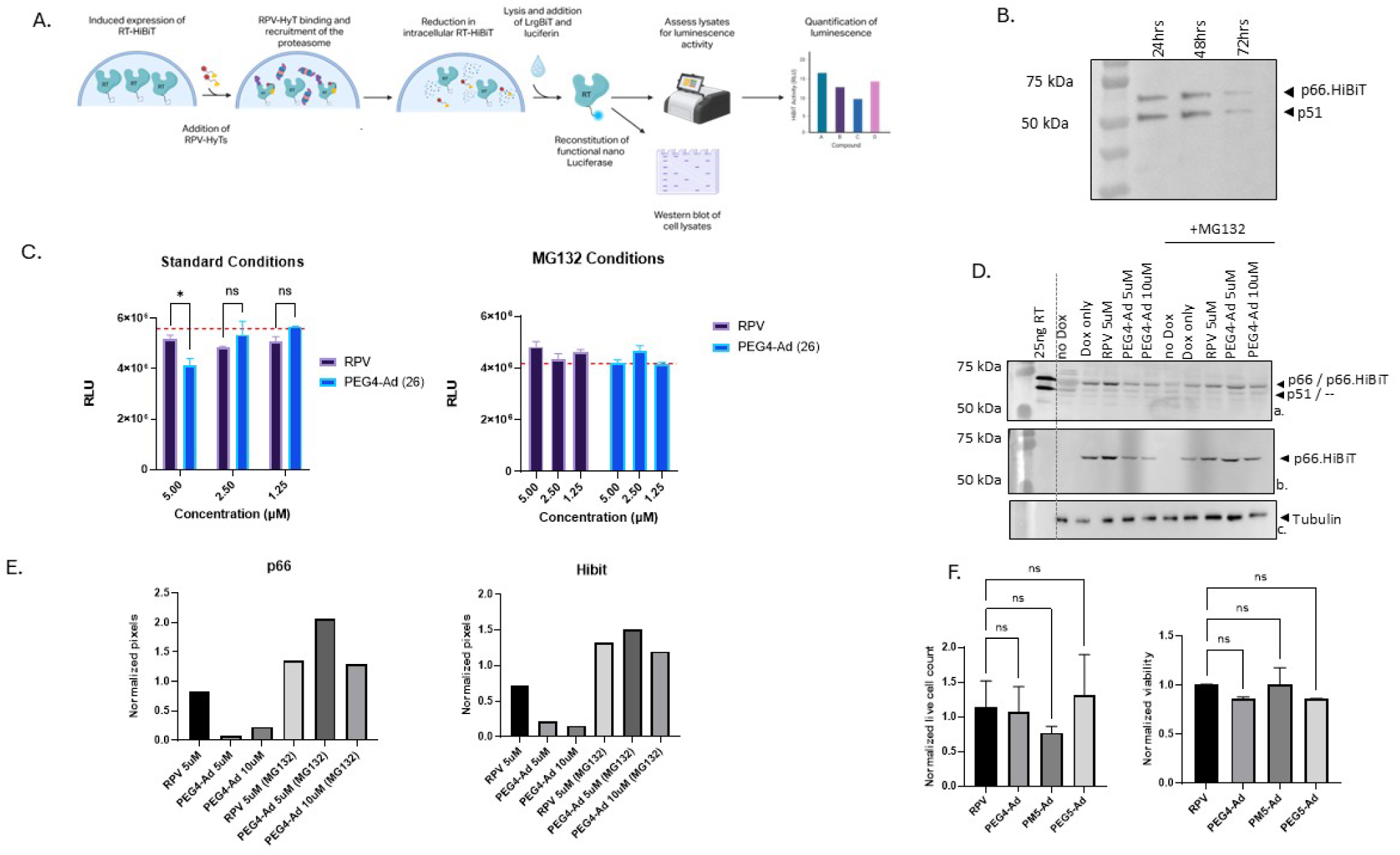
Evaluation of PEG4-Ad’s degradation mechanism. (A) Schematic representation of PEG4-Ad degradation assay: Expression of p51-P2A-p66.HiBiT in stably LVV-transduced cells is induced with 10 ng/mL doxycycline 24 hrs prior to treatment with RPV’-HyTs. 24hrs post treatment, cells are lysed and LrgBiT and luminescence substrate are added. Functional nano-luciferase is reconstituted upon bimolecular complementation of LrgBiT with HiBiT, and HibiT levels are then quantified as Relative Light Units (RLU) by luminescence; a Western blot is also conducted to visualize protein levels. Created in BioRender. Amanor, F. (2026) https://BioRender.com/2ggbqx8 (B) Equimolar expression of p51 and p66.HiBiT in 293T cells transiently transfected with p51-P2A-p66.HiBiT LVV plasmid as shown by Western blot. Lysates were collected at the indicated time points post transfection and 1µg/ml doxycycline induction and probed for the RT polymerase domain with the 8C6 anti-RT antibody (1:250). C) Relative light units (RLU) luminescence quantification of p66.HiBiT under degradation assay standard conditions as outlined .in (A) with RPV and PEG4-Ad (26) added at the three indicated concentrations in the absence (left graph), or presence (right graph) of 5µM MG132. Dotted red lines indicate the mean RLU of doxycycline induced controls (left) or doxycycline and MG132 treated controls (right), respectively. A two-way ANOVA followed by Sidak’s multiple comparison test was used to determine statistical significance. (D) Lysates used in (C) were used to probe for RT by Western blot (panel a) using pooled HIV-IG (1:100) and mouse anti-human-IgG-HRP, and for p66.HiBiT was via the HiBiT blotting assay (Promega) (panel b). Purified heterodimeric RT was used as control. (E) Semi-quantification of western blots in (D). RT and HiBiT band pixel intensities were first normalized to those of tubulin bands and then subsequently normalized to the pixel intensity of the dox-only treated control band. (F) HEK 293F cells stably transduced with p51-p2a-p66.HiBiT LVV and induced with doxycycline were treated with 5µM RPV or RPV’-HyTs for 72hrs and stained with Trypan blue. Cell counts and viability were assessed using a Cellometer. Cell counts and viability were normalized to untreated control samples. A 1-way ANOVA followed by Dunnett’s multiple comparison test was used to determine statistical significance.

Some HyTs have been reported to be dependent on HSP70 recognition for their mechanism of protein degradation [18, 64]. To determine if PEG4-Ad’s (**26**) mechanism of degradation was mediated by HSP70, we sought to upregulate HSP70 expression in stably transduced, RT/p51-p66.HiBiT expressing cells by treatment with the HSP90 inhibitor 17-AAG. We observed an upregulation in HSP70 expression with as little as 0.037 µM 17-AAG (**Figure 16A**). Co-treatment with RPV’-HyTs or RPV at 5 µM and 17-AAG at 1 µM did not disrupt HSP70 upregulation (**Figure 16A**). No cytotoxic effects were observed with 17-AAG treatment (**Figure 16B and 16C**). However, despite HSP70 upregulation in the presence of 17-AAG, we did not observe a significant increase in degradation of p66.HiBiT by PEG4-Ad (**26**) when co-administered with 0.33 µM 17-AAG by HiBiT readout **(Figure 16D**). This suggests that PEG4-Ad’s (**26**) proteasome dependent degradation is independent of HSP70 recognition.

**Figure 16.**
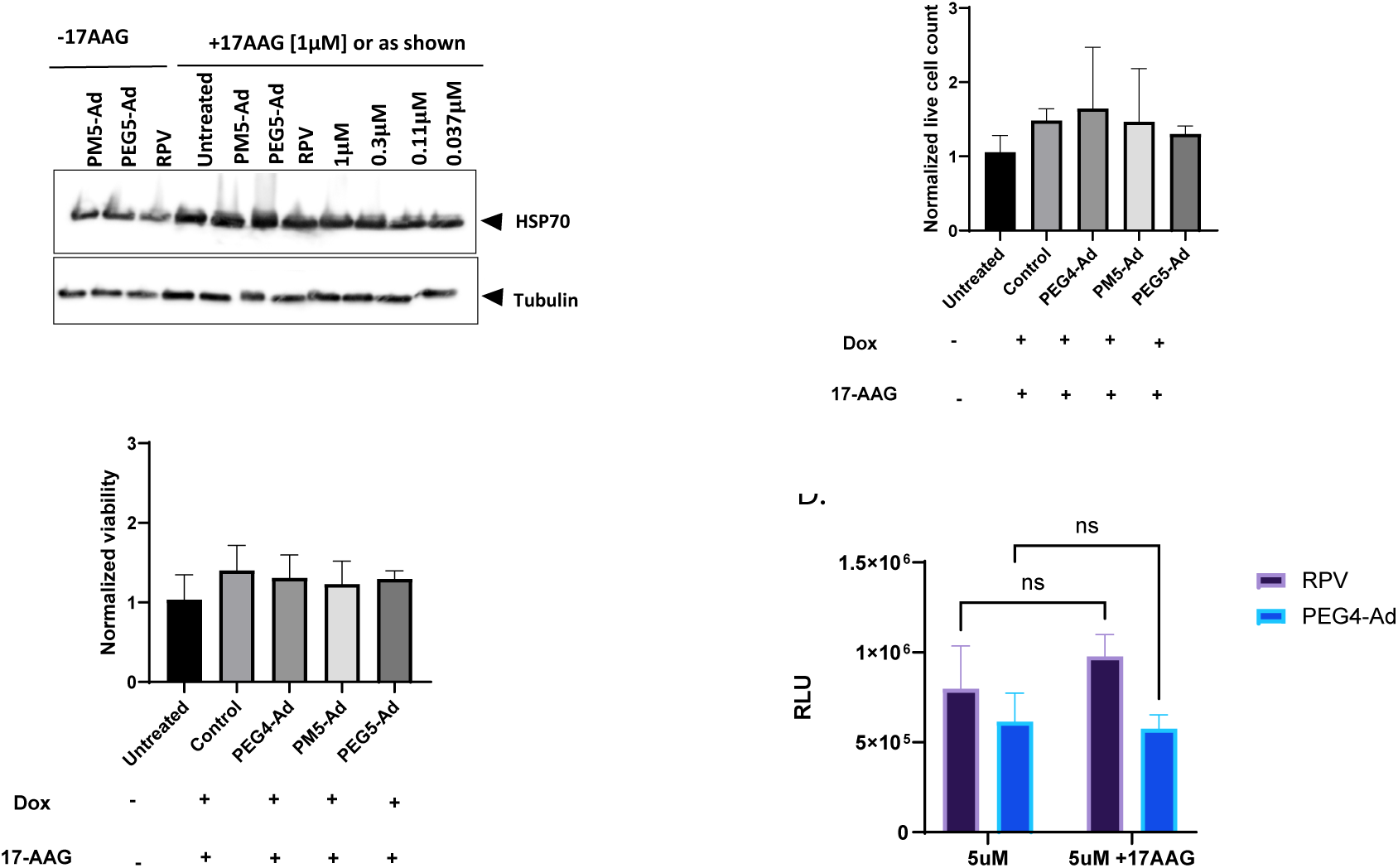
HSP70 is not involved in PEG4-Ad’s mechanism of degradation. (A) Western blot probing for HSP70 and Tubulin of 10ng/ml dox induced 293F cells stably transduced with p51-p2a-p66.HiBiT LVV. Cells were treated for 72 hr with two RPV’-HyTs or RPV at 5 µM (left 3 lanes); or, in the presence of 1 µM 17AAG, mock treated, treated with RPV or RPV’-HyTs, or treated with 17-AAG alone at the indicated concentrations (untreated refers to cells treated with 17-AAG but not with doxycycline). Live cell counts (B) and cell viability (C) after treatment with 5 µM RPV or RPV’-HyTs ± 1 µM 17-AAG for 72hrs, normalized to control. (D) Luminescence quantification of p66.HiBiT in a degradation assay conducted in the presence of 5 µM RPV or PEG4-Ad ± 0.33 µM 17-AAG. A two-way ANOVA followed by Sidak’s multiple comparison test was used to determine statistical significance.

Several studies have shown that TPDs exhibit improved activity against target proteins carrying resistance mutations relative to conventional occupancy-based inhibitors. This effect is attributed to the formation of productive ternary complexes, where additional protein–protein and ligand-mediated interactions can compensate for losses in binary target binding caused by the mutation. [25, 65, 66]. Additionally, some HyTs have also been shown to be effective against variants resistant of their parent drug inhibitors, although the mechanisms for these improved potencies have not been identified [67, 68]. Thus, we investigated whether PEG4-Ad (**26**) would exhibit higher potency compared to RPV against RT in the presence of three clinically relevant NNRTI resistance mutations (E138K, K103N/Y181C, and Y181V) by determining IC_50_ values in the single round infection TZMbl assay (**Figure 17**). Assessing PEG4-Ad (**26**) inhibition against the K103N/Y181C mutant, we found a similar reduction in potency as compared to RPV. Against the E138K variant, the reduction in potency was 2-fold greater for PEG4-Ad (**26**) than for RPV. Against the Y181V mutant, which confers a 20-fold reduction in RPV IC_50_, we did not reach an IC_50_ for PEG4-Ad (**Figure 17 B**). However, our dilution series of PEG4-Ad (**26**) did not cover a concentration range wide enough to allow determination of a Y181V IC₅₀ value 20-fold higher than the IC₅₀ against wild type RT

**Figure 17.**
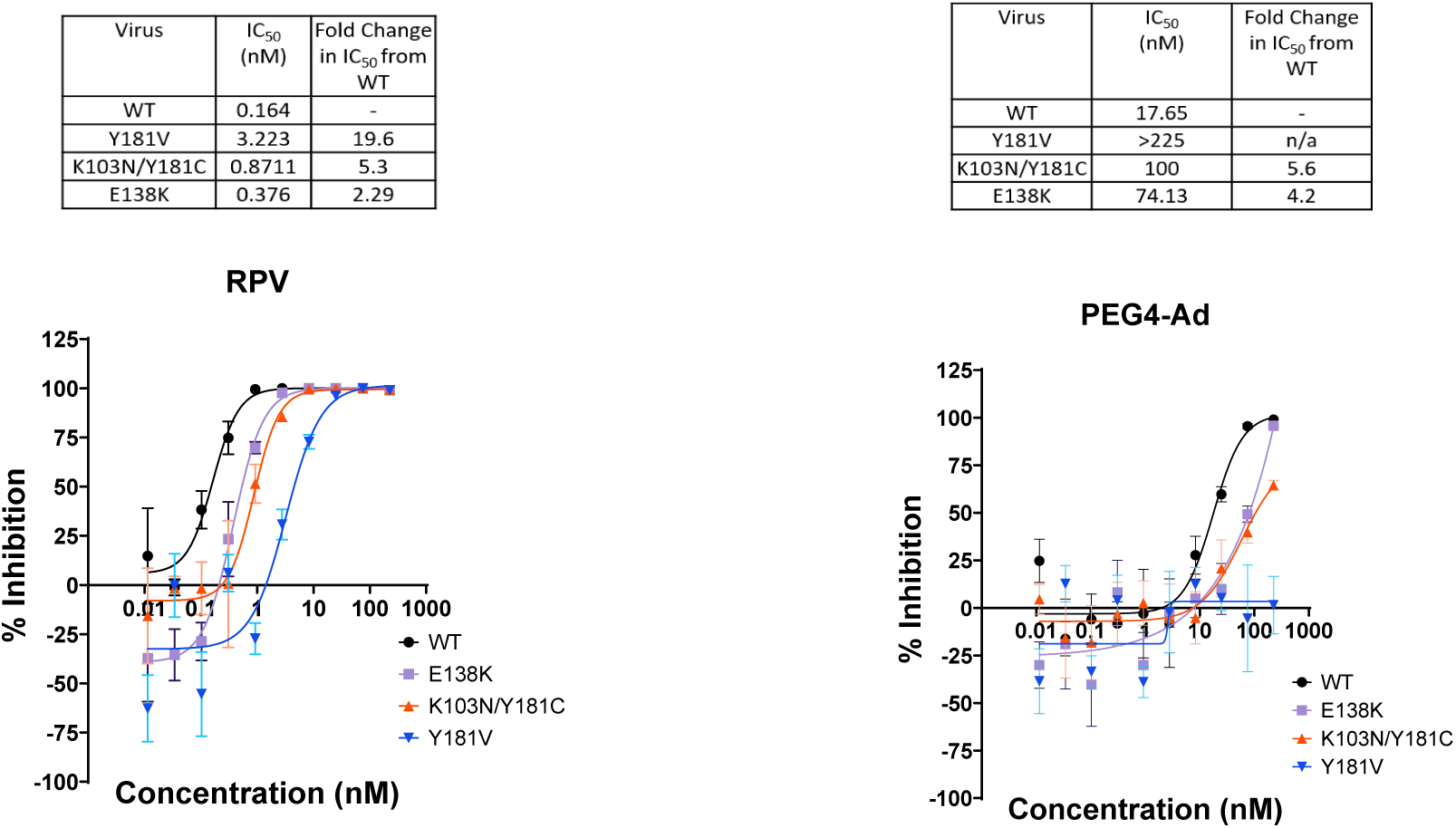
Screening of PEG4-Ad (26) against common NNRTI resistance mutations. Antiviral activity of RPV (**1**) (A) and PEG4-Ad (**26**) (B) against NL4-3 LucR IMCs encoding RTs with clinically relevant NNRTI resistance mutations in the TZMbl assay. N = 2 independent biological replicates.

## 4. Discussion

Here, we report our investigation of rilpivirine-based TPDs against HIV-1 reverse transcriptase (RT) using either E3 ligase–recruiting PROTACs or a hydrophobic tag (HyT) strategy. Among the compounds evaluated, the HyT molecule PEG4-Ad (**26**) displayed the most promising profile, exhibiting a low nanomolar IC₅₀ against the HIV-1 reference strain NL4-3, and outperforming its PROTAC counterparts in the TZMbl single round infection assay. PEG4-Ad (**26**) also inhibited viral spread across multiple rounds of replication in a CD4+ T cell line, demonstrating activity in a more physiologically relevant infection model. Mechanistic studies supported a degradation-based mode of action, as PEG4-Ad (**26**) induced loss of RT/p66-HiBiT in our p51-P2A-p66.HiBiT LVV expression system in a proteasome-dependent manner. This effect was abrogated by the proteasome inhibitor MG-132, supporting that the HyT approach successfully recruits the cellular protein quality control (PQC) machinery to RT. MD simulations also showed that over the equilibrated 20 to 50 ns interval, PEG4-Ad (**26**) exposed its adamantane to solvent in >12% of frames, roughly four-fold more than PEG2PM5-Ad (**31**) or PEG5-Ad (**27**) (**Figure 12A, B**). Given that hydrophobic tagging depends on presentation of the exposed hydrophobic moiety to the protein quality-control machinery, the sustained solvent exposure, visualized in the medoid overlay (**Figure 13**), provides a plausible basis for PEG4-Ad’s activity in our degradation assay. Given that there are multiple reports implicating HSP70 as the chaperone that recognizes HyT destabilized proteins to mediate target polyubiquitination and degradation [18, 60, 69, 70], we also investigated whether recruitment of the PQC machinery by PEG4-Ad (**26**) was dependent on HSP70. Upregulation of HSP70 by treatment with 17-AAG of RT/p51-p66.HiBiT expressing cells did not lead to a significant increase in PEG4-Ad (**26**)-mediated RT/p66.HiBiT degradation. HSP70-mediated recognition, however, is not the only mechanism by which HyTs can engage the PQC [69, 71, 72]. HSP70-independent protein degradation has also been observed in the development of an HBV core protein HyT degrader by Xu *et al.*, who found that 2 µM 17-AAG did not enhance the degradation effect of their NVR 3-778-derived degrader. Furthermore, Xu *et al.* determined that the autophagy-lysosome pathway, rather than the proteasome, was responsible for degradation of the target protein [59]. Alternate mechanisms for HSP70-independent, proteasome-dependent degradation may involve recognition of an artificially exposed hydrophobic surface by the BAG6-UBL4A-TRC35 quality-control complex, followed by RNF126-mediated ubiquitination [63[73, 74]. It is also conceivable that PEG4-Ad (**26**)-induced local destabilization of RT could facilitate direct engagement by the E3 ligase CHIP (STUB1), which possesses intrinsic chaperone-like activity and has been reported to bind non-native protein conformations, thereby promoting ubiquitination independently of classical HSP70-mediated substrate recognition [75]. Alternatively, although less directly supported for adamantyl HyTs, PEG4-Ad (**26**) binding could induce sufficient local unfolding to permit ubiquitin-independent degradation by the 20S proteasome, which is capable of directly degrading structurally disordered proteins in the absence of ubiquitination [76]. This pathway for degradation would be independent of chaperone recognition and would still be sensitive to inhibition by MG132. Future studies exploring the ubiquitination of RT and identifying binding partners in the presence of PEG4-Ad (**26**) would assist in clarifying the specific pathways utilized for PEG4-Ad (**26**)-mediated degradation.

Molecular dynamics simulations provided further insight into the structure–activity relationships within the HyT series. Specifically, PEG4-Ad compound (**26**) showed superior performance in computational simulations because the PEG4 linker length enabled episodic, productive exposure of the adamantane tag above PQC recognition thresholds, whereas shorter linkers lacked sufficient reach and longer linkers appeared prone to self-masking conformations, reducing tag accessibility (**Figure 12A, B; Figure 13**).

An unexpected observation was our inability to detect the p51 subunit in HEK293 cells stably transduced with the dox-inducible p51–P2A-p66–HiBiT LVV construct. Of note, prior NMR and small angle X-ray scattering (SAXS) studies have shown that the unliganded p51 monomer is intrinsically less stable than p66 due to a less compact fingers-to-thumb arrangement [77]. Moreover, in addition to its heterodimeric form, p66 can exist in both monomeric and homodimeric forms, and both species retain the ability to bind NNRTIs [77, 78]. It is possible that under the low protein expression level conditions optimized for the degradation assay p51 is more unstable – and harder to detect – despite the equimolar translation of p51 and p66.HiBiT. Given that the p66 homodimer more closely reflects the conformation of RT within the GagPol precursor prior to the proteolytic event that generates p51, our findings are consistent with a predominantly p66 like population in our cellular model system. Taken together, these considerations raise the possibility that PEG4-Ad (**26**) may engage RT in its precursor forms during the assembly stage late in the replication cycle, when RT is accessible prior to viral budding and maturation, thereby enabling HyT-mediated degradation [79–81]. Further experimental work is needed to determine the extent to which precursor engagement contributes to the antiviral activity of PEG4-Ad (**26**).

Despite significant PEG4-Ad (**26**) activities in the single and multi-round infectivity and degradation assays, PEG4-Ad (**26**) did not show improved antiviral potency over RPV against NL4-3 mutants carrying clinically relevant NNRTI resistance mutations. Although TPDs often outperform their parent inhibitors against resistant targets in other systems— particularly in cancer [65, 82] but also observed for HBV core protein and oseltamivir resistant IAV—HIV imposes unique structural constraints. Reverse transcription does not occur in the cytosol as originally proposed [83–85]. Instead, recent studies [86–88] demonstrate that the HIV capsid remains largely intact after fusion, traffics to the nuclear pore via FG nucleoporin interactions, and undergoes major uncoating only after nuclear entry, with reverse transcription proceeding inside the intact or partially uncoated core [89–91]. This encapsidated environment protects viral nucleic acids but also physically blocks access by the cellular PQC machinery.

Our data indicate that PEG4-Ad (**26**) can reach RT within this environment— consistent with its significant inhibition during the early stages of the HIV replication cycle—but PQC components required for HyT-mediated degradation cannot pass through the ∼8 Å pores of the native capsid lattice. This steric exclusion, together with the reduced affinity of PEG4-Ad (**26**) relative to RPV, likely explains the absence of enhanced antiviral activity relative to RPV. Nevertheless, PEG4-Ad activity in our degradation assay demonstrates that degradation of an exposed RT remains feasible, either in the context of a prematurely disassembled capsid or during viral assembly. The latter window of opportunity includes expression and assembly of GagPol during which p66 within the precursor adopts a p66/p51-like dimer, and RT driven dimerization of the precursor helps position and activate PR—a state in which RT is structurally exposed and ligand addressable (e.g., NNRTI binding enhances RT dimerization and modulates GagPol processing) [92]. Because the capsid has not yet formed at this stage, HyT engagement of the RT domain within GagPol or its derivatives resulting from the onset of PR-driven proteolytic maturation during budding but prior to virion release [79, 93] could recruit PQC machinery and drive precursor degradation prior to virion maturation. These alternative pre-maturation contexts illustrate viable mechanistic opportunities for future degrader optimization.

The generally superior performance of HyT compounds over PROTACs, despite the latter’s favorable ternary complex geometries predicted by modeling, may reflect a mechanistic advantage of HyTs over PROTACs. Since HyTs do not feature a second ligand that can facilitate the production of binary complexes between the PROTAC and POI or the PROTAC and E3 ligase (i.e., the hook effect), they prevent the sequestration of the TPD by the E3 ligase, which exists in much higher concentrations in the infected cell compared to RT. This mechanistic advantage would leave the entire intracellular pool of HyTs available to bind RT and inhibit its function leading to the superior performance observed in the TZMbl assays.

Several questions raised by this work merit further investigation. Clarifying the precise ubiquitination status of RT in the presence of PEG4-Ad (**26**) and identifying its binding partners under these conditions will be essential to delineating the specific PQC pathway engaged. The unexpected absence of detectable p51 in our expression system warrants follow-up, as it supports the intriguing possibility that PEG4-Ad (**26**) could engage RT within GagPol precursor forms during late-stage replication, a window of opportunity in which RT is structurally exposed and the capsid has not yet assembled, rendering PQC machinery accessible. Exploiting these pre-maturation contexts, alongside strategies to improve RT binding affinity relative to the parent RPV scaffold, represent the most promising avenues for next-generation RT-targeting degrader design. More broadly, the potential capsid access barrier may be a general constraint applicable to any RT-directed degrader and circumventing it may be a central challenge for the field going forward.

## Supporting Information

The following supporting information can be downloaded at: https://www.mdpi.com/article/doi/s1, Figure S1: title; Table S1: title; Video S1: title.

## Author Contributions

“Conceptualization, C.O., T.S.S., S.C.S., R.C.R and J.C.K.; methodology, F.A., G.D.C., R.K., A.F.H., D.S.M. ZY, K.G., C.O., J.C.K, T.S.S, and S.C.S; validation, F.A., G.D.C., R.K., A.F.H., D.S.M, C.O., T.S.S, S.C.S; formal analysis, F.A., G.D.C., R.K., A.F.H., D.S.M., K.G., Z.Y.; investigation, F.A., G.D.C., R.K., A.F.H., D.S.M., C.B., Z.Y., K.G., S.C.S.; resources, C.O., T.S.S., J.C.K., Z.Y., S.C.S.; writing—original draft preparation, F.A., R.K., G.D.C., T.S.S., A.F.H, D.S.M.; writing—review and editing, C.O., T.S.S., J.C.K, R.C.R, S.C.S.; visualization, F.A., R.K., T.S.S., G.D.C., A.F.H., D.S.M.; supervision, C.O., T.S.S., S.C.S., J.C.K.; project administration, R.C.R, C.O., T.S.S.; S.C.S.; funding acquisition, R.C.R., T.S.S, C.O., S.C.S., J.C.K. All authors have read and agreed to the published version of the manuscript.

## Funding

This work was supported by the National Institutes of Health (NIH) National Institute of Allergy and Infectious Diseases (NIAID), R21AI157362 (A New Paradigm for HIV Treatment: Targeted Degradation of HIV Reverse Transcriptase via the Ubiquitin-Proteasome Pathway) to RCR, and in part by the State of Florida Biomedical Research Program, Bankhead Coley Research Infrastructure grant 23B16.

## Data Availability Statement

The original contributions presented in this study are included in the article/supplementary material. Further inquiries can be directed to the corresponding author(s).

## Supporting information

Supplementary NMR Spectra File

Graphical Abstract

## Acknowledgments

SCS and RK thank the Sylvester Comprehensive Cancer Center for its support, and the Frost Institute for Data Science & Computing (IDSC), particularly Pedro Davila, for his technical expertise in maintaining servers and resolving Docker-related issues; we also thank ALAFIA.AI for providing our group with a desktop supercomputer and a dedicated server; and we are grateful to ChemAxon and OpenEye for providing access to advanced chemical informatics and computational chemistry software.

FKA, JCK and CO would like to thank Dr. Stefan G. Sarafianos for generously gifting us purified RT for our thermal stability assays. CO, JCK, and FKA thank Jennifer Jones for providing expertise and training in mammalian cell and HIV-1 culture to FKA; we also want to acknowledge Jie Zeng in grateful memory for her contributions of excellence in molecular cloning techniques to our laboratory.

TSS, AFH, and KG thank Dr. Marco Bonizzoni for assistance with the kinetic solubility studies and use of analytical equipment.

## Conflicts of Interest

The authors have no conflicts of interest to declare.

## Abbreviations

The following abbreviations are used in this manuscript:

ART: Antiretroviral Therapy
CHIP: Carboxy-terminus of Hsc70-Interacting Protein
CRBN: Cereblon
HIV-1: Human Immunodeficiency Virus-1
HSP70: Heat Shock Protein 70
HyT: Hydrophobic Tag
IDV: Indinavir
IMC: Infectious Molecular Clone
MD: Molecular Dynamics
MST: Microscale Thermophoresis
NNRTI: Non-Nucleoside Reverse Transcriptase Inhibitor
NRTI: Nucleoside Reverse Transcriptase Inhibitor
POI: Protein of Interest
PROTAC: Proteolysis Targeting Chimera
PWLH: People Living with HIV
RPV: Rilpivirine
RPV’: 1,3,5-triazine analog of Rilpivirine
RT: Reverse Transcriptase
snLUC: Secreted Nano-Luciferase
TPD: Targeted Protein Degrader
VHL: von Hippel-Lindau

## Disclaimer/Publisher’s Note

The statements, opinions and data contained in all publications are solely those of the individual author(s) and contributor(s) and not of MDPI and/or the editor(s). MDPI and/or the editor(s) disclaim responsibility for any injury to people or property resulting from any ideas, methods, instructions or products referred to in the content.

## Notes

### Competing Interest Statement

The authors have declared no competing interest.

## References

1. Asahchop EL, Wainberg MA, Sloan RD, Tremblay CL. Antiviral drug resistance and the need for development of new HIV-1 reverse transcriptase inhibitors. Antimicrob Agents Chemother. 2012;56(10):5000–8.

2. Saag MS, Gandhi RT, Hoy JF, Landovitz RJ, Thompson MA, Sax PE, et al. Antiretroviral Drugs for Treatment and Prevention of HIV Infection in Adults: 2020 Recommendations of the International Antiviral Society-USA Panel. JAMA. 2020;324(16):1651–69.

3. Das K, Arnold E. HIV-1 reverse transcriptase and antiviral drug resistance. Part 1. Curr Opin Virol. 2013;3(2):111–8.

4. Smith SJ, Pauly GT, Akram A, Melody K, Rai G, Maloney DJ, et al. Rilpivirine analogs potently inhibit drug-resistant HIV-1 mutants. Retrovirology. 2016;13:11.

5. Smith SJ, Pauly GT, Akram A, Melody K, Ambrose Z, Schneider JP, et al. Rilpivirine and Doravirine Have Complementary Efficacies Against NNRTI-Resistant HIV-1 Mutants. Jaids-J Acq Imm Def. 2016;72(5):485–91.

6. Das K, Arnold E. HIV-1 reverse transcriptase and antiviral drug resistance. Part 2. Curr Opin Virol. 2013;3(2):119–28.

7. Melikian GL, Rhee SY, Varghese V, Porter D, White K, Taylor J, et al. Non-nucleoside reverse transcriptase inhibitor (NNRTI) cross-resistance: implications for preclinical evaluation of novel NNRTIs and clinical genotypic resistance testing. J Antimicrob Chemother. 2014;69(1):12–20.

8. Das K, Bauman JD, Clark AD, Jr., Frenkel YV, Lewi PJ, Shatkin AJ, et al. High-resolution structures of HIV-1 reverse transcriptase/TMC278 complexes: strategic flexibility explains potency against resistance mutations. Proc Natl Acad Sci U S A. 2008;105(5):1466–71.

9. Wang Z, Rumrill S, Kang D, Guma SD, Feng D, De Clercq E, et al. Development of enhanced HIV-1 non-nucleoside reverse transcriptase inhibitors with improved resistance and pharmacokinetic profiles. Sci Adv. 2025;11(22):eadt8916.

10. Kumar A, Ali S, Shen Q, Zhou J. FDA Approval of the First-Ever PROTAC: Vepdegestrant (ARV-471) Marks a New Era in Targeted Protein Degradation. J Med Chem. 2026;69(14):16135–42.

11. Sakamoto KM, Kim KB, Kumagai A, Mercurio F, Crews CM, Deshaies RJ. Protacs: chimeric molecules that target proteins to the Skp1-Cullin-F box complex for ubiquitination and degradation. Proc Natl Acad Sci U S A. 2001;98(15):8554–9.

12. Faryal B, Ul Abideen Z, Irfan M, Ahmed H, Jalilov F, Abduraximova L, et al. Targeted Protein Degradation in Cancer: PROTACs, New Targets, and Clinical Mechanisms. Biomolecules. 2026;16(2).

13. Deshaies RJ. Protein degradation: Prime time for PROTACs. Nat Chem Biol. 2015;11(9):634–5.

14. Knoll N, Neamati N, Uren A. Mechanisms and Design Principles of Proteolysis-Targeting Chimeras and Their Emerging Applications. ACS Pharmacol Transl Sci. 2026;9(4):784–814.

15. Zhao L, Zhao J, Zhong KH, Tong AP, Jia D. Targeted protein degradation: mechanisms, strategies and application. Signal Transduct Tar. 2022;7(1).

16. Pan YC, Wang YJ, Gou SH. Proteolysis targeting chimera, molecular glue degrader and hydrophobic tag tethering degrader for targeted protein degradation: Mechanisms, strategies and application. Bioorg Chem. 2025;161.

17. Neklesa TK, Tae HS, Schneekloth AR, Stulberg MJ, Corson TW, Sundberg TB, et al. Small-molecule hydrophobic tagging-induced degradation of HaloTag fusion proteins. Nat Chem Biol. 2011;7(8):538–43.

18. Gustafson JL, Neklesa TK, Cox CS, Roth AG, Buckley DL, Tae HS, et al. Small-Molecule-Mediated Degradation of the Androgen Receptor through Hydrophobic Tagging. Angew Chem Int Edit. 2015;54(33):9659–62.

19. Bondeson DP, Mares A, Smith IE, Ko E, Campos S, Miah AH, et al. Catalytic in vivo protein knockdown by small-molecule PROTACs. Nat Chem Biol. 2015;11(8):611–7.

20. Alabi S, Jaime-Figueroa S, Yao Z, Gao YJ, Hines J, Samarasinghe KTG, et al. Mutant-selective degradation by BRAF-targeting PROTACs. Nature Communications. 2021;12(1).

21. Kostic M, Jones LH. Critical Assessment of Targeted Protein Degradation as a Research Tool and Pharmacological Modality. Trends Pharmacol Sci. 2020;41(5):305–17.

22. Bondeson DP, Smith BE, Burslem GM, Buhimschi AD, Hines J, Jaime-Figueroa S, et al. Lessons in PROTAC Design from Selective Degradation with a Promiscuous Warhead. Cell Chem Biol. 2018;25(1):78–87 e5.

23. Li YQ, Lannigan WG, Davoodi S, Daryaee F, Corrionero A, Alfonso P, et al. Discovery of Novel Bruton’s Tyrosine Kinase PROTACs with Enhanced Selectivity and Cellular Efficacy. J Med Chem. 2023;66(11):7454–74.

24. Pike A, Lee ECY, Michaelides IN, Schade M, Sharma A, Scott JS, et al. Lessons learned in linking PROTACs from discovery to the clinic. Nat Rev Chem. 2026;10(2):117–32.

25. de Wispelaere M, Du G, Donovan KA, Zhang T, Eleuteri NA, Yuan JC, et al. Small molecule degraders of the hepatitis C virus protease reduce susceptibility to resistance mutations. Nat Commun. 2019;10(1):3468.

26. Xu Z, Liu X, Ma X, Zou W, Chen Q, Chen F, et al. Discovery of oseltamivir-based novel PROTACs as degraders targeting neuraminidase to combat H1N1 influenza virus. Cell Insight. 2022;1(3):100030.

27. Alugubelli YR, Xiao J, Khatua K, Kumar S, Sun L, Ma YY, et al. Discovery of First-in-Class PROTAC Degraders of SARS-CoV-2 Main Protease. J Med Chem. 2024;67(8):6495–507.

28. Emert-Sedlak LA, Tice CM, Shi HB, Alvarado JJ, Shu ST, Reitz AB, et al. PROTAC-mediated degradation of HIV-1 Nef efficiently restores cell-surface CD4 and MHC-I expression and blocks HIV-1 replication. Cell Chemical Biology. 2024;31(4).

29. Luo D, Luo RH, Wang WL, Deng R, Wang SR, Ma XY, et al. Discovery of L15 as a novel Vif PROTAC degrader with antiviral activity against HIV-1. Bioorg Med Chem Lett. 2024;111.

30. Newton LS, Gathmann C, Ridewood S, Smith RJ, Wijaya AJ, Hornsby TW, et al. Macrocycle-based PROTACs selectively degrade cyclophilin A and inhibit HIV-1 and HCV. Nature Communications. 2025;16(1).

31. Friesner RA, Murphy RB, Zhang YQ, Xiong YY, Devlaminck PA, Tubert-Brohman I, et al. Glide WS: Methodology and Initial Assessment of Performance for Docking Accuracy and Virtual Screening. J Chem Theory Comput. 2025;21(24):12696–708.

32. Ge JX, Li SM, Weng GQ, Wang HT, Fang MJ, Sun HY, et al. PROTAC-DB 3.0: an updated database of PROTACs with extended pharmacokinetic parameters. Nucleic Acids Res. 2024;53(D1):D1510–D5.

33. van Zundert GCP, Rodrigues JPGLM, Trellet M, Schmitz C, Kastritis PL, Karaca E, et al. The HADDOCK2.2 Web Server: User-Friendly Integrative Modeling of Biomolecular Complexes. J Mol Biol. 2016;428(4):720–5.

34. Jofily P, Kalyaanamoorthy S. P4ward: An Automated Modeling Platform for Protac Ternary Complexes. J Chem Inf Model. 2025;65(16):8806–18.

35. Kevin J., Bowers EC, Huafeng Xu, Ron O. Dror, Michael P. Eastwood, Brent A. Gregersen, John L. Klepeis, Istvan Kolossvary, Mark A. Moraes, Federico D. Sacerdoti, John K. Salmon, Yibing Shan, David E. Shaw Scalable algorithms for molecular dynamics simulations on commodity clusters. Proceedings of the 2006 ACM/IEEE conference on Supercomputing. 2006:84–es.

36. Lu C, Wu CJ, Ghoreishi D, Chen W, Wang LL, Damm W, et al. OPLS4: Improving Force Field Accuracy on Challenging Regimes of Chemical Space. J Chem Theory Comput. 2021;17(7):4291–300.

37. Soto-Martinez DM, Clements GD, Díaz JE, Becher J, Reynolds RC, Ochsenbauer C, et al. Preparation of von Hippel-Lindau (VHL) E3 ubiquitin ligase ligands exploiting constitutive hydroxyproline for benzylic amine protection. Rsc Adv. 2024;14(24):17077–90.

38. Roche. Reverse Transcriptase Assay, colorimetric. 2020.

39. Alberti MO, Jones JJ, Miglietta R, Ding H, Bakshi RK, Edmonds TG, et al. Optimized Replicating Renilla Luciferase Reporter HIV-1 Utilizing Novel Internal Ribosome Entry Site Elements for Native Nef Expression and Function. AIDS Res Hum Retroviruses. 2015;31(12):1278–96.

40. Wei X, Decker JM, Liu H, Zhang Z, Arani RB, Kilby JM, et al. Emergence of Resistant Human Immunodeficiency Virus Type 1 in Patients Receiving Fusion Inhibitor (T-20) Monotherapy. Antimicrobial Agents and Chemotherapy. 2002;46(6):1896–905.

41. Sarzotti-Kelsoe M, Bailer RT, Turk E, Lin CL, Bilska M, Greene KM, et al. Optimization and validation of the TZM-bl assay for standardized assessments of neutralizing antibodies against HIV-1. J Immunol Methods. 2014;409:131–46.

42. Harris JR, Kitchen AD, Harrison JF, Tovey G. Viral release from HIV-I-induced syncytia of CD4+ C8166 cells. J Med Virol. 1989;28(2):81–9.

43. Krowicka H, Robinson JE, Clark R, Hager S, Broyles S, Pincus SH. Use of tissue culture cell lines to evaluate HIV antiviral resistance. AIDS Res Hum Retroviruses. 2008;24(7):957–67.

44. Michailidis E, Marchand B, Kodama EN, Singh K, Matsuoka M, Kirby KA, et al. Mechanism of inhibition of HIV-1 reverse transcriptase by 4’-Ethynyl-2-fluoro-2’-deoxyadenosine triphosphate, a translocation-defective reverse transcriptase inhibitor. J Biol Chem. 2009;284(51):35681–91.

45. Liu Z, Chen O, Wall JBJ, Zheng M, Zhou Y, Wang L, et al. Systematic comparison of 2A peptides for cloning multi-genes in a polycistronic vector. Sci Rep. 2017;7(1):2193.

46. Bai N, Riching KM, Makaju A, Wu H, Acker TM, Ou SC, et al. Modeling the CRL4A ligase complex to predict target protein ubiquitination induced by cereblon-recruiting PROTACs. Journal of Biological Chemistry. 2022;298(4).

47. Zhang S, Anang S, Zhang Z, Nguyen HT, Ding H, Kappes JC, et al. Conformations of membrane human immunodeficiency virus (HIV-1) envelope glycoproteins solubilized in Amphipol A18 lipid-nanodiscs. J Virol. 2024;98(10):e0063124.

48. Kerns EH, Di L, Carter GT. Solubility Assays in Drug Discovery. Curr Drug Metab. 2008;9(9):879–85.

49. Venturi A, Di Bona S, Desantis J, Eleuteri M, Bartalucci M, Baroni M, et al. Between Theory and Practice: Computational/Experimental Integrated Approaches to Understand the Solubility and Lipophilicity of PROTACs. J Med Chem. 2024;67(18):16355–80.

50. Janssen PA, Lewi PJ, Arnold E, Daeyaert F, de Jonge M, Heeres J, et al. In search of a novel anti-HIV drug: multidisciplinary coordination in the discovery of 4-[[4-[[4-[(1E)-2-cyanoethenyl]-2,6-dimethylphenyl]amino]-2-pyrimidinyl]amino]benzonitrile (R278474, rilpivirine). J Med Chem. 2005;48(6):1901–9.

51. Bollini M, Cisneros JA, Spasov KA, Anderson KS, Jorgensen WL. Optimization of diarylazines as anti-HIV agents with dramatically enhanced solubility. Bioorg Med Chem Lett. 2013;23(18):5213–6.

52. Adachi A, Gendelman HE, Koenig S, Folks T, Willey R, Rabson A, et al. Production of acquired immunodeficiency syndrome-associated retrovirus in human and nonhuman cells transfected with an infectious molecular clone. J Virol. 1986;59(2):284–91.

53. Edmonds TG, Ding H, Yuan X, Wei Q, Smith KS, Conway JA, et al. Replication competent molecular clones of HIV-1 expressing Renilla luciferase facilitate the analysis of antibody inhibition in PBMC. Virology. 2010;408(1):1–13.

54. Watson ER, Novick S, Matyskiela ME, Chamberlain PP, de la Pena AH, Zhu JY, et al. Molecular glue CELMoD compounds are regulators of cereblon conformation. Science. 2022;378(6619):549–53.

55. Lopez-Girona A, Mendy D, Ito T, Miller K, Gandhi AK, Kang J, et al. Cereblon is a direct protein target for immunomodulatory and antiproliferative activities of lenalidomide and pomalidomide. Leukemia. 2012;26(11):2326–35.

56. Fischer ES, Bohm K, Lydeard JR, Yang H, Stadler MB, Cavadini S, et al. Structure of the DDB1-CRBN E3 ubiquitin ligase in complex with thalidomide. Nature. 2014;512(7512):49–53.

57. Frost J, Rocha S, Ciulli A. Von Hippel-Lindau (VHL) small-molecule inhibitor binding increases stability and intracellular levels of VHL protein. J Biol Chem. 2021;297(2):100910.

58. Diehl CJ, Ciulli A. Discovery of small molecule ligands for the von Hippel-Lindau (VHL) E3 ligase and their use as inhibitors and PROTAC degraders. Chem Soc Rev. 2022;51(19):8216–57.

59. Shoda T, Ohoka N, Tsuji G, Fujisato T, Inoue H, Demizu Y, et al. Targeted Protein Degradation by Chimeric Compounds using Hydrophobic E3 Ligands and Adamantane Moiety. Pharmaceuticals-Base. 2020;13(3).

60. Choi SR, Wang HM, Shin MH, Lim HS. Hydrophobic Tagging-Mediated Degradation of Transcription Coactivator SRC-1. Int J Mol Sci. 2021;22(12).

61. Raina K, Lu J, Qian Y, Altieri M, Gordon D, Rossi AM, et al. PROTAC-induced BET protein degradation as a therapy for castration-resistant prostate cancer. Proc Natl Acad Sci U S A. 2016;113(26):7124–9.

62. Hartl FU, Bracher A, Hayer-Hartl M. Molecular chaperones in protein folding and proteostasis. Nature. 2011;475(7356):324–32.

63. Robins T, Plattner J. HIV protease inhibitors: their anti-HIV activity and potential role in treatment. J Acquir Immune Defic Syndr (1988). 1993;6(2):162–70.

64. Neklesa TK, Noblin DJ, Kuzin AP, Lew S, Seetharaman J, Acton TB, et al. A bidirectional system for the dynamic small molecule control of intracellular fusion proteins. ACS Chem Biol. 2013;8(10):2293–300.

65. Buhimschi AD, Armstrong HA, Toure M, Jaime-Figueroa S, Chen TL, Lehman AM, et al. Targeting the C481S Ibrutinib-Resistance Mutation in Bruton’s Tyrosine Kinase Using PROTAC-Mediated Degradation. Biochemistry. 2018;57(26):3564–75.

66. Guo M, He S, Cheng J, Li Y, Dong G, Sheng C. Hydrophobic Tagging-Induced Degradation of PDEdelta in Colon Cancer Cells. ACS Med Chem Lett. 2022;13(2):298–303.

67. Xu SJ, Wang Y, Shi DZ, Wang S, Qiao LJ, Yang G, et al. Discovery and mechanism verification of first-in-class hydrophobic tagging-based degraders of HBV core protein. Acta Pharm Sin B. 2025;15(4):2170–96.

68. Liu YQ, Li HB, Liang DZ, Chen YG, Lu KY, Tao HQ, et al. Harnessing hydrophobic tag technology to combat drug-resistant influenza: design, synthesis and potency of oseltamivir-derived HyTTDs. New J Chem. 2025;49(13):5489–504.

69. Xie SW, Zhan FY, Zhu JJ, Sun Y, Zhu HJ, Liu J, et al. Discovery of Norbornene as a Novel Hydrophobic Tag Applied in Protein Degradation. Angew Chem Int Edit. 2023;62(13).

70. Wang MY, Lin RK, Li JC, Suo YY, Gao J, Liu LP, et al. Discovery of LL-K8-22: A Selective, Durable, and Small-Molecule Degrader of the CDK8-Cyclin C Complex. J Med Chem. 2023;66(7):4932–51.

71. Balestrero FC, Gioiello L, Goutsiou G, Sangaletti S, Di Martino RMC, Condorelli F, et al. Versatile One-Pot Synthesis of Hydrophobic Tags by Multicomponent Reactions. ACS Omega. 2025;10(5):4745–53.

72. He Q, Zhao X, Wu D, Jia S, Liu C, Cheng Z, et al. Hydrophobic tag-based protein degradation: Development, opportunity and challenge. Eur J Med Chem. 2023;260:115741.

73. Line Pedersen JD, Kristoffer E. Johansson, Celeste M. Hackney, Isa K. Henrichs, Erna S. Sigmarsdóttir, Martin Grønbæk-Thygesen, Vasileios Voutsinos, Kresten Lindorff-Larsen, Rasmus Hartmann-Petersen. BAG6 and RNF126 are broadly involved in protein quality control of non-native missense protein variants. BioRxiv. 2026.

74. Krysztofinska EM, Martinez-Lumbreras S, Thapaliya A, Evans NJ, High S, Isaacson RL. Structural and functional insights into the E3 ligase, RNF126. Sci Rep. 2016;6:26433.

75. Rosser MF, Washburn E, Muchowski PJ, Patterson C, Cyr DM. Chaperone functions of the E3 ubiquitin ligase CHIP. J Biol Chem. 2007;282(31):22267–77.

76. Ben-Nissan G, Sharon M. Regulating the 20S proteasome ubiquitin-independent degradation pathway. Biomolecules. 2014;4(3):862–84.

77. Seetaha S, Kamonsutthipaijit N, Yagi-Utsumi M, Seako Y, Yamaguchi T, Hannongbua S, et al. Biophysical Characterization of p51 and p66 Monomers of HIV-1 Reverse Transcriptase with Their Inhibitors. Protein J. 2023;42(6):741–52.

78. Sharaf NG, Xi ZY, Ishima R, Gronenborn AM. The HIV-1 p66 homodimeric RT exhibits different conformations in the binding-competent and –incompetent NNRTI site. Proteins. 2017;85(12):2191–7.

79. Kleinpeter AB, Freed EO. Mechanisms of HIV-1 assembly, release and maturation. Nat Rev Microbiol. 2026.

80. Harrison JJEK, Passos DO, Bruhn JF, Bauman JD, Tuberty L, DeStefano JJ, et al. Cryo-EM structure of the HIV-1 Pol polyprotein provides insights into virion maturation. Sci Adv. 2022;8(27).

81. Balibar CJ, Klein DJ, Zamlynny B, Diamond TL, Fang Z, Cheney CA, et al. Potent targeted activator of cell kill molecules eliminate cells expressing HIV-1. Sci Transl Med. 2023;15(684):eabn2038.

82. Burke MR, Smith AR, Zheng G. Overcoming Cancer Drug Resistance Utilizing PROTAC Technology. Front Cell Dev Biol. 2022;10:872729.

83. Hu WS, Hughes SH. HIV-1 Reverse Transcription. Csh Perspect Med. 2012;2(10).

84. Bukrinsky M. A hard way to the nucleus. Mol Med. 2004;10(1-6):1–5.

85. Mamede JI, Cianci GC, Anderson MR, Hope TJ. Early cytoplasmic uncoating is associated with infectivity of HIV-1. P Natl Acad Sci USA. 2017;114(34):E7169–E78.

86. Müller TG, Zila V, Muller B, Kräusslich HG. Nuclear Capsid Uncoating and Reverse Transcription of HIV-1. Annu Rev Virol. 2022;9:261–84.

87. Dharan A, Bachmann N, Talley S, Zwikelmaier V, Campbell EM. Nuclear pore blockade reveals that HIV-1 completes reverse transcription and uncoating in the nucleus. Nat Microbiol. 2020;5(9):1088–+.

88. Selyutina A, Persaud M, Lee K, KewalRamani V, Diaz-Griffero F. Nuclear Import of the HIV-1 Core Precedes Reverse Transcription and Uncoating. Cell Rep. 2020;32(13).

89. Dickson CF, Hertel S, Tuckwell AJ, Li N, Ruan J, Al-Izzi SC, et al. The HIV capsid mimics karyopherin engagement of FG-nucleoporins. Nature. 2024;626(8000).

90. Fu LR, Weiskopf EN, Akkermans O, Swanson NA, Cheng SY, Schwartz TU, et al. HIV-1 capsids enter the FG phase of nuclear pores like a transport receptor. Nature. 2024;626(8000).

91. Christensen DE, Ganser-Pornillos BK, Johnson JS, Pornillos O, Sundquist WI. Reconstitution and visualization of HIV-1 capsid-dependent replication and integration in vitro. Science. 2020;370(6513).

92. Checkley MA, Luttge BG, Freed EO. HIV-1 Envelope Glycoprotein Biosynthesis, Trafficking, and Incorporation. J Mol Biol. 2011;410(4):582–608.

93. Tabler CO, Wegman SJ, Chen J, Shroff H, Alhusaini N, Tilton JC. The HIV-1 Viral Protease Is Activated during Assembly and Budding Prior to Particle Release. J Virol. 2022;96(9):e0219821.

