## Supplementary Information_Figures and Tables for "Performance of Rilpivirine-Based Hydrophobic Tags and PROTACs Directed Against HIV-1 Reverse Transcriptase"

**Abbreviations:**

|  |  |
| --- | --- |
| ACN | acetonitrile |
| BAIB | bis(acetoxiodo)benzene |
| Barton's base | 2-*tert*-butyl-1,1,3,3-tetramethylguanidine |
| *n*-BuLi | *n*-butyl lithium |
| CDT | 1,1'-carbonyl-di-(1,2,4-triazole) |
| COMU | 1-[1-(cyano-2-ethoxy-2-oxoethylideneaminoxy)-dimethylamino-morpholino]-uronium hexafluorophosphate |
| DCM | dichloromethane |
| DIPEA | *N,N*-diisopropylethylamine |
| DMAP | 4-dimethylaminopyridine |
| DMF | *N,N*-dimethylformamide |
| DMSO | dimethyl sulfoxide |
| EDC•HCl | 1-ethyl-3-(3-dimethylaminopropyl)carbodiimide hydrochloride |
| EtOAc | ethyl acetate |
| HATU | 1-[bis(dimethylamino)methylene]-1H-1,2,3-triazolo[4,5-b]pyridinium 3-oxide hexafluorophosphate |
| HOBt hydrate | 1-hydroxybenzotriazole hydrate |
| MeOH | methanol |
| 2-MeTHF | 2-methyltetrahydrofuran |
| MTBE | methyl *tert*-butyl ether |
| TBSCl | *tert*-butyldimethylsilyl chloride |
| TEA | triethylamine |
| TEMPO | (2,2,6,6-tetramethylpiperidin-1-yl)oxyl |
| TFA | trifluoroacetic acid |
| THF | tetrahydrofuran |
| TLC | thin-Layer Chromatography |
| TsCl | 4-toluenesulfonyl chloride |

### SECTION 1: Synthesis Routes, Methods, and Characterization Data

#### General Information

All reactants, reagents, and solvents were purchased from Sigma-Aldrich, Ambeed, Acros Organics, TCI Chemical or VWR suppliers. Tetrahydrofuran (THF) was dried by distillation from sodium benzophenone ketyl radical. Reactions were monitored by Thin-Layer chromatography (TLC) on pre-coated 0.25 mm silica gel plates (60F254) purchased from Silicycle and visualized using UV light (254 nm or 365 nm), ninhydrin, or $\text{KMnO}_4$ stain with mild charring. Flash chromatography was performed using silica gel (60 Å, 230–400 mesh) from Silicycle pre-dried in a 150 °C oven for at least 24 h with a manual column or a Teledyne ISCO Combiflash Rf 200i. $^1\text{H}$ and $^{13}\text{C}$ NMR spectra were recorded on a Bruker NEO-500 spectrometer with a cryoprobe. All reported $^1\text{H}$ NMR chemical shift values ($\delta\text{H}$) were referenced to the residual $^1\text{H}$ signal of deuterated solvents ($\text{CDCl}_3$: $^1\text{H}$ = 7.26 ppm; $(\text{CD}_3)_2\text{CO}$: $^1\text{H}$ = 2.05 ppm; $\text{CD}_3\text{OD}$: $^1\text{H}$ = 3.31 ppm; $\text{CD}_3\text{CN}$: $^1\text{H}$ = 1.94 ppm).<sup>1</sup> All reported $^{13}\text{C}$ chemical shift values ($\delta\text{C}$) were referenced to the residual $^{13}\text{C}$ signal of deuterated solvents ($\text{CDCl}_3$: $^{13}\text{C}$ = 77.16 ppm; $(\text{CD}_3)_2\text{CO}$: $^{13}\text{C}$ = 29.84 ppm; $\text{CD}_3\text{OD}$: $^{13}\text{C}$ = 49.00 ppm; $\text{CD}_3\text{CN}$: $^{13}\text{C}$ = 1.32 ppm, 118.26 ppm).<sup>1</sup> Mass spectra were recorded using a Waters Xevo G2-XS QToF with ACUITU UPLC M-Class equipped with ESI and a high-performance orthogonal-acceleration Time of Flight (oaTOF) mass analyzer (MS2). Rilpivirine and polysubstituted triazines are reported to exist as observable conformers in NMR spectra.<sup>2-3</sup>

#### General Procedures

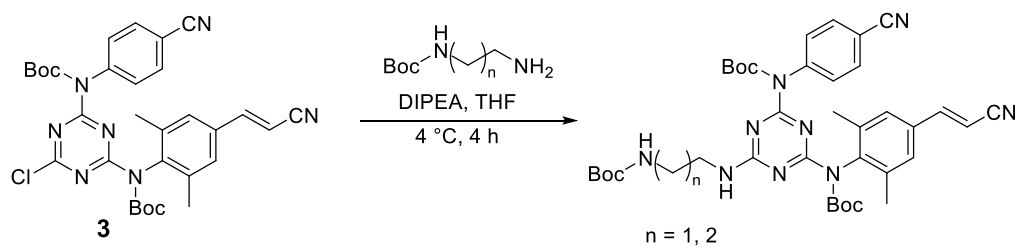

##### General Procedure I: Nucleophilic Aromatic Substitution of Disubstituted Chlorotriazines with Amines

To a flame-dried 100 mL round-bottom flask equipped with a magnetic stir bar under inert atmosphere ($\text{N}_2$) was added **3** (1.00 equiv), *tert*-butyl (2-aminoethyl)carbamate or *tert*-butyl (3-aminopropyl)carbamate (1.50 equiv) and distilled THF (0.10 M with respect to **3**). The flask was chilled in an ice bath for 10 minutes before adding DIPEA dropwise (2.0 equiv) over 5 minutes. The cold bath was maintained for 4 h, at which time the reaction was judged complete by TLC (1:1 hexanes:EtOAc). The solution was poured into ~150 mL of ice in a 250 mL Erlenmeyer flask using cold deionized water to transfer the residual. The flask was placed in an ice bath, and its contents were stirred with a spatula for 15 minutes to precipitate a solid. The suspension was filtered over a fine fritted funnel and washed with deionized water (3 x 50 mL) to remove residual THF and base. The filter cake was kept under vacuum filtration for 30 minutes forming a loose powder, then the solid was purified by flash column chromatography using a gradient of 0–40% EtOAc in hexanes. The desired materials eluted with 40% EtOAc in hexanes. The fractions were collected and concentrated by rotary evaporation at 35 °C to obtain a solid. The solid was dried overnight with stirring under reduced pressure to afford the product.

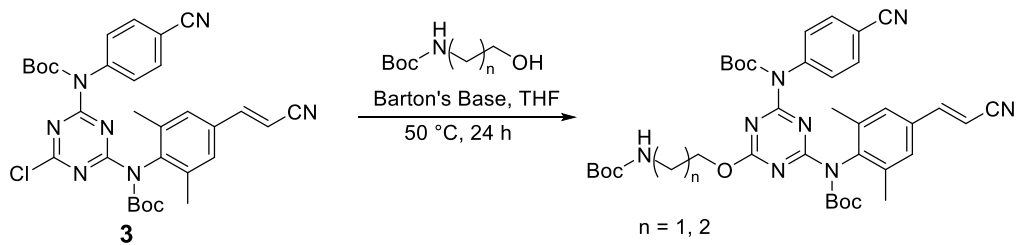

#### General Procedure II: Nucleophilic Aromatic Substitution of Disubstituted Chlorotriazines with Alcohols

To a flame-dried 100 mL round-bottom flask equipped with a magnetic stir bar under inert atmosphere ($N_2$) was added **3** (1.00 equiv), *tert*-butyl (2-hydroxyethyl)carbamate or *tert*-butyl (3-hydroxypropyl)carbamate (1.50 equiv), and distilled THF (0.10 M with respect to **3**). The flask was chilled in an ice bath for 10 minutes before Barton's base (2.00 equiv) was added dropwise over 5 minutes by syringe. The ice bath was maintained for 15 minutes then the solution was heated to 50 °C for 24 h, at which time the reaction was judged complete by TLC (1:1 hexanes:EtOAc). The flask was cooled to room temperature in a water bath then the solution was poured into ~150 mL of ice in a 250 mL Erlenmeyer flask using cold deionized water to transfer the residual. The flask was placed in an ice bath, and the contents were stirred for 15 minutes with a spatula to precipitate a solid. The suspension was filtered over a fine fritted funnel, and the solid was washed with deionized water (3 x 50 mL) to remove residual THF and base. The filter cake was kept under vacuum filtration on the fritted funnel for 30 minutes forming a loose powder. The solid was purified by column chromatography using a gradient of 0-40% EtOAc in hexanes. The desired material **4** and **S5** eluted with 40% EtOAc in hexanes. The fractions were concentrated by rotary evaporation at 35 °C to afford a solid. The solid was dried overnight with stirring under reduced pressure to afford the product.

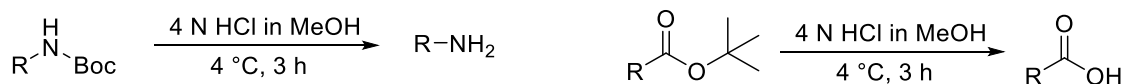

#### General Procedure III: Removal of a Boc Protecting Group or a *tert*-butyl ester via HCl to Liberate an Amine or Carboxylic Acid

To a 100 mL round-bottom flask equipped with a magnetic stir bar was added **x** (1.00 equiv **4**, **S5**, **S7**, **S8**, **S16**, **S18**, **S20**, **S21**, or **S29**) Transferring the flask to an ice bath, 6 N HCl in MeOH (0.10 M with respect to **x**) was added and the ice bath maintained for 5 minutes. After 5 minutes, the ice bath was removed and the flask was kept at room temperature for 3 h, at which time the reaction was judged complete by TLC (1:1 hexanes:EtOAc). The solution was chilled in an ice bath and aqueous sodium bicarbonate was added slowly (Note: $CO_2$ formation and solution foaming observed) to quench the HCl. Maintaining the ice bath, the pH of the solution was raised using a saturated aqueous solution of NaOH chilled with ice chips (Note: pH raised to an appropriate solution pH to bring the desired compound to an ideal protonation state for liquid:liquid extraction. In the case of carboxylic acid products, the pH was maintained at 1-2). Adding ice chips to maintain the solution temperature, the aqueous phase was extracted with EtOAc chilled in an ice bath (7 x 15 mL). The combined organic phase was dried with sodium sulfate, filtered through a fritted funnel using EtOAc, and concentrated by rotary evaporation at 35 °C to afford the product. The material was dried overnight with stirring under reduced pressure.

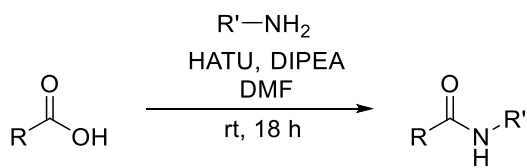

##### General Procedure IV: HATU-Mediated Amide Bond Formation

To a flame-dried 1-dram vial equipped with a magnetic stir bar under inert atmosphere ($\text{N}_2$) was added **5**, **S6**, **S9**, or **S10** (1.00 equiv), HATU (1.20 equiv) and 2-(2-(2-azidoethoxy)ethoxy)ethoxy)acetic acid (1.10 equiv), and anhydrous DMF (0.20 M with respect to **5**, **S6**, **S9**, or **S10**). Once the solution was homogenized, DIPEA (3.50 equiv) was added and the solution was maintained at room temperature for 18 h, at which time the reaction was judged complete by TLC (5% MeOH in $\text{CHCl}_3$). The solution was diluted with EtOAc (4 mL) and washed with an aqueous solution of $\text{NH}_4\text{Cl}$ (3 x 3 mL), an aqueous solution of sodium bicarbonate (2 x 3 mL), and brine (1 x 3 mL). The combined organic phase was dried with magnesium sulfate, filtered through a cotton plug using EtOAc, and concentrated by rotary evaporation at 35 °C to afford a solid. The solid was purified by preparative Thin-Layer Chromatography using 3% MeOH in $\text{CHCl}_3$. The material was desorbed from the silica by stirring the slurry in 20% MeOH in $\text{CHCl}_3$ for 18 h, filtering through a plug of Celite with $\text{CHCl}_3$, and concentrating by rotary evaporation at 25 °C. The solid was dried under reduced pressure to afford **8-11**, respectively.

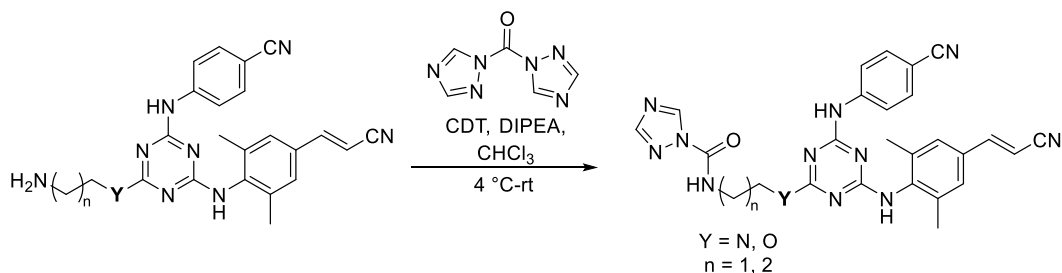

##### General Procedure V: Synthesis of Activated Carbamoyl Triazole Intermediates

To a flame-dried 1-dram vial equipped with a magnetic stir bar was added **5**, **S6**, **S9**, or **S10** (1.00 equiv) and $\text{CHCl}_3$ (0.10 M with respect to **5**, **S6**, **S9**, or **S10**) under standard atmospheric conditions. The solution was chilled in an ice bath for 5 minutes before the addition of CDT (2.00 equiv), then DIPEA (1.50 equiv) by syringe (Note: the CDT did not dissolve fully until the addition of DIPEA). The solution was gradually warmed to room temperature over 18 h, at which time the reaction was judged complete by TLC (5% MeOH in $\text{CHCl}_3$). The solution was diluted with EtOAc and washed with brine (3 x 3 mL). The combined organic phase was dried with magnesium sulfate, filtered through a cotton plug using EtOAc, and concentrated by rotary evaporation at 35 °C to afford **S12-S15**. Compounds **S12-S15** were used to generate **12-15** without further purification.

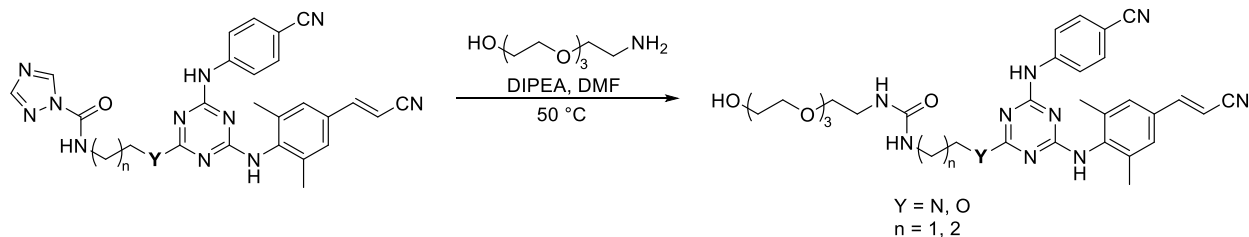

#### General Procedure VI: Urea Bond Formation via Aminolysis of Activated Carbamoyl Triazoles

To a flame-dried 1-dram vial equipped with a magnetic stir bar under inert atmosphere ($N_2$) was added **S12-S15** (1.00 equiv), 2-(2-(2-(2-aminoethoxy)ethoxy)ethoxy)ethan-1-ol (3.00 equiv), and anhydrous DMF (0.10 M with respect to **S12-S15**). Once the solution was homogeneous, DIPEA (2.00 equiv) was added by syringe and the solution was heated to 50 °C for 48 h, at which time the reaction was judged complete by TLC (5% MeOH in $CHCl_3$). The solution was cooled to room temperature before diluting with EtOAc (3 mL). The organics were washed with an aqueous solution of $NH_4Cl$ (1 x 3 mL) and brine (3 x 3 mL). The combined organic phase was dried with magnesium sulfate, filtered through a cotton plug using EtOAc, and concentrated by rotary evaporation at 35 °C to afford a solid. The solid was purified by preparative Thin-Layer Chromatography using 3% MeOH in $CHCl_3$. The material was desorbed from the silica by stirring the slurry in 20% MeOH in $CHCl_3$ for 18 h, filtering through a pad of celite with acetone, and concentrating by rotary evaporation at 25 °C to afford **12-15**, respectively.

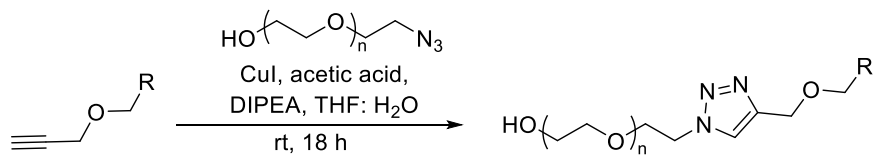

#### General Procedure VII: Formation of Triazole-Linked PEG Conjugates via Azide-Alkyne Cycloaddition

To a 10 mL round-bottom flask equipped with a magnetic stir bar was added **S17** (1.0 equiv), commercial azido-PEG<sub>n</sub>-alcohol (**a**, n = 2; **b**, n = 3;) (1.5 equiv), a catalytic amount of copper (I) iodide (0.10 equiv), a catalytic amount of glacial acetic acid (0.10 equiv), a catalytic amount of DIPEA (0.10 equiv), and a 5:1 v/v solution of THF:deionized water (0.12 M with respect to **S17** or **S23**). The reaction was doped with copper (I) iodide (3.0 mg, 0.014 mmol) at $t = 3$ h, 6 h, and 9 h. The reaction was judged complete in 18 h by TLC (1:1 hexanes:EtOAc). The reaction was quenched with saturated aqueous sodium bicarbonate (20 mL), and the aqueous phase was extracted with EtOAc (6 x 15 mL). The combined organic phase was dried with sodium sulfate, filtered through a plug of Celite using acetone, and concentrated by rotary evaporation at 30 °C to obtain a solid. The solid was purified by flash column chromatography using a gradient of 0-40% then 40-80% acetone in hexanes. The material eluted with 60-80% acetone in hexanes and the fractions were concentrated by rotary evaporation at 35 °C. The material was dried overnight with stirring under reduced pressure to afford the product.

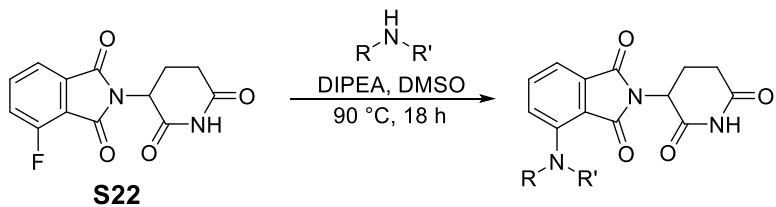

#### General Procedure VIII: Nucleophilic Aromatic Substitution of Amines into Fluorinated Pomalidomide

To a flame-dried 50 mL round-bottom flask equipped with a magnetic stir bar under inert atmosphere ($N_2$) was added 4-fluoropomalidomide (**S22**, 1.00 equiv), amino-PEG<sub>n</sub>-alcohol (1.20 equiv), anhydrous DMSO (0.10 M with respect to **S22**), and DIPEA (1.50 equiv). The solution was heated to 90 °C for 17 h, at which time the reaction was judged complete by TLC (1:1 hexanes:acetone). The solution was cooled to room temperature in a water bath for 20 minutes before the reaction was quenched with saturated aqueous sodium bicarbonate (20 mL), and the aqueous phase was extracted with EtOAc (7 x 10 mL). The combined organic phase was dried with sodium sulfate, filtered through a fritted funnel using EtOAc, and concentrated by rotary evaporation at 35 °C to afford a tacky oil. The oil was purified by flash column chromatography using a gradient of 0-45% acetone in hexanes, 30% acetone in hexanes eluted remaining starting material (**S22**), and a slow gradient up to 45% acetone in hexanes afforded the product. The fractions were concentrated by rotary evaporation at 35 °C, and the material was dried overnight under reduced pressure to afford the product.

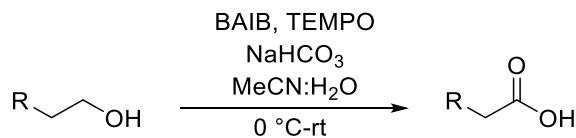

#### General Procedure IX: BAIB-Mediated Oxidation of Primary Alcohols to Carboxylic Acids

To a flame-dried 25 mL round-bottom flask equipped with a magnetic stir bar was added **x** (1.00 equiv, **x** = **S23**, **S24**, **S31**, **S36**, **S53**, **S58**), BAIB (2.30 equiv), TEMPO (0.25 equiv), sodium bicarbonate (5.50 equiv), and the solution was cooled in an ice bath. A 1:1 v/v solution of deionized water:ACN (0.10 M with respect to **x**) was added (Note: add the ACN first to assist dissolution of **S23** or **S24** before adding the deionized water). The cold bath was maintained for 4 h, at which time the reaction was judged complete by TLC (3:2 hexanes:acetone). The reaction was diluted with cold deionized water (35 mL) and extracted with $CHCl_3$ (4 x 10 mL) to selectively remove BAIB, TEMPO, ACN, and residual starting material into the organic phase. Maintaining the temperature of the aqueous phase with ice chips, the pH was lowered to 2 with aqueous 3 N HCl and extracted with EtOAc (7 x 20 mL). The combined organic phase was dried with sodium sulfate, filtered through a fritted funnel using EtOAc, and concentrated by rotary evaporation at 30 °C to obtain a solid. To remove residual acetic acid, the material was reconstituted in minimal acetone (<5 mL) and precipitated with hexanes (100 mL). The suspension was filtered through a fine fritted funnel, and the filter cake was rinsed with hexanes (3 x 20 mL). The solid was collected and reconstituted in acetone, dried with sodium sulfate, filtered through a fritted funnel using EtOAc, and concentrated by rotary evaporation at 30 °C to obtain the product. The material was dried under reduced pressure.

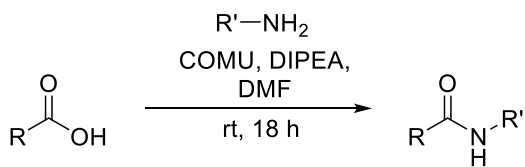

#### General Procedure X: COMU-Mediated Amide Bond Formation

To a flame-dried 10 mL round-bottom flask equipped with a magnetic stir bar under inert atmosphere ($\text{N}_2$) was added **5** (1.00 equiv), **S25**, **S26**, or **S30** (1.10 equiv), COMU (1.30 equiv), and distilled THF (0.10 M with respect to **5**). The solution was chilled in an ice bath for 5 minutes before DIPEA (3.30 equiv) was added. The ice bath was maintained for 30 minutes then held at room temperature for 18 h until the reaction was judged complete by TLC (2:3 hexanes:acetone). The reaction was quenched with saturated aqueous sodium bicarbonate (20 mL) and extracted with EtOAc (6 x 20 mL). To remove residual DIPEA and coupling agent byproducts from the combined organic phase, the combined organic phase was washed with saturated aqueous sodium bicarbonate (2 x 10 mL), and the aqueous washes were back extracted with EtOAc (2 x 10 mL). The combined organic phase was dried with sodium sulfate, filtered through a fritted funnel using EtOAc, and concentrated by rotary evaporation at 35 °C to afford a solid. The solid was purified by flash column chromatography using a gradient of 0-40% acetone in hexane to remove impurities, and slowly up to 85% acetone in hexanes to elute the product. The fractions were concentrated by rotary evaporation at 35 °C, and the product was dried overnight under reduced pressure to afford the product.

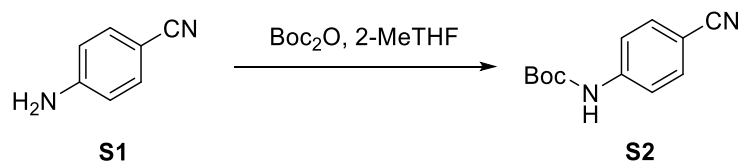

#### Scheme S1: Preparation of Compound S2

##### *tert*-Butyl (4-cyanophenyl)carbamate (**S2**)

To a flame-dried 100 mL cylindrical pressure vessel equipped with a magnetic stir bar was added di-*tert*-butyl dicarbonate (24.2258 g, 111.000 mmol), 4-aminobenzonitrile **S1** (3.5442 g, 30.000 mmol), and 2-MeTHF (16.3 mL, 1.88 M with respect to **S1**). The vessel was purged with nitrogen, sealed, and heated to 80 °C for 24 h, at which time the reaction was judged complete by TLC (7:3 hexanes:EtOAc). The solution was chilled in an ice bath for 20 minutes and the seal was released slowly—as the buildup of $\text{CO}_2$ could cause rapid depressurization. The solvent was removed by rotary evaporation at 60 °C to remove the *tert*-butanol byproduct and 2-MeTHF to afford an oil. The loose oil was poured into a 500 mL Erlenmeyer flask with residual transfer using hexanes. Additional hexanes was added (400 mL) to precipitate a white solid. The suspension was sonicated for 5 minutes then filtered over a fine fritted funnel to afford a white solid. The filter cake was rinsed with hexanes (3 x 100 mL) while agitating the solid each rinse with a spatula to ensure the removal of nonpolar impurities (i.e., excess di-*tert*-butyl dicarbonate). The filter cake was kept under vacuum filtration for 30 minutes forming a loose powder, then the solid was transferred to a 250 mL Erlenmeyer flask and dissolved with $\text{CHCl}_3$ (50 mL). The solution was dried using sodium sulfate, filtered through a fritted funnel using $\text{CHCl}_3$, and concentrated by rotary evaporation at 35 °C to afford a white, powdery solid. The solid was dried with stirring overnight under reduced pressure to afford **S2**. Yield: 5.3104 g, 24.331 mmol, white powder (81%).

**$^1\text{H}$ NMR** (500 MHz, $\text{CDCl}_3$) $\delta$ (ppm): 7.57 (d, $J$ = 8.8 Hz, 2H), 7.48 (d, $J$ = 8.8 Hz, 2H), 6.71 (br s, 1H), 1.52 (s, 9H). **$^{13}\text{C}$ NMR** (126 MHz, $\text{CDCl}_3$) $\delta$ (ppm): 152.07, 142.70, 133.44, 119.18, 118.21, 105.92, 81.84, 28.36. **HRMS QToF-ESI** calculated for $\text{C}_{12}\text{H}_{14}\text{N}_2\text{O}_2\text{Na}$ $[\text{M}+\text{Na}]^+$ $m/z$ 241.0948; found $m/z$ 241.0953.

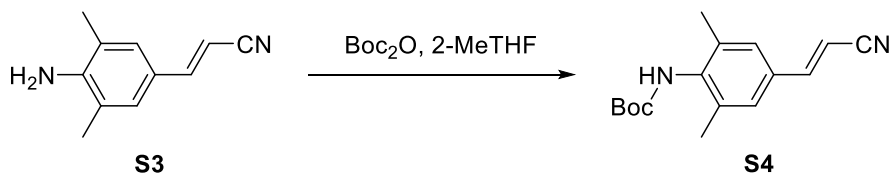

### Scheme S2: Preparation of Compound S4

#### *tert*-Butyl (*E*)-(4-(2-cyanovinyl)-2,6-dimethylphenyl)carbamate (**S4**)

To a flame-dried 50 mL pressure tube equipped with a magnetic stir bar was added di-*tert*-butyl dicarbonate (11.4038 g, 52.2557 mmol), (*E*)-3-(4-amino-3,5-dimethylphenyl)acrylonitrile **S3** (3.0000 g, 17.419 mmol), and 2-MeTHF (9.4 mL, 1.85 M with respect to **S3**). The vessel was purged with nitrogen, sealed, and heated to 80 °C for 24 h, at which time the reaction was judged complete by TLC (7:3 hexanes:EtOAc). The solution was chilled in an ice bath for 20 minutes and the seal was released slowly—as the buildup of $\text{CO}_2$ could cause rapid depressurization. The solvent was removed by rotary evaporation at 60 °C to remove the *tert*-butanol byproduct and 2-MeTHF to afford an oil. The loose oil was poured into a 500 mL Erlenmeyer flask with residual transfer using hexanes. Additional hexanes was added (400 mL) to precipitate a white solid. The suspension was sonicated for 5 minutes then filtered over a fine fritted funnel to afford a white solid. The filter cake was rinsed with hexanes (3 x 100 mL) while agitating the solid each rinse with a spatula to remove nonpolar impurities (i.e., excess di-*tert*-butyl dicarbonate). The filter cake was kept under vacuum filtration for 30 minutes forming a loose powder, then the solid was transferred to a 250 mL Erlenmeyer flask and dissolved with $\text{CHCl}_3$ (50 mL). The solution was dried using sodium sulfate, filtered through a fritted funnel using $\text{CHCl}_3$, and concentrated by rotary evaporation at 35 °C to afford an off-white powder. The solid was dried with stirring overnight under reduced pressure to afford **S4**. Yield: 4.0082 g, 14.717 mmol, off-white powder (84%).

**$^1\text{H}$ NMR** (500 MHz, $\text{CDCl}_3$) $\delta$ (ppm): $\delta$ 7.29 (d, $J$ = 16.6 Hz, 1H), 7.14 (s, 2H), 5.94 (br s, 1H), 5.80 (d, $J$ = 16.6 Hz, 1H), 2.28 (s, 6H), 1.49 (br s, 9H). **$^{13}\text{C}$ NMR** (126 MHz, $\text{CDCl}_3$) $\delta$ (ppm): 153.40,

150.32, 137.16, 136.53, 131.89, 127.33, 118.43, 95.98, 80.59, 28.39, 18.62. **HRMS QToF-ESI** calculated for C<sub>16</sub>H<sub>20</sub>N<sub>2</sub>O<sub>2</sub> [M+Na]<sup>+</sup> *m/z* 295.1418; found *m/z* 295.1422.

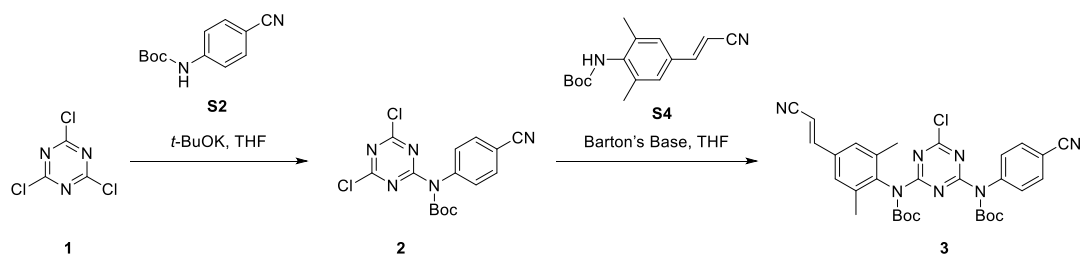

#### Scheme S3: Preparation of Compound 3

##### *tert*-Butyl (4-cyanophenyl)(4,6-dichloro-1,3,5-triazin-2-yl)carbamate (**2**)

To a flame-dried 250 mL round-bottom flask equipped with a magnetic stir bar under inert atmosphere (N<sub>2</sub>) (Flask A) was added **S2** (3.0000 g, 13.745 mmol). Under inert atmosphere, potassium *tert*-butoxide (3.8560 g, 34.363 mmol) was added to a fritted dropping funnel connected to Flask A and the potassium *tert*-butoxide was dissolved in distilled THF (82.5 mL, 0.17 M with respect to **S2**) (Note: the filtration was used to remove potassium hydroxide present in the potassium *tert*-butoxide formed from adventitious moisture, as potassium *tert*-butoxide is readily soluble in dry THF while potassium hydroxide has limited solubility). The resultant solution was filtered under inert atmosphere into Flask A which was maintained in an ice bath to limit exotherms. Into a separate flame-dried 500 mL round-bottom flask equipped with a magnetic stir bar under inert atmosphere (N<sub>2</sub>) (Flask B) was added cyanuric chloride **1** (7.6038 g, 41.235 mmol). The flask was chilled in a dry ice/EtOAc bath to -78 °C for 5 minutes before distilled THF (55 mL, 0.75 M with respect to **1**) was added to Flask B. After cooling Flask B for 15 minutes at -78 °C and stirring the contents of the flask until homogeneous, the contents of Flask A were slowly cannulated into Flask B over 35 minutes. The temperature of Flask B was maintained at -78 °C during this process. Once the contents of Flask A were fully transferred to Flask B, Flask B was maintained at -78 °C for an additional 4 h, at which time the reaction was judged complete by TLC (7:3 hexanes:EtOAc). The contents of Flask B were transferred to a 1 L Erlenmeyer flask and room temperature hexanes was added (900 mL) to precipitate a white solid. The contents of the Erlenmeyer flask were stirred for 15 minutes at room temperature to dissolve any remaining **1**. After 15 minutes, the suspension was filtered over a fine fritted funnel and the filter cake was rinsed with hexanes (3 x 150 mL), agitating the solid each time with a spatula to remove impurities and excess **1**. The filter cake was kept under vacuum filtration for 30 minutes on the fritted funnel forming a loose powder. To remove excess potassium *tert*-butoxide, the powder was transferred to a 250 mL Erlenmeyer flask and reconstituted with CHCl<sub>3</sub> (50 mL). Cold saturated aqueous sodium bicarbonate was added to the flask (100 mL), and the solution was transferred to a separatory funnel with CHCl<sub>3</sub>. The solution was washed with cold saturated aqueous sodium bicarbonate (2 x 100 mL) to remove the potassium *tert*-butoxide. The aqueous washes were combined and back extracted with CHCl<sub>3</sub> (3 x 50 mL). The combined organic phase was dried with sodium sulfate, filtered through a fritted funnel using CHCl<sub>3</sub>, and concentrated by rotary evaporation at 30 °C to afford a white solid. The solid was dried overnight by stirring under reduced pressure to afford **2**. Yield: 4.5200 g, 12.343 mmol, white powder (90%).

**<sup>1</sup>H NMR** (500 MHz, CDCl<sub>3</sub>) δ (ppm): 7.76 (d, *J* = 8.5 Hz, 2H), 7.33 (d, *J* = 8.5 Hz, 2H), 1.50 (s, 9H). **<sup>13</sup>C NMR** (126 MHz, CDCl<sub>3</sub>) δ (ppm): 171.61, 166.82, 150.38, 142.25, 133.57, 128.91, 118.11, 112.66, 86.11, 27.86. **HRMS QToF-ESI** calculated for C<sub>15</sub>H<sub>13</sub>N<sub>5</sub>O<sub>2</sub>Cl<sub>2</sub> [M+Na]<sup>+</sup> *m/z* 388.0345; found *m/z* 388.0344.

*tert*-Butyl (E)-4-((*tert*-butoxycarbonyl)(4-(2-cyanovinyl)-2,6-dimethylphenyl)amino)-6-chloro-1,3,5-triazin-2-yl)(4-cyanophenyl)carbamate (**3**)

To a flame-dried 250 mL round-bottom flask equipped with a magnetic stir bar under inert atmosphere (N<sub>2</sub>) was added **2** (3.0000 g, 8.1922 mmol), **S4** (2.2758 g, 8.3561 mmol), distilled THF (82.0 mL, 0.10 M with respect to **2**) and chilled in an ice bath for 15 minutes before adding Barton's base dropwise over 5 minutes (3.3 mL, 16 mmol). The ice bath was maintained for 4 h, at which time the reaction was judged complete by TLC (1:1 hexanes:EtOAc). The solution was poured over ~200 mL of ice in a 500 mL Erlenmeyer flask. The flask was placed in an ice bath and the solution was stirred with a spatula for 15 minutes to precipitate a solid. This suspension was filtered over a fine fritted funnel and washed with deionized water (3 x 100 mL) to remove residual THF and base. The filter cake was kept under vacuum filtration for 30 minutes forming a loose powder. The solid was purified by flash column chromatography using a gradient of 0-60% EtOAc in hexanes. Impurities and remaining excess **S4** eluted at ~20% EtOAc in hexanes, while **3** eluted slowly from 40-60% EtOAc in hexanes due to limited solubility. The fractions were collected and concentrated by rotary evaporation at 35 °C to obtain a white solid. The solid was dried overnight with stirring under reduced pressure to afford **3**. Yield: 3.7151 g, 6.1703 mmol, white powder (75%).

**<sup>1</sup>H NMR** (500 MHz, CDCl<sub>3</sub>) δ (ppm): 7.46 (d, *J* = 8.5 Hz, 2H), 7.34 (d, *J* = 16.7 Hz, 1H), 7.04-7.02 (m, 4H), 5.91 (d, *J* = 16.6 Hz, 1H), 1.96 (s, 6H), 1.40 (s, 9H), 1.38 (s, 9H). **<sup>13</sup>C NMR** (126 MHz, CDCl<sub>3</sub>) δ (ppm): 171.76, 166.71, 165.76, 150.45, 150.03, 149.51, 143.30, 139.94, 136.76, 133.31, 132.76, 128.75, 127.05, 118.24, 118.09, 111.46, 97.35, 85.02, 84.19, 27.84, 17.91. **HRMS QToF-ESI** calculated for C<sub>31</sub>H<sub>33</sub>N<sub>7</sub>O<sub>4</sub>Cl [M+H]<sup>+</sup> *m/z* 602.2283; found *m/z* 602.2263.

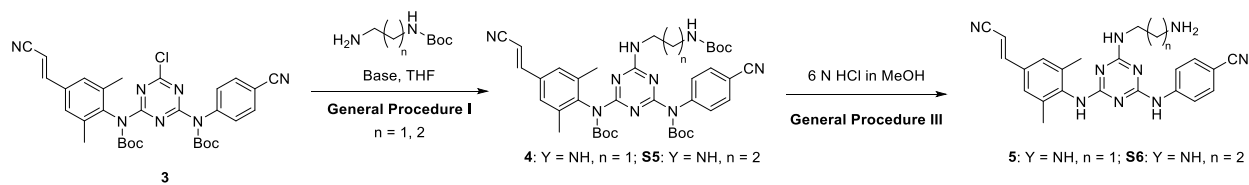

##### Scheme S4: Preparation of Compounds 5 and S6

*tert*-Butyl (E)-4-((*tert*-butoxycarbonyl)(4-(2-cyanovinyl)-2,6-dimethylphenyl)amino)-6-((2-((*tert*-butoxycarbonyl)amino)ethyl)amino)-1,3,5-triazin-2-yl)(4-cyanophenyl)carbamate (**4**)

**See General Procedure I to generate 4.** Quantities Used: **3** (1.5000 g, 2.4913 mmol), *tert*-butyl (2-aminoethyl)carbamate (580 μL, 3.74 mmol), distilled THF (25 mL, 0.10 M with respect to **3**), and DIPEA (870 μL, 5.00 mmol). Yield: 1.2781 g, 1.7608 mmol, white powder (71%).

**<sup>1</sup>H NMR** (500 MHz, CDCl<sub>3</sub>, major and minor rotamer) $\delta$ (ppm): 7.38 (dd, $J$ = 8.2, 2.7 Hz, 2H), 7.33 (dd, $J$ = 16.7, 2.2 Hz, 1H), 6.98 – 6.94 (m, 4H), 5.90 – 5.87 (m, 2H), 5.24 (dt, $J$ = 70.7, 5.7 Hz, 1H), 3.50 (d, $J$ = 5.9 Hz, 2H), 3.28 (dd, $J$ = 19.9, 5.9 Hz, 2H), 1.93 (d, $J$ = 7.4 Hz, 6H), 1.40 – 1.30 (m, 27H). **<sup>13</sup>C NMR** (126 MHz, CDCl<sub>3</sub>, major and minor rotamer) $\delta$ (ppm): 167.21, 167.15, 166.39, 166.35, 165.23, 165.06, 156.33, 151.16, 151.14, 150.68, 150.65, 149.82, 144.56, 144.54, 141.14, 141.05, 136.80, 136.72, 132.54, 132.50, 132.25, 128.77, 126.67, 118.55, 118.27, 110.31, 110.30, 96.60, 96.54, 83.43, 82.70, 82.63, 79.38, 79.29, 41.16, 40.97, 40.77, 28.48, 28.47, 27.90, 27.87, 27.85, 17.88. **HRMS QToF-ESI** calculated for C<sub>38</sub>H<sub>48</sub>N<sub>9</sub>O<sub>6</sub> [M+H]<sup>+</sup> $m/z$ 726.3728; found $m/z$ 726.3705.

*tert*-Butyl (E)-4-((*tert*-butoxycarbonyl)(4-(2-cyanovinyl)-2,6-dimethylphenyl)amino)-6-((3-((*tert*-butoxycarbonyl)amino)propyl)amino)-1,3,5-triazin-2-yl)(4-cyanophenyl)carbamate (**S5**)

**See General Procedure I to generate S5.** Quantities Used: **3** (600.0 mg, 0.9965 mmol), *tert*-butyl (3-aminopropyl)carbamate (260 μL, 1.49 mmol), distilled THF (10.0 mL, 0.10 M with respect to **5**), and DIPEA (350 μL, 2.00 mmol). Yield: 374.7 mg, 0.5064 mmol, white powder (51%).

**<sup>1</sup>H NMR** (500 MHz, CDCl<sub>3</sub>, major and minor rotamer) $\delta$ (ppm): 7.38 – 7.29 (m, 3H), 6.96 – 6.92 (m, 4H), 5.88 (br d, $J$ = 16.5 Hz, 2H), 5.49 (d, $J$ = 18.3 Hz, 1H), 3.46 (dd, $J$ = 10.1, 5.7 Hz, 2H), 3.12 – 3.10 (m, 2H), 1.91 (d, $J$ = 8.4 Hz, 6H), 1.69 – 1.64 (m, 2H), 1.41-1.29 (m, 27H). **<sup>13</sup>C NMR** (126 MHz, CDCl<sub>3</sub>, major and minor rotamer) $\delta$ (ppm): 167.19, 166.48, 166.22, 165.43, 164.92, 156.52, 151.13, 151.01, 150.66, 149.78, 144.58, 144.54, 141.19, 141.04, 136.75, 136.67, 132.49, 132.45, 132.19, 132.16, 128.71, 126.63, 126.61, 118.53, 118.23, 110.24, 110.22, 96.55, 96.53, 83.35, 83.33, 82.57, 82.54, 79.05, 37.80, 37.33, 30.52, 30.49, 28.54, 27.91, 27.85, 27.82, 17.84, 17.82. **HRMS QToF-ESI** calculated for C<sub>39</sub>H<sub>50</sub>N<sub>9</sub>O<sub>6</sub> [M+H]<sup>+</sup> $m/z$ 740.3884; found $m/z$ 740.3882.

(E)-4-((4-((2-aminoethyl)amino)-6-((4-(2-cyanovinyl)-2,6-dimethylphenyl)amino)-1,3,5-triazin-2-yl)amino)benzonitrile (**5**)

**See General Procedure III to generate 5.** Quantities Used: **4** (1.1500 g, 1.5857 mmol), and 6 N HCl in MeOH (15.8 mL, 0.10 M with respect to **4**). Crude yield: 585.4 mg, 1.376 mmol, white solid (87%). Compound **5** was used in subsequent reactions without further purification.

**<sup>1</sup>H NMR** (500 MHz, CD<sub>3</sub>OD, NH signals not evident in <sup>1</sup>H spectrum) $\delta$ (ppm): 7.91 (br s, 1H), 7.57 (br s, 2H), 7.44 (d, $J$ = 16.5 Hz, 1H), 7.30 (s, 3H), 6.13 (d, $J$ = 16.6 Hz, 1H), 3.45 (br s, 2H), 2.80 (br s, 2H), 2.21 (s, 6H). **<sup>13</sup>C NMR** (126 MHz, CD<sub>3</sub>OD) $\delta$ (ppm): 167.99, 166.89, 165.93, 151.90, 146.17, 140.22, 138.72, 133.71, 133.52, 128.24, 120.39, 120.17, 119.60, 104.67, 96.60, 44.04, 42.20, 18.69. **HRMS QToF-ESI** calculated for C<sub>23</sub>H<sub>24</sub>N<sub>9</sub> [M+H]<sup>+</sup> $m/z$ 426.2155; found $m/z$ 426.2157.

(*E*)-4-((4-((3-aminopropyl)amino)-6-((4-(2-cyanovinyl)-2,6-dimethylphenyl)amino)-1,3,5-triazin-2-yl)amino)benzonitrile (**S6**)

See General Procedure III to generate **14**. Quantities Used: **S5** (350.0 mg, 0.4731 mmol), and 6 N HCl in MeOH (4.8 mL, 0.10 M with respect to **S5**). Crude yield: 198.6 mg, 0.4518 mmol, white solid (96%). Compound **S6** was used to generate **11** and **S13** without further purification.

**<sup>1</sup>H NMR** (500 MHz, CD<sub>3</sub>OD, NH signals not evident in <sup>1</sup>H spectrum) δ (ppm): 7.96 (br s, 1H), 7.61 (s, 2H), 7.49 (d, *J* = 16.6 Hz, 1H), 7.35 (s, 3H), 6.18 (d, *J* = 16.6 Hz, 1H), 3.49 (br s, 2H), 2.72 (br s, 2H), 2.25 (br s, 6H), 1.77 (br s, 2H). **<sup>13</sup>C NMR** (126 MHz, CD<sub>3</sub>OD) δ (ppm): 167.85, 166.89, 165.91, 151.93, 146.20, 140.23, 138.72, 133.71, 133.53, 128.26, 120.39, 119.61, 104.74, 96.60, 39.47, 38.66, 33.27, 18.70. **HRMS QToF-ESI** calculated for C<sub>24</sub>H<sub>26</sub>N<sub>9</sub> [M+H]<sup>+</sup> *m/z* 440.2311; found *m/z* 440.2319.

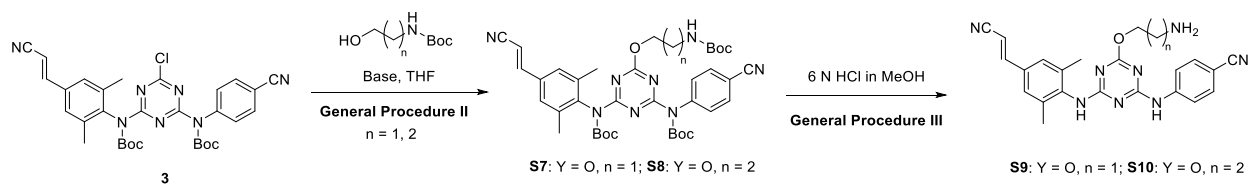

#### Scheme S5: Preparation of Compounds S9 and S10

*tert*-Butyl (*E*)-4-((*tert*-butoxycarbonyl)(4-(2-cyanovinyl)-2,6-dimethylphenyl)amino)-6-(2-((*tert*-butoxycarbonyl)amino)ethoxy)-1,3,5-triazin-2-yl)(4-cyanophenyl)carbamate (**S7**)

See General Procedure II to generate **S7**. Quantities Used: **3** (600.0 mg, 0.9965 mmol), *tert*-butyl (2-hydroxyethyl)carbamate (241.0 mg, 1.495 mmol), distilled THF (10.0 mL, 0.10 M with respect to **3**), and Barton's base (400 μL, 2.00 mmol). Yield: 373.6 mg, 0.5140 mmol, white solid (52%).

**<sup>1</sup>H NMR** (500 MHz, CDCl<sub>3</sub>) δ (ppm): 7.40 (d, *J* = 8.2 Hz, 2H), 7.32 (d, *J* = 16.7 Hz, 1H), 6.98 – 6.95 (m, 4H), 5.89 (d, *J* = 16.7 Hz, 1H), 5.17 (t, *J* = 6.0 Hz, 1H), 4.38 (t, *J* = 5.0 Hz, 2H), 3.47 (q, *J* = 5.4 Hz, 2H), 1.91 (s, 6H), 1.40 – 1.32 (m, 27H). **<sup>13</sup>C NMR** (126 MHz, CDCl<sub>3</sub>) δ (ppm): 171.57, 167.59, 166.45, 155.84, 150.56, 150.15, 149.56, 143.95, 140.51, 136.59, 132.84, 132.44, 128.78, 126.79, 118.30, 118.11, 110.88, 96.92, 84.08, 83.27, 79.41, 67.79, 39.67, 28.40, 27.79, 17.76. **HRMS QToF-ESI** calculated for C<sub>38</sub>H<sub>47</sub>N<sub>8</sub>O<sub>7</sub> [M+H]<sup>+</sup> *m/z* 727.3568; found *m/z* 727.3548.

*tert*-Butyl (*E*)-4-((*tert*-butoxycarbonyl)(4-(2-cyanovinyl)-2,6-dimethylphenyl)amino)-6-(3-((*tert*-butoxycarbonyl)amino)propoxy)-1,3,5-triazin-2-yl)(4-cyanophenyl)carbamate (**S8**)

See General Procedure II to generate **S8**. Quantities Used: **3** (600.0 mg, 0.9965 mmol), *tert*-butyl (3-hydroxypropyl)carbamate (261.9 mg, 1.495 mmol), distilled THF (10.0 mL, 0.10 M with respect to **3**), and Barton's base (400 μL, 2.00 mmol). Yield: 291.8 mg, 0.3939 mmol, white solid (40%).

**$^1\text{H}$ NMR** (500 MHz, $\text{CDCl}_3$) $\delta$ (ppm): 7.40 (d, $J$ = 8.1 Hz, 2H), 7.32 (d, $J$ = 16.7 Hz, 1H), 6.98 – 6.96 (m, 4H), 5.89 (d, $J$ = 16.6 Hz, 1H), 5.10 (br s, 1H), 4.42 (t, $J$ = 6.0 Hz, 2H), 3.19 (d, $J$ = 6.4 Hz, 2H), 1.90 (m, 8H), 1.41 – 1.32 (m, 27H). **$^{13}\text{C}$ NMR** (126 MHz, $\text{CDCl}_3$) $\delta$ (ppm): 171.80, 167.63, 166.52, 156.13, 150.61, 150.20, 149.61, 144.04, 140.61, 136.64, 132.83, 132.44, 128.80, 126.80, 118.34, 118.14, 110.83, 96.90, 84.01, 83.21, 79.18, 65.89, 37.30, 29.19, 28.50, 27.82, 17.80. **HRMS QToF-ESI** calculated for $\text{C}_{39}\text{H}_{49}\text{N}_8\text{O}_7$ $[\text{M}+\text{H}]^+$ $m/z$ 741.3724; found $m/z$ 741.3716.

*(E)-4-((4-(2-aminoethoxy)-6-((4-(2-cyanovinyl)-2,6-dimethylphenyl)amino)-1,3,5-triazin-2-yl)amino)benzonitrile (S9)*

**See General Procedure III to generate S9.** Quantities Used: **S7** (350.0 mg, 0.4815 mmol), and 6 N HCl in MeOH (4.8 mL, 0.10 M with respect to **S7**). Crude yield: 153.6 mg, 0.3601 mmol, white solid (75%). Compound **S9** was used to generate **8** and **S14** without further purification.

**$^1\text{H}$ NMR** (500 MHz, $\text{CD}_3\text{OD}$, major and minor rotamer, NH signals not evident in $^1\text{H}$ spectrum) $\delta$ (ppm): 8.01 – 7.35 (m, 7H), 6.26 – 6.16 (m, 1H), 4.45 – 4.23 (m, 2H), 3.04 – 2.90 (m, 2H), 2.26 (s, 6H). **$^{13}\text{C}$ NMR** (126 MHz, $\text{CD}_3\text{OD}$, major and minor rotamer) $\delta$ (ppm): 172.31, 167.96, 166.83, 151.78, 145.36, 138.69, 133.98, 133.68, 128.35, 121.02, 120.58, 120.08, 119.52, 105.66, 97.06, 69.47, 41.40, 18.53. **HRMS QToF-ESI** calculated for $\text{C}_{23}\text{H}_{23}\text{N}_8\text{O}$ $[\text{M}+\text{H}]^+$ $m/z$ 427.1995; found $m/z$ 427.2008.

*(E)-4-((4-(3-aminopropoxy)-6-((4-(2-cyanovinyl)-2,6-dimethylphenyl)amino)-1,3,5-triazin-2-yl)amino)benzonitrile (S10)*

**See General Procedure III to generate S10.** Quantities Used: **S8** (280.0 mg, 0.3779 mmol), and 6 N HCl in MeOH (3.8 mL, 0.10 M with respect to **S8**). Crude yield: 116.1 mg, 0.2636 mmol, white solid (70%). Compound **S10** was used to generate **9** and **S15** without further purification.

**$^1\text{H}$ NMR** (500 MHz, $\text{CD}_3\text{OD}$, major and minor rotamer, NH signals not evident in $^1\text{H}$ spectrum) $\delta$ (ppm): 8.00 – 7.32 (m, 7H), 6.25 – 6.14 (m, 1H), 4.49 – 4.29 (m, 2H), 2.84 – 2.69 (m, 2H), 2.25 (s, 6H), 1.96 – 1.80 (m, 2H). **$^{13}\text{C}$ NMR** (126 MHz, $\text{CD}_3\text{OD}$, major and minor rotamer) $\delta$ (ppm): 172.28, 167.94, 166.80, 151.78, 145.36, 139.74, 138.68, 133.97, 133.67, 128.33, 120.98, 120.54, 119.53, 105.60, 97.03, 66.16, 39.36, 32.75, 18.56. **HRMS QToF-ESI** calculated for $\text{C}_{24}\text{H}_{25}\text{N}_8\text{O}$ $[\text{M}+\text{H}]^+$ $m/z$ 441.2151; found $m/z$ 441.2168.

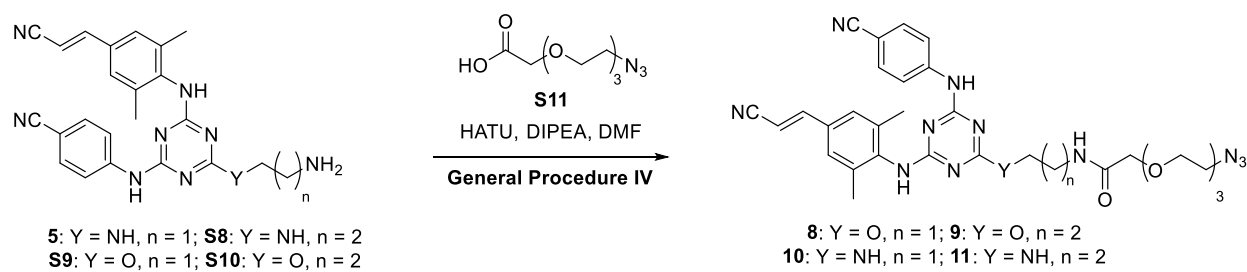

#### Scheme S6: Preparation of Compounds 8-11

*(E)*-2-(2-(2-(2-azidoethoxy)ethoxy)ethoxy)-*N*-(2-((4-((4-cyanophenyl)amino)-6-((4-(2-cyanovinyl)-2,6-dimethylphenyl)amino)-1,3,5-triazin-2-yl)amino)ethyl)acetamide (**8**)

**See General Procedure IV to generate 8.** Quantities Used: **5** (30.0 mg, 0.0705 mmol), 2-(2-(2-(2-azidoethoxy)ethoxy)ethoxy)acetic acid (26.0 mg, 0.110 mmol), HATU (45.0 mg, 0.110 mmol), DMF (500 μL, 0.2 M with respect to **5**), and DIPEA (60 μL, 0.33 mmol). Yield: 15.0 mg, 0.0234 mmol, white solid (33%). **R<sub>f</sub>**: 0.26 (3% MeOH in CHCl<sub>3</sub>).

**<sup>1</sup>H NMR** (500 MHz, CD<sub>3</sub>OD, major and minor rotamer, NH signals not evident in <sup>1</sup>H spectrum) $\delta$ (ppm): 8.12 – 7.98 (m, 1H), 7.62 – 7.38 (m, 6H), 6.21 (d, *J* = 14.1 Hz, 1H), 3.94 (br s, 1H), 3.65 – 3.31 (br m, 10H), 2.26 (s, 6H), 1.90 (m, 2H). **<sup>13</sup>C NMR** (126 MHz, CD<sub>3</sub>OD, major and minor rotamer) $\delta$ (ppm): 173.71, 173.28, 167.94, 167.82, 166.90, 166.50, 165.89, 165.67, 151.93, 146.21, 138.76, 133.63, 128.28, 120.37, 120.18, 119.60, 104.82, 96.66, 71.88, 71.50, 71.39, 71.20, 71.02, 51.73, 50.72, 40.88, 18.71, 18.66. **HRMS QToF-ESI** calculated for C<sub>31</sub>H<sub>36</sub>N<sub>12</sub>O<sub>4</sub> [M+H]<sup>+</sup> *m/z*: 641.3061; found *m/z* 641.3044.

*(E)*-2-(2-(2-(2-azidoethoxy)ethoxy)ethoxy)-*N*-(3-((4-((4-cyanophenyl)amino)-6-((4-(2-cyanovinyl)-2,6-dimethylphenyl)amino)-1,3,5-triazin-2-yl)amino)propyl)acetamide (**9**)

**See General Procedure IV to generate 9.** Quantities Used: **S8** (35.0 mg, 0.0796 mmol), 2-(2-(2-(2-azidoethoxy)ethoxy)ethoxy)acetic acid (22.8 mg, 0.0978 mmol), HATU (37.2 mg, 0.0978 mmol), DMF (350 μL, 0.25 M with respect to **S8**), and DIPEA (50 μL, 0.28 mmol). Yield: 22.1 mg, 0.0338 mmol, white solid (42%). **R<sub>f</sub>**: 0.20 (3% MeOH in CHCl<sub>3</sub>).

**<sup>1</sup>H NMR** (500 MHz, CD<sub>3</sub>OD, major and minor rotamer, NH signals not evident in <sup>1</sup>H spectrum) $\delta$ (ppm): 8.00 – 7.32 (m, 7H), 6.25 – 6.14 (m, 1H), 4.49 – 4.29 (m, 2H), 2.84–2.69 (m, 2H), 2.25 (s, 6H), 1.96 – 1.80 (m, 2H). **<sup>13</sup>C NMR** (126 MHz, CD<sub>3</sub>OD, major and minor rotamer) $\delta$ (ppm): 172.28, 167.94, 166.80, 151.78, 145.36, 139.74, 138.68, 133.97, 133.67, 128.33, 120.98, 120.54, 119.53, 105.60, 97.03, 66.16, 39.36, 32.75, 18.56. **HRMS QToF-ESI** calculated for C<sub>24</sub>H<sub>25</sub>N<sub>8</sub>O [M+H]<sup>+</sup> *m/z* 441.2151; found *m/z* 441.2168.

*(E)*-2-(2-(2-(2-azidoethoxy)ethoxy)ethoxy)-*N*-(2-((4-((4-cyanophenyl)amino)-6-((4-(2-cyanovinyl)-2,6-dimethylphenyl)amino)-1,3,5-triazin-2-yl)oxy)ethyl)acetamide (**10**)

**See General Procedure IV to generate 10.** Quantities Used: **S9** (35.0 mg, 0.0821 mmol), 2-(2-(2-(2-azidoethoxy)ethoxy)ethoxy)acetic acid (22.4 mg, 0.0962 mmol), HATU (36.6 mg, 0.0962 mmol), DMF (400 μL, 0.20 M with respect to **S9**), and DIPEA (50 μL, 0.28 mmol). Yield: 21.0 mg, 0.0327 mmol, white solid (40%). **R<sub>f</sub>**: 0.31 (3% MeOH in CHCl<sub>3</sub>).

**<sup>1</sup>H NMR** (500 MHz, acetone-d<sub>6</sub>, major and minor rotamer) $\delta$ (ppm): 8.72 – 8.60 (br m, 1H), 8.01 – 7.95 (m, 2H), 7.79 – 7.73 (m, 2H), 7.45 – 7.36 (m, 7H), 6.57 (br s, 1H), 6.19 (d, $J$ = 11.2 Hz, 1H), 3.81 (m, 2H), 3.53 – 3.28 (m, 10H), 2.18 and 2.15 (s, 7H), 1.77 (m, 4H). **<sup>13</sup>C NMR** (126 MHz, acetone-d<sub>6</sub>, major and minor rotamer) $\delta$ (ppm): 170.76, 167.76 and 167.54, 166.61 and 166.35, 165.66 and 165.42, 151.01, 145.86, 145.78, 139.96, 138.08, 133.52, 133.34, 132.93, 127.94, 119.94, 119.20, 104.59, 104.44, 96.80, 71.54, 71.10, 70.96, 70.90, 70.55, 69.70, 55.47, 51.32, 41.50, 40.04, 38.72, 18.81. **HRMS QToF-ESI** calculated for C<sub>31</sub>H<sub>35</sub>N<sub>11</sub>O<sub>5</sub> [M+H]<sup>+</sup> $m/z$ 642.2892; found $m/z$ 642.2901.

*(E)-2-(2-(2-(2-azidoethoxy)ethoxy)ethoxy)-N-(3-((4-((4-cyanophenyl)amino)-6-((4-(2-cyanovinyl)-2,6-dimethylphenyl)amino)-1,3,5-triazin-2-yl)oxy)propyl)acetamide (11)*

**See General Procedure IV to generate 11.** Quantities Used: **S10** (30.0 mg, 0.0681 mmol), 2-(2-(2-(2-azidoethoxy)ethoxy)ethoxy)acetic acid (19.0 mg, 0.0815 mmol), HATU (31.2 mg, 0.0820 mmol), DMF (350 μL, 0.20 M with respect to **S10**), and DIPEA (40 μL, 0.24 mmol). Yield: 13.2 mg, 0.0201 mmol, white solid (30%). **R<sub>f</sub>**: 0.25 (3% MeOH in CHCl<sub>3</sub>).

**<sup>1</sup>H NMR** (500 MHz, acetone-d<sub>6</sub>, major and minor rotamer), $\delta$ (ppm): 8.72-8.56 (br m, 1H), 8.00-7.78 (m, 4H), 7.44-7.36 (m, 6H), 7.12-7.11 (m, 1H), 6.51 (s, 1H), 6.19 (s, $J$ = 14.5 Hz, 1H), 3.89 (s, 1H), 3.62-3.54 (m, 4H), 3.34-3.08 (m, 7H), 2.18 and 2.16 (s, 6H, major and minor rotamer), 1.78 (br s, 2H), 1.69-1.66 (m, 2H); **<sup>13</sup>C NMR** (126 MHz, acetone-d<sub>6</sub>, major and minor rotamer) $\delta$ (ppm): 167.36 and 167.20, 166.53 and 166.10, 165.58 and 165.35, 151.00, 145.77, 139.89, 138.09, 133.47, 133.35, 132.96, 127.94, 119.92, 119.20, 104.64 and 104.42, 96.81, 71.75, 71.11, 71.03, 70.92, 70.56, 69.72, 68.04, 55.45, 51.29, 38.67 and 38.28, 37.23 and 37.12, 22.96, 18.79. **HRMS QToF-ESI** calculated for C<sub>32</sub>H<sub>37</sub>N<sub>11</sub>O<sub>5</sub> [M+Na]<sup>+</sup> 678.2877; found $m/z$ 678.2836.

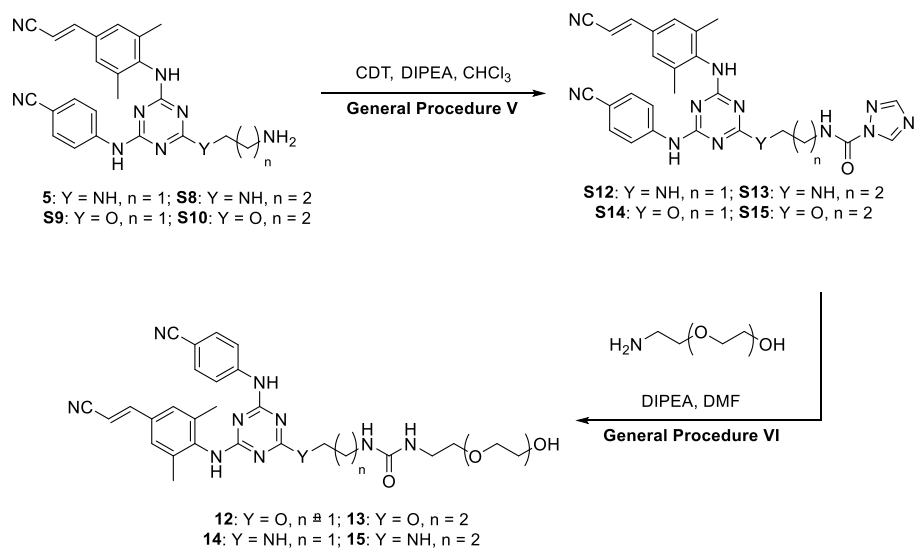

#### Scheme S7: Preparation of Compounds 12-15

*(E)*-1-(2-((4-((4-cyanophenyl)amino)-6-((4-(2-cyanovinyl)-2,6-dimethylphenyl)amino)-1,3,5-triazin-2-yl)amino)ethyl)-3-(2-(2-(2-(2-hydroxyethoxy)ethoxy)ethoxy)ethyl)urea (**14**)

**See General Procedure V to generate S12.** Quantities Used: **5** (60.0 mg, 0.140 mmol), CDT (45.0 mg, 0.280 mmol), DCM (1.4 mL, 0.10 M with respect to **5**), and DIPEA (40 μL, 0.21 mmol). Crude Yield: 62.2 mg, 0.119 mmol, white solid. Compound **S12** was used to generate **14** without further purification.

**See General Procedure VI to generate 14.** Quantities Used: **S12** (62.2 mg, 0.119 mmol), DMF (1.2 mL, 0.10 M with respect to **S12**), DIPEA (20 μL, 0.24 mmol), and 2-(2-(2-(2-hydroxyethoxy)ethoxy)ethoxy)ethan-1-ol (69.8 mg, 0.361 mmol). Yield: 26.1 mg, 0.0405 mmol, white solid (29% over two steps). **R<sub>f</sub>**: 0.31 (3% MeOH in CHCl<sub>3</sub>).

**<sup>1</sup>H NMR** (500 MHz, CD<sub>3</sub>CN, major and minor rotamer) $\delta$ (ppm): 8.19 (s, 1H), 7.92 (br s, 1H), 7.55-7.20 (m, 6H), 6.32 (br s, 1H), 6.06 (d, $J$ = 16.1 Hz, 1H), 5.63 (br s, 2H), 3.59-3.37 (m, 14H), 3.28-3.21 (m, 5H), 2.29 (br s, 2H), 2.23 and 2.21 (s, 6H). **<sup>13</sup>C NMR** (126 MHz, CD<sub>3</sub>CN, major and minor rotamer) $\delta$ (ppm): 167.41, 167.29, 166.42, 166.06, 165.45, 165.14, 160.14, 151.19, 145.44, 139.51, 138.29, 133.73, 133.09, 127.99, 120.27, 119.94, 119.56, 104.72, 96.92, 73.14, 71.21, 70.96, 70.91, 70.81, 70.75, 61.73, 55.18, 42.55, 40.89, 32.19, 29.70, 18.82. **HRMS QToF-ESI** calculated for C<sub>32</sub>H<sub>40</sub>N<sub>10</sub>O<sub>5</sub> [M+H]<sup>+</sup> $m/z$ 645.3211; found $m/z$ 645.3245.

*(E)*-1-(3-((4-((4-cyanophenyl)amino)-6-((4-(2-cyanovinyl)-2,6-dimethylphenyl)amino)-1,3,5-triazin-2-yl)amino)propyl)-3-(2-(2-(2-(2-hydroxyethoxy)ethoxy)ethoxy)ethyl)urea (**15**)

**See General Procedure V to generate S13.** Quantities Used: **S8** (60.0 mg, 0.137 mmol), CDT (30.0 mg, 0.180 mmol), DCM (0.90 mL, 0.15 M with respect to **S8**), DIPEA (30 μL, 0.17 mmol). Crude Yield: 60.0 mg, 0.112 mmol, white solid. Compound **S13** was used to generate **15** without further purification.

**See General Procedure VI 15.** Quantities Used: **S13** (60.0 mg, 0.112 mmol), DMF (1.1 mL, 0.10 M with respect to **S13**), DIPEA (40 μL, 0.22 mmol) and 2-(2-(2-(2-aminoethoxy)ethoxy)ethoxy)ethan-1-ol (64.0 mg, 0.331 mmol). Yield: 34.0 mg, 0.0516 mmol, white solid (38% over two steps). **R<sub>f</sub>**: 0.13 (3% MeOH in CHCl<sub>3</sub>).

**<sup>1</sup>H NMR** (500 MHz, CD<sub>3</sub>OD, major and minor rotamer, **NH** and **OH** signals not evident in <sup>1</sup>H spectrum) $\delta$ (ppm): 7.99 (br s, 1H), 7.64 (br s, 2H), 7.52 (d, *J* = 16.7 Hz, 1H), 7.38 (br s, 3H), 6.22 (d, *J* = 16.3 Hz, 1H), 3.65-3.60 (m, 8H), 3.55-3.51 (m, 5H), 2.26 (s, 6H), 1.76 (br s, 2H). **<sup>13</sup>C NMR** (126 MHz, CD<sub>3</sub>OD, major and minor rotamer) $\delta$ (ppm): 167.18, 166.47, 166.06, 165.50, 165.18, 160.17, 151.20, 145.58, 139.55, 138.34, 133.70, 133.08, 127.99, 120.28, 119.91, 119.56, 104.66, 96.91, 73.09, 71.49, 70.92, 70.71, 61.69, 40.73, 37.64, 31.75, 31.36, 18.80. **HRMS QToF-ESI** calculated for C<sub>33</sub>H<sub>42</sub>N<sub>10</sub>O<sub>5</sub> [M+H]<sup>+</sup> *m/z* 660.3340; found *m/z* 660.3350.

*(E)*-1-(2-((4-((4-cyanophenyl)amino)-6-((4-(2-cyanovinyl)-2,6-dimethylphenyl)amino)-1,3,5-triazin-2-yl)oxy)ethyl)-3-(2-(2-(2-(2-hydroxyethoxy)ethoxy)ethoxy)ethyl)urea (**12**)

**See General Procedure V to generate S14.** Quantities Used: **S9** (60.0 mg, 0.139 mmol), CDT (46.0 mg, 0.281 mmol), DCM (1.4 mL, 0.10 M with respect to **S9**), and DIPEA (40 μL, 0.21 mmol). Crude Yield: 60.0 mg, 0.112 mmol, white solid. Compound **S14** was used to generate **12** without further purification.

**See General Procedure VI to generate 12.** Quantities Used: **S14** (60.0 mg, 0.112 mmol), DMF (1.2 mL, 0.10 M with respect to **S14**), DIPEA (40 μL, 0.24 mmol), and 2-(2-(2-(2-aminoethoxy)ethoxy)ethoxy)ethan-1-ol (66.0 mg, 0.342 mmol). Yield: 22.1 mg, 0.0335 mmol, white solid (24% over two steps). **R<sub>f</sub>**: 0.25 (3% MeOH in CHCl<sub>3</sub>).

**<sup>1</sup>H NMR** (500 MHz, CD<sub>3</sub>OD, major and minor rotamer, **NH** and **OH** signals not evident in <sup>1</sup>H spectrum) $\delta$ (ppm): 8.01 (d, *J* = 6.8 Hz, 1H), 7.65 (d, *J* = 7.9 Hz, 1H), 7.56-7.33 (m, 5H), 6.24 and 6.17 (each d, *J* = 16.4 Hz, 1H), 4.44 and 4.24 (each s, 1H), 3.66-3.47 (m, 16H), 2.25 (s, 6H). **<sup>13</sup>C NMR** (126 MHz, CD<sub>3</sub>OD, major and minor rotamer) $\delta$ (ppm): 172.34, 172.27, 168.07, 167.91, 166.77, 161.04, 160.93, 151.86, 151.77, 145.54, 145.34, 139.70, 139.11, 138.64, 138.48, 133.99, 133.96, 133.88, 133.70, 128.34, 120.93, 120.52, 120.24, 120.10, 119.54, 105.73, 105.58, 97.04, 96.83, 73.61, 71.48, 71.32, 71.18, 67.57, 62.13, 40.98, 40.33, 18.60. **HRMS QToF-ESI** calculated for C<sub>32</sub>H<sub>39</sub>N<sub>9</sub>O<sub>6</sub> [M+H]<sup>+</sup> *m/z* 646.3023; found *m/z* 646.3088.

*(E)*-1-(3-((4-((4-cyanophenyl)amino)-6-((4-(2-cyanovinyl)-2,6-dimethylphenyl)amino)-1,3,5-triazin-2-yl)oxy)propyl)-3-(2-(2-(2-(2-hydroxyethoxy)ethoxy)ethoxy)ethyl)urea (**13**)

**See General Procedure V to generate S15.** Quantities Used: **S10** (60.0 mg, 0.137 mmol), CDT (44.8 mg, 0.272 mmol), DCM (1.4 mL, 0.10 M with respect to **S10**), and DIPEA (40 μL, 0.20 mmol). Crude Yield: 60.2 mg, 0.112 mmol, white solid. Compound **S15** was used to generate **13** without further purification.

**See General Procedure VI to generate 13.** Quantities Used: **S15** (60.2 mg, 0.112 mmol), DMF (1.1 mL, 0.10 M with respect to **S15**), DIPEA (30 μL, 0.17 mmol), and 2-(2-(2-(2-

aminoethoxy)ethoxy)ethoxy)ethan-1-ol (66.0 mg, 0.342 mmol). Yield: 23.2 mg, 0.0352 mmol, white solid (26% over two steps). **R<sub>f</sub>**: 0.28 (3% MeOH in CHCl<sub>3</sub>).

**<sup>1</sup>H NMR** (500 MHz, CD<sub>3</sub>CN, major and minor rotamer) $\delta$ (ppm): 8.49-8.37 (m, 1H), 7.95 (br s, 1H), 7.85 (br s, 1H), 7.66 (d, *J* = 6.5 Hz, 1H), 7.58-7.16 (m, 7H), 6.10-6.04 (each d, *J* = 16.7 Hz, 1H), 5.44 (s, 1H), 5.30 (s, 1H), 4.34 (s, 1H), 4.17 (s, 1H), 3.58-3.44 (m, 15H), 3.28-3.10 (m, 5H), 2.20 (s, 8H), 1.85 and 1.75 (br s, 2H). **<sup>13</sup>C NMR** (126 MHz, CD<sub>3</sub>CN, major and minor rotamer) $\delta$ (ppm): 172.15, 171.99, 167.70, 166.48, 159.62, 159.52, 153.24, 153.14, 151.12, 144.87 and 144.72, 138.28, 133.97, 133.71, 133.47, 129.38, 128.07, 120.58, 120.10, 119.53, 105.59 and 105.41, 97.21, 73.22, 71.18, 71.08, 70.77, 65.66, 61.84, 59.67, 55.21, 40.75, 37.52, 30.50, 18.70. **HRMS QToF-ESI** calculated for C<sub>33</sub>H<sub>41</sub>N<sub>9</sub>O<sub>6</sub> [M+H]<sup>+</sup> *m/z* 659.3418; found *m/z* 659.3408.

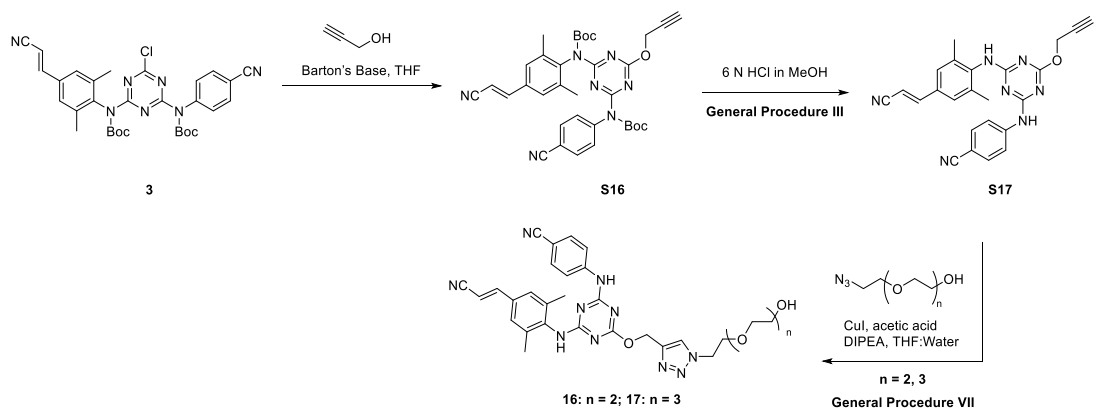

### Scheme S8: Preparation of Compounds 16-17

*tert*-Butyl (*E*)-(4-((*tert*-butoxycarbonyl)(4-cyanophenyl)amino)-6-(prop-2-yn-1-yloxy)-1,3,5-triazin-2-yl)(4-(2-cyanovinyl)-2,6-dimethylphenyl)carbamate (**S16**)

To a flame-dried 50 mL round-bottom flask equipped with a magnetic stir bar under inert atmosphere (N<sub>2</sub>) was added **3** (600.0 mg, 0.9965 mmol), propargyl alcohol (100 μL, 1.73 mmol), and distilled THF (8.3 mL, 0.10 M with respect to **3**). The solution was chilled in an ice bath for 10 minutes before adding Barton's base (330 μL, 1.66 mmol) dropwise over 5 minutes. The cold bath was maintained 15 minutes, then allowed to warm to room temperature for 4 h, at which time the reaction was judged complete by TLC (1:1 hexanes:EtOAc). The solution was poured into ~100 mL of ice in a 125 mL Erlenmeyer flask, placed in an ice bath, and stirred with a spatula for 10 minutes to precipitate a solid. This suspension was filtered over a fine fritted funnel and washed with deionized water (3 x 50 mL) to remove residual THF, excess propargyl alcohol, and base. Vacuum filtration of the filter cake on the fritted funnel formed a loose powder after 30 minutes, and the solid was purified by flash column chromatography using a gradient of 0-40% EtOAc in hexanes. Compound **S16** eluted with 40% EtOAc in hexanes. The fractions were concentrated by rotary evaporation at 35 °C to afford a white solid. The solid was dried overnight with stirring under reduced pressure to afford **S16**. Yield: 496.3 mg, 0.7983 mmol, off-white solid (80%).

**<sup>1</sup>H NMR** (500 MHz, CDCl<sub>3</sub>) δ (ppm): 7.48 (d, *J* = 8.5 Hz, 2H), 7.34 (d, *J* = 16.6 Hz, 1H), 7.07 (d, *J* = 8.6 Hz, 2H), 7.04 (s, 2H), 5.89 (d, *J* = 16.6 Hz, 1H), 4.90 (d, *J* = 2.5 Hz, 2H), 2.44 (t, *J* = 2.4 Hz, 1H), 1.99 (s, 6H), 1.40 (s, 9H), 1.37 (s, 9H). **<sup>13</sup>C NMR** (126 MHz, CDCl<sub>3</sub>) δ (ppm): 170.91, 167.75, 166.78, 150.90, 150.38, 149.77, 143.96, 140.60, 136.92, 133.02, 132.63, 128.87, 126.98, 118.42, 118.20, 111.03, 96.94, 84.28, 83.43, 75.49, 55.48, 27.88, 17.94. **HRMS QToF-ESI** calculated for C<sub>34</sub>H<sub>36</sub>N<sub>7</sub>O<sub>5</sub> [M+H]<sup>+</sup> *m/z* 622.2778; found *m/z* 622.2770.

*(E)*-4-((4-((4-(2-cyanovinyl)-2,6-dimethylphenyl)amino)-6-(prop-2-yn-1-yloxy)-1,3,5-triazin-2-yl)amino)benzonitrile (**S17**)

**See General Procedure III to generate S17.** Quantities Used: **S16** (350.0 mg, 0.5630 mmol), and 6 N HCl in MeOH (5.6 mL, 0.10 M with respect to **S16**). Crude Yield: 177.4 mg, 0.4218 mmol, off-white solid (75%). Compound **S17** was used to generate **16** and **17** without further purification.

*(E)*-4-((4-((4-(2-cyanovinyl)-2,6-dimethylphenyl)amino)-6-((1-(2-(2-(2-hydroxyethoxy)ethoxy)ethyl)-1H-1,2,3-triazol-4-yl)methoxy)-1,3,5-triazin-2-yl)amino)benzonitrile (**16**)

**See General Procedure VII to generate 16.** Quantities Used: **S17** (50.0 mg, 0.120 mmol), 2-(2-(2-azidoethoxy)ethoxy)ethan-1-ol (31.0 mg, 0.180 mmol), copper (I) iodide (3.0 mg, 0.012 mmol), glacial acetic acid (1 drop), DIPEA (1 drop), 5:1 v/v THF:deionized water (1.0 mL, 0.12 M with respect to **16**). The reaction was doped with copper (I) iodide (3.0 mg, 0.014 mmol) at *t* = 3 h, 6 h, and 9 h. Yield: 51.9 mg, 0.0870 mmol, light yellow solid (34%).

**<sup>1</sup>H NMR** (500 MHz, CDCl<sub>3</sub>, major and minor rotamer) δ (ppm): 7.99 (s, 1H), 7.82 – 7.34 (m, 6H), 7.24 and 7.19 (each s, 2H), 5.91 and 5.85 (each d, *J* = 16.4 Hz and *J* = 17.4 Hz, 1H), 5.54 and 5.32 (each s, 2H), 4.55 and 4.48 (each s, 2H), 3.86 – 3.53 (m, 11H), 2.25 (s, 7H). **<sup>13</sup>C NMR** (126 MHz, CDCl<sub>3</sub>, major and minor rotamer) δ (ppm): 170.30, 166.45, 165.20, 149.93, 143.08, 142.81, 137.37, 133.22, 133.02, 132.63, 127.22, 125.12, 120.04, 119.33, 119.14, 118.20, 105.72, 96.81, 72.55, 70.48, 70.30, 69.31, 61.69, 60.81, 50.46, 18.82. **HRMS QToF-ESI** calculated for C<sub>30</sub>H<sub>33</sub>N<sub>10</sub>O<sub>4</sub> [M+H]<sup>+</sup> *m/z* 597.2686; found *m/z* 597.2681.

*(E)*-4-((4-((4-(2-cyanovinyl)-2,6-dimethylphenyl)amino)-6-((1-(2-(2-(2-(2-hydroxyethoxy)ethoxy)ethoxy)ethyl)-1H-1,2,3-triazol-4-yl)methoxy)-1,3,5-triazin-2-yl)amino)benzonitrile (**17**)

**See General Procedure VII to generate 17.** Quantities Used: **S17** (50.0 mg, 0.120 mmol), 2-(2-(2-(2-azidoethoxy)ethoxy)ethoxy)ethan-1-ol (40.0 mg, 0.180 mmol), copper (I) iodide (3.0 mg, 0.012 mmol), glacial acetic acid (1 drop), DIPEA (1 drop), 5:1 v/v THF:deionized water (1.0 mL, 0.12 M with respect to **S17**). The reaction was doped with copper (I) iodide (3.0 mg, 0.014 mmol) at *t* = 3 h, 6 h, and 9 h. Yield: 60.0 mg, 0.0936 mmol, light yellow solid (78%).

**<sup>1</sup>H NMR** (500 MHz, CDCl<sub>3</sub>, major and minor rotamer) δ (ppm): 8.55 (br s, 1H), 7.93 – 6.95 (m, 10H), 5.91 and 5.85 (each d, *J* = 16.6 Hz, 16.7 Hz, 1H), 5.50 and 5.29 (each br s, 2H), 4.53 and 4.46 (each br s, 2H), 3.85 and 3.80 (each t, *J* = 5.0 Hz, 5.0 Hz, 2H), 3.71 – 3.55 (m, 13H), 2.36 (br s, 1H), 2.23 (s, 6H). **<sup>13</sup>C NMR** (126 MHz, CDCl<sub>3</sub>, major and minor rotamer) δ (ppm): 170.70, 170.26, 166.62, 165.21, 149.99, 143.02, 137.63, 137.33, 137.21, 137.00, 133.11, 132.91, 132.41, 127.34, 127.15, 125.36, 120.07, 119.34, 119.23, 118.27, 105.61, 105.36, 96.61, 96.45, 72.56, 70.52, 70.44, 70.34, 70.23, 69.35, 61.54, 60.46, 50.49, 29.78, 27.90, 18.79. **HRMS QToF-ESI** calculated for C<sub>32</sub>H<sub>37</sub>N<sub>10</sub>O<sub>5</sub> [M+H]<sup>+</sup> *m/z* 641.2948; found *m/z* 641.2946.

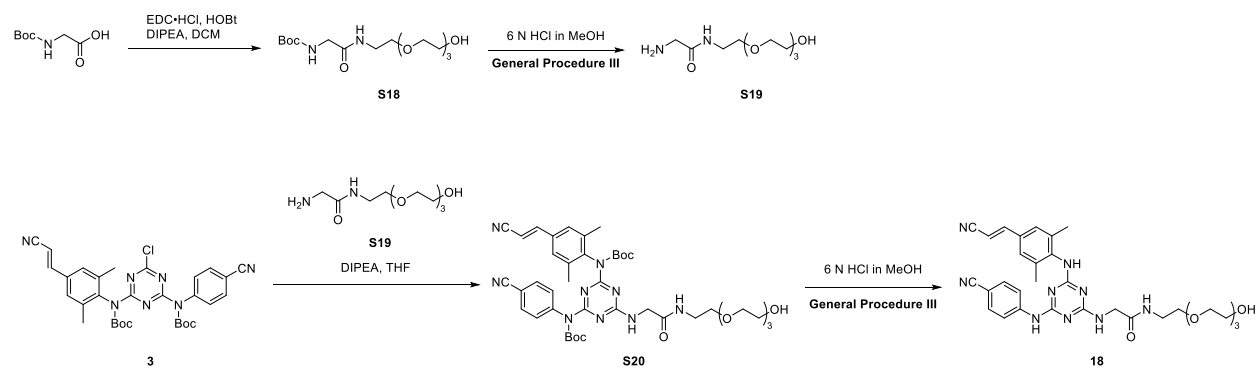

#### Scheme S9: Preparation of Compound 18

##### *tert*-Butyl (14-hydroxy-2-oxo-6,9,12-trioxa-3-azatetradecyl)carbamate (**S18**)

To a flame-dried 25 mL round-bottom flask equipped with a magnetic stir bar under inert atmosphere (N<sub>2</sub>) was added 2-(2-(2-(2-aminoethoxy)ethoxy)ethoxy)ethan-1-ol (500.0 mg, 2.587 mmol), (*tert*-butoxycarbonyl)glycine (543.9 mg, 3.105 mmol), EDC·HCl (644.7 mg, 3.363 mmol), HOBt hydrate (515.0 mg, 3.363 mmol), and DCM (8.6 mL, 0.30 M with respect to 2-(2-(2-(2-aminoethoxy)ethoxy)ethoxy)ethan-1-ol). The solution was chilled in an ice bath for 5 minutes before DIPEA (1.6 mL, 9.1 mmol) was added by syringe. The solution was stirred for 30 minutes in the ice bath then at room temperature for 18 h, at which time the reaction was judged complete by TLC (1:1 hexanes:diethyl ether, KMnO<sub>4</sub> stain). The reaction was quenched with saturated aqueous sodium bicarbonate (20 mL) and extracted with EtOAc (6 x 15 mL). To remove residual base and coupling agent byproducts, the combined organic phase was washed with the saturated aqueous sodium bicarbonate (2 x 10 mL), and the washes were back extracted with EtOAc (2 x 10 mL). The organic phase was dried with sodium sulfate, filtered through a fritted funnel using EtOAc, and concentrated by rotary evaporation at 35 °C to obtain an oil. The oil was purified by flash column chromatography using a gradient of 0-60% diethyl ether in hexanes. The material eluted with 40-60% diethyl ether in hexanes to afford **S18**. Yield: 718.0 mg, 2.049 mmol, colorless oil (79%).

**<sup>1</sup>H NMR** (500 MHz, CDCl<sub>3</sub>) δ (ppm): 7.41 (d, *J* = 7.1 Hz, 1H), 5.42 (br s, 1H), 3.82-3.46 (m, 18H), 1.44 (s, 9 H). **<sup>13</sup>C NMR** (126 MHz, CDCl<sub>3</sub>) δ (ppm): 169.40, 156.04, 79.85, 72.73, 70.83, 70.78, 70.69, 70.64, 70.53, 70.40, 70.15, 70.10, 70.08, 69.80, 69.07, 61.58, 39.38, 39.34, 28.50. **HRMS QToF-ESI** calculated for C<sub>15</sub>H<sub>31</sub>N<sub>2</sub>O<sub>7</sub> [M+H]<sup>+</sup> *m/z* 351.2131; found *m/z* 351.2148.

*2-amino-N-(2-(2-(2-(2-hydroxyethoxy)ethoxy)ethoxy)ethyl)acetamide (S19)*

**See General Procedure III to generate S19.** Quantities Used: **S18** (480.0 mg, 1.370 mmol), and 6 N HCl in MeOH (13.7 mL, 0.10 M with respect to **S18**). The solution was concentrated by rotary evaporation at 30 °C with a dry ice/EtOAc cold finger rather than subjecting the solution to an aqueous workup (Note: this was done due to the zwitterionic character of the amino-PEG-alcohol product lending poor partitioning into the organic phase during extraction over a pH range of 8 to 12). Crude Yield: 333.6 mg, 1.333 mmol, red oil (97%). Compound **S19** was used to generate **S20** without further purification.

*tert-Butyl (E)-(4-((tert-butoxycarbonyl)(4-(2-cyanovinyl)-2,6-dimethylphenyl)amino)-6-((14-hydroxy-2-oxo-6,9,12-trioxa-3-azatetradecyl)amino)-1,3,5-triazin-2-yl)(4-cyanophenyl)carbamate (S20)*

**See General Procedure I to generate S20.** Quantities Used: **3** (545.0 mg, 0.9052 mmol), **S19** (333.6 mg, 1.333 mmol), distilled THF (9.1 mL, 0.10 M with respect to **3**), and DIPEA (480 μL, 2.72 mmol). The material was purified by flash column chromatography using a gradient of 0-40% acetone in hexanes, then 40-75% acetone in hexanes where the product eluted with 75% acetone in hexanes. The fractions were concentrated by rotary evaporation at 35 °C to afford a solid. The material was dried overnight under reduced pressure to afford **S20**. Yield: 263.4 mg, 0.3228 mmol, white tacky solid (36%).

**<sup>1</sup>H NMR** (500 MHz, CDCl<sub>3</sub>, major and minor rotamer) δ (ppm): 7.84 (d, *J* = 26.1 Hz, 1H), 7.39 (d, *J* = 6.1 Hz, 2H), 7.34 (d, *J* = 16.7 Hz, 1H), 6.98 – 6.95 (m, 4H), 6.73 (d, *J* = 6.3 Hz, 1H), 5.90 (d, *J* = 16.6 Hz, 1H), 4.09 (d, *J* = 4.1 Hz, 2H), 3.78 (p, *J* = 4.1 Hz, 2H), 3.71 – 3.54 (m, 10H), 3.55 (q, *J* = 4.7 Hz, 3H), 3.46 (q, *J* = 5.8 Hz, 4H), 2.17 (br s, 2H), 1.93 (d, *J* = 9.1 Hz, 6H), 1.34 (dd, *J* = 17.4, 10.7 Hz, 18H). **<sup>13</sup>C NMR** (126 MHz, CDCl<sub>3</sub>, major and minor rotamer) δ (ppm): 169.02, 168.96, 166.73, 165.42, 151.42, 151.18, 150.94, 150.81, 149.89, 149.86, 144.54, 141.02, 136.96, 136.79, 132.63, 132.32, 128.92, 128.75, 126.76, 118.62, 118.32, 110.49, 110.36, 96.67, 83.67, 82.78, 72.69, 72.67, 70.81, 70.79, 70.49, 70.15, 70.12, 70.09, 69.93, 61.53, 61.47, 44.86, 44.72, 39.48, 39.44, 27.95, 27.91, 18.00, 17.93. **HRMS QToF-ESI** calculated for C<sub>41</sub>H<sub>54</sub>N<sub>9</sub>O<sub>9</sub> [M+H]<sup>+</sup> *m/z* 816.4044; found *m/z* 816.4028.

*(E)-2-((4-((4-cyanophenyl)amino)-6-((4-(2-cyanovinyl)-2,6-dimethylphenyl)amino)-1,3,5-triazin-2-yl)amino)-N-(2-(2-(2-(2-hydroxyethoxy)ethoxy)ethoxy)ethyl)acetamide (18)*

**See General Procedure III to generate 18.** Quantities Used: **S20** (260.0 mg, 0.3187 mmol), and 6 N HCl in MeOH (3.2 mL, 0.10 M with respect to **S20**). The material was purified by flash column chromatography using a gradient of 0-40% acetone in hexanes then 40-85% acetone in hexanes where the product eluted with 85% acetone in hexanes, then the fractions were concentrated by rotary evaporation at 35 °C to afford a solid. The material was dried overnight under reduced pressure to afford **18**. Yield: 186.0 mg, 0.3021 mmol, white solid (95%).

**<sup>1</sup>H NMR** (500 MHz, DMSO-d<sub>6</sub>, major and minor rotamer) $\delta$ (ppm): 9.57 (d, $J$ = 52.7 Hz, 1H), 8.67 (t, $J$ = 55.5 Hz, 1H), 8.00 – 7.41 (m, 8H), 7.24 (s, 1H), 6.42 (d, $J$ = 16.7 Hz, 1H), 4.58 (t, $J$ = 5.5 Hz, 1H), 3.97 – 3.77 (m, 2H), 3.49 – 3.22 (m, 18H), 2.18 (d, $J$ = 9.5 Hz, 6H). **<sup>13</sup>C NMR** (126 MHz, DMSO-d<sub>6</sub>, major and minor rotamer) $\delta$ (ppm): 169.58, 165.98, 164.98, 164.11, 150.46, 144.98, 139.17, 136.74, 132.50, 131.60, 127.19, 119.58, 119.09, 102.38, 95.94, 72.34, 69.83, 69.74, 69.71, 69.61, 69.56, 69.05, 67.04, 60.21, 43.73, 18.41. **HRMS QToF-ESI** calculated for C<sub>31</sub>H<sub>38</sub>N<sub>9</sub>O<sub>5</sub> [M+H]<sup>+</sup> $m/z$ 616.2996; found $m/z$ 616.2994.

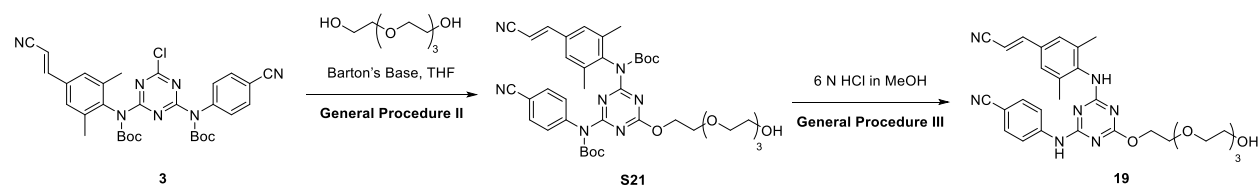

##### Scheme S10: Preparation of Compound 19

*tert*-Butyl (*E*)-4-((*tert*-butoxycarbonyl)(4-(2-cyanovinyl)-2,6-dimethylphenyl)amino)-6-(2-(2-(2-(2-hydroxyethoxy)ethoxy)ethoxy)ethoxy)-1,3,5-triazin-2-yl)(4-cyanophenyl)carbamate (**S21**)

**See General Procedure II to generate S21.** Quantities Used: **3** (200.0 mg, 0.3322 mmol), tetraethylene glycol (350 μL, 2.00 mmol), distilled THF (3.3 mL, 0.10 M with respect to **3**), and Barton's base (140 μL, 0.664 mmol). The material was purified by flash column chromatography using a gradient of 0-40% acetone in hexanes where **S21** eluted with 30% acetone in hexanes. The fractions were concentrated by rotary evaporation at 35 °C and the material was dried overnight with stirring under reduced pressure to afford **S21**. Yield: 131.2 mg, 0.1727 mmol, white tacky solid (52%).

**<sup>1</sup>H NMR** (500 MHz, CDCl<sub>3</sub>) $\delta$ (ppm): 7.43 (d, $J$ = 8.1 Hz, 2H), 7.31 (d, $J$ = 16.7 Hz, 1H), 7.00 (br s, 4H, overlap of Aryl CH and Aryl CH), 5.87 (d, $J$ = 16.7 Hz, 1H), 4.41 (s, 2H), 3.74 – 3.57 (m, 16 h), 2.73 (s, 1H), 2.25 (s, 1H), 1.94 (s, 6H), 1.34 (d, $J$ = 12.0 Hz, 9H). **<sup>13</sup>C NMR** (126 MHz, CDCl<sub>3</sub>) $\delta$ (ppm): 171.58, 167.70, 166.67, 150.82, 150.33, 149.68, 144.06, 140.63, 136.83, 132.89, 132.46, 128.76, 126.85, 118.32, 118.09, 110.80, 96.84, 83.94, 83.14, 72.56, 70.69, 70.63, 70.54, 70.36, 68.74, 67.34, 61.72, 27.80, 17.80. **HRMS QToF-ESI** calculated for C<sub>39</sub>H<sub>50</sub>N<sub>7</sub>O<sub>9</sub> [M+H]<sup>+</sup> $m/z$ 760.3670; found $m/z$ 760.3655.

(*E*)-4-((4-((4-(2-cyanovinyl)-2,6-dimethylphenyl)amino)-6-(2-(2-(2-(2-hydroxyethoxy)ethoxy)ethoxy)ethoxy)-1,3,5-triazin-2-yl)amino)benzonitrile (**19**)

**See General Procedure III to generate 19.** Quantities Used: **S21** (131.2 mg, 0.1727 mmol), and 6 N HCl in MeOH (1.8 mL, 0.10 M with respect to **S21**). The material was purified by flash column chromatography with a gradient of 0-40% acetone in hexanes to remove impurities, then a 40-75% gradient of acetone in hexanes to elute compound **19**. Yield: 22.0 mg, 0.039 mmol, white tacky solid (23%).

**<sup>1</sup>H NMR** (500 MHz, acetone-d<sub>6</sub>, major and minor rotamer) $\delta$ (ppm): 9.14 (s, 1H), 8.66 – 8.51 (m, 1H), 8.13 – 7.49 (m, 7H), 6.34 and 6.25 (each d, $J$ = 16.7 Hz and $J$ = 16.7 Hz, 1H), 4.47 and 4.27 (each t, $J$ = 4.8 Hz and $J$ = 4.9 Hz, 2H), 3.86 and 3.76 (each t, $J$ = 6.1 Hz and $J$ = 4.7 Hz, 2H), 3.62 – 3.59 (m, 13H), 3.53 – 3.51 (m, 5H), 2.93 (s, 1H), 2.28 (s, 6H). **<sup>13</sup>C NMR** (126 MHz, acetone-d<sub>6</sub>, major and minor rotamer) $\delta$ (ppm): 171.86, 167.64, 166.58, 150.90, 144.96, 139.31, 138.91, 138.09, 137.97, 133.66, 133.47, 133.33, 128.07, 128.02, 120.64, 120.24, 119.78, 119.70, 119.15, 105.55, 105.37, 97.19, 97.08, 73.48, 73.46, 71.26, 71.23, 71.21, 71.14, 71.06, 70.84, 69.73, 69.57, 67.01, 66.80, 61.94, 61.87, 18.69. **HRMS QToF-ESI** calculated for C<sub>29</sub>H<sub>34</sub>N<sub>7</sub>O<sub>5</sub> [M+H]<sup>+</sup> $m/z$ 560.2621; found $m/z$ 560.2612.

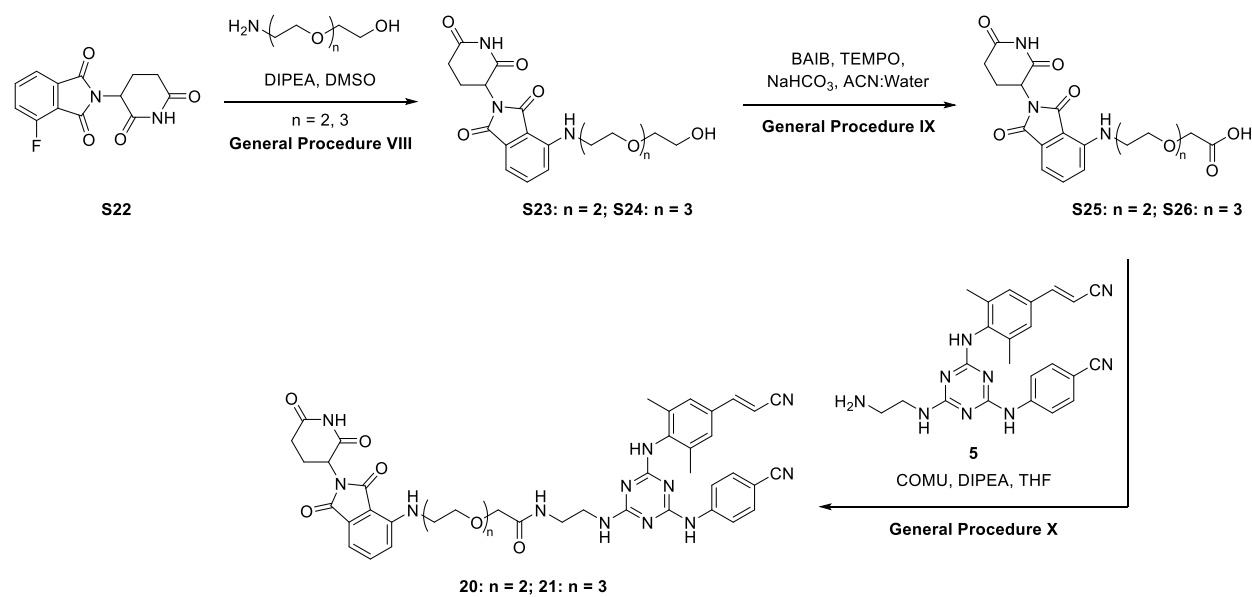

#### Scheme S11: Preparation of Compounds 20-21

**2-(2,6-dioxopiperidin-3-yl)-4-((2-(2-(2-hydroxyethoxy)ethoxy)ethyl)amino)isoindoline-1,3-dione (**S23**)**

**See General Procedure VIII to generate S23.** Quantities Used: **S22** (550.0 mg, 1.991 mmol), 2-(2-(2-aminoethoxy)ethoxy)ethan-1-ol (356.4 mg, 2.389 mmol), DMSO (20.0 mL, 0.10 M with respect to **S22**), and DIPEA (60 μL, 0.30 mmol). Yield: 456.0 mg, 1.125 mmol, yellow/green oil (56%).

**<sup>1</sup>H NMR** (500 MHz, CDCl<sub>3</sub>) $\delta$ (ppm): 8.62 (s, 1H), 7.47 (t, $J$ = 7.8 Hz, 1H), 7.08 (d, $J$ = 7.1 Hz, 1H), 6.89 (d, $J$ = 8.5 Hz, 1H), 6.54 (br s, 1H), 4.91 (dd, $J$ = 12.0, 5.5 Hz, 1H), 3.72 – 3.67 (m, 8H), 3.60 – 3.59 (m, 2H), 3.46 (br s, 2H), 2.87 – 2.68 (m, 3H), 2.32 (br s, 1H), 2.16 – 2.08 (m, 1H). **<sup>13</sup>C NMR**

(126 MHz, CDCl<sub>3</sub>) $\delta$ (ppm): 171.57, 169.57, 168.75, 167.74, 146.89, 136.21, 132.57, 116.89, 111.81, 110.38, 72.75, 70.41, 69.33, 61.84, 48.98, 42.34, 31.50, 22.85. **HRMS QToF-ESI** calculated for C<sub>19</sub>H<sub>23</sub>N<sub>3</sub>O<sub>7</sub> [M+Na]<sup>+</sup> $m/z$ 428.1430; found $m/z$ 428.1434.

*2-(2,6-dioxopiperidin-3-yl)-4-((2-(2-(2-(2-hydroxyethoxy)ethoxy)ethoxy)ethyl)amino)isoindoline-1,3-dione (S24)*

**See General Procedure VIII to generate S24.** Quantities Used: **S22** (550.0 mg, 1.991 mmol), 2-(2-(2-(2-aminoethoxy)ethoxy)ethoxy)ethan-1-ol (461.7 mg, 2.389 mmol), DMSO (20.0 mL, 0.10 M with respect to **S22**), and DIPEA (60 μL, 0.30 mmol). Yield: 544.5 mg, 1.211 mmol, yellow/green oil (61%).

**<sup>1</sup>H NMR** (500 MHz, CDCl<sub>3</sub>) $\delta$ (ppm): 8.87 (s, 1H), 7.47 – 7.45 (m, 1H), 7.06 (d, $J$ = 7.1 Hz, 1H), 6.89 (d, $J$ = 8.5 Hz, 1H), 6.48 (t, $J$ = 5.7 Hz, 1H), 4.91 (dd, $J$ = 12.0, 5.4 Hz, 1H), 3.70 – 3.64 (m, 12H), 3.59 – 3.57 (m, 2H), 3.45 (q, $J$ = 5.5 Hz, 2H), 2.85 – 2.70 (m, 3H), 2.31 (br s, 1H), 2.15 – 2.07 (m, 1H). **<sup>13</sup>C NMR** (126 MHz, CDCl<sub>3</sub>) $\delta$ (ppm): 171.69, 169.32, 168.79, 167.69, 146.81, 136.05, 132.50, 116.83, 111.63, 110.24, 72.51, 70.69, 70.64, 70.51, 70.28, 69.45, 61.64, 48.85, 42.33, 31.41, 22.79. **HRMS QToF-ESI** calculated for C<sub>21</sub>H<sub>27</sub>N<sub>3</sub>O<sub>8</sub> [M+Na]<sup>+</sup> $m/z$ 472.1699; found $m/z$ 472.1696.

*2-(2-(2-((2-(2,6-dioxopiperidin-3-yl)-1,3-dioxoisindolin-4-yl)amino)ethoxy)ethoxy)acetic acid (S25)*

**See General Procedure IX to generate S25.** Quantities Used: **S23** (710.0 mg, 1.751 mmol), BAIB (1.2973 g, 4.0275 mmol), TEMPO (68.4 mg, 0.4378 mmol), sodium bicarbonate (809.2 mg, 9.632 mmol), and 1:1 v/v deionized water:ACN (17.5 mL, 0.10 M with respect to **S23**). Crude Yield: 572.9 mg, 1.366 mmol, yellow/green solid (78%). Compound **S25** was used to generate **20** without further purification.

**<sup>1</sup>H NMR** (500 MHz, DMSO-d<sub>6</sub>, Crude Spectrum) $\delta$ (ppm): 12.56 (br s, 1H), 11.09 (s, 1H), 7.58 (dd, $J$ = 8.6, 7.1 Hz, 1H), 7.15 (d, $J$ = 8.6 Hz, 1H), 7.04 (d, $J$ = 7.0 Hz, 1H), 6.61 (t, $J$ = 5.9 Hz, 1H), 5.05 (dd, $J$ = 12.8, 5.4 Hz, 1H), 4.02 (s, 2H), 3.63-3.57 (m, 6H), 3.47 (q, $J$ = 5.6 Hz, 2H), 2.88 (ddd, $J$ = 16.7, 13.7, 5.4 Hz, 1H), 2.60-2.53 (m, 2H), 2.05-2.01 (m, 1H). **<sup>13</sup>C NMR** (126 MHz, DMSO-d<sub>6</sub>, Crude Spectrum) $\delta$ (ppm): 172.81, 171.66, 170.09, 168.93, 167.30, 146.41, 136.23, 132.10, 117.46, 110.67, 109.25, 69.87, 69.65, 68.88, 67.61, 48.55, 41.69, 30.98, 22.13. **HRMS QToF-ESI** calculated for C<sub>19</sub>H<sub>22</sub>N<sub>3</sub>O<sub>8</sub> [M+Na]<sup>+</sup> $m/z$ 420.1407; found $m/z$ 420.1407.

*2-(2-(2-(2-((2-(2,6-dioxopiperidin-3-yl)-1,3-dioxoisindolin-4-yl)amino)ethoxy)ethoxy)ethoxy)acetic acid (S26)*

**See General Procedure IX to generate S26.** Quantities Used: **S24** (620.0 mg, 1.379 mmol), BAIB (1.0220 g, 3.1731 mmol), TEMPO (53.9 mg, 0.345 mmol), sodium bicarbonate (637.2 mg, 7.585 mmol), and 1:1 v/v deionized water:ACN (13.8 mL, 0.10 M with respect to **S24**). Crude

Yield: 377.7 mg, 0.8150 mmol, yellow/orange solid (59%). Compound **S26** was used to generate **21** without further purification.

**<sup>1</sup>H NMR** (500 MHz, DMSO-d<sub>6</sub>, Crude Spectrum) $\delta$ (ppm): 12.53 (br s, 1H), 11.09 (s, 1H), 7.58 (dd, $J$ = 8.6, 7.1 Hz, 1H), 7.15 (d, $J$ = 8.6 Hz, 1H), 7.04 (d, $J$ = 7.0 Hz, 1H), 6.61 (t, $J$ = 5.9 Hz, 1H), 5.05 (dd, $J$ = 12.8, 5.4 Hz, 1H), 4.00 (s, 2H), 3.63-3.45 (m, 12H), 2.88 (ddd, $J$ = 16.8, 13.7, 5.4 Hz, 1H), 2.61-2.53 (m, 2H), 2.04-2.01 (m, 1H). **<sup>13</sup>C NMR** (126 MHz, DMSO-d<sub>6</sub>, Crude Spectrum) $\delta$ (ppm): 172.81, 171.65, 170.08, 168.93, 167.30, 146.41, 136.23, 132.10, 117.46, 110.67, 109.25, 69.83, 69.79, 69.75, 68.88, 67.57, 48.56, 41.69, 30.98, 22.13. **HRMS QToF-ESI** calculated for C<sub>21</sub>H<sub>26</sub>N<sub>3</sub>O<sub>9</sub> [M+Na]<sup>+</sup> $m/z$ 464.1674; found $m/z$ 464.1669.

*(E)-N-(2-((4-((4-cyanophenyl)amino)-6-((4-(2-cyanovinyl)-2,6-dimethylphenyl)amino)-1,3,5-triazin-2-yl)amino)ethyl)-2-(2-(2-((2-(2,6-dioxopiperidin-3-yl)-1,3-dioxoisindolin-4-yl)amino)ethoxy)ethoxy)acetamide (20)*

**See General Procedure X to generate 20.** Quantities Used: **5** (100.0 mg, 0.2350 mmol), **S25** (108.4 mg, 0.2585 mmol), COMU (130.8 mg, 0.3055 mmol), distilled THF (2.4 mL, 0.10 M with respect to **5**), and DIPEA (140 μL, 0.776 mmol). Yield: 157.2 mg, 0.1901 mmol, yellow solid (81%)

**<sup>1</sup>H NMR** (500 MHz, acetone-d<sub>6</sub>, major and minor rotamer) $\delta$ (ppm): 11.33 (d, $J$ = 50.8 Hz, 1H), 8.83 (br s, 1H), 8.28 (br s, 1H), 7.77 – 7.34 (m, 8H), 7.00 (d, $J$ = 6.6 Hz, 2H), 6.55 (br s, 1H), 6.26 (br s, 1H), 5.13 – 5.11 (m, 1H), 3.89 (s, 2H), 3.68 – 3.44 (m, 12H), 2.98 – 2.79 (m, 3H), 2.24 (d, $J$ = 16.5 Hz, 7H), 1.41 – 1.20 (m, 1H). **<sup>13</sup>C NMR** (126 MHz, acetone-d<sub>6</sub>, major and minor rotamer) $\delta$ (ppm): 174.43, 171.75, 170.86, 170.13, 168.13, 167.23, 165.28, 164.96, 150.93, 150.88, 147.54, 145.49, 145.38, 139.59, 137.83, 136.81, 133.34, 132.88, 127.91, 119.92, 119.22, 117.74, 111.65, 110.91, 104.68, 96.76, 79.14, 71.47, 71.44, 71.09, 70.65, 69.89, 68.00, 49.66, 42.79, 41.26, 39.62, 31.90, 26.09, 23.35, 18.86. **HRMS QToF-ESI** calculated for C<sub>42</sub>H<sub>43</sub>N<sub>12</sub>O<sub>7</sub> [M+H]<sup>+</sup> $m/z$ 827.3378; found $m/z$ 827.3341.

*(E)-N-(2-((4-((4-cyanophenyl)amino)-6-((4-(2-cyanovinyl)-2,6-dimethylphenyl)amino)-1,3,5-triazin-2-yl)amino)ethyl)-2-(2-(2-(2-((2-(2,6-dioxopiperidin-3-yl)-1,3-dioxoisindolin-4-yl)amino)ethoxy)ethoxy)ethoxy)acetamide (21)*

**See General Procedure X to generate 21.** Quantities Used: **5** (110.0 mg, 0.2585 mmol), **S26** (131.7 mg, 0.2844 mmol), COMU (143.9 mg, 0.3361 mmol), distilled THF (2.6 mL, 0.10 M with respect to **5**), and DIPEA (150 μL, 0.856 mmol). Yield: 184.8 mg, 0.2122 mmol, yellow/orange solid (82%).

**<sup>1</sup>H NMR** (500 MHz, acetone-d<sub>6</sub>, major and minor rotamer) $\delta$ (ppm): 11.35 – 10.91 (m, 1H), 8.92 (d, $J$ = 62.7 Hz, 1H), 8.26 – 8.05 (m, 1H), 7.77 – 7.32 (m, 8H), 7.00 (d, $J$ = 7.3 Hz, 2H), 6.55 (br s, 1H), 6.26 (br s, 1H), 5.13 (dd, $J$ = 12.7, 5.4 Hz, 1H), 3.89 (d, $J$ = 18.5 Hz, 2H), 3.67 – 3.21 (m, 16H), 2.99 – 2.93 (m, 3H), 2.82 – 2.76 (m, 2H), 2.24 (d, $J$ = 18.1 Hz, 7H), 1.41 – 1.20 (m, 1H). **<sup>13</sup>C NMR** (126 MHz, acetone-d<sub>6</sub>, major and minor rotamer) $\delta$ (ppm): 173.74, 171.07, 170.46, 169.45, 167.52, 166.68, 165.41, 164.67, 164.38, 150.28, 146.92, 144.90, 144.80, 139.00, 137.23,

136.18, 132.72, 127.27, 119.28, 118.59, 117.09, 110.93, 110.20, 104.00, 96.12, 78.51, 70.94, 70.90, 70.47, 70.33, 70.29, 69.28, 69.25, 54.76, 48.99, 42.19, 40.55, 39.03, 31.25, 22.74, 18.21.
**HRMS QToF-ESI** calculated for $C_{44}H_{46}N_{12}O_8Na$ $[M+Na]^+$ $m/z$ 893.3459; found $m/z$ 893.3419.

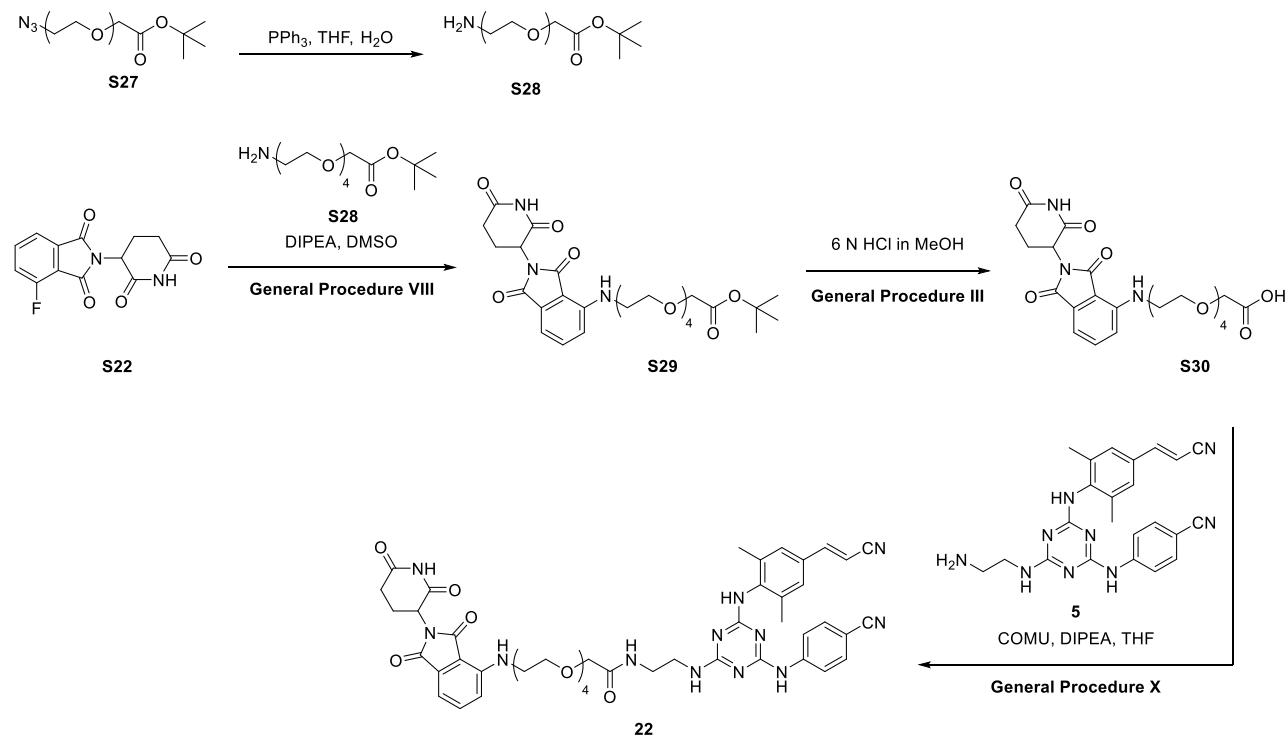

### Scheme S12: Preparation of Compound 22

#### *tert*-Butyl 14-amino-3,6,9,12-tetraoxatetradecanoate (**S28**)

To a flame-dried round-bottom flask equipped with a magnetic stir bar was added **S27** (487.0 mg, 1.461 mmol), $PPh_3$ (459.5 mg, 1.752 mmol), and THF (6.6 mL, 0.20 M with respect to **S27**). Once the solution was homogeneous, deionized water (0.70 mL) was added and the solution was maintained at room temperature for 48 h, at which time the reaction was judged complete by TLC (1:1 hexanes:EtOAc). The solvent was removed by rotary evaporation at 35 °C to afford an oil. The oil was reconstituted in EtOAc, and the solution was washed with aqueous $NH_4Cl$ (3 x 2 mL). The pH of the aqueous phase was adjusted to 10 with aqueous 1 N NaOH and back extracted with $CHCl_3$ (5 x 3 mL). The combined organic phase was dried with magnesium sulfate, filtered through a pad of Celite using $CHCl_3$, and concentrated by rotary evaporation at 35 °C to afford **S28**. Crude Yield: 281.0 mg, mmol, 0.9142 mmol, light-yellow oil (63%). Compound **S28** was used to generate **S29** without further purification.

Characterization data matches that reported previously.<sup>4</sup>

**<sup>1</sup>H NMR** (500 MHz, CDCl<sub>3</sub>) δ (ppm): 4.01 (s, 2H), 3.69-3.62 (m, 12H), 3.51 (t, *J* = 5.19, 2H), 2.86 (t, *J* = 5.24, 2H), 1.74 (s, 2H), 1.46 (s, 9H). **<sup>13</sup>C NMR** (126 MHz, CDCl<sub>3</sub>) δ (ppm): 169.82, 81.70, 73.35, 70.82, 70.72, 70.70, 70.68, 70.66, 70.38, 69.15, 41.89, 28.23.

*tert*-Butyl 14-((2-(2,6-dioxopiperidin-3-yl)-1,3-dioxoisindolin-4-yl)amino)-3,6,9,12-tetraoxatetradecanoate (**S29**)

**See General Procedure VIII to generate S29.** Quantities Used: **S22** (90.0 mg, 0.325 mmol), **S28** (120.0 mg, 0.3904 mmol), anhydrous DMSO (3.3 mL, 0.10 M with respect to **S22**), and DIPEA (110 μL, 0.651 mmol). Yield: 101 mg, 0.180 mmol, yellow/green oil (55%).

**<sup>1</sup>H NMR** (500 MHz, CDCl<sub>3</sub>) δ (ppm): 8.67 (s, 1H), 7.45 (dd, *J* = 8.6, 7.1 Hz, 1H), 7.06 (d, *J* = 7.1 Hz, 1H), 6.89 (d, *J* = 8.5 Hz, 1H), 6.46 (t, *J* = 5.6 Hz, 1H), 4.89 (dd, *J* = 12.0, 5.4 Hz, 1H), 3.99 (s, 2H), 3.70 – 3.63 (m, 14H), 3.44 (q, *J* = 5.5 Hz, 2H), 2.85 – 2.69 (m, 3H), 2.14 – 2.07 (m, 2H), 1.43 (s, 9H). **<sup>13</sup>C NMR** (126 MHz, CDCl<sub>3</sub>) δ (ppm): 171.47, 169.75, 169.32, 168.62, 167.70, 146.88, 136.05, 132.55, 132.18, 132.10, 128.59, 128.50, 116.84, 111.63, 110.31, 81.60, 70.78, 70.74, 70.65, 70.63, 70.54, 69.52, 69.03, 48.91, 42.42, 31.47, 28.15, 22.82. **HRMS QToF-ESI** calculated for C<sub>27</sub>H<sub>38</sub>N<sub>3</sub>O<sub>10</sub> [M+H]<sup>+</sup> *m/z* 564.2557; found *m/z* 564.2556.

14-((2-(2,6-dioxopiperidin-3-yl)-1,3-dioxoisindolin-4-yl)amino)-3,6,9,12-tetraoxatetradecanoic acid (**S30**)

**See General Procedure III to generate S30.** Quantities Used: **S29** (94.8 mg, 0.168 mmol), and 6 N HCl in MeOH (1.7 mL, 0.10 M with respect to **S29**). The extraction was performed cold (ice chips) at pH 2 to extract the acid product. Crude Yield: 83.7 mg, 0.165 mmol, yellow oil (98%). Compound **S30** was used to generate **22** without further purification.

(*E*)-*N*-(2-((4-((4-cyanophenyl)amino)-6-((4-(2-cyanovinyl)-2,6-dimethylphenyl)amino)-1,3,5-triazin-2-yl)amino)ethyl)-14-((2-(2,6-dioxopiperidin-3-yl)-1,3-dioxoisindolin-4-yl)amino)-3,6,9,12-tetraoxatetradecanamide (**22**)

**See General Procedure X to generate 22.** Quantities Used: **5** (64.0 mg, 0.150 mmol), **S30** (83.9 mg, 0.165 mmol), COMU (84.0 mg, 0.195 mmol), distilled THF (1.5 mL, 0.10 M with respect to **5**), and DIPEA (90 μL, 0.50 mmol). Yield: 20.0 mg, 0.0219 mmol, yellow solid (15% over two steps)

**<sup>1</sup>H NMR** (500 MHz, acetone-d<sub>6</sub>, major and minor rotamer) δ (ppm): 11.20 (br s, 1H), 8.81 (s, 1H), 8.09 (s, 1H), 7.83 – 7.46 (m, 6H), 7.07 (d, *J* = 20.8 Hz, 2H), 6.75 (s, 1H), 6.61 (s, 1H), 6.28, (s, 1H), 5.72 (s, 1H), 5.11 (s, 1H), 3.84 – 3.59 (m, 18H), 3.00 (m, 2H), 2.28 and 2.25 (each s, 6H), 143-120 (m, 4H). **<sup>13</sup>C NMR** (126 MHz, acetone-d<sub>6</sub>, major and minor rotamer) δ (ppm): 173.89, 170.97, 170.22, 168.17, 167.47, 166.19, 165.48, 165.21, 151.01, 150.99, 147.73, 145.60, 139.80, 138.04, 136.92, 133.54, 133.37, 133.00, 127.98, 119.98, 119.22, 117.86, 111.56, 110.99, 104.71, 96.85, 79.19, 71.58, 71.25, 71.20, 71.08, 71.02, 70.92, 70.79, 70.03, 70.00, 55.45, 49.72, 42.94,

41.21, 39.72, 31.91, 23.49, 18.85. **HRMS QToF-ESI** calculated for $C_{46}H_{51}N_{12}O_9$ $[M+H]^+$ $m/z$ 915.3902; found $m/z$ 915.3885.

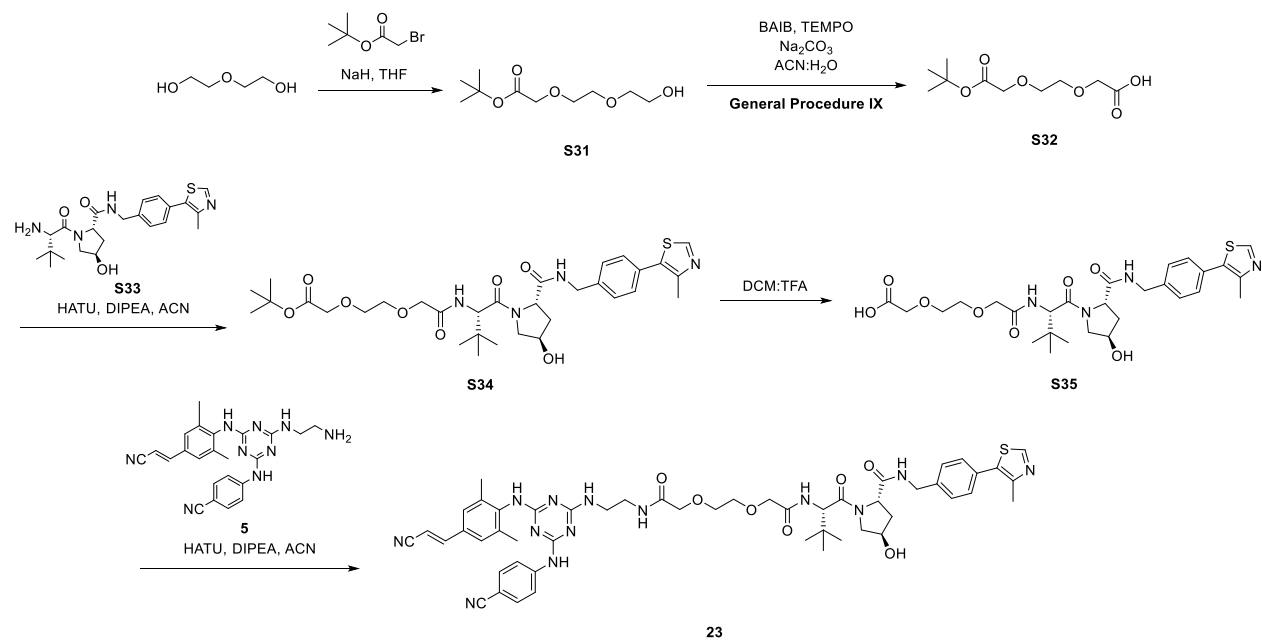

#### Scheme S13: Preparation of Compound 23

##### *tert*-Butyl 2-(2-(2-hydroxyethoxy)ethoxy)acetate (**S31**)

To a flame-dried round-bottom flask equipped with a magnetic stir bar under inert atmosphere ($N_2$) was added a suspension of 60% NaH in mineral oil (745.0 mg, 18.63 mmol). The mineral oil was removed from the NaH by the addition of portions of distilled THF (5.0 mL) as follows. The suspension was stirred for 5 minutes then allowed to stand until the solid NaH settled on the bottom of the flask. The THF and mineral oil were decanted by syringe, and the process was repeated a total of three times. To the flask was added distilled THF (94 mL, 0.15 M with respect to *tert*-butyl 2-bromoacetate). The solution was cooled in an ice bath, and diethylene glycol (4.0000 g, 37.693 mmol) was added dropwise by syringe (Note: gas evolution). The ice bath was maintained for 10 minutes until effervescence was no longer observed. *tert*-Butyl 2-bromoacetate (2.26 mL, 14.1 mmol) was added, and the ice bath was allowed to gradually warm to room temperature over 1 h at which time the reaction was judged complete by TLC (1:1 hexanes:EtOAc). The reaction was quenched with an aqueous solution of $NH_4Cl$ (0.2 mL). The solution was stirred for 15 minutes, transferred to a round bottom flask with $CHCl_3$, and concentrated by rotary evaporation at 35 °C. The concentrate was reconstituted in diethyl ether, transferred to a separatory funnel, and washed with brine (3 x 3 mL). The combined organic phase was dried with magnesium sulfate, filtered through a pad of Celite using diethyl ether, and concentrated by rotary evaporation at 30 °C to afford **S31**. Crude Yield: 1.24 g, 5.63 mmol, colorless oil (40%). Compound **S31** was used to generate **S32** without further purification.

Characterization data identical to that previously reported.<sup>5</sup>

**<sup>1</sup>H NMR** (500 MHz, CDCl<sub>3</sub>) δ (ppm): 4.01 (s, 2H), 3.73-3.71 (m, 8H), 3.62 (t, *J* = 4.2 Hz, 2H), 2.47 (br s, 1H), 1.47 (s, 9H). **<sup>13</sup>C NMR** (126 MHz, CDCl<sub>3</sub>) 169.75, 81.90, 72.70, 70.99, 70.49, 69.11, 61.88, 28.24.

*2-(2-(2-(tert-butoxy)-2-oxoethoxy)ethoxy)acetic acid (S32)*

**See General Procedure IX to generate S32.** Quantities Used: BAIB (531.5 mg, 1.650 mmol), TEMPO (23.4 mg, 0.150 mmol), sodium bicarbonate (346.5 mg, 4.125 mmol), and 1:1 v/v ACN:deionized water (1.5 mL, 0.50 M with respect to **S31**). The solution was chilled in an ice bath for 10 minutes before **S31** (250.0 mg, 1.135 mmol) was added. The reaction was judged complete by TLC at 18 h (1:1 hexanes:EtOAc). Crude Yield: 167.3 mg, 0.7142 mmol, light yellow oil (63%). Compound **S32** was used to generate **S34** without further purification.

Characterization data identical to that previously reported.<sup>6</sup>

**<sup>1</sup>H NMR** (500 MHz, CDCl<sub>3</sub>) δ (ppm): 7.70 (s, 1H), 4.19 (s, 2H), 4.03 (s, 2H), 3.79-3.78 (m, 2H), 3.75-3.74 (m, 2H), 1.47 (s, 9H). **<sup>13</sup>C NMR** (126 MHz, CDCl<sub>3</sub>) δ (ppm): 172.95, 169.56, 82.23, 71.42, 70.70, 69.06, 68.83, 28.24.

*tert-Butyl 2-(2-(2-(((S)-1-((2S,4R)-4-hydroxy-2-((4-(4-methylthiazol-5-yl)benzyl)carbamoyl)pyrrolidin-1-yl)-3,3-dimethyl-1-oxobutan-2-yl)amino)-2-oxoethoxy)ethoxy)acetate (S34)*

To a flame-dried 25 mL round-bottom flask equipped with a magnetic stir bar under inert atmosphere (N<sub>2</sub>) was added **S32** (75.8 mg, 0.290 mmol), HATU (145.6 mg, 0.3829 mmol), and anhydrous ACN (1.6 mL, 0.18 M with respect to **S32**). The solution was chilled in an ice/acetone bath, and the contents of the flask were stirred for 5 minutes before DIPEA (170 μL, 0.976 mmol) was added dropwise. The bath was maintained for 30 min, then allowed to gradually warm to room temperature with subsequent addition of **S33** (125.8 mg, 0.2922 mmol). At 2 h, the reaction was judged complete by TLC (5% MeOH in DCM). The reaction was quenched by adding deionized water (1.5 mL), and the aqueous phase was extracted with EtOAc (5 x 1 mL). The combined organic phase was washed with an aqueous solution of sodium bicarbonate (1 x 2 mL) dried with sodium sulfate, filtered through a fritted funnel using EtOAc, and concentrated by rotary evaporation at 35 °C to afford an oil. The oil was purified by Combiflash using a silica cartridge of 24 g and a gradient of 3-10% MeOH in DCM. The fractions were concentrated by rotary evaporation at 35 °C and further dried under reduced pressure to afford **S34**. Yield: 125 mg, 0.193 mmol, off-white solid (66%).

**<sup>1</sup>H NMR** (500 MHz, CD<sub>3</sub>CN, major and minor rotamer) δ (ppm): 8.73 (s, 1H), 7.42 (d, *J* = 8.6 Hz, 2H), 7.39 (d, *J* = 8.6 Hz, 2H), 7.28 (d, *J* = 9.4 Hz, 1H), 7.20 (t, *J* = 6.3 Hz, 1H), 4.60 (d, *J* = 9.3 Hz, 1H), 4.51-4.44 (m, 2H), 4.41 (br s, 1H), 4.30 (dd, *J* = 15.6, 5.7 Hz, 1H), 4.02 (s, 1H), 4.00 (s, 1H), 3.96 (s, 2H), 3.78 (br s, 0.3H), 3.76 (br s, 0.6H), 3.70-3.66 (m, 5H), 3.29 (d, *J* = 4.2 Hz, 1H), 2.46 (s, 3H), 2.10 (dd, *J* = 8.4, 3.6 Hz, 2H), 1.42 (s, 9H), 1.33-1.30 (m, 1H), 0.96 (s, 9H). **<sup>13</sup>C NMR** (126

MHz, CD<sub>3</sub>CN, major and minor rotamer) $\delta$ (ppm): 172.69, 171.40, 170.60, 170.25, 151.64, 149.39, 140.28, 132.40, 131.48, 130.10, 128.63, 81.97, 71.93, 71.08, 70.87, 70.76, 69.49, 60.09, 57.62, 57.36, 55.33, 43.17, 38.34, 36.42, 28.28, 26.68, 16.42. **HRMS QToF-ESI** calculated for C<sub>32</sub>H<sub>47</sub>N<sub>4</sub>O<sub>8</sub>S [M+H<sup>+</sup>] $m/z$ 647.3115; found $m/z$ 647.3108.

*2-(2-(2-(((S)-1-((2S,4R)-4-hydroxy-2-((4-(4-methylthiazol-5-yl)benzyl)carbamoyl)pyrrolidin-1-yl)-3,3-dimethyl-1-oxobutan-2-yl)amino)-2-oxoethoxy)ethoxy)acetic acid (S35)*

In a 1-neck 50 mL round-bottom flask equipped with a magnetic stir bar in an ice bath was added **S34** (44.2 mg, 0.0682 mmol) and TFA:DCM (0.5:1.5 v/v, 5.7 mL, 0.01 M with respect to **S34**). The bath was maintained for 1.5 h, at which time the reaction was judged complete by TLC (5% MeOH in DCM). The volatile residues were removed by rotary evaporation at 33 °C, and the resulting oil was dried overnight under reduced pressure to give the corresponding acid **S35** (TFA salt). Crude Yield: 47.1 mg, 0.0669 mmol, light brown oil (98%). Compound **S35** was used to generate **23** without further purification.

*(2S,4R)-1-((S)-2-(tert-butyl)-14-(((4-(4-cyanophenyl)amino)-6-((4-((E)-2-cyanovinyl)-2,6-dimethylphenyl)amino)-1,3,5-triazin-2-yl)amino)-4,11-dioxo-6,9-dioxo-3,12-diazatetradecanoyl)-4-hydroxy-N-(4-(4-methyl-thiazol-5-yl)benzyl)pyrrolidine-2-carboxamide (23)*

To a flame-dried 25 mL round-bottom flask equipped with a magnetic stir bar under inert atmosphere (N<sub>2</sub>) was added **S35** (40.3 mg, 0.0680 mmol) and DMF (350 μL, 0.20 M with respect to **S35**), followed by DIPEA (50 μL, 0.28 mmol) at -45 °C. The solution was transferred to an ice/acetone bath for 30 minutes before HATU (52.5 mg, 0.138 mmol) was added. The bath was maintained for 20 minutes before **5** (29.6 mg, 0.0683 mmol) was added, and the ice bath was maintained for another 30 minutes. The solution was gradually warmed to room temperature over 19 h, at which time the reaction was judged complete by TLC (5% MeOH in DCM). The reaction was quenched with deionized water (0.65 mL), and the aqueous phase was extracted with EtOAc (4 x 1 mL). The combined organic phase was washed with an aqueous solution of sodium bicarbonate (1 x 1 mL), dried with sodium sulfate, filtered through a fritted funnel using EtOAc, and concentrated by rotary evaporation at 35 °C. The material was purified by Combiflash using a gradient of 3-10% MeOH in DCM to afford **23**. Yield: 42.9 mg, 0.0726 mmol, off-white solid (63% over 2 steps). **R<sub>f</sub>**: 0.45 (5% MeOH/CH<sub>2</sub>Cl<sub>2</sub>)

**<sup>1</sup>H NMR** (500 MHz, DMSO-d<sub>6</sub>, major and minor rotamer, OH signal not evident by <sup>1</sup>H NMR) $\delta$ (ppm): 9.57-9.40 (m, 2H), 8.98 and 8.97 (s, 1H), 8.58 (t, $J$ = 6.0 Hz, 1H), 8.05 (br s, 1H), 7.89 (br s, 1H), 7.81-7.78 (m, 2H), 7.68 (br s, 1H), 7.61-7.56 (m, 2H), 7.51-7.45 (m, 2H), 7.43-7.36 (m, 6H), 7.13-7.08 (m, 1H), 6.41 (d, $J$ = 16.3 Hz, 1H), 5.16 (d, $J$ = 3.5 Hz, 1H), 4.58 (d, $J$ = 9.5 Hz, 1H), 4.45 (t, $J$ = 8.3 Hz, 1H), 4.36 – 4.26 (m, 4H), 3.99 – 3.92 (m, 5H), 3.68 – 3.60 (m, 7H), 3.48-3.99 (m, 2H), 2.43 (s, 3H), 2.17 (s, 6H), 2.08-2.04 (m, 1H), 1.90 (ddd, $J$ = 13.0, 8.6, 4.4 Hz, 1H), 0.93 (s, 9H). **<sup>13</sup>C NMR** (126 MHz, DMSO-d<sub>6</sub>, major and minor rotamer) $\delta$ (ppm): 171.69, 169.44, 169.23, 168.59, 166.00, 165.78, 164.20, 163.97, 151.43, 150.42, 147.75, 145.07, 139.37, 136.74, 132.49, 131.54, 131.11, 129.72, 128.87, 128.70, 128.13, 127.45, 127.14, 119.56, 119.04, 118.82, 102.32, 102.20, 95.90, 70.17, 70.10, 69.99, 68.84, 58.72, 56.61, 55.74, 41.68, 40.09, 37.96,

35.70, 26.17, 18.38, 18.35, 15.92. **HRMS QToF-ESI:** calculated for $C_{51}H_{60}N_{13}O_7S$ $[M+H]^+$ $m/z$ 998.4459; found $m/z$ 998.4465.

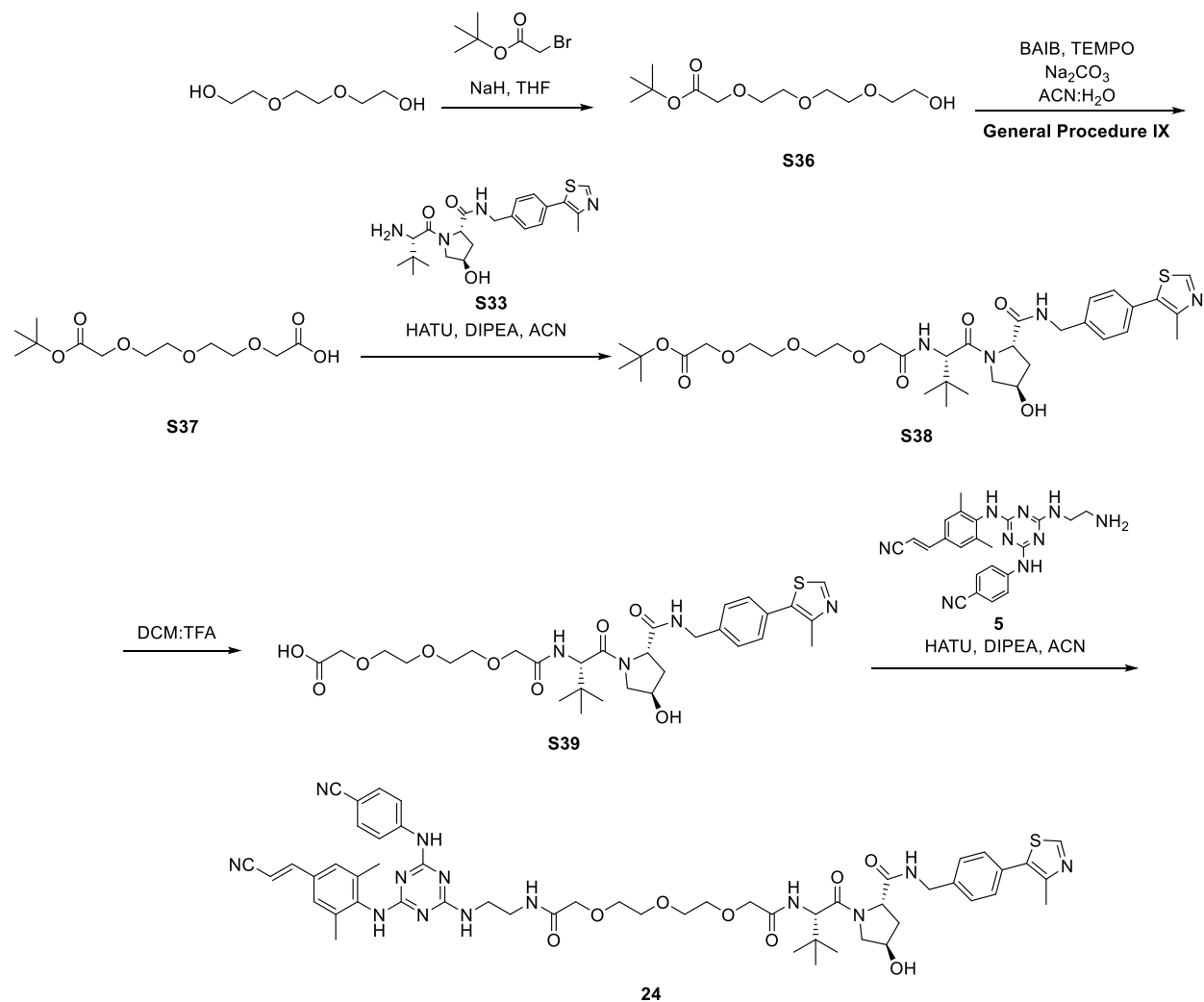

### Scheme S14: Preparation of Compound 24

#### *tert*-Butyl 2-(2-(2-(2-hydroxyethoxy)ethoxy)ethoxy)acetate (**S36**)

To a flame-dried round-bottom flask equipped with a magnetic stir bar under inert atmosphere ($N_2$) was added a suspension of 60% NaH in mineral oil (480.0 mg, 11.95 mmol). The mineral oil was removed from the NaH by the addition of portions of distilled THF (5.0 mL) as follows. The suspension was stirred for 5 minutes then allowed to stand until the solid NaH settled on the bottom of the flask. The THF and mineral oil were decanted by syringe, and the process was repeated a total of three times. To the flask was added distilled THF (51 mL, 0.20 M with respect to *tert*-butyl 2-bromoacetate) and the solution was chilled in an ice bath. Triethylene glycol (3.0935 g, 20.600 mmol) was added to the solution dropwise (Note: gas evolution), and the ice bath was maintained for 10 minutes. *tert*-Butyl 2-bromoacetate (1.50 mL, 10.3 mmol) was added, and the ice bath was allowed to gradually warm to room temperature over 1 h, at which time the reaction was judged complete by TLC (1:1 hexanes:EtOAc). The reaction was quenched with aqueous NH<sub>4</sub>Cl (0.2 mL), stirred for 15 minutes, then concentrated by rotary evaporation at 25 °C. The concentrate was reconstituted in diethyl ether and transferred to a separatory funnel, and the solution was washed with brine (3 x 3 mL). The combined organic phase was dried with magnesium sulfate, filtered through a pad of Celite using diethyl ether, and concentrated by rotary evaporation at 25 °C to afford **S36**. Crude Yield: 1.01 g, 3.81 mmol, clear, colorless oil (37%). Compound **S36** was used to generate **S37** without further purification.

Characterization data matches that previously reported.<sup>5</sup>

**<sup>1</sup>H NMR** (500 MHz, CDCl<sub>3</sub>) δ (ppm): 3.92 (s, 2H), 3.60-3.57 (m, 12H), 2.90 (br s, 1H), 1.38 (s, 9H). **<sup>13</sup>C NMR** (126 MHz, CDCl<sub>3</sub>) δ (ppm): 169.56, 81.54, 72.52 70.56, 70.50, 70.46, 70.36, 70.11, 68.82, 61.47, 27.97.

##### *13,13-dimethyl-11-oxo-3,6,9,12-tetraoxatetradecanoic acid (S37)*

**See General Procedure IX to generate S37.** Quantities Used: BAIB (1.800 g, 5.588 mmol), TEMPO (79.3 mg, 0.508 mmol), sodium bicarbonate (853.0 mg, 10.15 mmol), and a solution of 1:1 v/v ACN:deionized water (5.2 mL, 0.50 M with respect to **S36**). The solution was chilled in an ice bath for 10 minutes before **S36** (671.4 mg, 2.540 mmol) was added. The reaction was judged complete by TLC at 18 h (1:1 hexanes:EtOAc). Crude Yield: 239.3 mg, 0.8599 mmol, light yellow oil (34%). Compound **S37** was used to generate **S38** without further purification.

Characterization data matches that previously reported.<sup>6</sup>

**<sup>1</sup>H NMR** (500 MHz, CDCl<sub>3</sub>) δ (ppm): 9.95 (s, 1H), 4.16 (s, 2H), 4.02 (s, 2H), 3.79-3.71 (m, 8H), 1.48 (s, 9H). **<sup>13</sup>C NMR** (126 MHz, CDCl<sub>3</sub>) 173.47, 169.93, 81.86, 71.12, 70.55, 70.52, 70.43, 68.96, 68.61, 28.08.

##### *tert-Butyl (S)-13-((2S,4R)-4-hydroxy-2-((4-(4-methylthiazol-5-yl)benzyl)carbamoyl)pyrrolidine-1-carbonyl)-14,14-dimethyl-11-oxo-3,6,9-trioxa-12-azapentadecanoate (S38)*

To a flame-dried 25 mL two-neck round-bottom flask equipped with a magnetic stir bar under inert atmosphere (N<sub>2</sub>) was added a solution of acid **S37** (139.2 mg, 0.5000 mmol), HATU (247.9 mg, 0.6520 mmol) and anhydrous ACN (2.6 mL, 0.20 M with respect to **S37**). The solution was chilled in an ice/acetone bath for 5 minutes before DIPEA (300 µL, 1.70 mmol) was added dropwise. The bath was maintained for 30 minutes before adding **S33** (217.4 mg, 0.5050 mmol). The solution was gradually warmed to room temperature over 19 h, at which time the reaction was judged complete by TLC (5% MeOH in DCM). The reaction was quenched with deionized water (2.5 mL), and the aqueous phase was extracted with EtOAc (5 x 2 mL). The combined organic phase was washed with an aqueous solution of sodium bicarbonate (1 x 2 mL), the combined organic phase was dried with sodium sulfate, filtered through a fritted funnel using EtOAc, and concentrated by rotary evaporation at 35 °C. The material was purified by Combiflash using a gradient of 5% MeOH in DCM and concentrated by rotary evaporation at 35 °C to afford **S38**. Yield: 254.6 mg, 0.3685 mmol, off-white solid (74%).

**<sup>1</sup>H NMR** (500 MHz, CD<sub>3</sub>CN, major and minor rotamer) $\delta$ (ppm): 8.74 (s, 1H), 7.42 (d, $J$ = 8.5 Hz, 2H), 7.39 (d, $J$ = 8.5 Hz, 2H), 7.25-7.19 (m, 2H), 4.60 (d, $J$ = 9.4 Hz, 1H), 4.50-4.46 (m, 2H), 4.42 (br s, 1H), 4.31 (dd, $J$ = 15.5, 5.7 Hz, 1H), 3.96-3.95 (m, 2H), 3.93 (s, 2H), 3.78 (br s, 0.4H), 3.76 (br s, 0.7H), 3.69 (dd, $J$ = 11.0, 3.9 Hz, 1H), 3.66-3.59 (m, 10H), 3.36 (d, $J$ = 4.1 Hz, 1H), 2.46 (s, 3H), 2.11-2.09 (m, 2H), 1.43 (s, 9H), 0.96 (s, 9H). **<sup>13</sup>C NMR** (126 MHz, CD<sub>3</sub>CN, major and minor rotamer) $\delta$ (ppm): 172.72, 171.37, 170.70, 170.26, 151.67, 149.38, 140.26, 132.40, 131.47, 130.10, 129.00, 128.66, 81.97, 71.92, 71.28, 71.12, 70.91, 70.86, 70.76, 69.53, 60.12, 57.64, 57.32, 43.19, 38.34, 36.49, 28.29, 26.81, 26.71, 16.43. **HRMS QToF-ESI** calculated for C<sub>34</sub>H<sub>51</sub>N<sub>4</sub>O<sub>9</sub>S [M+H<sup>+</sup>] $m/z$ 691.3377; found: 691.3370.

*(S)-13-((2S,4R)-4-hydroxy-2-((4-(4-methylthiazol-5-yl)benzyl)carbamoyl)pyrrolidine-1-carbonyl)-14,14-dimethyl-11-oxo-3,6,9-trioxa-12-azapentadecanoic acid carboxamide (S39)*

To a one-neck 50 mL round-bottom flask equipped with a magnetic stir bar in an ice bath was added **S38** (70.4 mg, 0.102 mmol) and TFA:DCM (0.5:1.5 v/v, 8.5 mL, 0.01 M with respect to **S38**). The bath was maintained for 1 h, at which time the reaction was judged complete by TLC (5% MeOH in DCM). The volatile residues were removed by rotary evaporation at 33 °C, and the resultant oil was dried overnight under reduced pressure to give the corresponding acid **S39** (TFA salt). Crude Yield: 75.8 mg, 0.101 mmol, light brown oil (99%). Compound **S39** was used to generate **24** without further purification.

*(2S,4R)-1-((S)-2-(tert-butyl)-17-(((4-((4-cyanophenyl)amino)-6-((4-((E)-2-cyanovinyl)-2,6-dimethylphenyl)amino)-1,3,5-triazin-2-yl)amino)-4,14-dioxo-6,9,12-trioxa-3,15-diazaheptadecanoyl)-4-hydroxy-N-(4-(4-methylthiazol-5-yl)benzyl)pyrrolidine-2-carboxamide (24)*

To a flame-dried two-neck 25 mL round-bottom flask equipped with a magnetic stir bar under inert atmosphere (N<sub>2</sub>) in an ice/ acetone bath was added a solution of **S39** (64.7 mg, 0.102 mmol) in DMF (510 μL, 0.20 M with respect to **S39**), followed by DIPEA (70 μL, 0.41 mmol). The bath was maintained for 15 minutes before HATU (46.8 mg, 0.122 mmol) was added. The solution was transferred to an ice bath for 20 minutes before **5** (44.3 mg, 0.103 mmol) was added and the bath was maintained for an additional 30 minutes. The solution gradually warmed to room temperature over 18 h, at which time the reaction was judged complete by TLC (5% MeOH in DCM). The reaction was quenched with deionized water (0.5 mL), and the aqueous phase was extracted with EtOAc (5 x 1.5 mL). The combined organic phase was washed with an aqueous solution of sodium bicarbonate (1 x 1 mL), dried with sodium sulfate, filtered through a fritted funnel using EtOAc, and concentrated by rotary evaporation at 35 °C to afford a solid. The solid was purified by Combiflash using a gradient of 3-10% MeOH in DCM and concentrated by rotary evaporation at 35 °C to afford **24**. Yield: 67.7 mg, 0.0650 mmol, off-white solid (64% over 2 steps). **R<sub>f</sub>**: 0.37 (5% MeOH in hexanes)

**<sup>1</sup>H NMR** (500 MHz, DMSO-d<sub>6</sub>, major and minor rotamer, OH signal not evident by <sup>1</sup>H NMR) δ (ppm): 9.59-9.41 (m, 1H), 8.97 (s, 1H), 8.58 (t, *J* = 6.1 Hz, 1H), 8.05 (br s, 1H), 7.83-7.56 (m, 5H), 7.43-7.39 (m, 7H), 7.14 (br s, 1H), 6.41 (d, *J* = 16.5 Hz, 1H), 5.15 (br s, 1H), 4.56 (d, *J* = 9.5 Hz, 1H), 4.45-4.39 (m, 2H), 4.37-4.34 (m, 1H), 4.26, (dd, *J* = 15.8, 5.8 Hz, 1H), 3.96 (s, 2H), 3.89-3.83 (m, 2H), 3.66 (dd, *J* = 10.7, 4.0 Hz, 1H), 3.62-3.51 (m, 10H), 3.44-3.24 (m, 8H), 2.44 (s, 3H), 2.17 (s, 6H) 2.05 (dd, *J* = 12.9, 7.8 Hz, 1H), 1.90 (ddd, *J* = 13.0, 8.9, 4.6 Hz, 1H), 0.93 (s, 9H). **<sup>13</sup>C NMR** (126 MHz, DMSO-d<sub>6</sub>, major and minor rotamer) δ (ppm): 171.72, 169.58, 169.11, 168.55, 165.93, 165.75, 164.17, 163.95, 151.43, 150.40, 147.72, 145.09, 139.41, 136.73, 132.49, 131.56, 131.11, 129.69, 128.85, 128.67, 128.11, 127.45, 127.13, 119.54, 119.03, 118.84, 102.33, 102.21, 95.90, 70.36, 70.15, 69.97, 69.58, 69.52, 68.85, 58.72, 56.54, 55.68, 54.90, 41.66, 40.0, 37.90, 35.69, 26.14, 18.37, 18.33, 15.90. **HRMS QToF-ESI**: calculated for C<sub>53</sub>H<sub>64</sub>N<sub>13</sub>O<sub>8</sub>S [M+H<sup>+</sup>] *m/z* 1042.4722; found *m/z* 1042.4701.

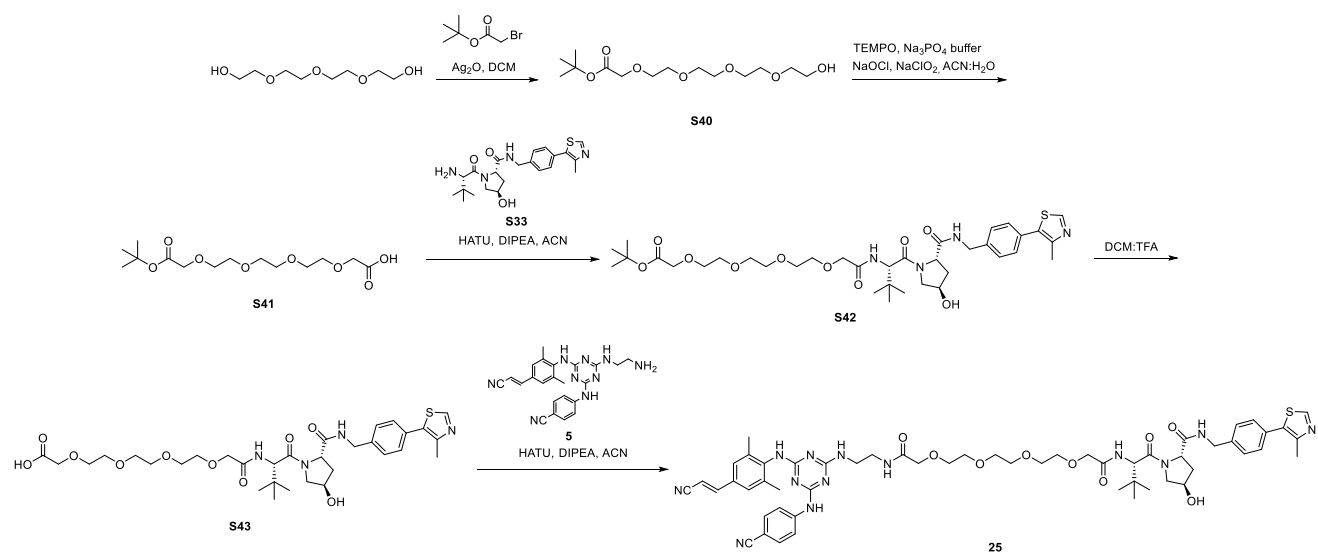

#### Scheme S15: Preparation of Compound 25

##### *tert*-Butyl 14-hydroxy-3,6,9,12-tetraoxatetradecanoate (**S40**)

To a flame-dried 4-dram vial equipped with a magnetic stir bar was added potassium iodide (85.5 mg, 0.521 mmol), Ag<sub>2</sub>O (290.0 mg, 1.251 mmol), tetraethylene glycol (540 μL, 3.10 mmol), and DCM (10 mL, 0.30 M with respect to tetraethylene glycol). The solution was chilled in an ice bath for 5 minutes before *tert*-butyl 2-bromoacetate (150 μL, 1.00 mmol) was added by syringe. The ice bath was allowed to warm to room temperature over 2 h, at which time the reaction was judged complete by TLC (1:1 hexanes:EtOAc). The reaction mixture was filtered through a pad of Celite using DCM, concentrated by rotary evaporation at 25 °C to afford an oil, and the oil was reconstituted in diethyl ether (10 mL). The solution was transferred to a separatory funnel and washed with brine (3 x 3 mL). The combined organic phase was dried with magnesium sulfate, filtered through a pad of Celite with diethyl ether, and concentrated by rotary evaporation at 35 °C

to afford **S40**. Crude Yield: 220 mg, 0.713 mmol, clear, colorless oil (71%). Compound **S40** was used to generate **S41** without further purification.

Characterization data matches that previously reported.<sup>7</sup>

**<sup>1</sup>H NMR** (500 MHz, CDCl<sub>3</sub>) $\delta$ (ppm): 4.00 (s, 2H), 3.69-3.65 (m, 13H), 3.60 (m, 2H), 2.31 (br s, 1H), 1.45 (s, 9H). **<sup>13</sup>C NMR** (126 MHz, CDCl<sub>3</sub>) $\delta$ (ppm): 169.76, 81.73, 72.74, 70.79, 70.68, 70.61, 70.34, 70.07, 69.08, 61.78, 28.20.

*16,16-dimethyl-14-oxo-3,6,9,12,15-pentaoxaheptadecanoic acid (S41)*

To a one-neck 25 mL round-bottom flask equipped with a magnetic stir bar was added **S40** (93.5 mg, 0.300 mmol), TEMPO (4.1 mg, 0.026 mmol), ACN (500 μL, 0.6 M with respect to **S40**) and 0.67 M sodium phosphate buffer (500 μL, 1:1 v/v NaH<sub>2</sub>PO<sub>4</sub>:Na<sub>2</sub>HPO<sub>4</sub>). The contents of the flask were stirred for 10 minutes at room temperature before a solution of sodium chlorite (55.2 mg, 0.610 mmol in 200 μL H<sub>2</sub>O) was added. The solution was heated to 35 °C before a 0.3 % sodium hypochlorite solution was added in five separate 40 μL portions at $t = 0, 2, 4, 7, 11$ h. An additional 100 μL portion was added at 15 h. After an additional 1 h, the reaction was judged complete by TLC (1:1 hexanes:EtOAc). The reaction was quenched with a sodium sulfite solution (105 mg in 1.0 mL deionized water). Diethyl ether was added, followed by a potassium carbonate solution (2 x 1.2 mL), and the mixture was extracted with diethyl ether (1 x 2 mL) to remove TEMPO and other impurities. The pH of the aqueous phase was adjusted to 3 with 3 M HCl, and the aqueous phase was extracted with diethyl ether (3 x 8 mL). The pH 3 combined organic phase was dried with magnesium sulfate, filtered through a pad of Celite with diethyl ether, and concentrated by rotary evaporation at 35 °C to afford **S41**. Crude Yield: 53.1 mg, 0.165 mmol, colorless oil (54%). Compound **S41** was used to generate **S42** without further purification.

Characterization data matches that previously reported.<sup>8</sup>

**<sup>1</sup>H NMR** (500 MHz, CDCl<sub>3</sub>) $\delta$ (ppm): 9.30 (br s, 1H), 4.15 (s, 2H), 4.00 (s, 2H), 3.74-3.64 (m, 12H), 1.45 (s, 9H). **<sup>13</sup>C NMR** (126 MHz, CDCl<sub>3</sub>) $\delta$ (ppm): 173.01, 169.80, 81.80, 71.40, 70.81, 70.69, 70.56, 70.47, 70.40, 69.03, 68.83, 28.17.

*tert-Butyl (S)-16-((2S,4R)-4-hydroxy-2-((4-(4-methylthiazol-5-yl)benzyl)carbamoyl)pyrrolidine-1-carbonyl)-17,17-dimethyl-14-oxo-3,6,9,12-tetraoxa-15-azaoctadecanoate (S42)*

To a flame-dried, two-neck 25 mL round-bottom flask equipped with a magnetic stir bar under inert atmosphere (N<sub>2</sub>) was added **S41** (64.6 mg, 0.192 mmol), HATU (98.3 mg, 0.258 mmol) and anhydrous ACN (1.0 mL, 0.20 M with respect to **S41**). The solution was chilled in an ice/acetone bath for 5 minutes before DIPEA (120 μL, 0.705 mmol) was added dropwise by syringe. The bath was maintained for 30 minutes before the adding **S33** (85.4 mg, 0.198 mmol). The bath gradually warmed to room temperature over 7 h, at which time the reaction was judged complete by TLC (5% MeOH in DCM). The reaction was quenched with deionized water (1 mL), and the aqueous phase was extracted with EtOAc (3 x 1 mL). The combined organic phase was washed with an aqueous solution of sodium bicarbonate (1 x 1 mL), dried with sodium sulfate, filtered through a fritted funnel using EtOAc, and concentrated by rotary evaporation at 35 °C. The material was purified by Combiflash using a gradient of 1-10% MeOH in DCM and concentrated by rotary evaporation at 35 °C to afford **S42**. Yield: 71.5 mg, 0.0995 mmol, off-white solid (52%).

**<sup>1</sup>H NMR** (500 MHz, CD<sub>3</sub>CN, major and minor rotamer) $\delta$ (ppm): 8.74 (s, 1H), 7.39 (m, 4H), 7.31 (t, $J$ = 6.0 Hz, 1H), 7.19 (d, $J$ = 9.3 Hz, 1H), 4.60 (d, $J$ = 9.3 Hz, 1H), 4.50- 4.45 (m, 2H), 4.42 (br s, 1H), 4.31 (dd, $J$ = 15.6, 5.5 Hz, 1H), 3.96-3.95 (m, 2H), 3.93 (s, 2H), 3.77 (br s, 0.3H), 3.75 (br s, 0.7H), 3.69 (dd, $J$ = 11.0, 3.8 Hz, 1H), 3.65-3.53 (m, 12H), 2.46 (s, 3H), 2.11-2.08 (m, 2H, signal is overlapping by solvent water), 1.43 (s, 9H), 0.96 (s, 9H). **<sup>13</sup>C NMR** (126 MHz, CD<sub>3</sub>CN, major and minor rotamer) $\delta$ (ppm): 172.74, 171.29, 170.82, 170.33, 151.68, 149.34, 140.25, 132.39, 131.44, 130.07, 128.65, 82.08, 71.85, 71.20, 71.12, 70.97, 70.89, 70.81, 70.79, 79.73, 69.44, 60.14, 57.65, 57.36, 43.18, 38.39, 36.52, 28.30, 26.72, 16.44. **HRMS QToF-ESI** calculated for C<sub>36</sub>H<sub>55</sub>N<sub>4</sub>O<sub>10</sub>S [M+H<sup>+</sup>] $m/z$ 735.3639; found: 735.3608.

*(S)-16-((2S,4R)-4-hydroxy-2-((4-(4-methylthiazol-5-yl)benzyl)carbamoyl)pyrrolidine-1-carbonyl)-17,17-dimethyl-14-oxo-3,6,9,12-tetraoxa-15-azaoctadecanoic acid (S43)*

In a one-neck 50 mL round-bottom flask equipped with a magnetic stir bar in an ice bath was added **S42** (61.1 mg, 0.0849 mmol) and TFA:DCM (0.5:1.5 v/v, 7.1 mL, 0.01 M with respect to **S42**). The ice bath was maintained for 1.5 h, at which time the reaction was judged complete by TLC (5% MeOH in DCM)). The volatile residues were removed by rotary evaporation at 33 °C to afford an oil, and the oil was dried overnight under reduced pressure to afford the corresponding acid **S43** (TFA salt). Crude Yield: 62.3 mg, 0.0803 mmol, light brown oil (95%). Compound **S43** was used to form **25** without further purification.

*N-1-(2-((4-((4-cyanophenyl)amino)-6-((4-((E)-2-cyanovinyl)-2,6-dimethylphenyl)amino)-1,3,5-triazin-2-yl)amino)ethyl)-N14-((S)-1-((2S,4R)-4-hydroxy-2-((4-(4-methylthiazol-5-yl)benzyl)carbamoyl)pyrrolidin-1-yl)-3,3-dimethyl-1-oxobutan-2-yl)-3,6,9,12-tetraoxatetradecanediarnide (25)*

To a flame-dried two-neck 15 mL round-bottom flask equipped with a magnetic stir bar under inert atmosphere (N<sub>2</sub>) in an ice/acetone bath was added **S43** (56.3 mg, 0.0845 mmol), DMF (420 μL, 0.20 M with respect to **S43**), and DIPEA (60 μL, 0.34 mmol). The bath was maintained for 5 minutes before adding HATU (39.4 mg, 0.102 mmol), and another 20 minutes before adding **5** (36.5 mg, 0.0849 mmol). The bath was maintained for an additional 30 minutes before gradually warming to room temperature over 19 h, at which time the reaction was judged complete by TLC (5% MeOH in DCM). The reaction was quenched with deionized water (0.43 mL), and the solution was extracted with EtOAc (3 x 1 mL). The combined organic phase was washed with an aqueous solution of sodium bicarbonate (1 x 1 mL), dried with sodium sulfate, filtered through a fritted funnel using EtOAc, and concentrated by rotary evaporation at 35 °C. The material was purified by Combiflash using a gradient of 5% MeOH in DCM and concentrated by rotary evaporation at

35 °C to afford **25**. Yield: 40.1 mg, 0.0369 mmol, off-white solid (43% over 2 steps). **R<sub>f</sub>**: 0.37 (5% MeOH/CH<sub>2</sub>Cl<sub>2</sub>)

**<sup>1</sup>H NMR** (500 MHz, DMSO-d<sub>6</sub>, major and minor rotamer, OH signal not evident by <sup>1</sup>H NMR) δ (ppm): 9.58 – 9.41 (m, 1H), 8.97 (s, 1H), 8.59 (t, *J* = 6.1 Hz, 1H), 8.05 (br s, 1H), 7.85 – 7.56 (m, 5H), 7.47 – 7.44 (m, 1H), 7.39 (s, 7H), 7.14 (br s, 1H), 6.41 (d, *J* = 16.4 Hz, 1H), 5.17 (br s, 1H), 4.56 (d, *J* = 9.5 Hz, 1H), 4.44 (t, *J* = 7.9 Hz, 1H), 4.40–4.35 (m, 2H), 4.25 (dd, *J* = 15.9, 5.8 Hz, 1H), 3.95 (s, 2H), 3.91 – 3.83 (m, 2H), 3.67 – 3.59 (m, 15H), 3.43 – 3.40 (m, 2H, overlapping with solvent water), 2.43 (s, 3H), 2.17 (s, 6H), 2.06 (dd, *J* = 13.1, 7.9 Hz, 1H), 1.90 (ddd, *J* = 13.0, 8.7, 4.4 Hz, 1H), 0.93 (s, 9H). **<sup>13</sup>C NMR** (126 MHz, DMSO-d<sub>6</sub>, major and minor rotamer) δ (ppm): 171.75, 169.61, 169.12, 168.56, 166.03, 165.79, 164.21, 163.98, 151.45, 150.43, 147.74, 145.13, 139.44, 136.75, 132.85, 132.46, 131.56, 131.13, 129.69, 128.87, 128.68, 128.13, 127.46, 127.14, 119.57, 119.05, 118.82, 102.33, 102.33, 95.91, 70.40, 70.14, 69.98, 69.79, 69.75, 69.58, 69.51, 68.86, 58.73, 56.57, 55.68, 41.67, 40.02, 37.92, 35.70, 26.16, 18.38, 18.35, 15.91. **HRMS QToF-ESI**: calculated for C<sub>55</sub>H<sub>68</sub>N<sub>13</sub>O<sub>9</sub>S [M+H<sup>+</sup>] *m/z* 1086.4984; found *m/z* 1086.4991.

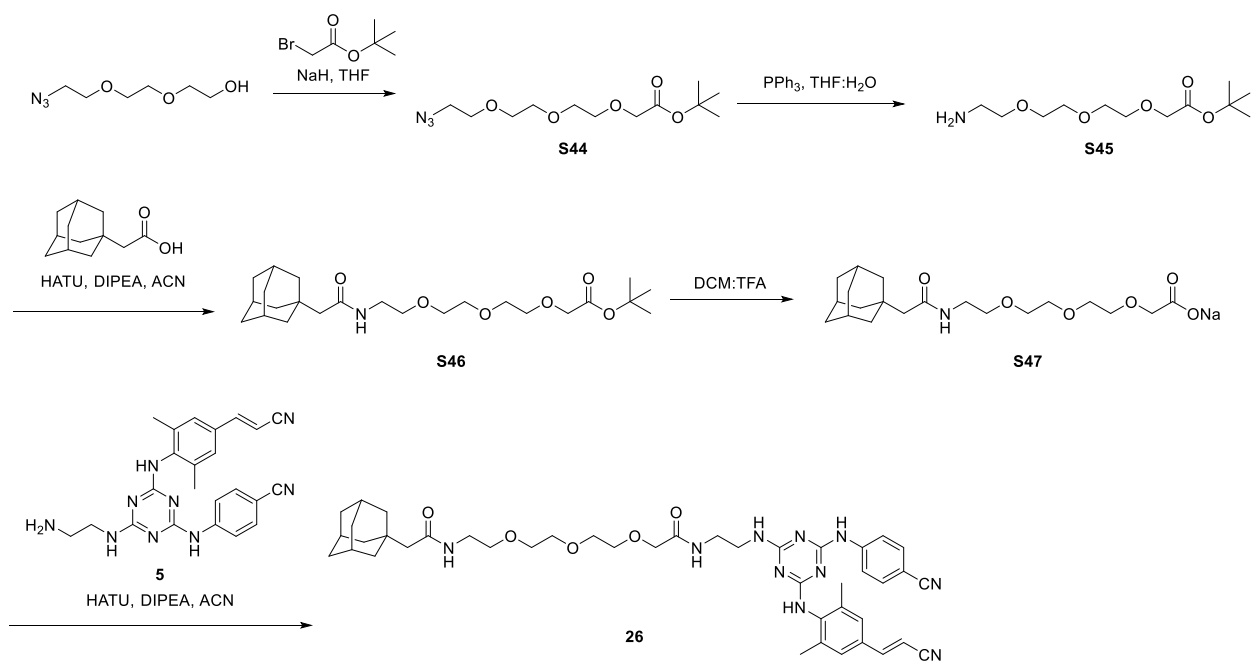

### Scheme S16: Preparation of Compound 26

#### *tert*-Butyl 2-(2-(2-(2-azidoethoxy)ethoxy)ethoxy)acetate (**S44**)

To a flame-dried round-bottom flask equipped with a magnetic stir bar under inert atmosphere (N<sub>2</sub>) was added a suspension of 60% NaH in mineral oil (86.0 mg, 2.15 mmol). The mineral oil was removed from the NaH by the addition of portions of distilled THF (1.0 mL) as follows. The suspension was stirred for 5 minutes then allowed to stand until the solid NaH settled on the bottom of the flask. The THF and mineral oil were decanted by syringe, and the process was repeated a total of three times. To the flask was added distilled THF (4.7 mL, 0.30 M with respect to 2-(2-(2-azidoethoxy)ethoxy)ethan-1-ol), and the solution was chilled in an ice bath for 10 minutes before 2-(2-(2-azidoethoxy)ethoxy)ethan-1-ol (250.0 mg, 1.427 mmol) was added dropwise by syringe (Note: gas evolution). The cold bath was maintained for 10 minutes before *tert*-butyl 2-bromoacetate (220 μL, 1.49 mmol) was added dropwise by syringe. The solution warmed to room temperature over 1 h, at which time the reaction was judged complete by TLC (1:1 hexanes:EtOAc). The reaction was quenched with aqueous $\text{NH}_4\text{Cl}$ (1 mL), and the mixture was stirred for 15 minutes before the solvent was removed by rotary evaporation at 30 $^\circ\text{C}$. The concentrate was reconstituted in diethyl ether and washed with brine (3 x 2.0 mL). The combined organic phase was dried with magnesium sulfate, filtered through a pad of Celite using diethyl ether, and concentrated by rotary evaporation at 30 $^\circ\text{C}$ to afford **S44**. Crude Yield: 237.1 mg, 0.8195 mmol, clear, colorless oil (55%). Compound **S44** was used to generate **S45** without further purification.

Characterization data matches that reported previously.<sup>11</sup>

**$^1\text{H}$ NMR** (500 MHz, $\text{CDCl}_3$) $\delta$ (ppm): 3.92 (s, 2H), 3.60-3.57 (m, 12H), 2.90 (br s, 1H), 1.38 (s, 9H); **$^{13}\text{C}$ NMR** (126 MHz, $\text{CDCl}_3$) $\delta$ (ppm): 169.56, 81.54, 72.52, 70.56, 70.50, 70.46, 70.36, 70.11, 68.82, 61.47, 27.97.

*tert*-Butyl 2-(2-(2-(2-aminoethoxy)ethoxy)ethoxy)acetate (**S45**)

To a 2-dram vial equipped with a magnetic stir bar was added **S44** (237.1 mg, 0.8195 mmol), $\text{PPh}_3$ (430.2 mg, 1.640 mmol), and THF (2.6 mL, 0.30 M with respect to **S44**). Once the solution was homogeneous, deionized water (0.25 mL) was added and the solution was maintained at room temperature for 48 h, at which time the reaction was judged complete by TLC (1:1 hexanes:EtOAc). The solvent was removed by rotary evaporation at 35 $^\circ\text{C}$, and the concentrate was reconstituted in EtOAc and washed with aqueous $\text{NH}_4\text{Cl}$ (3 x 2 mL). The pH of the aqueous phase was adjusted to 10 with aqueous 1 N NaOH and extracted with $\text{CHCl}_3$ (6 x 3 mL). The combined organic phase was dried with magnesium sulfate, filtered through a pad of Celite using $\text{CHCl}_3$, and concentrated by rotary evaporation at 35 $^\circ\text{C}$ to afford **S45**. Crude Yield: 147.7 mg, 0.5609 mmol, light yellow oil (68%). Compound **S45** was used to generate **S46** without further purification.

Characterization data matches that reported previously.<sup>7</sup>

**$^1\text{H}$ NMR** (500 MHz, $\text{CDCl}_3$) $\delta$ (ppm): 4.01 (s, 2H), 3.70-3.62 (m, 8H), 3.53 (t, $J$ = 10.36, 2H), 2.87 (t, $J$ = 10.35, 2H), 2.14 (br s, 4H), 1.46 (s, 9H). **$^{13}\text{C}$ NMR** (126 MHz, $\text{CDCl}_3$) $\delta$ (ppm): 169.85, 81.81, 72.98, 70.77, 70.65, 70.34, 69.13, 41.72, 28.22.

*tert*-Butyl 1-((3*r*,5*r*,7*r*)-adamantan-1-yl)-2-oxo-6,9,12-trioxa-3-azatetradecan-14-oate (**S46**)

To a flame-dried round-bottom flask equipped with a magnetic stir bar under inert atmosphere (N<sub>2</sub>) was added adamantane acetic acid (90.0 mg, 0.467 mmol), HATU (211.4 mg, 0.5560 mmol), DIPEA (240 µL, 1.38 mmol), and anhydrous ACN (2.4 mL, 0.20 M with respect to **S45**). The solution was stirred for 15 minutes before adding **S45** (147.7 mg, 0.5609 mmol). At 18 h, the reaction was judged complete by TLC (4:1 hexanes:EtOAc). The solvent was removed by rotary evaporation at 35 °C, and the concentrate was reconstituted in EtOAc. The solution was washed with aqueous NH<sub>4</sub>Cl (3 x 3 mL), aqueous sodium bicarbonate (2 x 3 mL), and brine (1 x 3 mL). The combined organic phase was dried with magnesium sulfate, filtered through a pad of Celite using EtOAc, and concentrated by rotary evaporation at 35 °C to afford **S46**. Crude Yield: 168.1 mg, 0.3824 mmol, yellow oil (78%). Compound **S46** was used to generate **S47** without further purification.

**<sup>1</sup>H NMR** (500 MHz, CDCl<sub>3</sub>) δ (ppm): 6.04 (br s, 1H), 4.01-3.99 (m, 2H), 3.70-3.60 (m, 8H), 3.52 (t, *J* = 4.9 Hz, 2H), 3.43-3.41 (m, 2H), 1.93-1.91 (m, 4H), 1.66-1.60 (m, 11H), 1.46-1.45 (m, 12H). **<sup>13</sup>C NMR** (126 MHz, CDCl<sub>3</sub>) δ (ppm) 171.12, 169.71, 81.75, 70.77, 70.64, 70.62, 70.28, 70.18, 69.09, 51.74, 42.68, 39.10, 36.89, 32.79, 28.75, 28.20. **HRMS QToF-ESI**: calculated for C<sub>24</sub>H<sub>42</sub>NO<sub>6</sub> *m/z*: 440.3012 [M+H]<sup>+</sup>; found *m/z*: 440.3001.

*1-((3*r*,5*r*,7*r*)-adamantan-1-yl)-2-oxo-6,9,12-trioxa-3-azatetradecan-14-oic acid (**S47**)*

In a 2-dram vial equipped with a magnetic stir bar under inert atmosphere (N<sub>2</sub>) was added a solution of **S46** (168.1 mg, 0.3824 mmol) in DCM (2.6 mL, 0.15 M with respect to **S46**). The solution was chilled in an ice bath for 10 minutes before TFA (1.3 mL) was added slowly. The solution gradually warmed to room temperature over 3 hours, at which time the reaction was judged complete by TLC (4:1 hexanes:EtOAc). The volatile residues were removed by rotary evaporation at 25 °C to afford an oil. The oil was reconstituted in EtOAc and extracted with aqueous sodium bicarbonate (3 x 2 mL). The pH of the aqueous phase was adjusted to 3 with aqueous 1 N HCl and extracted with CHCl<sub>3</sub> (5 x 3 mL). The combined organic phase was dried with magnesium sulfate, filtered through a pad of Celite using EtOAc, and concentrated by rotary evaporation at 35 °C to afford **S47** (TFA salt). Crude yield: 47.0 mg, 0.123 mmol, yellow oil (20%). Compound **S47** was used to generate **26** without further purification.

**<sup>1</sup>H NMR** (500 MHz, CDCl<sub>3</sub>) δ (ppm): 9.05 (br s, 1H), 6.30 (br s, 1H), 4.15 (s, 2H), 3.76-2.74 (m, 2H), 3.68-3.65 (m, 2H), 3.62-3.60 (m, 2H), 3.55-3.53 (m, 2H), 3.45-3.42 (m, 2H), 1.97-1.94 (m, 5H), 1.68-1.60 (m, 11H); **<sup>13</sup>C NMR** (126 MHz, CDCl<sub>3</sub>) δ (ppm): 172.16, 171.92, 71.34, 70.66, 70.36, 70.26, 70.02, 68.99, 51.42, 42.61, 39.23, 36.91, 32.88, 28.79. **HRMS QToF-ESI**: calculated for C<sub>20</sub>H<sub>32</sub>NO<sub>6</sub>Na *m/z*: 404.0567 [M+Na]<sup>+</sup>; found *m/z*: 404.0559.

*2-((3*r*,5*r*,7*r*)-adamantan-1-yl)-N-(1-((4-((4-cyanophenyl)amino)-6-((4-((*E*)-2-cyanovinyl)-2,6-dimethylphenyl)amino)-1,3,5-triazin-2-yl)amino)-4-oxo-6,9,12-trioxa-3-azatetradecan-14-yl)acetamide (**26**)*

To a flame-dried round-bottom flask equipped with a magnetic stir bar under inert atmosphere (N<sub>2</sub>) was added **S47** (47.0 mg, 0.123 mmol), HATU (53.0 mg, 0.140 mmol), DIPEA (70 µL, 0.41

mmol), and anhydrous ACN (600 μL, 0.20 M with respect to **5**). The contents of the flask were stirred for 15 minutes at room temperature before adding **5** (50.0 mg, 0.117 mmol). At 18 h, the reaction was judged complete by TLC (5% MeOH in $\text{CHCl}_3$). The reaction solvent was removed by rotary evaporation at 35 $^{\circ}\text{C}$, and the concentrate was reconstituted in EtOAc (5 mL). The solution was washed with aqueous $\text{NH}_4\text{Cl}$ (3 x 3 mL), aqueous sodium bicarbonate (2 x 3 mL), and brine (1 x 3 mL). The combined organic phase was dried with magnesium sulfate, filtered through a pad of Celite using EtOAc, and concentrated to afford a solid. The solid was purified by flash chromatography using a gradient of 3-4% MeOH in $\text{CHCl}_3$ and the fractions were concentrated by rotary evaporation 35 $^{\circ}\text{C}$ to afford **26**. Yield: 59.0 mg, 0.0746 mmol, white solid (20% over two steps); $R_f$: 0.24 (4% MeOH in $\text{CHCl}_3$).

**$^1\text{H}$ NMR** (500 MHz, $\text{CDCl}_3$ major and minor rotamer) $\delta$ (ppm): 7.79-7.35 (m, 5H), 7.22 (br s, 2H), 6.11 (s, 1H), 5.88 (d, $J$ = 16.3 Hz, 1H), 4.00 (s, 2H), 3.63-3.45 (m, 14H), 2.26 (s, 6H), 1.91-1.80 (m, 11H), 1.68-1.47 (m, 13H). **$^{13}\text{C}$ NMR** (126 MHz, $\text{CDCl}_3$, major and minor rotamer) $\delta$ (ppm): 171.42, 170.92, 166.31, 165.05, 164.42, 150.10, 143.70, 137.24, 133.04, 132.22, 128.84, 127.18, 119.45, 118.91, 118.22, 104.85, 96.37, 70.87, 70.53, 70.35, 70.19, 70.00, 51.64, 42.68, 40.96, 39.52, 39.09, 36.85, 32.82, 28.74, 18.89. **HRMS QToF-ESI**: calculated for $\text{C}_{43}\text{H}_{54}\text{N}_{10}\text{O}_5$ $[\text{M}+\text{H}]^+$ $m/z$ 791.4356; found $m/z$ 791.4349.

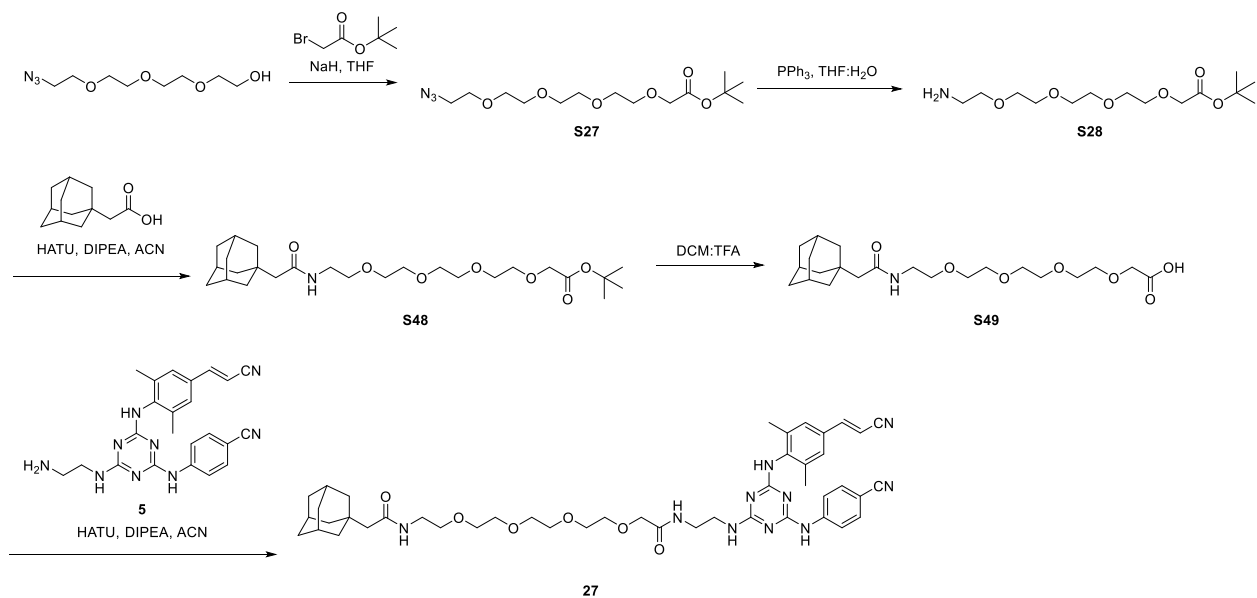

#### Scheme S17: Preparation of Compound 27

##### *tert*-Butyl 1-((3*r*,5*r*,7*r*)-adamantan-1-yl)-2-oxo-6,9,12,15-tetraoxa-3-azaheptadecan-17-oate (**S48**)

To a flame-dried round-bottom flask equipped with a magnetic stir bar under inert atmosphere ($\text{N}_2$) was added adamantane acetic acid (77.0 mg, 0.360 mmol), EDC·HCl (87.0 mg, 0.455 mmol), HOBT hydrate (62.0 mg, 0.455 mmol), $\text{CHCl}_3$ (2.0 mL, 0.20 M with respect to **S28**), and DIPEA (230 μL, 0.379 mmol). The contents of the flask were stirred for 15 minutes at room temperature before **S28** (116.5 mg, 0.3790 mmol) was added. At 18 h, the reaction was judged complete by TLC (4:1 hexanes:EtOAc). The solvent was removed by rotary evaporation at 35 °C to afford an oil which was reconstituted in EtOAc. The solution was washed with aqueous NH<sub>4</sub>Cl (3 x 3 mL), aqueous sodium bicarbonate (2 x 3 mL), and brine (1 x 3 mL). The combined organic phase was dried with magnesium sulfate, filtered through a pad of Celite using EtOAc, and concentrated by rotary evaporation at 35 °C to afford **S48**. Crude Yield 138 mg, 0.286 mmol, yellow oil (79%). Compound **S48** was used to generate **S49** without further purification.

**<sup>1</sup>H NMR** (500 MHz, CDCl<sub>3</sub>) δ (ppm): 6.40 (br s, 1H), 4.13 (s, 2H), 3.74-3.72 (m, 2H), 3.67-3.63 (m, 10H), 3.56 (t, *J* = 10.0 Hz, 2H), 3.43 (t, *J* = 5.46 Hz, 2H), 2.79 (s, 2H), 1.95-1.93 (m, 5H), 1.68-1.59 (m, 12H). **<sup>13</sup>C NMR** (126 MHz, CDCl<sub>3</sub>) δ (ppm): 172.56, 172.00, 71.09, 70.52, 70.47, 70.40, 70.34, 70.30, 70.22, 68.88, 51.47, 42.63, 39.14, 38.72, 36.86, 32.85, 28.74. **HRMS QToF-ESI**: calculated for C<sub>26</sub>H<sub>46</sub>NO<sub>7</sub> *m/z*: 484.3274 [M+H]<sup>+</sup>; found *m/z*: 484.3265.

*1-((3*r*,5*r*,7*r*)-adamantan-1-yl)-2-oxo-6,9,12,15-tetraoxa-3-azaheptadecan-17-oic acid (**S49**)*

To a 2-dram vial equipped with a magnetic stir bar placed in an ice bath was added a solution of **S48** (138.2 mg, 0.2857 mmol) and DCM (2.2 mL, 0.15 M with respect to **S48**). The contents of the flask were stirred 10 minutes before the slow addition of TFA (1.1 mL, 0.30 M with respect to **S48**) by syringe. The cold bath was maintained for 3 hours, at which time the reaction was judged complete by TLC (4:1 hexanes:EtOAc). The volatile residues were removed by rotary evaporation at 25 °C to afford **S49** (TFA salt). Crude Yield: 118.1 mg, 0.2828 mmol, yellow oil (99%). Compound **S49** was used to generate **27** without further purification.

**<sup>1</sup>H NMR** (500 MHz, CDCl<sub>3</sub>) δ (ppm): 10.33 (br s, 1H), 6.40 (br s, 1H), 4.13 (s, 2H), 3.74-3.72 (m, 2H), 3.67-3.61 (m, 11H), 3.57-3.55 (m, 2H), 3.43-3.42 (m, 2H), 2.79 (s, 2H), 1.95-1.93 (m, 5H), 1.66-1.59 (m, 12H); **<sup>13</sup>C NMR** (126 MHz, CDCl<sub>3</sub>) δ (ppm): 172.56, 172.00, 71.09, 70.52, 70.47, 70.40, 70.34, 70.30, 70.22, 68.88, 51.47, 42.63, 39.14, 38.72, 36.86, 32.85, 28.74. **HRMS QToF-ESI**: calculated for C<sub>22</sub>H<sub>37</sub>NO<sub>7</sub> *m/z*: 427.5448 [M+H]<sup>+</sup>; found *m/z*: 427.5441.

*14-(2-((3*r*,5*r*,7*r*)-adamantan-1-yl)acetamido)-N-(2-((4-(4-cyanophenyl)-6-((4-((*E*)-2-cyanovinyl)-2,6-dimethylphenyl)amino)-1,3,5-triazin-2-yl)amino)ethyl)-3,6,9,12-tetraoxatetradecanamide (**27**)*

To a flame-dried flask equipped with a magnetic stir bar under inert atmosphere (N<sub>2</sub>) was added **S49** (58.0 mg, 0.140 mmol), EDC•HCl (34.1 mg, 0.180 mmol), HOBT hydrate (24.2 mg, 0.181 mmol), DIPEA (50 μL, 0.31 mmol), and anhydrous DMF (600 μL, 0.20 M with respect to **S49**). The contents of the flask were stirred for 15 minutes at room temperature before the addition of **5** (63.0 mg, 0.148 mmol). At 18 h, the reaction was judged complete by TLC (5% MeOH in CHCl<sub>3</sub>). The reaction solvent was removed by rotary evaporation at 35 °C to afford a tacky solid, and the material was reconstituted in EtOAc. The solution was washed with aqueous NH<sub>4</sub>Cl (3x 3 mL), aqueous sodium bicarbonate (2 x 3 mL), and brine (1 x 3 mL). The combined organic phase was dried with sodium sulfate, filtered through a pad of Celite using EtOAc, and concentrated by rotary evaporation at 35 °C to afford a solid. The material was purified by flash chromatography using

5% MeOH in CHCl<sub>3</sub>, and the fractions were concentrated by rotary evaporation at 35 °C to afford **27**. Yield: 80.0 mg, 0.0958 mmol, white solid (34% over two steps); **R<sub>f</sub>**: 0.37 (2% MeOH in CHCl<sub>3</sub>).

**<sup>1</sup>H NMR** (500 MHz, CDCl<sub>3</sub>, major and minor rotamer) $\delta$ (ppm): 7.91-7.35 (m, 7H), 7.35-7.32 (m, 2H), 6.94 (br s, 2H), 6.12 (br s, 2H), 5.86 (d, *J* = 16.0 Hz, 1H), 3.95 (s, 2H), 3.63-3.40 (m, 19H), 2.25-2.23 (m, 6H), 1.90-1.85 (m, 5H), 1.66-1.55 (m, 11H). **<sup>13</sup>C NMR** (126 MHz, CDCl<sub>3</sub>, major and minor rotamer) $\delta$ (ppm): 171.28, 170.89, 165.94, 164.72, 164.22, 150.07, 143.70, 137.92, 137.24, 133.00, 132.10, 128.78, 127.13, 119.41, 119.07, 118.30, 104.75, 96.25, 70.91, 70.45, 70.15, 51.63, 42.64, 40.72, 39.39, 39.00, 36.80, 32.74, 28.68, 18.85. **HRMS QToF-ESI** calculated for C<sub>45</sub>H<sub>58</sub>N<sub>10</sub>O<sub>6</sub> [M+H]<sup>+</sup> *m/z* 835.4620; found *m/z* 835.4619.

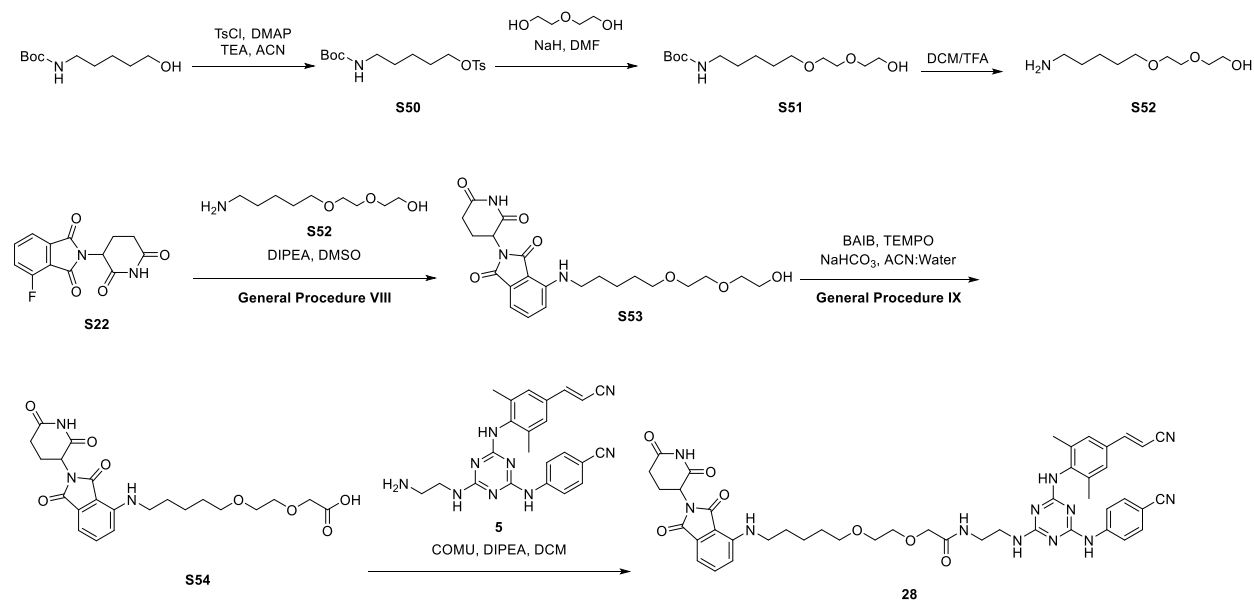

### Scheme S18: Preparation of Compound 28

#### 5-((*tert*-butoxycarbonyl)amino)pentyl 4-methylbenzenesulfonate (**S50**)

To a 250 mL flame-dried round-bottom flask equipped with a magnetic stir bar under inert atmosphere (N<sub>2</sub>) was added DMAP (200.0 mg, 1.637 mmol), TsCl (1.0200 g, 5.3501 mmol), and ACN (25 mL, 0.20 M with respect to *tert*-butyl (5-hydroxypentyl)carbamate). The solution was chilled in an ice bath for 15 minutes before adding TEA (1.0 mL, 7.4 mmol). Once the contents were homogenous, *tert*-butyl (5-hydroxypentyl)carbamate (1.00 g, 4.92 mmol) was added dropwise and the solution was gradually warmed to room temperature over 18 h, at which time the reaction was judged complete by TLC (3:1 hexanes:EtOAc). The solvent was removed by rotary evaporation at 35 °C to afford an oil. The oil was reconstituted in EtOAc and washed with aqueous NH<sub>4</sub>Cl (3 x 10 mL) and brine (3 x 10 mL). The combined organic phase was dried with magnesium sulfate, filtered through a pad of Celite using EtOAc, and concentrated by rotary evaporation at 35 °C to afford an oil. The oil was purified by flash chromatography using 15%

EtOAc in hexanes and concentrated by rotary evaporation at 35 °C to afford **S50**. Yield: 1.3012 g, 3.6402 mmol, clear, colorless oil (75%).

**<sup>1</sup>H NMR** (500 MHz, CDCl<sub>3</sub>) δ (ppm): 7.76 (d, *J* = 7.76 Hz, 2H), 7.33 (d, *J* = 7.33 Hz, 2H), 4.54 (br s, 1H), 3.99 (t, *J* = 6.36 Hz, 2H), 3.04 (dd, *J* = 6.26 Hz, 2H), 2.42 (s, 3H), 1.66-1.60 (m, 2H), 1.41-1.29 (m, 13H). **<sup>13</sup>C NMR** (126 MHz, CDCl<sub>3</sub>) δ (ppm): 156.03, 144.83, 133.16, 129.92, 127.93, 79.15, 70.44, 40.27, 29.47, 28.54, 28.47, 22.72, 21.69. **HRMS QToF-ESI**: calculated for C<sub>17</sub>H<sub>27</sub>NO<sub>5</sub>S [M+H<sup>+</sup>] *m/z* 380.1508; found *m/z* 380.1509.

*tert*-Butyl (5-(2-(2-hydroxyethoxy)ethoxy)pentyl)carbamate (**S51**)

To a flame-dried 25 mL round-bottom flask equipped with a magnetic stir bar under inert atmosphere (N<sub>2</sub>) was added a 60% suspension of NaH in mineral oil (175.0 mg, 4.375 mmol), and anhydrous DMF (9.3 mL, 0.30 M with respect to **S50**). The solution was chilled in an ice bath before diethylene glycol (1.33 mL, 14.0 mmol) was added dropwise by syringe, and the bath was maintained for 15 minutes before **S50** (1.0000 g, 2.7975 mmol) was added by syringe. The solution gradually warmed to room temperature over 18 h, at which time the reaction was judged complete by TLC (3:1 hexanes:EtOAc). The reaction was quenched with aqueous NH<sub>4</sub>Cl (0.20 mL), stirred for 20 minutes, and the mixture was transferred to a round-bottom flask and concentrated azeotropically with toluene by rotary evaporation at 35 °C. The concentrate was reconstituted in CHCl<sub>3</sub> (10 mL) and washed with brine (2 x 5 mL). The combined organic phase was dried with magnesium sulfate, filtered through a pad of Celite using CHCl<sub>3</sub>, and concentrated by rotary evaporation at 35 °C. The material was purified by flash chromatography using 20% EtOAc in hexanes, and the fractions were concentrated by rotary evaporation at 35 °C to afford **S51**. Yield: 782.0 mg, 2.684 mmol, light yellow oil (96%).

**<sup>1</sup>H NMR** (500 MHz, CDCl<sub>3</sub>) δ (ppm): 4.69 (br s, 1H), 3.70-3.55 (m, 6H), 3.44 (t, *J* = 6.36, 2H), 3.09 (d, *J* = 5.14, 2H), 2.80 (br s, 1H), 1.59-1.56 (m, 2H), 1.48-1.35 (m, 11H). **<sup>13</sup>C NMR** (126 MHz, CDCl<sub>3</sub>) δ (ppm): 156.13, 79.08, 72.65, 71.28, 70.53, 70.27, 61.85, 40.51, 29.84, 29.22, 28.51, 23.41. **HRMS QToF-ESI**: calculated for C<sub>14</sub>H<sub>29</sub>NO<sub>5</sub>S [M+H<sup>+</sup>] *m/z* 318.2280; found *m/z* 318.2283.

2-(2-((5-aminopentyl)oxy)ethoxy)ethan-1-ol (**S52**)

To a 4-dram vial equipped with a magnetic stir bar was added **S51** (466.2 mg, 1.600 mmol) and 2-MeTHF (5.2 mL, 0.30 M with respect to **S51**). The vial was chilled in an ice bath for 5 minutes before TFA (2.6 mL, 0.60 M with respect to **S51**) was added. The bath was maintained for 3 h, at which time the reaction was judged complete by TLC (1:1 hexanes:EtOAc). The solvent was removed by rotary evaporation at 30 °C to afford **S52** (TFA salt). Crude Yield: 300.1 mg, 1.569 mmol, yellow oil (98%). Compound **S52** was used to generate **S53** without further purification.

2-(2,6-dioxopiperidin-3-yl)-4-((5-(2-(2-hydroxyethoxy)ethoxy)pentyl)amino)isoindoline-1,3-dione (**S53**)

**See General Procedure VIII to generate S53.** Quantities Used: **S22** (100.0 mg, 0.3620 mmol), **S52** (81.7 mg, 0.398 mmol), anhydrous DMSO (2.4 mL, 0.15 M with respect to **S22**), and DIPEA (200 μL, 1.09 mmol). The material was purified by column chromatography using 20% acetone in hexanes and concentrated by rotary evaporation at 35 °C to afford **S53**. Yield: 82.0 mg, 0.180 mmol, yellow solid (45%).

**<sup>1</sup>H NMR** (500 MHz, CDCl<sub>3</sub>) $\delta$ (ppm): 8.47 (br s, 1H), 7.47 (t, $J$ = 8.0 Hz, 1H), 7.07 (d, $J$ = 7.1 Hz, 1H), 6.87 (d, $J$ = 8.5 Hz, 1H), 6.23 (t, $J$ = 5.3 Hz, 1H), 4.92-4.89 (m, 1H), 3.73-3.71 (m, 2H), 3.67-3.66 (m, 2H), 3.62-3.57 (m, 4H), 3.48 (t, $J$ = 7.5 Hz, 2H), 3.28-3.24 (m, 2H), 2.88-2.71 (m, 4H), 2.13-2.09 (m, 2H), 1.71-1.61 (m, 4H), 1.50-1.44 (m, 2H). **<sup>13</sup>C NMR** (126 MHz, CDCl<sub>3</sub>) $\delta$ (ppm): 171.39, 169.62, 168.65, 167.76, 147.05, 136.24, 132.58, 116.76, 111.50, 109.93, 72.61, 71.23, 70.52, 70.34, 61.89, 48.95, 42.64, 31.52, 29.36, 29.13, 23.67, 22.91. **HRMS QToF-ESI:** calculated for C<sub>22</sub>H<sub>29</sub>N<sub>3</sub>O<sub>7</sub> [M+H]<sup>+</sup> $m/z$ 448.2084; found $m/z$ 448.2079.

*2-(2-((6-((2-(2,6-dioxopiperidin-3-yl)-1,3-dioxoisindolin-4-yl)amino)hexyl)oxy)ethoxy)acetic acid (**S54**)*

**See General Procedure IX to generate S54.** Quantities used: BAIB (170.2 mg, 0.5285 mmol), TEMPO (9.1 mg, 0.058 mmol), sodium bicarbonate (110.0 mg, 1.310 mmol), 1:1 v/v ACN:deionized water (2.4 mL, 0.1 M with respect to **S53**), and **S53** (110.0 mg, 0.2458 mmol). The reaction was judged complete by TLC at 4 h (5:2 hexanes:acetone). Crude Yield: 84.0 mg, 0.197 mmol orange solid (80%). Compound **S54** was used to generate **28** without further purification.

*(E)-N-(2-((4-((4-cyanophenyl)amino)-6-((4-(2-cyanovinyl)-2,6-dimethylphenyl)amino)-1,3,5-triazin-2-yl)amino)ethyl)-2-(2-((5-((2-(2,6-dioxopiperidin-3-yl)-1,3-dioxoisindolin-4-yl)amino)pentyl)oxy)ethoxy)acetamide (**28**)*

To a flame-dried 10 mL round-bottom flask equipped with a magnetic stir bar under inert atmosphere (N<sub>2</sub>) was added **S54** (60.4 mg, 0.131 mmol), **5** (61.2 mg, 0.144 mmol), HATU (64.6 mg, 0.170 mmol), and DCM (1.3 mL, 0.01 M with respect to **S54**). The flask was chilled in an ice bath for 5 minutes before DIPEA (80 μL, 0.44 mmol) was added. The ice bath was maintained for 30 minutes before the solution gradually warmed to room temperature over 18 h, at which time the reaction was judged complete by TLC (1:1 hexanes:acetone). The reaction was quenched with saturated aqueous sodium bicarbonate (20 mL) and extracted with EtOAc (6 x 10 mL). The combined organic phase was washed with aqueous sodium bicarbonate (3 x 10 mL) then dried with sodium sulfate, filtered through a pad of Celite using EtOAc, and concentrated by rotary evaporation at 35 °C to afford a solid. The solid was purified by preparative Thin-Layer Chromatography using 50% acetone in hexanes. The material was desorbed from the silica by stirring the slurry in acetone for 3 h, filtered through a pad of Celite using acetone, and concentrated by rotary evaporation at 35 °C to afford **28**. Yield: 28.1 mg, 0.0323 mmol, yellow solid (13% over two steps). **R<sub>f</sub>**: 0.17 (1:1 acetone in hexane)

**<sup>1</sup>H NMR** (500 MHz, CDCl<sub>3</sub>, major and minor rotamer) δ (ppm): 8.35 – 7.35 (m, 6H), 7.20 – 6.86 (m, 4H), 6.39 – 6.26 (m, 1H), 5.87 (d, *J* = 16.2 Hz, 1H), 5.40 – 4.98 (m, 1H), 3.94 (s, 2H), 3.60 – 3.30 (m, 12H), 2.91 – 2.63 (m, 3H), 2.26 (s, 7H), 1.68 – 1.43 (m, 11H), 1.25 (s, 4H). **<sup>13</sup>C NMR** (126 MHz, CDCl<sub>3</sub>, major and minor rotamer) δ (ppm): 174.37, 171.40, 170.68, 169.65, 168.37, 167.70, 166.09, 165.27, 164.04, 150.33, 147.17, 144.10, 143.44, 137.91, 137.38, 136.36, 133.10, 131.93, 128.83, 127.14, 119.67, 118.93, 117.91, 117.01, 111.72, 104.92, 96.07, 71.08, 70.94, 70.71, 69.77, 69.66, 53.90, 48.88, 48.81, 42.23, 40.75, 39.22, 31.90, 31.45, 29.86, 29.41, 28.94, 28.30, 23.29, 22.85, 18.93. **HRMS QToF-ESI**: calculated for C<sub>45</sub>H<sub>49</sub>N<sub>12</sub>O<sub>7</sub> [M+H]<sup>+</sup> *m/z* 869.3847; found *m/z* 869.3844.

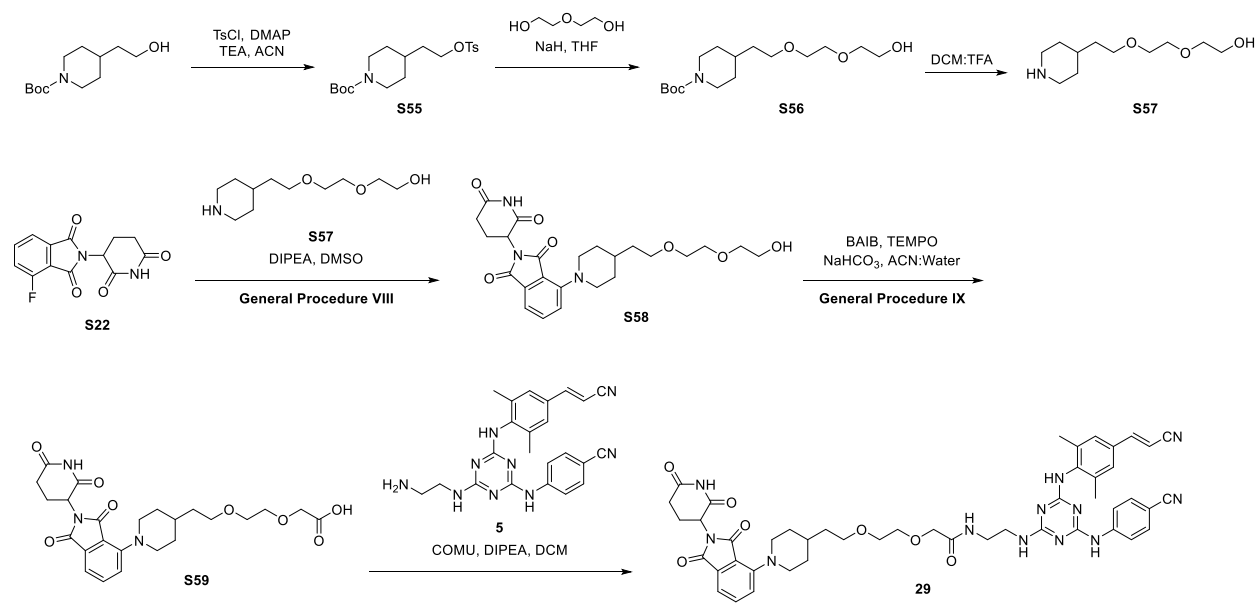

### Scheme S19: Preparation of Compound 29

#### *tert*-Butyl 4-(2-(tosyloxy)ethyl)piperidine-1-carboxylate (S55)

To a 25 mL flame-dried round-bottom flask equipped with a magnetic stir bar under inert atmosphere (N<sub>2</sub>) in an ice bath was added *tert*-butyl 4-(2-hydroxyethyl)piperidine-1-carboxylate (500.0 mg, 2.180 mmol), TsCl (457.6 mg, 2.400 mmol), DMAP (53.2 mg, 0.435 mmol), and anhydrous ACN (7.3 mL, 0.30 M with respect to *tert*-butyl 4-(2-hydroxyethyl)piperidine-1-carboxylate). The bath was maintained for 5 minutes before dropwise addition of TEA (0.90 mL, 6.5 mmol) by syringe. The solution was gradually warmed to room temperature over 3 h, at which time the reaction was judged complete by TLC (2:1 hexanes:EtOAc). The solvent was removed by rotary evaporation at 35 °C to afford an oil, and the oil was reconstituted in EtOAc and washed with aqueous NH<sub>4</sub>Cl (3 x 5 mL). The combined organic phase was dried with magnesium sulfate, filtered through a pad of Celite using EtOAc, and concentrated by rotary evaporation at 35 °C to afford a yellow oil. The oil was purified by flash column chromatography using 20% EtOAc in hexanes, and the fractions were concentrated by rotary evaporation at 35 °C to afford **S55**. Yield: 599.6 mg, 1.563 mmol, clear, colorless oil (72%).

**<sup>1</sup>H NMR** (500 MHz, CDCl<sub>3</sub>) δ (ppm): 7.78-7.75 (m, 2H), 7.34-7.26 (m, 2H), 4.05-4.00 (m, 4H), 2.60 (t, *J* = 12.5 Hz, 2H), 2.42 (m, 3H), 2.15-2.14 (m, 1H), 1.56-1.50 (m, 5H), 1.46-1.42 (m, 9H), 1.02-1.00 (m, 2H). **<sup>13</sup>C NMR** (126 MHz, CDCl<sub>3</sub>) δ (ppm): 154.86, 144.92, 133.20, 129.96, 127.99, 79.43, 68.09, 43.83, 35.37, 32.31, 31.74, 28.54, 21.72. **HRMS QToF-ESI**: calculated for C<sub>19</sub>H<sub>29</sub>NO<sub>5</sub>S [M+H<sup>+</sup>] *m/z* 384.1845; found *m/z* 384.1855.

*tert*-Butyl 4-(2-(2-(2-hydroxyethoxy)ethoxy)ethyl)piperidine-1-carboxylate (**S56**)

To a 25 mL flame-dried round-bottom flask equipped with a magnetic stir bar under inert atmosphere (N<sub>2</sub>) was added a 60% suspension of NaH in mineral oil (80.0 mg, 2.00 mmol). The mineral oil was removed from the NaH by the addition of portions of distilled THF (2.0 mL) as follows. The suspension was stirred for 5 minutes then allowed to stand until the solid NaH settled on the bottom of the flask. The THF and mineral oil were decanted by syringe, and the process was repeated a total of three times. To the flask was added distilled THF (6.7 mL, 0.60 M with respect to diethylene glycol). The solution was chilled in an ice bath before diethylene glycol (380 μL, 4.00 mmol) was added dropwise (Note: gas evolution). The ice bath was maintained for 20 minutes before **S55** (575.3 mg, 1.500 mmol) was added, and the solution was gradually warmed to room temperature over 18 h, at which time the reaction was judged complete by TLC (3:1 hexanes:EtOAc). The reaction was quenched with aqueous NH<sub>4</sub>Cl (20 mL) and was stirred for 15 minutes before the solvent was removed azeotropically with toluene by rotary evaporation at 55 °C. The concentrate was reconstituted in 2-MeTHF (5 mL), transferred to a separatory funnel, and washed with brine (5 x 5 mL) to remove excess diethylene glycol. The combined organic phase was dried with magnesium sulfate, filtered through a pad of Celite using 2-MeTHF, and concentrated by rotary evaporation at 35 °C to afford a yellow oil. The oil was purified by column chromatography using a gradient of 20-100% EtOAc in hexanes and concentrated by rotary evaporation at 35 °C to afford **S56**. Yield: 364.6 mg, 1.148 mmol, yellow oil (77%).

**<sup>1</sup>H NMR** (500 MHz, CDCl<sub>3</sub>) δ (ppm): 4.05 (d, *J* = 13.2 Hz, 2H), 3.72-3.71 (m, 2H), 3.67-3.65 (m, 2H), 3.61-3.59 (m, 2H), 3.58-3.56 (m, 2H), 3.52-3.49 (m, 2H), 2.70-2.64 (m, 2H), 2.26 (br s, 1H), 1.64 (d, *J* = 12.6 Hz, 2H), 1.56-1.50 (m, 3H), 1.44 (s, 9H), 1.13-1.06 (m, 2H). **<sup>13</sup>C NMR** (126 MHz, CDCl<sub>3</sub>) δ (ppm): 155.01, 79.34, 72.60, 70.57, 70.35, 68.99, 61.95, 44.06, 36.19, 33.04, 32.22, 28.59. **HRMS QToF-ESI**: calculated for C<sub>16</sub>H<sub>31</sub>NO<sub>5</sub> [M+H<sup>+</sup>] *m/z* 318.2280; found *m/z* 318.2283.

2-(2-(2-(piperidin-4-yl)ethoxy)ethoxy)ethan-1-ol (**S57**)

To a 25 mL flame-dried round-bottom flask equipped with a magnetic stir bar in an ice bath was added **S56** (364.6 mg, 1.148 mmol) and 2-MeTHF (7.3 mL, 0.16 M with respect to **S56**). Once the solution was homogeneous, TFA (3.7 mL) was added, and the cold bath maintained for 4 h, at which time the reaction was judged complete by TLC (2:1 hexanes:EtOAc). The volatiles were removed by rotary evaporation at 25 °C, and the material was dried under reduced pressure to afford **S57**. Crude Yield: 250.2 mg, 0.1082 mmol, yellow oil (99%). Compound **S57** was used to generate **S58** without further purification.

*2-(2,6-dioxopiperidin-3-yl)-4-(4-(2-(2-(2-hydroxyethoxy)ethoxy)ethyl)piperidin-1-yl)isoindoline-1,3-dione (S58)*

**See General Procedure VIII to generate S58.** Quantities Used: **S22** (100.0 mg, 0.3620 mmol), anhydrous DMSO (2.4 mL, 0.15 M with respect to **S57**), **S57** (92.1 mg, 0.398 mmol), and DIPEA (200 μL, 1.1 mmol). The material was purified by column chromatography using 25% acetone in hexanes and concentrated by rotary evaporation at 35 °C to afford **S58**. Yield: 139.2 mg, 0.2940 mmol, yellow solid (26% over two steps).

**<sup>1</sup>H NMR** (500 MHz, CDCl<sub>3</sub>) $\delta$ (ppm): 8.22-8.20 (m, 1H), 7.55 (t, $J$ = 7.5 Hz, 1H), 7.35 (d, $J$ = 7.0 Hz, 1H), 7.16 (d, $J$ = 8.2 Hz, 1H), 4.97-4.94 (m, 1H), 3.74-3.55 (m, 12H), 2.89-2.71 (m, 5H), 2.17-2.09 (m, 4H), 1.84 (d, $J$ = 12.5 Hz, 2H), 1.63-1.61 (m, 4H), 1.51-1.49 (m, 2H). **<sup>13</sup>C NMR** (126 MHz, CDCl<sub>3</sub>) $\delta$ (ppm): 171.18, 168.40, 167.56, 166.82, 151.02, 135.60, 134.24, 123.82, 117.26, 115.40, 72.60, 70.59, 70.37, 69.01, 61.98, 52.22, 51.88, 49.22, 36.14, 32.50, 32.38, 31.55, 22.82. **HRMS QToF-ESI**: calculated for C<sub>24</sub>H<sub>31</sub>N<sub>3</sub>O<sub>7</sub> [M+H<sup>+</sup>] $m/z$ 473.2162; found $m/z$ 473.2162.

*2-(2-(2-(1-(2-(2,6-dioxopiperidin-3-yl)-1,3-dioxoisoindolin-4-yl)piperidin-4-yl)ethoxy)ethoxy)acetic acid (S59)*

**See General Procedure IX to generate S59.** Quantities Used: BAIB (129.3 mg, 0.331 mmol), TEMPO (5.7 mg, 0.037 mmol), sodium bicarbonate (48.4 mg, 0.576 mmol), 1:1 v/v ACN:deionized water (720 μL, 0.20 M with respect to **S58**) and **S58** (70.0 mg, 0.147 mmol). The reaction was judged complete by TLC at 4 h (5:2 hexanes:acetone). Crude Yield: 25.0 mg, 0.0513 mmol, yellow solid (35%). Compound **S59** was used to generate **29** without further purification.

*(E)-N-(2-((4-((4-cyanophenyl)amino)-6-((4-(2-cyanovinyl)-2,6-dimethylphenyl)amino)-1,3,5-triazin-2-yl)amino)ethyl)-2-(2-(2-(1-(2-(2,6-dioxopiperidin-3-yl)-1,3-dioxoisoindolin-4-yl)piperidin-4-yl)ethoxy)ethoxy)acetamide (29)*

To a 1-dram vial equipped with a magnetic stir bar under inert atmosphere (N<sub>2</sub>) was added **S59** (25.0 mg, 0.0513 mmol), **5** (24.0 mg, 0.0565 mmol), COMU (24.4 mg, 0.0565 mmol), and anhydrous ACN (0.70 mL, 0.08 M with respect to **5**). The flask was transferred to an ice bath for 10 minutes before DIPEA (40 μL, 0.18 mmol) was added dropwise by syringe. The bath gradually warmed to room temperature over 18 h, at which time the reaction was judged complete by TLC (3% MeOH in DCM). The solvent was removed by rotary evaporation at 25 °C to afford a solid, which was reconstituted in EtOAc. The organic phase was washed with aqueous NH<sub>4</sub>Cl (3 x 3.0 mL), aqueous sodium bicarbonate (2 x 3.0 mL), and brine (1 x 3.0 mL). The organic phase was dried with magnesium sulfate, filtered through a pad of Celite using EtOAc, and concentrated by rotary evaporation at 35 °C to afford a solid. The solid was purified by preparative Thin-Layer Chromatography using 100% EtOAc. The material was desorbed from the silica by stirring the slurry in 20% MeOH in CHCl<sub>3</sub> for 3 h. The slurry was filtered through a pad of Celite using CHCl<sub>3</sub>,

**<sup>1</sup>H NMR** (500 MHz, CD<sub>3</sub>CN, major and minor rotamer) δ (ppm): 9.93 (br s, 1H), 8.02-7.91 (m, 2H), 7.55-7.28 (m, 8H), 6.99 (d, *J* = 6.8 Hz, 1H), 6.91 (d, *J* = 7.8 Hz, 1H), 6.22 (br s, 2H), 6.04 (d, *J* = 16.2 Hz, 1H), 4.96-4.93 (m, 1H), 3.84 (s, 2H), 3.53-3.37 (m, 9H), 3.20 (s, 2H), 2.78-2.64 (m, 3H), 2.21 (s, 6H), 2.19-2.12 (m, 10H), 1.55-1.50 (m, 4H), 1.37-1.36 (m, 4H). **<sup>13</sup>C NMR** (126 MHz, CD<sub>3</sub>CN, major and minor rotamer) δ (ppm): 173.89, 171.40, 170.51, 168.58, 167.63, 166.17, 165.39, 151.15, 147.85, 145.29, 139.39, 138.15, 137.11, 133.78, 133.51, 133.03, 129.32, 127.98, 120.22, 119.79, 119.55, 111.55, 110.59, 104.83, 96.89, 71.75, 71.55, 71.06, 70.42, 49.78, 43.00, 41.20, 39.96, 32.00, 30.01, 29.41, 24.13, 23.36, 18.93, 18.83. **HRMS QToF-ESI**: calculated for C<sub>47</sub>H<sub>51</sub>N<sub>12</sub>O<sub>7</sub> [M+H]<sup>+</sup> *m/z* 895.4004; found *m/z* 895.3995.

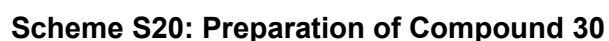

50

To a flame-dried, three-neck 100 mL round-bottom flask equipped with a magnetic stir bar under inert atmosphere (N<sub>2</sub>) was added 1,5-pentanediol (845.7 mg, 8.120 mmol), and distilled THF (12.0 mL, 0.68 M with respect to 1,5-pentanediol). The solution was cooled to -72 °C for 10 minutes before adding n-BuLi (2.20 mL, 5.41 mmol, 2.5 M in hexanes, *titrated using N-benzyl benzamide*).<sup>9</sup> The bath was maintained for 10 minutes before a solution of TBSCl (851.0 mg, 5.640 mmol) in distilled THF (4.0 mL, 1.4 M with respect to TBSCl) was added dropwise by syringe. The bath was maintained for 1 h before gradually warming to room temperature over 3.5 h, at which time the reaction was judged complete by TLC (1:4 hexanes:EtOAc). The reaction was quenched with an aqueous solution of NH<sub>4</sub>Cl (16 mL), and extracted with diethyl ether (5 × 9 mL). The combined organic phases was dried with magnesium sulfate, filtered through a fritted funnel using diethyl ether, and concentrated by rotary evaporation at 35 °C. The material was purified by flash column chromatography using a gradient of 20-35% diethyl ether in hexanes and concentrated by rotary evaporation at 35 °C to afford **S60**. Yield: 855.4 mg, 3.916 mmol, colorless oil (70%).

Characterization data matches that reported previously.<sup>10</sup>

**<sup>1</sup>H NMR** (500 MHz, CDCl<sub>3</sub>) δ (ppm): 3.67-3.61 (m, 4H), 1.62-1.52 (m, 4H), 1.44-1.37 (m, 2H), 1.31-1.30 (m, 1H), 0.89 (s, 9H), 0.05 (s, 6H). **<sup>13</sup>C NMR** (126 MHz, CDCl<sub>3</sub>) δ (ppm): 63.26, 63.12, 32.65, 26.12, 22.18, 18.52, -5.13.

##### 5-((*tert*-butyldimethylsilyl)oxy)pentyl 4-methylbenzenesulfonate (**S61**)

To a one-neck 50 mL round-bottom flask equipped with a magnetic stir bar in an ice bath was added DMAP (83.8 mg, 0.666 mmol), **S60** (1.520 g, 6.959 mmol), DCM (8.0 mL, 0.80 M with respect to **S60**), and a solution of TsCl (1.510 g, 7.880 mmol) in DCM (5.4 mL, 1.46 M with respect to TsCl). The ice bath was maintained while TEA (1.80 mL, 13.1 mmol) was added by syringe. The solution was gradually warmed to room temperature over 5 h, at which time the reaction was judged complete by TLC (9:1 hexanes:EtOAc). The reaction was quenched with aqueous NH<sub>4</sub>Cl (10 mL), and the aqueous phase was extracted with DCM (3 × 6 mL). The combined organic phase was dried with magnesium sulfate, filtered through a fritted funnel using DCM, and concentrated by rotary evaporation at 35 °C to afford an oil. The oil was purified by flash column chromatography using 10% EtOAc in hexanes and concentrated by rotary evaporation at 35 °C to afford **S61**. Yield: 2.1710 g, 5.8267 mmol, colorless oil (84%).

**<sup>1</sup>H NMR** (500 MHz, CDCl<sub>3</sub>) δ (ppm): 7.79 (d, *J* = 8.3 Hz, 2H), 7.34 (d, *J* = 8.6 Hz, 2H), 4.03 (t, *J* = 6.5 Hz, 2H), 3.55 (t, *J* = 6.3 Hz, 2H), 2.45 (s, 3H), 1.69-1.63 (m, 2H), 1.49-1.43 (m, 2H), 1.38-1.32 (m, 2H), 0.87 (s, 9H), 0.02 (s, 6H). **<sup>13</sup>C NMR** (126 MHz, CDCl<sub>3</sub>) δ (ppm): 144.77, 133.39, 129.95, 128.03, 70.72, 62.89, 32.19, 28.79, 26.09, 21.94, 21.78, 18.47, -5.18. **HRMS QToF-ESI** calculated for C<sub>18</sub>H<sub>33</sub>O<sub>4</sub>SiS [M+H<sup>+</sup>] *m/z* 373.1869; found: 373.1868.

##### 2-((5-((*tert*-butyldimethylsilyl)oxy)pentyl)oxy)ethan-1-ol (**S62**)

To a flame-dried 2-neck 25 mL round-bottom flask equipped with a magnetic stir bar under inert atmosphere (N<sub>2</sub>) was added a suspension of NaH 60% in mineral oil (85.8 mg, 2.15 mmol) and anhydrous DMF (2.20 mL, 1.46 M with respect to ethylene glycol). The solution was chilled in an ice/acetone bath for 10 minutes before ethylene glycol (200 µL, 3.22 mmol) was added dropwise by syringe (Note: gas evolution). The bath was maintained for 1 h before a solution of **S61** (417.3 mg, 1.120 mmol) in anhydrous DMF (1.0 mL, 1.2 M with respect to **S61**) was added dropwise. The bath was maintained for 1 h before gradually warming to room temperature over 23 h, at which time the reaction was judged complete by TLC (4:1 hexanes:EtOAc). The reaction was quenched with aqueous NH<sub>4</sub>Cl (3 mL), the solution was stirred for 15 minutes, and the aqueous phase was extracted with diethyl ether (8 x 4 mL). The combined organic phase was dried with magnesium sulfate, filtered through a fritted funnel using diethyl ether, and concentrated by rotary evaporation at 35 °C. Residual DMF was removed azeotropically with toluene by rotary evaporation at 50 °C to afford an oil. The oil was purified by flash column chromatography using a gradient of 10-20% EtOAc in hexanes and concentrated by rotary evaporation at 35 °C to afford **S62**. Yield: 154.7 mg, 0.5997 mmol, light-yellow oil (54%).

**<sup>1</sup>H NMR** (500 MHz, CDCl<sub>3</sub>) δ (ppm): 3.74-3.71 (m, 2H), 3.61 (t, *J* = 6.5 Hz, 2H), 3.54-3.52 (m, 2H), 3.48 (t, *J* = 6.6 Hz, 2H), 1.90 (br s, 1H), 1.57 (dq, *J* = 8.0, 6.7 Hz, 2H), 1.54 (dt, *J* = 14.8, 6.6 Hz, 2H), 1.42-1.36 (m, 2H), 0.89 (s, 9H), 0.05 (s, 6H). **<sup>13</sup>C NMR** (126 MHz, CDCl<sub>3</sub>) δ (ppm): 71.84, 71.44, 63.23, 62.05, 32.76, 29.58, 26.12, 22.53, 18.52, -5.12. **HRMS QToF-ESI** calculated for C<sub>13</sub>H<sub>30</sub>O<sub>3</sub>NaSi [M+Na<sup>+</sup>] *m/z* 285.1862; found: 285.1852.

*tert*-Butyl 2,2,3,3-tetramethyl-4,10,13-trioxa-3-silapentadecan-15-oate (**S63**)

To a flame-dried 3-neck 25 mL round-bottom flask equipped with a magnetic stir bar under inert atmosphere (N<sub>2</sub>) in an ice/acetone bath was added a suspension of NaH 60% in mineral oil (30.0 mg, 1.25 mmol). The mineral oil was removed from the NaH by the addition of portions of distilled THF (1.0 mL) as follows. The suspension was stirred for 5 minutes then allowed to stand until the solid NaH settled on the bottom of the flask. The THF and mineral oil were decanted by syringe, and the process was repeated a total of three times. To the flask was added distilled THF (1.0 mL, 0.70 M with respect to NaH). The bath was maintained for 5 minutes before the dropwise addition of a solution of **S62** (142.5 mg, 0.540 mmol) in distilled THF (1.8 mL, 0.30 M with respect to **S62**). *tert*-Butyl 2-bromoacetate (170 µL, 1.08 mmol) was added dropwise, and the bath was maintained for 20 minutes before gradually warming to room temperature over 18 h, at which time the reaction was judged complete by TLC (4:1 hexanes:EtOAc). The reaction was quenched with aqueous NH<sub>4</sub>Cl (2.5 mL), and the aqueous phase was extracted with diethyl ether (8 x 6 mL). The combined organic phase was dried with magnesium sulfate, filtered through a fritted funnel using diethyl ether, and concentrated by rotary evaporation at 35 °C. The residual DMF was removed azeotropically with toluene by rotary evaporation at 50 °C to afford an oil. The oil was purified by flash column chromatography using a gradient of 10-25% EtOAc in hexanes and concentrated by rotary evaporation at 35 °C to afford **S63**. Yield: 105.1 mg, 0.2791 mmol, colorless oil (52%).

**<sup>1</sup>H NMR** (500 MHz, CDCl<sub>3</sub>) δ (ppm): 4.03 (s, 2H), 3.71-3.69 (m, 2H), 3.62-3.59 (m, 4H), 3.46 (t, *J* = 6.8 Hz, 2H), 1.63-1.49 (m, 4H), 1.48 (s, 9H), 1.40-1.35 (m, 2H), 0.89 (s, 9H), 0.04 (s, 6H).

**<sup>13</sup>C NMR** (126 MHz, CDCl<sub>3</sub>) $\delta$ (ppm): 169.88, 81.64, 71.63, 70.91, 70.27, 69.23, 63.29, 32.82, 29.56, 28.27, 28.26, 26.13, 22.48, 18.51, -5.12. **HRMS QToF-ESI** calculated for C<sub>19</sub>H<sub>40</sub>O<sub>5</sub>NaSi [M+Na<sup>+</sup>] $m/z$ 399.2543; found: 399.2548.

*tert*-Butyl 2-(2-((5-hydroxypentyl)oxy)ethoxy)acetate (**S64**)

To a 3-dram vial equipped with a magnetic stir bar under inert atmosphere (N<sub>2</sub>) was added **S63** (436.9 mg, 1.160 mmol) and distilled THF (700 μL, 1.66 M with respect to **S63**). The contents of the flask were stirred for 5 minutes before the addition of a TBAF solution (3.4 mL, 0.29 mmol, 1 M in THF). At 5 h, the reaction was judged complete by TLC (1:1 hexanes:EtOAc). The reaction was quenched with aqueous NH<sub>4</sub>Cl (5 mL) and extracted with EtOAc (3 x 1 mL). The combined organic phase was dried with magnesium sulfate, filtered through a fritted funnel using EtOAc, and concentrated by rotary evaporation at 35 °C to afford **S64**. Crude Yield: 225.9 mg, 0.8611 mmol, colorless oil (74%). Compound **S64** was used to generate **S65** without further purification.

**<sup>1</sup>H NMR** (500 MHz, CDCl<sub>3</sub>) $\delta$ (ppm): 4.02 (s, 2H), 3.70–3.69 (m, 2H), 3.64 (t, $J$ = 6.5 Hz, 2H), 3.62–3.61 (m, 2H), 3.48 (t, $J$ = 6.6 Hz, 2H), 1.65–1.56 (m, 4H), 1.47 (s, 9H), 1.46–1.39 (m, 3H). **<sup>13</sup>C NMR** (126 MHz, CDCl<sub>3</sub>) $\delta$ (ppm): 169.86, 81.69, 71.46, 70.89, 70.27, 69.20, 62.99, 32.64, 29.42, 28.26, 22.48. **HRMS QToF-ESI** calculated for C<sub>13</sub>H<sub>26</sub>O<sub>5</sub>Na [M+Na<sup>+</sup>] $m/z$ 285.1678; found: 285.1672.

5-(2-(2-(*tert*-butoxy)-2-oxoethoxy)ethoxy)pentanoic acid (**S65**)

To a one-neck 15 mL round-bottom flask equipped with a magnetic stir bar was added **S64** (85.2 mg, 0.325 mmol), TEMPO (3.7 mg, 0.024 mmol), and ACN (450 μL, 0.70 M with respect to **S64**) and 0.67 M sodium phosphate buffer (450 μL, 1:1 v/v NaH<sub>2</sub>PO<sub>4</sub>:Na<sub>2</sub>HPO<sub>4</sub>). The contents of the flask were stirred for 10 minutes at room temperature before a solution of sodium chlorite (55.8 mg, 0.617 mmol in 300 μL deionized water) was added. The solution was warmed in an oil bath to 35 °C before 0.3 % sodium hypochlorite was added in three separate 30 μL portions at $t$ = 0 h, 2 h and 18 h. At 20 h, the reaction was judged complete by TLC (1:1 hexanes:EtOAc). The reaction was quenched with a sodium sulfite solution (244 mg in 1.5 mL water). Aqueous sodium bicarbonate was added (1.2 mL), and the aqueous phase was extracted with MTBE (1 x 2 mL) to remove TEMPO and other impurities. The pH of the aqueous phase was adjusted to 4 with aqueous 3 M HCl and was extracted with MTBE (6 x 5 mL). The pH 4 combined organic phase was dried with magnesium sulfate, filtered through a fritted funnel using MTBE, and concentrated by rotary evaporation at 35 °C to afford **S65**. Crude Yield: 118 mg, 0.427 mmol, colorless oil (94%). Compound **S65** was used to generate **S67** without further purification.

**<sup>1</sup>H NMR** (500 MHz, CDCl<sub>3</sub>) $\delta$ (ppm): 10.11 (br s, 1H), 4.02 (s, 2H), 3.71–3.69 (m, 2H), 3.62–3.60 (m, 2H), 3.50 (t, $J$ = 6.2 Hz, 2H), 2.40 (t, $J$ = 7.2 Hz, 2H), 1.76–1.70 (m, 2H), 1.69–1.63 (m, 2H), 1.47 (s, 9H). **<sup>13</sup>C NMR** (126 MHz, CDCl<sub>3</sub>) $\delta$ (ppm): 178.05, 169.98, 81.84, 71.06, 70.87, 70.30, 69.21, 33.75, 28.99, 28.28, 21.79. **HRMS QToF-ESI**: calculated for C<sub>13</sub>H<sub>24</sub>O<sub>6</sub>Na [M+Na<sup>+</sup>] $m/z$ 299.1471; found: 299.1465.

*tert-Butyl-2-(2-((5-(((S)-1-((2S,4R)-4-hydroxy-2-(((S)-1-(4-(4-methylthiazol-5-yl)phenyl)ethyl)carbamoyl)pyrrolidin-1-yl)-3,3-dimethyl-1-oxobutan-2-yl)amino)-5-oxopentyl)oxy)ethoxy)acetate (S67)*

To a flame-dried two-neck 15 mL round-bottom flask equipped with a magnetic stir bar under inert atmosphere (N<sub>2</sub>) was added **S65** (64.2 mg, 0.232 mmol), DCM (640 µL, 0.036 M with respect to **S65**), EDC•HCl (47.5 mg, 0.301 mmol), HOBt hydrate (41.9 mg, 0.301 mmol), a solution of **S66** (115 mg, 0.258 mmol) in DCM (640 µL, 0.40 with respect to **S65**) and anhydrous DMF (150 µL, 1.72 M with respect to **S65**). The solution was chilled in an ice/acetone bath for 5 minutes before the addition of DIPEA by syringe (100 µL, 0.574 mmol). The cold bath was maintained for 1 h before gradually warming to room temperature over 24 h, at which time the reaction was judged complete by TLC (5% MeOH in DCM). The reaction was quenched with deionized water (0.8 mL), and the aqueous phase extracted with EtOAc (4 x 1 mL). The combined organic phase was washed with an aqueous solution of sodium bicarbonate (1 x 1 mL), dried with sodium sulfate, filtered through a fritted funnel using EtOAc, and concentrated by rotary evaporation at 35 °C to afford a solid. The material was purified by flash column chromatography using 5% MeOH in DCM and concentrated by rotary evaporation at 35 °C to afford **S67**. Yield: 139 mg 0.198 mmol, off-white solid (85%).

**<sup>1</sup>H NMR** (500 MHz, CDCl<sub>3</sub>, major and minor rotamer) δ (ppm): 8.63 (s, 1H), 7.48 (d, *J* = 7.9 Hz, 1H), 7.34 (d, *J* = 8.4 Hz, 2H), 7.31 (d, *J* = 8.4 Hz, 2H), 6.44 (d, *J* = 8.8 Hz, 1H), 5.03 (p, *J* = 7.0 Hz, 1H), 4.63 (t, *J* = 7.8 Hz, 1H), 4.54 (d, *J* = 8.8 Hz, 1H), 4.44 (br s, 1H), 4.19 (br s, 1H), 3.96-3.94 (m, 3H), 3.62 (dd, *J* = 5.8, 3.6 Hz, 2H), 3.59 (dd, *J* = 11.1, 4.0 Hz, 1H), 3.54 (dd, *J* = 5.8, 3.6 Hz, 2H), 3.41 (td, *J* = 6.3, 1.4 Hz, 2H), 2.46 (s, 3H), 2.35 (ddd, *J* = 12.7, 7.6, 4.7 Hz, 1H), 2.15 (hept, *J* = 7.3 Hz, 2H), 2.02-1.97 (m, 1H), 1.63-1.58 (m, 2H), 1.56-1.51 (m, 2H), 1.42 (d, *J* = 6.7 Hz, 3H), 1.41 (s, 9H), 0.98 (s, 9H). **<sup>13</sup>C NMR** (126 MHz, CDCl<sub>3</sub>, major and minor rotamer) δ (ppm): 173.37, 171.84, 169.99, 169.66, 150.40, 148.36, 143.27, 131.58, 130.76, 129.48, 126.45, 81.55, 70.93, 70.64, 69.94, 69.80, 68.93, 58.69, 57.46, 56.67, 48.73, 35.95, 35.85, 35.31, 28.80, 28.08, 26.51, 22.51, 22.16, 16.03. **HRMS QToF-ESI** calculated for C<sub>36</sub>H<sub>55</sub>N<sub>4</sub>O<sub>8</sub>S [M+H<sup>+</sup>] *m/z* 703.3741; found: 703.3746.

*2-(2-((5-(((S)-1-((2S,4R)-4-hydroxy-2-(((S)-1-(4-(4-methylthiazol-5-yl)phenyl)ethyl)carbamoyl)pyrrolidin-1-yl)-3,3-dimethyl-1-oxobutan-2-yl)amino)-5-oxopentyl)oxy)ethoxy)acetic acid (S68)*

To a one-neck round-bottom flask equipped with a magnetic stir bar in an ice bath was added **S67** (138.9 mg, 0.1976 mmol) and TFA:DCM (1:3 v/v, 16.4 mL, 0.01 M with respect to **S67**). The cold bath was maintained for 1.5 h, at which time the reaction was judged complete by TLC (5% MeOH in DCM). The volatile residues were removed by rotary evaporation at 33 °C to afford an oil, and the oil was dried overnight under reduced pressure to afford the corresponding acid **S68** (TFA salt). Crude Yield: 145.2 mg, 0.1911 mmol, light brown oil (97%). Compound **S68** was used to generate **30** without further purification.

(2*S*,4*R*)-1-((*S*)-16-(*tert*-butyl)-1-((4-((4-cyanophenyl)amino)-6-((4-((*E*)-2-cyanovinyl)-2,6-dimethylphenyl)amino)-1,3,5-triazin-2-yl)amino)-4,14-dioxo-6,9-dioxo-3,15-diazaheptadecan-17-oyl)-4-hydroxy-*N*-((*S*)-1-(4-(4-methylthiazol-5-yl)phenyl)ethyl)pyrrolidine-2-carboxamide (**30**)

To a flame-dried 10 mL round-bottom flask equipped with a magnetic stir bar under inert atmosphere (N<sub>2</sub>) in an ice/acetone bath was added **5** (93.4 mg, 0.220 mmol), EDC•HCl (41.4 mg, 0.255 mmol), HOBt hydrate (35.2 mg, 0.255 mmol), and anhydrous DMF (200 µL, 1.1 M with respect to **5**). A solution of **S68** (129.4 mg, 0.2000 mmol) and DIPEA (60 µL, 0.34 mmol) was added to the reaction mixture dropwise, maintaining the cold bath temperature. The cold bath was maintained for an additional 30 minutes before the solution was gradually warmed to room temperature over 15 h, at which time the reaction was judged complete by TLC (5% MeOH in DCM). The reaction was quenched with deionized water (1 mL), and the aqueous phase was extracted with EtOAc (4 x 1 mL). The combined organic phase was washed with aqueous sodium bicarbonate (1 x 1 mL), dried with sodium sulfate, filtered through a fritted funnel using EtOAc, and concentrated by rotary evaporation at 35 °C. The material was purified by flash column chromatography using a gradient of 4-10% MeOH in DCM and concentrated by rotary evaporation at 35 °C to afford **30**. Yield: 156.5 mg, 0.1489 mmol, off-white solid (76% over two steps). *R*<sub>f</sub>: 0.42 (5% MeOH in DCM)

**<sup>1</sup>H NMR** (500 MHz, DMSO-d<sub>6</sub>, major and minor rotamer, OH signal not evident by <sup>1</sup>H NMR) δ (ppm): 9.59-9.41 (m, 1H), 8.98 (s, 1H), 8.63-8.55 (m, 1H), 8.36 (d, *J* = 7.8 Hz, 1H), 8.05-7.97 (m, 1H), 7.85-7.67 (m, 4H), 7.61-7.53 (m, 2H), 7.48-7.37 (m, 7H), 7.14 (br s, 1H), 6.41 (d, *J* = 16.5 Hz, 1H), 5.09 (d, *J* = 3.6 Hz, 1H), 4.91 (p, *J* = 7.5 Hz, 1H), 4.51 (d, *J* = 9.3 Hz, 1H), 4.42 (t, *J* = 8.0 Hz, 1H), 4.28 (br s, 1H), 3.90-3.85 (m, 2H), 3.60-3.46 (m, 7H), 2.45 (s, 3H), 2.27-2.22 (m, 1H), 2.18 (s, 6H), 2.13-2.08 (m, 1H), 2.03-1.98 (m, 1H), 1.79 (ddd, *J* = 12.9, 8.5, 4.6 Hz, 1H), 1.49 (m, 5H), 1.37 (d, *J* = 6.9 Hz, 3H), 1.24 (m, 1H), 0.92 (s, 10H). **<sup>13</sup>C NMR** (126 MHz, DMSO-d<sub>6</sub>, major and minor rotamer) δ (ppm): 171.92, 170.60, 169.59, 166.04, 165.80, 164.20, 163.99, 151.48, 150.43, 147.75, 145.13, 144.65, 139.19, 136.75, 132.48, 131.56, 131.11, 129.69, 128.85, 128.82, 127.14, 126.37, 119.57, 119.05, 118.85, 102.33, 102.22, 95.91, 70.19, 70.04, 70.02, 69.20, 68.75, 58.53, 56.36, 56.26, 47.69, 39.94 (overlapping with deuterated solvent signal), 37.73, 35.19, 34.59, 28.73, 26.43, 22.43, 22.14, 28.39, 18.35, 15.98. **HRMS QToF-ESI**: calculated for C<sub>55</sub>H<sub>68</sub>N<sub>13</sub>O<sub>7</sub>S [M+H<sup>+</sup>] *m/z* 1054.5085; found *m/z* 1054.5073.

**Scheme S21: Preparation of Compound 31**

##### *2-(2-((5-aminopentyl)oxy)ethoxy)acetic acid (S69)*

To a 1-neck 25 mL round-bottom flask equipped with a magnetic stir bar was added a solution of **S51** (206.9 mg, 0.7100 mmol), TEMPO (8.0 mg, 0.050 mmol), and a 1:1 v/v solution of ACN (1.0 mL, 0.70 M with respect to **S51**) and 0.67 M sodium phosphate buffer (1.0 mL, 1:1 v/v $\text{NaH}_2\text{PO}_4\text{:Na}_2\text{HPO}_4$, 0.70 M with respect to **S51**). The contents of the flask were stirred for 10 minutes at room temperature before a solution of sodium chlorite (128.5 mg, 1.421 mmol in 800 $\mu\text{L}$ deionized water) was added. The solution was heated to 35 °C before 0.3% sodium hypochlorite solution was added in two separate 30 $\mu\text{L}$ portions at $t = 0$ and 4 h. At 21 h, the reaction was judged complete by TLC (1:1 hexanes:EtOAc). The reaction was quenched with a sodium sulfate solution (412.0 mg in 2.0 mL deionized water), then aqueous sodium bicarbonate (0.90 mL) was added and the pH was adjusted to 9 using cold aqueous concentrated NaOH. The aqueous phase was extracted with MTBE (1 x 2 mL) to remove TEMPO and other impurities. The pH of the aqueous phase was adjusted to 4 with aqueous 1 M HCl and extracted with MTBE (5 x 3 mL). The pH 4 combined organic phase was dried with magnesium sulfate, filtered through a fritted funnel using MTBE, and concentrated by rotary evaporation at 35 °C to afford **S69**. Crude Yield: 130.1 mg, 0.4260 mmol, colorless oil (60%). Compound **S69** was used to generate **S70** without further purification.

**$^1\text{H}$ NMR** (500 MHz, $\text{CDCl}_3$) $\delta$ (ppm): 7.94 (br s, 1H), 4.14 (s, 2H), 3.72-3.70 (m, 2H), 3.60-3.58 (m, 2H), 3.46 (t, $J = 6.5$ Hz, 2H), 3.10-3.02 (m, 2H), 1.58 (p, $J = 6.8$ Hz, 2H), 1.49-1.31 (m, 14H). **$^{13}\text{C}$ NMR** (126 MHz, $\text{CDCl}_3$) $\delta$ (ppm): 173.23, 156.29, 79.35, 71.32, 71.26, 70.02, 68.75, 40.51, 29.80, 29.11, 28.48, 23.28. **HRMS QToF-ESI** calculated for $\text{C}_{14}\text{H}_{27}\text{NO}_6\text{Na}$ [$\text{M}+\text{Na}^+$] $m/z$ 328.1736; found: 328.1724.

*tert-Butyl (E)-5-(2-(2-((2-((4-((4-cyanophenyl)amino)-6-((4-(2-cyanovinyl)-2,6-dimethylphenyl)amino)-1,3,5-triazin-2-yl)amino)ethyl)amino)-2-oxoethoxy)ethoxy)pentyl)carbamate (S70)*

To a flame-dried 25 mL round-bottom flask equipped with a magnetic stir bar under inert atmosphere (N<sub>2</sub>) was added **S69** (196.0 mg, 0.6485 mmol), DCM (1.6 mL, 0.040 M with respect to **S69**), EDC•HCl (161.6 mg, 0.8431 mmol), HOBt hydrate (129.1 mg, 0.8431 mmol), and a solution of **5** (271.5 mg, 0.6380 mmol) in 5:3 v/v DCM:anhydrous DMF (800 µL, 0.80 M with respect to **5**). The solution was cooled using an ice/acetone bath for 10 minutes before DIPEA (300 µL, 1.47 mmol) was added dropwise. The cold bath was maintained for 1 h before gradually warming to room temperature over 18 h, at which time the reaction was judged complete by TLC (5% MeOH/DCM). The reaction was quenched with deionized water (2.4 mL), the aqueous phase was extracted with EtOAc (5 x 4 mL), and the combined organic phase was washed with an aqueous solution of sodium bicarbonate (1 x 8 mL). The organic phase was dried with sodium sulfate, filtered through a fritted funnel using EtOAc, and concentrated by rotary evaporation at 35 °C to afford a solid. The material was purified by Combiflash using a 5% MeOH in DCM and concentrated by rotary evaporation at 35 °C to afford **S70**. Yield: 292.3 mg, 0.4100 mmol, off-white solid (64%).

**<sup>1</sup>H NMR** (500 MHz, DMSO-d<sub>6</sub>, major and minor rotamer) δ (ppm): 9.58-9.41 (m, 1H), 8.71-8.51 (m, 1H), 8.06-7.39 (m, 9H), 7.13 (br s, 1H), 6.73 (t, *J* = 5.7 Hz, 1H), 6.41 (d, *J* = 16.5 Hz, 1H), 3.89-3.84 (m, 2H), 3.59-3.39 (m, 6H), 3.32-3.30 (m, 2H, overlapping signal of H of CH<sub>2</sub> with water solvent) 2.89-2.84 (m, 2H), 2.18 (s, 6H), 1.46-1.41 (m, 3H), 1.36 (s, 9H), 1.36-1.32 (m, 2H), 1.24-1.19 (m, 3H). **<sup>13</sup>C NMR** (126 MHz, DMSO-d<sub>6</sub>, major and minor rotamer) δ (ppm): 169.61, 166.05, 165.80, 164.20, 163.99, 155.56, 150.42, 145.13, 139.17, 136.75, 132.74, 132.49, 131.57, 127.14, 119.56, 119.04, 118.85, 102.33, 102.21, 95.92, 77.29, 70.29, 70.22, 70.19, 70.03, 69.21, 39.98 (overlapping with deuterated solvent signal), 39.74 (overlapping with deuterated solvent signal), 29.28, 28.81, 28.26, 22.91, 18.34. **HRMS QToF-ESI** calculated for C<sub>37</sub>H<sub>49</sub>N<sub>10</sub>O<sub>5</sub> [M+H<sup>+</sup>] *m/z* 713.3887; found: 713.3889.

*(E)-2-(2-((5-aminopentyl)oxy)ethoxy)-N-(2-((4-((4-cyanophenyl)amino)-6-((4-(2-cyanovinyl)-2,6-dimethyl phenyl)amino)-1,3,5-triazin-2-yl)amino)ethyl)acetamide (S71)*

To a 1-neck 25 mL round-bottom flask equipped with a magnetic stir bar in an ice bath was added **S70** (282.0 mg, 0.3956 mmol) and DCM (1.8 mL, 0.20 M with respect to **S70**). The contents of the flask were stirred for 10 minutes before TFA (1.8 mL) was added slowly by syringe. The solution was gradually warmed to room temperature over 3 h, at which time the reaction was judged complete by TLC (5% MeOH/DCM). The volatile residues were removed by rotary evaporation at 30 °C to afford an oil, which was dried overnight under reduced pressure to give the corresponding acid **S70** (TFA salt) as a light brown oil. In an Erlenmeyer flask, the oil was reconstituted in a mixture of DCM:deionized water (7:1 v/v) in an ice bath, and a concentrated aqueous NaOH solution was slowly added with rapid mixing until the pH reached 12. The aqueous phase was extracted using DCM (6 x 5 mL). The combined organic phase was dried with magnesium sulfate, filtered through a fritted funnel using DCM, and concentrated by rotary evaporation at 33 °C to afford **S71**. Crude Yield: 175.1 mg, 0.2858 mmol, white solid (72%). Compound **S71** was used to generate **31** without further purification.

**<sup>1</sup>H NMR** (500 MHz, DMSO-d<sub>6</sub>, major and minor rotamer) δ (ppm): 9.58-9.41 (m, 1H), 8.05-7.98 (m, 1H), 7.84-7.41 (m, 7H), 7.14 (br s, 1H), 6.41 (d, *J* = 16.6 Hz, 1H), 3.90-3.84 (m, 2H), 3.57-3.25 (m, 12H, overlapping signal of H from CH<sub>2</sub> with water solvent), 2.54-2.52 (m, 2H, overlapping signal of H from CH<sub>2</sub> with solvent signal), 2.88 (q, *J* = 6.7 Hz, 1H), 2.18 (s, 6H), 1.47-1.42 (m, 2H), 1.36-1.31 (m, 2H), 1.28-1.23 (m, 2H). **<sup>13</sup>C NMR** (126 MHz, DMSO-d<sub>6</sub>, major and minor rotamer) δ (ppm): 169.61, 166.04, 165.79, 164.20, 163.99, 157.09, 150.42, 145.10, 139.15, 136.74, 132.51, 131.56, 127.14, 119.57, 119.04, 118.83, 102.32, 102.21, 95.91, 70.31, 70.19, 70.03, 69.26, 40.97, 39.75 (overlapping with deuterated solvent signal), 31.72, 29.42, 28.99, 28.86, 22.93, 18.34. **HRMS QToF-ESI** calculated for C<sub>32</sub>H<sub>41</sub>N<sub>10</sub>O<sub>3</sub> [M+H<sup>+</sup>] *m/z* 613.3363; found: 613.3359.

*2-((3*r*,5*r*,7*r*)-adamantan-1-yl)-N-(5-(2-(2-((2-((4-(4-cyanophenyl)amino)-6-((4-((*E*)-2-cyanovinyl)-2,6-dimethylphenyl)amino)-1,3,5-triazin-2-yl)amino)ethyl)amino)-2-oxoethoxy)ethoxy)pentyl)acetamide (**31**)*

To a flame-dried 15 mL round-bottom flask equipped with a magnetic stir bar under inert atmosphere (N<sub>2</sub>) was added **S71** (53.0 mg, 0.0860 mmol), 1-adamantylacetic acid (18.3 mg, 0.0941 mmol), EDC•HCl (17.8 mg, 0.111 mmol), HOBt hydrate (15.7 mg, 0.111 mmol), and 9:1 v/v DCM:anhydrous DMF (510 μL, 0.17 M with respect to **S71**). The solution was cooled using an ice/acetone bath for 10 minutes before DIPEA (52 μL, 0.30 mmol) was added. The temperature of the bath was maintained for 30 minutes before gradually warming to room temperature over 19 hours, at which time the reaction was judged complete by TLC (5% MeOH/DCM). The reaction was quenched with deionized water (0.50 mL), the aqueous phase was extracted with EtOAc (4 x 1 mL), and the combined organic phase was washed with an aqueous solution of sodium bicarbonate (1 x 3 mL). The organic phase was dried with sodium sulfate, filtered through a fritted funnel using EtOAc, and concentrated by rotary evaporation at 35 °C to afford a solid. The material was purified by flash column chromatography using a gradient of 4-5% MeOH in DCM, and the fractions were concentrated by rotary evaporation at 35 °C to afford **31**. Yield: 41.7 mg, 0.0529 mmol, off-white solid (13% over two steps). **R<sub>f</sub>**: 0.55 (5% MeOH in DCM)

**<sup>1</sup>H NMR** (500 MHz, DMSO-d<sub>6</sub>, major and minor rotamer) δ (ppm): 9.59-9.41 (m, 1H), 8.71 (br s, 1H), 8.56-8.50 (m, 1H), 8.05-7.98 (m, 1H), 7.84-7.41 (m, 9H), 7.14 (br s, 1H), 6.41 (d, *J* = 16.5 Hz, 1H), 3.89-3.84 (m, 2H), 3.57-3.45 (m, 6H), 2.97 (q, *J* = 6.5 Hz, 2H), 2.18 (s, 6H), 1.88 (br s, 3H), 1.78 (s, 2H), 1.64-1.62 (m, 4H), 1.56-1.53 (m, 9H), 1.44 (t, *J* = 7.1 Hz, 3H), 1.38-1.32 (m, 3H), 1.27-1.22 (m, 3H). **<sup>13</sup>C NMR** (126 MHz, DMSO-d<sub>6</sub>, major and minor rotamer) δ (ppm): 169.66, 166.05, 165.80, 164.20, 164.00, 150.42, 145.12, 139.20, 136.74, 132.60, 131.57, 127.14, 119.56, 119.04, 118.84, 102.32, 102.21, 95.91, 79.17, 70.33, 70.18, 70.04, 69.23, 50.08, 42.12, 38.14, 39.99 (overlapping with deuterated solvent signal), 36.46, 32.12, 29.01, 28.80, 28.03, 23.10, 18.34. **HRMS QToF-ESI**: calculated for C<sub>44</sub>H<sub>57</sub>N<sub>10</sub>O<sub>4</sub> [M+H<sup>+</sup>] *m/z* 789.4564; found *m/z* 789.4554.

### **SECTION 2: Computational Design of Targeted Protein Degraders and Hydrophobic Tags**

#### **2.1 Overall Computational Design and Workflow**

The overall computational design workflow is summarized in manuscript **Figure 3** and described in the main Materials and Methods. Detailed computational protocols, software settings, and analysis procedures are provided in the Sections **2.2–2.9** below.

#### **2.2 Linker Enumeration and Derivative Generation**

The 232-member library described in the main text comprised RPV' triazine analogs bearing linker attachment at the 6-position and varied connector and linker compositions. Seventeen control compounds from previously published work<sup>12</sup> were included for benchmarking.

Linker variants were prepared in Schrodinger LigPrep with the following parameters: OPLS4 force field, maximum ligand size of 500 atoms, ionization states generated at pH 7.40 ± 2.00 using Epik<sup>13,15</sup>, (Schrodinger Release 2020-1 OR 2024-4), desalting enabled, tautomer generation enabled, stereochemistry set to retain specified chirality while allowing variation at unspecified centers, and a maximum of 32 conformers per ligand. Six connector chemotypes (amide, amine, sulfonamide, triazole, urea, polyether-only) that demonstrated favorable docking scores and predicted solubility<sup>16</sup> (QPlogS > -5) were used to compile a virtual library of 35 linker variants using polyethylene glycol (PEG) chains of varying lengths (one to six PEG units) (**Figure S1**).

#### **2.3 Constrained Molecular Docking into HIV-1 Reverse Transcriptase**

The above structures were docked using Schrodinger Glide SP (standard precision) with a core constrained protocol in which the warhead orientation was fixed in its crystallographic or consensus pose while the appended linker was allowed to sample conformational space (Schrodinger Release 2020-1 OR 2024-4)<sup>15</sup>. This constraint reduced pose degeneracy across the linker series and enabled direct comparison of docking scores across the different linker assemblies. The receptor grid was centered on the co-crystallized rilpivirine in PDB 2ZD1<sup>17</sup> and expanded to 36 Å to accommodate larger linker constructs and ensure access to the RT inter subunit channel. Core constraints were applied using maximum common substructure matching, restricted to the reference position with an RMSD tolerance of 0.10 Å. Additional docking parameters included flexible ligand sampling with nitrogen inversion and ring conformation sampling enabled, torsion bias sampling for all predefined functional groups, Epik state penalties added to docking scores, van der Waals radii scaling factor of 0.80 with partial charge cutoff of 0.15, and exclusion of ligands exceeding 500 atoms or 100 rotatable bonds. Poses were visually inspected to confirm satisfaction of the constraint, preservation of key interactions, and productive linker threading toward the inter-subunit channel. Docking scores were used as the primary ranking metric.

#### **2.4 Physicochemical Property Prediction and Triage**

For each LigPrep output, Schrodinger QikProp was run to extract two descriptors: QPlogPo/w (predicted octanol/water partition coefficient; QPlogP) and QPlogS (predicted aqueous solubility)<sup>16</sup>. Synthesis decisions were based on the triad of docking score, QPlogP, and QPlogS, along with synthetic accessibility. We prioritized designs that paired strong, constraint-consistent docking with moderate QPlogP (to avoid excessive hydrophobicity) and less-negative QPlogS (to avoid extreme predicted insolubility). No additional property triage (e.g., PSA/HBD/HBA cutoffs or external percentile benchmarks) was applied to this linker set. Amine and sulfonamide linkers were excluded from synthesis due to poor docking scores. Docking scores, QPlogP, QPlogS, and synthetic accessibility were used to prioritize 35 linker candidates, of which 12 were selected for synthesis, as summarized in manuscript **Figure 3** and **Table S1**.

### 2.5 Assembly and Enumeration of Full Bi-functional TPDs

Using ChemAxon Marvin (ChemAxon. *MarvinSketch*, versions 20.1, 21.1, and Marvin JS web application, v2025. Budapest, Hungary; 2020, 2021, and 2025), TPD building blocks were sketched and 120 TPDs were enumerated by combining optimized connector-linker chemotypes and lengths, and Pom and VH032 E3 recruiting ligands. 3D conformations were generated using Schrödinger LigPrep (OPLS4 force field, maximum 32 conformers per ligand), and protonation states were generated with Epik at pH 7.4 ± 1.0 with desalting enabled and tautomer generation disabled to retain only the most probable physiological state. For each TPD, Epik states were collapsed to a single representative "best" state by ranking first on total formal charge (preferring neutrality), second on Epik state penalty (lower is better), and third on population estimate (higher is better). The resulting best-state set was evaluated with QikProp using default settings, retaining the following properties: molecular weight (MW), topological polar surface area (PSA), hydrogen-bond donor count (donorHB), hydrogen-bond acceptor count (accptHB), rotatable bond count (#rotor), QPlogPo/w, and QPlogS. An internal database was maintained with global compound identifiers and all computed properties.

To place TPD properties on modality-appropriate scales, empirical anchors were derived from PROTAC-DB<sup>18</sup> by computing the 10th, 50th, and 90th percentiles (p10, p50, p90) across MW, PSA, donorHB, accptHB, and #rotor. Visualization used a desirability mapping in which values ≤ p50 were treated as fully favorable and values ≥ p90 as least favorable, with linear interpolation between (**Table S2**).

### 2.6 Protein–Protein Docking for Ternary Complex Modeling

Crystal structures of HIV-1 reverse transcriptase (RT) and the E3 ligase cereblon (CRBN) were downloaded from the Protein Data Bank (RT: PDB 2ZD1, chains A/B, co-crystallized with rilpivirine; CRBN: PDB 4CI3, chain B, co-crystallized with pomalidomide; VHL: PDB 4W9H, chain C, co-crystallized with VH032). Structures were prepared in Schrödinger Maestro (Release 2021-1 OR 2024-4) using the Protein Preparation Wizard with the OPLS4 force field. Bond assignment, protonation state optimization, and the addition of missing hydrogens were carried out at pH 7.4 followed by restrained minimization. RT chains were merged, with chain B (p51) residues renumbered +1000 for downstream use. For CRBN, the zinc ion was removed because its inclusion produced an error during HADDOCK processing.

Initial interface predictions were generated using CPORT<sup>19</sup>, integrating cons-PPISP<sup>20</sup> and SPPIDER<sup>21</sup>, however, this approach did not produce complexes with reciprocally oriented ligand-binding pockets. Interface residues were therefore defined semi-manually using PyMOL. For each protein, seed residues proximal to the bound ligand and exposed to solvent were selected, expanded by 5 Å, and intersected with surface-exposed residues (10 Å cutoff). This expand–

intersect procedure was repeated once, yielding 90 surface residues for RT and 95 surface residues for CRBN, each placed proximal to their respective ligand binding pockets.

Protein-protein docking was performed on the HADDOCK 2.4<sup>22</sup> web server following the standard three-stage workflow: (i) rigid-body docking (it0) using interface restraints, producing 1,000 models; (ii) semi-flexible refinement (it1) of the top 200 solutions with side-chain and limited backbone flexibility under simulated annealing; and (iii) explicit-solvent refinement with a TIP3P water shell and final minimization. Clusters were formed by fraction of common contacts and ranked by the standard HADDOCK scoring function.

Clusters were manually inspected to ensure reciprocal orientation of the RT NNRTI pocket and the CRBN thalidomide-binding pocket. The top-quality instance from Cluster 6 (model 1) was selected as the best docking solution. The TPD was constructed by cloning the co-crystallized ligands (rilpivirine from 2ZD1 and pomalidomide from 4CI3) into the respective binding sites of the docked protein complex, followed by manual assembly of the linker along the two exit vectors. The resulting ternary complex was energy-minimized in Schrödinger Prime to remove residual strain and clashes.

P4ward<sup>23</sup> was used for post-hoc PROTAC modeling starting from pre-prepared receptor and E3 ligase structures generated in Schrödinger Maestro (Protein Preparation Wizard); accordingly, P4ward's internal protein preparation steps (e.g., PDBFixer/OpenMM-based fixing/minimization) were disabled. The Maestro-prepared receptor and ligase PDBs, along with their corresponding bound ligand structures, were provided as inputs, and candidate PROTAC designs were supplied as a SMILES list for RDKit-based sanitization and conformer handling. P4ward then generated an ensemble of receptor-ligase orientations using MEGADOCK, followed by post-hoc filtering to retain geometries compatible with PROTAC formation (e.g., ligand-ligand proximity and steric compatibility/CRL clash criteria and E3-context lysine accessibility screening). The top-ranked filtered poses were then passed to P4ward's RDKit-based PROTAC sampling module, which attempts to generate linker conformations compatible with the fixed receptor- and ligase-bound ligand geometries (within a defined RMSD tolerance), discarding poses that fail sampling and optionally filtering by RDKit internal energy and/or rescoring with RXdock for prioritization.

### 2.7 Molecular Dynamics Simulations of RT-Degrader-E3 Complexes

Explicit-solvent molecular dynamics simulations were performed using Desmond (via Schrodinger Maestro). Systems were built using the System Builder with the following parameters: TIP4P water model, orthorhombic box of 10 Å length in each dimension, OPLS4 force field, neutralization with Na<sup>+</sup> ions, and 0.15 M NaCl<sup>14,24</sup>. Simulations were run in the NPT ensemble at 300 K and 1.01325 bar using the default Desmond relaxation protocol prior to production. Production trajectories were recorded at 150 ps intervals (~1,000 frames per trajectory) for 150-250 ns. Binary complexes (RT-E3 without TPD) were simulated as controls to evaluate ternary versus binary stability.

Five RT-E3 ternary complexes were simulated: N-C2-Am-PEG3 (**20**), N-C2-Am-PEG4 (**21**), N-C2-Am-PEG5 (**22**), N-C2-Am-PEG2PM5 (**28**), and O-C1-Am-Tri-PEG5 (**32**), assembled from RT (Chain A; PDB 2ZD1) and E3 (Chain B; PDB 4CI3). Per-trajectory CSVs were exported from the

Maestro Trajectory Analysis panel, capturing degrader heavy-atom RMSD (TPD RMSD), Cα RMSD for RT and E3 (RT Cα RMSD, E3 Cα RMSD), the complex radius of gyration across all protein residues (RT-E3 R<sub>gyr</sub>), and degrader solvent-accessible surface area (TPD SASA). RMSD calculations used frame superimposition with reference frame = 1 (**Figure S3**).

### 2.8 Hydrophobic-Tag Modeling and Molecular Dynamics Assessment

Four adamantyl-tagged HyT ligands – PEG3-Ad (theoretical analog only), PEG4-Ad (**26**), PEG5-Ad (**27**), and PEG2PM5-Ad (**31**) – were prepared using Schrodinger LigPrep with the same parameters as linker preparation (OPLS4 force field, pH 7.40 ± 2.00 using Epik, desalting and tautomer generation enabled, maximum 32 conformers per ligand). Physicochemical properties (MW, PSA, donorHB, accptHB, #rotor, QPlogPo/w, QPlogS) were computed using Schrodinger QikProp. (**Table S3**). Each construct was docked into RT and subjected to 50 ns all-atom explicit-water molecular dynamics using Desmond 2024-4 (TIP4P, orthorhombic box with 10 Å buffer, OPLS4, 0.15 M NaCl, NPT at 300 K). Per-trajectory CSVs were exported from the Maestro Trajectory Analysis panel, capturing HyT (excluding the RPV warhead) heavy-atom RMSD, SASA, and radius of gyration (R<sub>g</sub>). Analyses were restricted to an equilibrated 20–50 ns window.

### 2.9 Comparative Stability and Exposure Analysis

**Ternary Complex Analysis.** All ternary complex molecular dynamics simulation analyses used a uniform 50–150 ns window (**Figure S3**). To reduce short-lag autocorrelation and place metrics on a common timescale, each trajectory was partitioned into non-overlapping 5-ns blocks ([50–55), ..., [145–150) ns), and arithmetic block means were computed, yielding $n = 20$ block means per metric per complex. Between-complex differences within each metric were assessed by Kruskal-Wallis, followed (when significant) by pairwise Wilcoxon rank-sum tests with Benjamini-Hochberg FDR control (q-values reported). Compact-letter displays (CLD) summarize pairwise outcomes (complexes sharing a letter are not different at 5% FDR).

For conformational clustering, frames within the 50–150 ns analysis window were first aligned on the protein backbone (Cα of RT and E3) and clustered by heavy-atom RMSD of the degrader plus protein interface residues (side chains within 6 Å of the degrader in the reference). Trajectories were thinned to ~1–2 ns/frame to limit short-time correlation. Clustering was performed in Schrödinger Maestro using the Desmond MD analysis tools based on RMSD and agglomerative average-linkage; the number of clusters was chosen by the elbow in the within-cluster sum of squares and confirmed by silhouette score. For each cluster, the medoid (lowest average RMSD to all members) was extracted as the representative pose and cluster populations were reported. As a robustness check, clustering was repeated using k-means on the same aligned coordinates, yielding concordant dominant poses and populations.

**HyT Analysis.** Conformational sampling was quantified using heavy-atom RMSD of the linker+adamantane fragment (warhead excluded), measured relative to frame 1 without structural realignment to capture scalar excursion from the starting pose. RMSD time-series were clustered in R using DBSCAN in one-dimensional value space with $\epsilon$ in the 1.6–2.0 Å range (tuned per system via density-valley heuristics) and minPts = 12; a 3-frame rolling median was applied to suppress high-frequency flicker. For each cluster, the medoid was defined as the frame whose RMSD is closest to the cluster mean. Medoid frame indices were exported from Maestro to capture representative structures. As a robustness check, modest parameter variations ($\epsilon \pm 0.2$ Å; minPts 10–20) did not change qualitative conclusions. Time series were visualized with a short rolling mean; distributions over 20–50 ns were summarized with violin and box-and-whisker statistics using identical axes across systems. To examine coordinated changes in compactness and exposure, two-dimensional density maps of Rg versus SASA were generated from 1,000 uniformly subsampled frames per system.

**Visualization.** Figures were generated in R (tidyverse, zoo, patchwork, multcompView).

### Tables & Figures

**Table S1: Predicted physicochemical properties and docking scores of connector chemotypes, and measured IC50 values for Compounds 8- 19 (or: 8-15 and 18-19).**

| Comp_ID | Codename | Comp_No. | RT Assay<br>IC50 (μM) | TZMbl<br>IC50 (nM) | #rotor | Plog | PlogS | accptHB | donorHB | mol W | docking score |
| --- | --- | --- | --- | --- | --- | --- | --- | --- | --- | --- | --- |
| D_01 | RPV-O-C2-Am-PEG3-CH3 |  |  |  | 20 | 2.996 | -7.102 | 14.6 | 3 | 586.649 | -13.502 |
| D_02 | RPV-O-C2-Am-PEG4-CH3 |  |  |  | 23 | 2.853 | -6.085 | 16.3 | 3 | 630.702 | -12.845 |
| D_03 | RPV-O-C2-Am-PEG5-CH3 |  |  |  | 26 | 3.402 | -7.965 | 18 | 3 | 674.755 | -13.534 |
| D_04 | RPV-O-C2-Am-PEG6-CH3 |  |  |  | 29 | 3.457 | -8.2 | 19.7 | 3 | 718.808 | -9.976 |
| D_05 | RPV-O-C2-Am-PEG4-N3 | 8 | 5.84 | 2.603 | 23 | 1.374 | -5.067 | 17.6 | 3 | 641.688 | -14.087 |
| D_06 | RPV-O-C3-Am-PEG4-N3 | 9 | 16.82 | 2.709 | 24 | 2.055 | -5.974 | 17.6 | 3 | 655.715 | -13.507 |
| D_07 | RPV-N-C2-Am-PEG4-N3 | 10 | 1.75 | 2.629 | 23 | 1.44 | -6.995 | 18.1 | 4 | 640.703 | -14.285 |
| D_08 | RPV-N-C3-Am-PEG4-N3 | 11 | 7.11 | 3.019 | 24 | 1.661 | -7.226 | 18.1 | 4 | 654.73 | -13.053 |
| D_09 | RPV-O-C2-Ur-PEG3-CH3 |  |  |  | 21 | 3.02 | -6.223 | 14.1 | 4 | 615.691 | -12.97 |
| D_10 | RPV-O-C2-Ur-PEG4-CH3 |  |  |  | 24 | 3.23 | -6.3 | 15.8 | 4 | 659.744 | -13.897 |
| D_11 | RPV-O-C2-Ur-PEG5-CH3 |  |  |  | 27 | 3.947 | -9.143 | 17.5 | 4 | 703.797 | -14.925 |
| D_12 | RPV-O-C2-Ur-PEG6-CH3 |  |  |  | 30 | 4.284 | -9.866 | 19.2 | 4 | 747.85 | -13.483 |
| D_13 | RPV-O-C2-Ur-PEG4-OH | 12 | 4.72 | 2.269 | 24 | 2.46 | -5.819 | 15.8 | 5 | 645.717 | -13.199 |
| D_14 | RPV-O-C3-Ur-PEG4-OH | 13 | 4.36 | 2.486 | 25 | 2.05 | -4.337 | 15.8 | 5 | 659.744 | -13.287 |
| D_15 | RPV-N-C2-Ur-PEG4-OH | 14 | 10.11 | 1.618 | 24 | 2.19 | -7.557 | 16.3 | 6 | 644.732 | -14.625 |
| D_16 | RPV-N-C3-Ur-PEG4-OH | 15 | 23.98 | 3.11 | 25 | 2.367 | -5.138 | 16.3 | 6 | 658.759 | -14.01 |
| D_17 | RPV-O-C1-Tri-PEG3-CH3 |  |  |  | 20 | 4.106 | -8.696 | 14.6 | 2 | 610.674 | -13.73 |
| D_18 | RPV-O-C1-Tri-PEG4-CH3 |  |  |  | 23 | 4.08 | -8.626 | 16.3 | 2 | 654.727 | -13.424 |
| D_19 | RPV-O-C1-Tri-PEG5-CH3 |  |  |  | 26 | 4.749 | -9.79 | 18 | 2 | 698.78 | -14.202 |
| D_21 | RPV-O-C1-Tri-PEG3-OH | 16 | >10 | >50 | 20 | 3.308 | -8.434 | 14.6 | 3 | 596.647 | -13.905 |
| D_22 | RPV-O-C1-Tri-PEG4-OH | 17 | >10 | >50 | 23 | 3.522 | -8.887 | 16.3 | 3 | 640.7 | -13.922 |
| D_23 | RPV-O-C2-Amn-PEG3-CH3 |  |  |  | 21 | 3.673 | -7.034 | 13.6 | 3 | 572.666 | -12.858 |
| D_24 | RPV-O-C2-Amn-PEG4-CH3 |  |  |  | 24 | 3.411 | -4.954 | 15.3 | 3 | 616.719 | -9.144 |
| D_25 | RPV-O-C2-Amn-PEG5-CH3 |  |  |  | 27 | 3.455 | -4.499 | 17 | 3 | 660.772 | -13.474 |
| D_26 | RPV-O-C2-Amn-PEG6-CH3 |  |  |  | 30 | 4.181 | -7.979 | 18.7 | 3 | 704.825 | -9.162 |
| D_27 | RPV-O-C2-Sul-PEG3-CH3 |  |  |  | 21 | 2.138 | -4.481 | 16.6 | 3 | 622.698 | -12.992 |
| D_28 | RPV-O-C2-Sul-PEG4-CH3 |  |  |  | 24 | 2.955 | -7.354 | 18.3 | 3 | 666.751 | -14.122 |
| D_29 | RPV-O-C2-Sul-PEG5-CH3 |  |  |  | 27 | 2.546 | -5.384 | 20 | 3 | 710.804 | -9.799 |
| D_30 | RPV-O-C2-Sul-PEG6-CH3 |  |  |  | 30 | 3.375 | -8.443 | 21.7 | 3 | 754.857 | -9.247 |
| D_31 | RPV-N-C1-Am(inv)-PEG4-OH | 18 | 4.47 | 3.857 | 23 | 1.495 | -2.573 | 16.05 | 4.25 | 615.691 | -13.51 |
| D_32 | RPV-O-PEG3-OH |  |  |  | 18 | 3.183 | -7.523 | 12.1 | 3 | 515.571 | -12.893 |
| D_33 | RPV-O-PEG4-OH | 19 | 39.77 | 4.463 | 21 | 3.213 | -7.464 | 13.8 | 3 | 559.624 | -13.115 |
| D_34 | RPV-O-PEG5-OH |  |  |  | 24 | 3.341 | -7.05 | 15.5 | 3 | 603.677 | -13.199 |
| D_35 | RPV-O-PEG6-OH |  |  |  | 27 | 3.973 | -9.489 | 17.2 | 3 | 647.73 | -13.264 |

**Table S2: Physicochemical properties of the RPV-based synthesized PROTACs benchmarked against PROTAC-DB 3.0.** Six descriptors (MW, cLogP, TPSA, HBD, HBA, rotatable bonds) were computed from SMILES with RDKit, identically for the compounds and the reference set. Each cell is shaded by the compound's percentile within the PROTAC-DB 3.0 distribution (n = 6,153): blue = low, gray = median, red = high.

| Comp No. | Codename | RT IC50 (μM) | TZM-bl IC50 (nM) | MW | cLogP | TPSA | HBD | HBA | Rot. bonds |
| --- | --- | --- | --- | --- | --- | --- | --- | --- | --- |
| 20 | RPV'-N-C2-Am-PEG3-AnPom | 6.398 | >225 | 827 | 3.5 | 265 | 6 | 16 | 19 |
| 21 | RPV'-N-C2-Am-PEG4-AnPom | 11.17 | 23.98 | 871 | 3.5 | 275 | 6 | 17 | 22 |
| 22 | RPV'-N-C2-Am-PEG5-AnPom | – | – | 915 | 3.5 | 284 | 6 | 18 | 25 |
| 23 | RPV'-N-C2-Am-PEG3-VH032 | 32.91 | 100 | 998 | 5.2 | 282 | 7 | 17 | 22 |
| 24 | RPV'-N-C2-Am-PEG4-VH032 | >40 | 100 | 1042 | 5.2 | 291 | 7 | 18 | 25 |
| 25 | RPV'-N-C2-Am-PEG5-VH032 | – | >225 | 1086 | 5.2 | 300 | 7 | 19 | 28 |
| 28 | RPV'-N-C2-Am-PEG2PM5-AnPom | ~20 | 30.89 | 869 | 4.6 | 265 | 6 | 16 | 22 |
| 29 | RPV'-N-C2-Am-PEG3Pip-Dpom | >40 | 70.95 | 895 | 4.7 | 257 | 5 | 16 | 19 |
| 30 | RPV'-N-C2-Am-PEG2PM4-Me032 | >40 | 73.6 | 1054 | 6.9 | 282 | 7 | 17 | 25 |

**Table S3: Predicted Physicochemical properties and docking scores of HyTs.**

| Comp No. | Codename | RT IC50 (μM) | TZMBL IC50 (nM) | Docking Score | Plog | PlogS | #rotor | PSA | accptHB | donorHB | mol MW |
| --- | --- | --- | --- | --- | --- | --- | --- | --- | --- | --- | --- |
|  | RPV'-N-C2-Am-PEG3-Ad |  |  | -14.146 | 4.215 | -8.848 | 22 | 195.371 | 15.9 | 5 | 746.911 |
| 26 | RPV'-N-C2-Am-PEG4-Ad | 22.93 | 18.24 | -14.319 | 4.201 | -9.346 | 25 | 209.265 | 17.6 | 5 | 790.964 |
| 27 | RPV'-N-C2-Am-PEG5-Ad | - | - | -14.111 | 4.34 | -9.56 | 28 | 220.244 | 19.3 | 5 | 835.017 |
| 31 | RPV'-N-C2-Am-PEG2PM5-Ad | >40 | 12.96 | -14.283 | 4.534 | -8.923 | 24 | 194.568 | 15.9 | 5 | 774.964 |

**Figure S1.** Predicted aqueous solubility and docking scores of RPV-linker assemblies (connector chemotypes). Scatter plot showing predicted aqueous solubility, expressed as predicted logS, plotted against docking score for rilpivirine-containing linker assemblies. Each point represents an individual RPV-linker candidate. Circles indicate synthesized compounds, whereas triangles indicate theoretical designs. Point colors correspond to the linker chemistry class: amide-PEG, amino-PEG, PEG, sulfonamide-PEG, triazole-PEG, or urea-PEG. More negative docking scores indicate stronger predicted binding, while lower predicted logS values indicate lower predicted aqueous solubility. Selected candidates are annotated by compound number or codename to highlight representative designs considered during candidate prioritization.

**Figure S2. Protein-protein docking and MD-based selection of HIV-1 RT ternary complexes for TPD selection.** Left: Overall model of the selected ternary complex, showing HIV-1 reverse transcriptase (RT; gray surface) associated with the recruited protein partner/E3 ligase component (purple surface). Compound 21 is shown in green and bridges the two protein surfaces, supporting formation of the modeled ternary complex. Right: Close-up view of the ternary-interface region, highlighting the predicted binding pose of compound 21 and neighboring protein residues. Dashed lines indicate key polar contacts or hydrogen-bonding interactions that stabilize the modeled complex.

TPD/Backbone metrics overview (window: 50-150 ns)

**Figure S3. MD analysis – comparison of ternary complexes (50–150 ns).** Left: TPD SASA vs TPD RMSD scatter plot (one facet per complex; points are 5-ns block means). Middle: Distributions of 5-ns block means for each metric shown as violins with embedded boxplots (white box = IQR; center line = median; dots = individual block means). Grey letters at the top of each facet indicate compact-letter display (CLD) groupings from tests on block means by Kruskal-Wallis omnibus followed, when significant, by pairwise Wilcoxon rank-sum tests with Benjamini–Hochberg FDR control ($q \leq 0.05$); complexes sharing a letter are not significantly different. Right: Time series within the 50–150 ns window (light = raw per-frame values; bold = $\sim 2$-ns rolling mean); y-axes are free per metric. Colors denote the TPD complex. Metrics: RT Cα RMSD (Chain A/2ZD1), E3 Cα RMSD (Chain B/4CI3), RT–E3 Rgyr (protein-pair radius of gyration), TPD RMSD (degrader heavy atoms), and TPD SASA (degrader solvent-accessible surface area).

#### SECTION 3. Optimization of Doxycycline Induction for the Expression of p66.HiBiT

**Figure S4.** Titration of doxycycline on cells transduced with p51-p66.HiBiT lentiviral vector. Cells were induced with the indicated doxycycline concentration to express p51-p66.HiBiT for 48 hours with doxycycline replenished every 24 hours. Background RLU of uninduced cells was subtracted from each condition. The blue arrow indicates the doxycycline concentration used in the degradation assay.

### SECTION 4. Kinetic Solubility Determination by UV–Vis Spectroscopy

#### 4.1 General Considerations and Instrumentation

Kinetic solubility measurements were performed using a UV–visible supersaturation assay adapted from the method of Kerns and co-workers,<sup>1</sup> with modifications to accommodate bifunctional degraders and hydrophobic tag (HyT) molecules. All measurements were conducted at room temperature (22–25 °C) in phosphate-buffered saline (PBS, pH 7.4) containing 1% (v/v) DMSO.

UV–vis absorbance measurements were collected using a Hewlett Packard 8452A diode array spectrophotometer equipped with 1 cm path length quartz cuvettes. pH adjustments were performed using a Mettler Toledo FiveGo pH meter calibrated immediately prior to use. Sonication was carried out using a VWR ultrasonic cleaner (model 97043-964). Sample filtration employed 0.22 µm hydrophobic PTFE syringe filters.

#### 4.2 Buffer Preparation

Phosphate-buffered saline (PBS; 10 mM phosphate, 150 mM NaCl, pH 7.4) was prepared using Na<sub>2</sub>HPO<sub>4</sub>·2H<sub>2</sub>O, NaH<sub>2</sub>PO<sub>4</sub>, and NaCl in deionized water. The pH was adjusted to 7.40 ± 0.02 using 1 M HCl or NaOH and confirmed with a calibrated pH meter.

#### 4.3 Stock Solutions

Stock solutions of compounds RPV'-N-C2-PEG4-Pom (**21**), RPV'-N-C2-PEG4-VH032 (**24**), and RPV'-N-C2-PEG4-Ad (**26**) were prepared in DMSO at concentrations ranging from 0.6–19 mM via gravimetric dilution. All stock solutions were sonicated for 3 min to ensure complete dissolution prior to use.

#### 4.4 Calibration Curves

Calibration standards were prepared in PBS containing 1% (v/v) DMSO over the indicated concentration ranges. Samples were equilibrated, filtered (0.22 µm PTFE), and analyzed by UV–vis spectroscopy at the respective $\lambda_{\text{max}}$ values. Each concentration was prepared in duplicate, and the resulting average absorbance values were plotted against concentration. Molar absorptivities were determined by linear regression using Beer's Law.

• **RPV'-N-C2-PEG4-VH032 (24):** $\lambda_{\text{max}} = 294 \text{ nm}$; $\epsilon = 24,090 \text{ M}^{-1} \text{ cm}^{-1}$
• **RPV'-N-C2-PEG4-Ad (26):** $\lambda_{\text{max}} = 292 \text{ nm}$; $\epsilon = 1486 \text{ M}^{-1} \text{ cm}^{-1}$

**Figure S5.** Calculated calibration curve for **24** from 5.1-90.9 μM and an overlaid UV-vis spectra.

**Figure S6.** Calculated calibration curve for **26** from 0.8-24.3 μM and an overlaid UV-vis spectra. Concentrations exceeding the specified ranges resulted in visible precipitation and were excluded from linear regression analyses.

For **RPV'-N-C2-PEG4-Pom (21)**, reliable post-filtration absorbance could not be obtained at low micromolar concentrations due to extensive precipitation following dilution into aqueous buffer exceeding 3 mM. Accordingly, a calibration curve could not be generated under these conditions.<sup>2</sup>

##### 4.5 Kinetic Solubility Determination

Kinetic solubility was defined as the maximum concentration of compound remaining in solution following overnight equilibration of supersaturated samples under aqueous conditions.

Supersaturated solutions (2.5 mL total volume) were prepared by dilution of concentrated DMSO stock solutions into PBS containing 1% (v/v) DMSO. Samples were equilibrated with gentle stirring for 1 h, followed by static incubation for 18 h at room temperature. After equilibration, samples were filtered through 0.22 $\mu\text{m}$ PTFE syringe filters to remove undissolved material, and the dissolved compound concentration was quantified by UV-vis spectroscopy using the corresponding calibration curve. Each concentration was prepared in triplicate, and the resulting average absorbance values were used to calculate the dissolved concentration.

##### RPV'-N-C2-PEG4-VH032 (24)

Following equilibration, visible precipitation was observed in all supersaturated samples. Quantitative UV-vis analysis of the filtered solutions indicated a kinetic solubility of $101.8 \pm 0.65 \mu\text{M}$ under the conditions employed.

**Figure S7.** Solubility curve of **24** depicting the theoretical dissolved concentration of compound ($\mu\text{M}$) versus the calculated dissolved concentration ($\mu\text{M}$)

#### RPV'-N-C2-PEG4-Ad (26)

Supersaturated samples of **26** similarly exhibited precipitation after overnight equilibration. UV-vis analysis of filtered solutions indicated a kinetic solubility of $16.06 \pm 0.88 \mu\text{M}$.

**Figure S8.** Solubility curve of **26** depicting the theoretical dissolved concentration of compound ($\mu\text{M}$) versus the calculated dissolved concentration ($\mu\text{M}$)

#### RPV'-N-C2-PEG4-Pom (21)

**21** showed extensive aggregation and precipitation upon dilution into PBS at concentrations exceeding $3 \mu\text{M}$, with post-filtration absorbance values falling below the reliable detection limit of the instrument. Representative UV-vis spectra collected before and after filtration demonstrate substantial loss of soluble material, consistent with rapid precipitation.

Based on these observations, the kinetic solubility of **21** under the conditions employed was estimated to be $< 3 \mu\text{M}$.

**Figure S9.** Representative UV–vis spectra of **21** before and after filtration.

#### Calculation of Dissolved Concentration

Dissolved compound concentrations were calculated using Beer–Lambert law:

$$A = \varepsilon l c$$

$$c \text{ (}\mu\text{M)} = \frac{A}{(\varepsilon \times l)} \times 10^6$$

where $A$ is absorbance, $\varepsilon$ is molar absorptivity ($\text{M}^{-1} \text{cm}^{-1}$), $l$ is path length (1 cm), and $c$ is calculated kinetic concentration ($\mu\text{M}$).

### References:

1. Babij, N. R.; McCusker, E. O.; Whiteker, G. T.; Canturk, B.; Choy, N.; Creemer, L. C.; Amicis, C. V. D.; Hewlett, N. M.; Johnson, P. L.; Knobelsdorf, J. A.; Li, F.; Lorsbach, B. A.; Nugent, B. M.; Ryan, S. J.; Smith, M. R.; Yang, Q. NMR Chemical Shifts of Trace Impurities: Industrially Preferred Solvents Used in Process and Green Chemistry. *Org. Process Res. Dev.*, **2016**, *20*, 661-667. DOI: 10.1021/acs.oprd.5b00417.
2. Delacroix, K.; Fours, B.; Kowalkowski, S.; Revil-Baudard, V.; Perez, M.; Petit, L. Process Development of a Novel Route to Rilpivirine Hydrochloride. *Org. Process Res. Dev.* **2024**, *28* (2), 524–531. https://doi.org/10.1021/acs.oprd.3c00326
3. Dobrikov, G. M.; Nikolova, Y.; Slavchev, I.; Dangalov, M.; Deneva, V.; Antonov, L.; Vassilev, N. G. Structure and Conformational Mobility of OLED-Relevant 1,3,5-Triazine Derivatives. *Molecules* **2023**, *28* (3), 1248. https://doi.org/10.3390/molecules28031248.
4. Yue, X.; Feng, Y.; Yu, Y. B. Synthesis and Characterization of Fluorinated Conjugates of Albumin. *J. Fluor. Chem.* **2013**, *152*, 173-181. DOI: 10.1016/j.fluchem.2013.01.026
5. Lu, T.; Chen, F.; Yao, J.; Bu, Z.; Kyani, A.; Liang, A.; Liang, B.; Chen, S.; Zheng, Y.; Liang, H.; Neamati, N.; Liu, Y. Design of FK866-Based Degraders for Blocking the Nonenzymatic Functions of Nicotinamide Phosphoribosyltransferase. *J. Med. Chem.* **2024**, *67*, 10, 8099–8121. DOI: 10.1021/acs.jmedchem.4c00193
6. Girardinia, M.; Maniacia, C.; Hughesa, S. J.; Testa, A.; Ciulli, A. Cereblon versus VHL: Hijacking E3 ligases against each other using PROTACs. *Bioorg. Med. Chem.* **2019**, *27*, 12, 2466-2479. DOI: 10.1016/j.bmc.2019.02.048
7. Ericsson, E.; Enander, K.; Bui, L.; Lundström, I.; Konradsson, P.; Liedberg, B. Site-Specific and Covalent Attachment of His-Tagged Proteins by Chelation Assisted Photoimmobilization: A Strategy for Microarraying of Protein Ligands. *Langmuir*. **2013**, *29*, 37, 11687–11694. DOI: 10.1021/la4011778
8. Mainolfi, N.; Ji, N.; Kluge, A. F.; Weiss, M. M.; Zhang, Y.; Zheng, X. IRAK Degradation and Uses Thereof. US WO 2020/113233 A1, 2020.
9. Nielsen, N. Chong J. M., Burchat, A. F. Titration of alkullithiums with a simple reagent to a blue endpoint. *J. Organomet. Chem.* **1997**, *542*, 281-283. DOI: 10.1016/S0022-328X(97)00143-5
10. Costello, J. P.; Ferreira, E. M. Regioselectivity Influences in Platinum-Catalyzed Intramolecular Alkyne O-H and N-H Additions. *Org. Lett.* **2019**, *21*, 9934-9939. DOI: 10.1021/acs.orglett.9b03557.
11. Champagne, P.L.; Ester, D.; Polan, D.; Williams, V.; Thangadurai, V.; Ling, C. C. Amphiphilic Cyclodextrin-Based Liquid Crystals for Proton Conduction. *J. Am. Chem. Soc.* **2019**, *141*, 9217-9224. DOI: 10.1021/jacs.8b13888

12. Bollini, M.; Cisneros, J. A.; Spasov, K. A.; Anderson, K. S.; Jorgensen, W. L., Optimization of Diarylazines as Anti-HIV Agents with Dramatically Enhanced Solubility. *Bioorg. Med. Chem. Lett.* 2013, 23 (18), 5213–5216.
13. Shelley, J. C.; Cholleti, A.; Frye, L. L.; Greenwood, J. R.; Timlin, M. R.; Uchimaya, M., Epik: A Software Program for pK<sub>a</sub> Prediction and Protonation State Generation for Drug-Like Molecules. *J. Comput. Aided Mol. Des.* 2007, 21 (12), 681–691.
14. Lu, C.; Wu, C.; Ghoreishi, D.; Chen, W.; Wang, L.; Damm, W.; Ross, G. A.; Dahlgren, M. K.; Russell, E.; Von Bargen, C. D.; Abel, R.; Friesner, R. A.; Harder, E., OPLS4: Improving Force Field Accuracy on Challenging Regimes of Chemical Space. *J. Chem. Theory Comput.* 2021, 17 (7), 4291–4300.
15. Friesner, R. A.; Banks, J. L.; Murphy, R. B.; Halgren, T. A.; Klicic, J. J.; Mainz, D. T.; Repasky, M. P.; Knoll, E. H.; Shelley, M.; Perry, J. K.; Shaw, D. E.; Francis, P.; Shenkin, P. S., Glide: A New Approach for Rapid, Accurate Docking and Scoring. 1. Method and Assessment of Docking Accuracy. *J. Med. Chem.* 2004, 47 (7), 1739–1749.
16. Jorgensen, W. L.; Duffy, E. M., Prediction of Drug Solubility from Structure. *Adv. Drug Deliv. Rev.* 2002, 54 (3), 355–366.
17. Das, K.; Bauman, J. D.; Clark, A. D., Jr.; Frenkel, Y. V.; Lewi, P. J.; Shatkin, A. J.; Hughes, S. H.; Arnold, E., High-Resolution Structures of HIV-1 Reverse Transcriptase/TMC278 Complexes: Strategic Flexibility Explains Potency against Resistance Mutations. *Proc. Natl. Acad. Sci. U.S.A.* 2008, 105 (5), 1466–1471.
18. Ge, J.; Li, S.; Weng, G.; Wang, H.; Fang, M.; Sun, H.; Deng, Y.; Hsieh, C.-Y.; Li, D.; Hou, T., PROTAC-DB 3.0: An Updated Database of PROTACs with Extended Pharmacokinetic Parameters. *Nucleic Acids Res.* 2025, 53 (D1), D1510–D1515.
19. de Vries, S. J.; Bonvin, A. M. J. J., CPORT: A Consensus Interface Predictor and Its Performance in Prediction-Driven Docking with HADDOCK. *PLoS One* 2011, 6 (3), e17695.
20. Chen, H.; Zhou, H.-X., Prediction of Interface Residues in Protein–Protein Complexes by a Consensus Neural Network Method: Test against NMR Data. *Proteins* 2005, 61 (1), 21–35.
21. Porollo, A.; Meller, J., Prediction-Based Fingerprints of Protein–Protein Interactions. *Proteins* 2007, 66 (3), 630–645.
22. Honorato, R. V.; Trellet, M. E.; Jiménez-García, B.; Schaarschmidt, J. J.; Giulini, M.; Reys, V.; Koukos, P. I.; Rodrigues, J. P. G. L. M.; Karaca, E.; van Zundert, G. C. P.; Roel-Touris, J.; van Noort, C. W.; Jandová, Z.; Melquiond, A. S. J.; Bonvin, A. M. J. J., The HADDOCK2.4 Web Server for Integrative Modeling of Biomolecular Complexes. *Nat. Protoc.* 2024, 19 (11), 3219–3241.
23. Jofily, P.; Kalyaanamoorthy, S., P4ward: An Automated Modeling Platform for Protac Ternary Complexes. *J. Chem. Inf. Model.* 2025, 65 (16), 8806–8818.
24. Bowers, K. J.; Chow, E.; Xu, H.; Dror, R. O.; Eastwood, M. P.; Gregersen, B. A.; Klepeis, J. L.; Kolossváry, I.; Moraes, M. A.; Sacerdoti, F. D.; Salmon, J. K.; Shan, Y.; Shaw, D. E.,

Scalable Algorithms for Molecular Dynamics Simulations on Commodity Clusters.
Proceedings of the 2006 ACM/IEEE Conference on Supercomputing (SC '06) 2006, 84.

25. Edward, H. K.; Li, D.; Guy, T. C., In Vitro Solubility Assays in Drug Discovery. *Curr. Drug Metab.* **2008**, 9 (9), 879-885.
26. Venturi, A.; Di Bona, S.; Desantis, J.; Eleuteri, M.; Bartalucci, M.; Baroni, M.; Benedetti, P.; Goracci, L.; Cruciani, G., Between Theory and Practice: Computational/Experimental Integrated Approaches to Understand the Solubility and Lipophilicity of PROTACs. *J. Med. Chem.* **2024**, 67 (18), 16355-16380.
